# Deriving genetic codes for molecular phenotypes from first principles

**DOI:** 10.1101/2022.08.15.503769

**Authors:** Salil S. Bhate, Anna Seigal, Juan Caicedo

## Abstract

The genetic code is a formal principle that determines which proteins an organism can produce from only its genome sequence, without mechanistic modeling. Whether similar formal principles govern the relationship between genome sequence and phenotype across scales – from molecules to cells to tissues – is unknown. Here, we show that a single formal principle – structural correspondence — underlies the relationship between phenotype and genome sequence across scales. We represent phenotypes and the genome as graphs and find mappings between them using structure preservation as the sole constraint. Combinatorial richness in phenotypes more tightly constrains which mappings preserve that structure. Thus, phenotypic structure predicts genetic associations independently of covariation with genotype. This principle rediscovers the amino acid code without prior knowledge of translation or coding sequences, using just one protein and genome sequence as input. We benchmark this principle: applied to phenotypes at the cell, tissue and organ scales, the mappings correctly predict established associations and are driven by transcription factor motifs. Applied to cancer tissue images, we find regulators of spatial gene expression in immune cells. We thus offer a first-principles framework to relate genome sequence with phenotypic structure and guide mechanistic discovery across scales.

## Introduction

Understanding how genome sequence constrains the phenotypes an organism can produce is a central question in genetics. Currently, this question is addressed through direct measurement and predictive modeling: genetic perturbation and population studies measure genotype-phenotype associations (Engreitz et al., 2024; Rood et al., 2025); sequence-to-function (Toneyan and Koo, 2024) and virtual cell models learn from this data to predict novel associations and perturbation effects (Bunne et al., 2024). However, mapping how genetic effects propagate through intermediates at cellular and tissue scales is restricted by sample availability and cost (Lappalainen and MacArthur, 2021). Thus, answering how the genome constrains phenotypes – even through scaling machine learning models – requires biological principles that provide the inductive biases to generalize across scales and systems.

At the protein scale, a principle exists that enables this generalization. The genetic code articulates which amino acid sequences can be expressed from any given genome. Beyond its fundamental biological role, this principle has a practical consequence: the genomic loci defining the sequence of any expressed protein can be found *in silico* by sequence alignment, without relying on genotype-phenotype covariation or mechanistic knowledge of the protein at hand. However, there does not exist a unifying, first-principles framework that similarly formalizes across scales, from molecules to cells to tissues, the relationship between phenotypic structures and the sequences in the genome that shape them. We propose that formally modeling the structural relationship between phenotypes and the genome sequence will enable identifying upstream genetic mechanisms and perturbation effect prediction in contexts where large-scale genotype-phenotype covariation cannot be measured.

Since organismal form and genome sequence are mutually shaped by evolutionary processes, we reasoned that comparing their formal structures might enable deciphering their connections without requiring covariation of genotype and phenotype. This intuition parallels the decipherment of Egyptian hieroglyphs from the Rosetta stone: consistent relationships between symbols across languages enabled translation from just three inscriptions. Here, we investigate whether a similar assumption of consistent structural relationships across phenotype and genome sequence is sufficient to correctly translate between them.

We consider a class of phenotypes we term compositional: those defined by a set of constituent biological units and the relationships between them. For example, proteins, tissues and organs are composed of amino acid residues, cells, and anatomical compartments as units respectively. This class of phenotypes includes single-cell and spatial atlases. Phenotypes such as height or disease state are not compositional, because they do not measure constituent biological units and relationships.

We formalize a principle – Phenotype Sequence Alignment Sufficiency (PSA-sufficiency) – that articulates how the relationships between units of a compositional phenotype can enable identification of the genomic sequences that influence those units, without requiring genotype-phenotype associations or modeling mechanisms.

PSA, without sufficiency, is the biologically plausible assumption that there is a correspondence between the similarity structure of a compositional phenotype’s units and the genomic sequences that influence them. This is expected (regardless of scale or mechanism) if similar phenotypic units arise, on average, from similar genetic programs, and similar sequences, on average, enact similar genetic programs.

Concretely, PSA is a search constraint for assigning genomic loci to compositional phenotypes. This computational task, which we term PSA-search, is to define phenotypic and sequence similarity measures and then group and match phenotypic units with genomic loci in a possibly many-to-many way that preserves these similarity measures. PSA-search is a general optimization task: the specific algorithm implementing it corresponds to a prior on the structure of the assignment and depends on the phenotype at hand.

PSA-search searches for structure-preserving assignments. It leverages phenotypic complexity, instead of covariation between sequences and measurements (from functional genomics or population measurements) or prior correspondence between phenotypic measurements and the genome (such as gene annotations), to assign genomic loci to phenotypic units. We refer to the output of PSA-search as a code: an assignment of genomic loci to phenotypic units.

The central question is whether PSA-search, when applied to a compositional phenotype, produces a code that recapitulates empirically established ground truth (e.g., population genetic associations or functional genomics). When PSA-search produces a code that recapitulates ground truth, we say that the phenotype is PSA-sufficient. PSA-sufficiency is a biological property of the phenotype: something we empirically investigate by comparing the output of a PSA-search to measured genotype-phenotype associations.

Our hypothesis is that many phenotypes, across scales, are PSA-sufficient. That is, they are sufficiently combinatorially rich that PSA is enough of a constraint – with an appropriate algorithmic parameterization of PSA-search – to recover the loci in the genome that affect a phenotype. We evaluated PSA-sufficiency of phenotypes across scales – from molecules to cells to tissues to organs – by comparing the output of PSA-search to known codes, associations from population genetics, causal perturbations and enriched motifs. We investigated the specificity of the output codes with ablations and permutation tests.

We first evaluated PSA-sufficiency in the context of protein sequences, where a ground truth code is available. Here, PSA-search rediscovers the amino acid code, *de novo*, using just one protein and one genome. Crucially, PSA-search only resolves the correct code when the input protein sequence is sufficiently combinatorially complex.

Like protein sequences, organs and single-cell distributions are combinatorially complex. We therefore investigated whether PSA-sufficiency held at these scales. We initially restricted our attention to codes in enhancers, because context-specific regulation of gene expression by enhancers is sequence-determined. We formulated PSA-search to assign enhancers in the ENCODE atlas (Moore et al., 2020) to compositional phenotypes.

First, we assigned enhancers to brain regions in the Allen brain atlas using preservation of phenotypic similarity between regions. The assignments were consistent across replicates and biologically coherent as determined by both region-specific gene expression and genome-wide association study hits. Next, we assign sets of enhancers to cells in the Tabula Sapiens single-cell atlas (Consortium* et al., 2022) using preservation of phenotypic similarity between cells. Transcription factor binding sites were correctly enriched in the enhancers assigned to different immune cell types. We found a correspondence between T cell transcriptional diversity and TCF7 motif diversity and established its evolutionary conservation, suggesting that the implementation of PSA-sufficiency is a fundamental biological property.

Given the success at finding immune cell codes while pre-specifying enhancers as loci of interest, we statistically benchmarked the ability of PSA-search to recover genetic associations without prior genomic annotations. We formulated PSA-search with neural networks to find continuous assignments of loci to single-cell gene expression in blood cells in the human genome to enable benchmarking with prediction of cell type specific expression quantitative trait loci (eQTLs).

The code we found outperformed baselines such as enhancer annotations and GC content, but did not outperform TSS proximity (which requires knowledge of where a gene is encoded) for predicting eQTLs. Permutation tests and ablations confirmed gene and cell type specificity of these predictions. Crucially, neural network attributions showed that CD8+ T cell naïve/effector diversity was also, in this annotation-free and continuous setting, encoded by TCF7 motifs. The code, as a result, causally predicts CRISPR screen hits for interferon-gamma production from TSS sequence, by editing interferon-gamma in the phenotypic data in-silico. Further investigating which in-silico phenotypic edits predict the screen hits correctly enriches for pathways downstream of interferon-gamma, highlighting how PSA-search can leverage structural correspondence between genome sequence and phenotype to make hypotheses about upstream regulators.

Since we observed the structural correspondence between T cell diversity and sequence motifs in multiple distinct contexts, we formulated PSA-search to find a code for a translationally important, higher-order phenotype in the human genome without prior annotations: CD8+ T cell neighborhoods in colorectal cancer. The code was driven by sequence patterns in enhancers corresponding to enrichment of TBX21, EOMES, STAT4, and PLAGL1 motifs (fate-determining TFs) that changed smoothly across neighborhoods. The code generated the hypotheses that (a) there is neighborhood-specific STAT4-dependent activation of the natural killer receptor KLRC3 active when CD8+ T cells are surrounded by tumor cells, and (b) that microenvironmental gene regulatory programs converge on ZBTB20. Hypothesis (a) is consistent with the STAT4 determined expression of KLRC3 in tumor-surrounded CD8+ T cells in (Fesneau et al., 2024) and hypothesis (b) is consistent with the central role of ZBTB20 in metabolic regulation of CD8+ T cell differentiation in tumors determined in (Sun et al., 2020).

PSA-sufficiency is thus a structural correspondence principle underlying the relationship between genotype and phenotype across levels of biological organization, and a framework for discovering genomic loci with emergent effects on phenotypic structure.

## Results

### Defining PSA, PSA-search, and PSA-sufficiency

A compositional phenotype is a phenotype with a collection of units and relationships between them (**Figure 1A, upper, black points indicate units and colored edges indicate relationships**). Examples of units are protein residues (**Figure 1**), brain regions (**Figure 2**), cells (**Figure 3-4**), and cellular neighborhoods (**Figure 5**). The relationships between units are proximity with respect to some notion of phenotypic distance.

**Figure 1.**
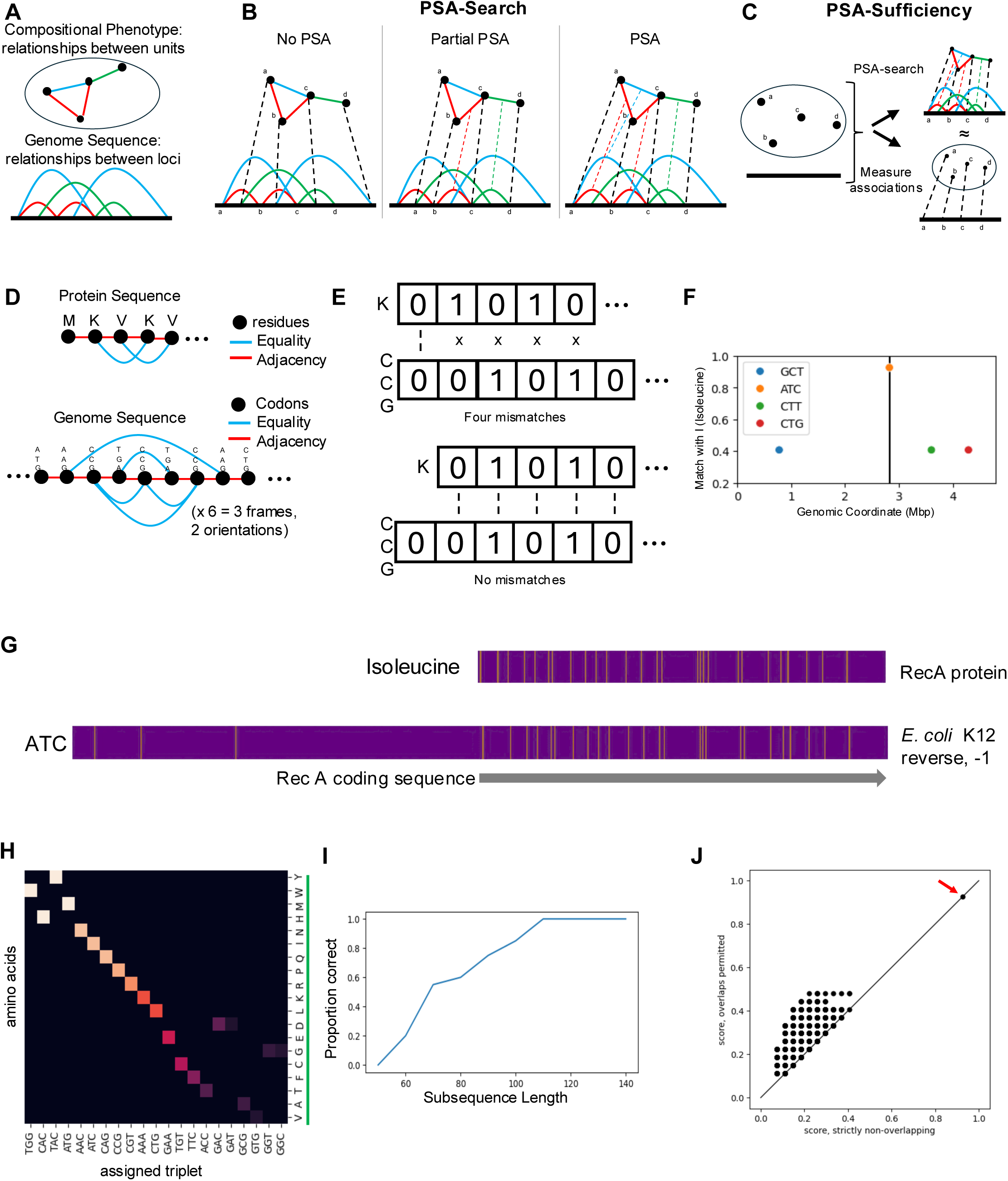
Expressed proteins are PSA-sufficient. A. Schematic illustrating compositional phenotypes consisting of units (black dots) with relationships (colors); the genome consists of loci (points on line) with sequence relationships (colors). B. Schematic illustrating PSA and PSA-search. Three different codes preserve relationships to different extents. Each code is an assignment of genomic loci to phenotypic units, represented by dashed black lines. A code (rightmost) exhibits PSA when relationships are preserved (dashed colored lines between edges). C. Schematic illustrating PSA-sufficiency. A phenotype respects PSA-sufficiency when a code exhibiting PSA is a sufficient constraint for it to recapitulate experimentally measured genotype-phenotype associations. D. Illustration of units and relationships in the context of amino acids relative to genomic 3-mers. E. Schematic showing how convolving one-hot encoded vectors for amino acids with those for 3-mers along the genome scores putative codes for PSA. F. Proportion of isoleucine equality relationships in RecA matched by putative assignment to indicated codons at different positions in *E. coli* K12 genome (maximum over frames/ orientations). Only positions/codons with >40% matching are shown. Black vertical line indicates coding sequence of RecA. G. Upper: binary vector with placement of amino acid I in Rec A. Lower: binary vector with placement of nucleotide triplet ATC near RecA coding sequence. H. Assignments of nucleotides to amino acids (rows) induced by PSA-search. Entry color indicates frequency of observed assignment (only those nucleotides assigned >50% of the time to one AA are shown). Green indicates correct recovery of the genetic code for all entries. I. Proportion of subsequences of RecA for which PSA-search recovers the correct coding sequence J. Comparing match proportion for isoleucine in RecA at each position/codon in E.coli genome allowing for non-uniformly spaced (i.e. overlapping) codes (y-axis) with non-overlapping codes (x-axis). Dashed line indicates same score with overlapping/ nonoverlapping assignments. Red arrow indicates best match across genome.

**Figure 2.**
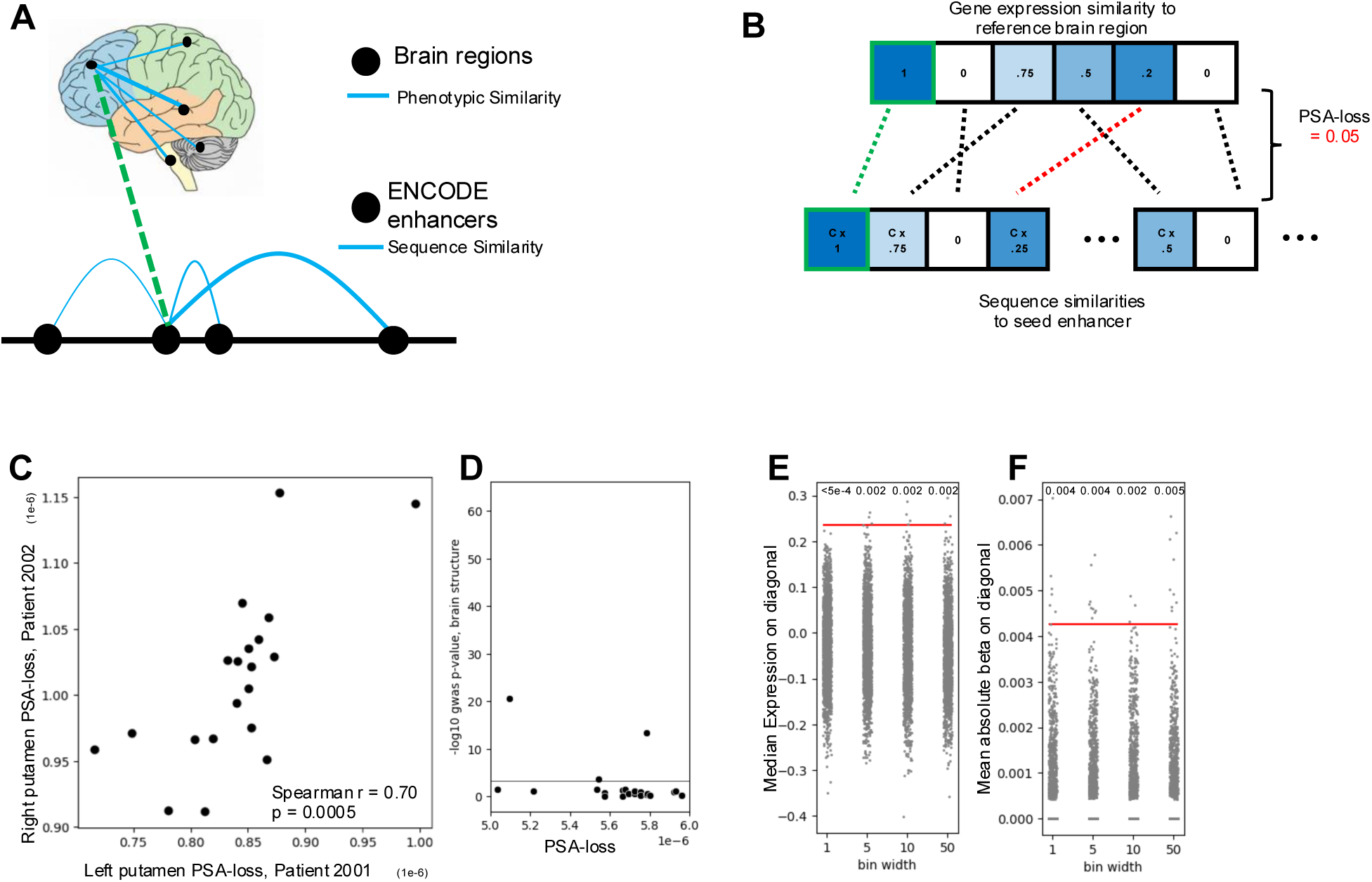
Brain anatomy is PSA-sufficient. A. Schematic illustrating the units and relationships in the context of brain regions (above) and enhancer sequences (below). Dashed green line indicates a seed assignment of an enhancer to a brain region. B. Schematic illustrating PSA-loss for a seed assignment of enhancer to a reference brain region by finding mismatch with best rescaling/reordering between vectors of phenotypic similarities to brain region and sequence similarities to seed enhancer. C. PSA-loss for left putamen in patient 2001 and PSA-loss for right putamen in patient 2002 for top twenty enhancers for left putamen. D. Maximum −log10 p-value for brain structure GWAS amongst two nearest SNPs for top 20 enhancers with lowest left putamen PSA-loss; line is GWAS p-value < 1e-4. E. Empirical null distributions across sampled codes of median expression on diagonal, controlling for sequence similarity at differing granularity. Red line indicates diagonal expression with PSA-code. F. Empirical null distributions across sampled codes of mean region-specific GWAS effect size, controlling for sequence similarity at differing granularity (x-axis). Red line indicates mean region-specific GWAS effect size with PSA code.

**Figure 3.**
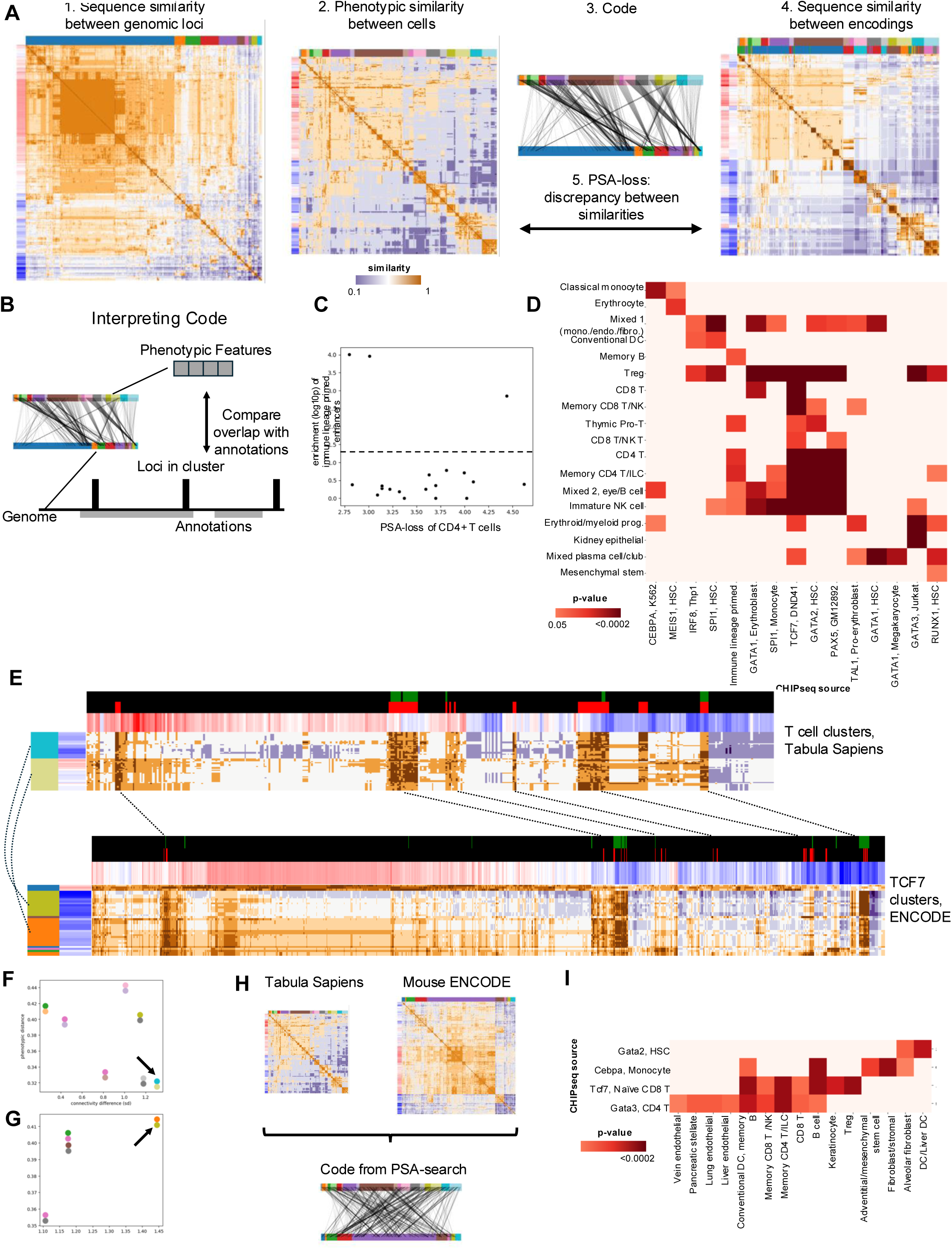
Single-cell transcriptional diversity is PSA-sufficient. A. Schematic illustrating optimal transport identification of subset-codes. 1. Pairwise similarity between clusters of ENCODE enhancers clustered by sequence similarity. Column colors indicate groupings by similarity relationships to other clusters, row colors are overall connectivity. 2. Pairwise similarity between cell clusters (metacells) from Tabula Sapiens atlas, row/column colors as with sequences 3. A code maps cell clusters to enhancer clusters (indicated by black arrows) 4. The code induces sequence similarities between cell clusters (row/column colors combined from 2A.1 and 2A.2. 5. Semirelaxed OT finds code such that phenotypic similarity between cell clusters matches sequence similarity between assigned enhancer clusters. B. Schematic illustrating interpretation of subset codes. Annotations are obtained for the phenotypic features (top) for each cell cluster and the genome (lower grey bars under black line). The loci contained in each locus cluster are overlapped with annotations and these are compared to phenotypic annotations. C. Log10 fisher exact p-value for enrichment of hematopoietic lineage primed enhancers (y-axis) in enhancer cluster assigned to CD4+ T cells across codes obtained from different clustering initializations, scored by PSA-loss on CD4+ T cells (x-axis) D. GC-adjusted enrichment (nominal p< 0.05) for CHIPseq peaks for immune TFs in the enhancer cluster assigned to each cell type in Tabula Sapiens atlas, color indicates p-value. E. Visual depiction of structural correspondence between TCF7 enhancer clusters and T cell clusters. Upper: Similarity between T cell metacells (rows) in Tabula Sapiens to other metacells (columns). Lower: similarity between TCF7 enriching loci clusters in ENCODE to other loci clusters. Black arrows indicate matching by code. F. Comparing pairs of blocks of cell clusters by phenotypic difference (y-axis) and average connectivity difference (x-axis). Blocks of T cell clusters in E indicated with arrow; only pairs of blocks with average phenotypic difference < 0.45 are shown. G. Comparing pairs of blocks of enhancer clusters by sequence difference (y-axis) and average connectivity difference (x-axis). Blocks of TCF7 cell clusters in E indicated with arrow; only pairs of blocks with average phenotypic difference < 0.45 are shown. H. Schematic showing how a PSA-search is applied to find a code for Tabula Sapiens within mouse ENCODE enhancers. I. GC-adjusted enrichment (nominal p<0.05) for for CHIPseq peaks for mouse TFs (columns) in the mouse enhancer cluster assigned to each Tabula Sapiens cell type, color indicates p-value.

**Figure 4.**
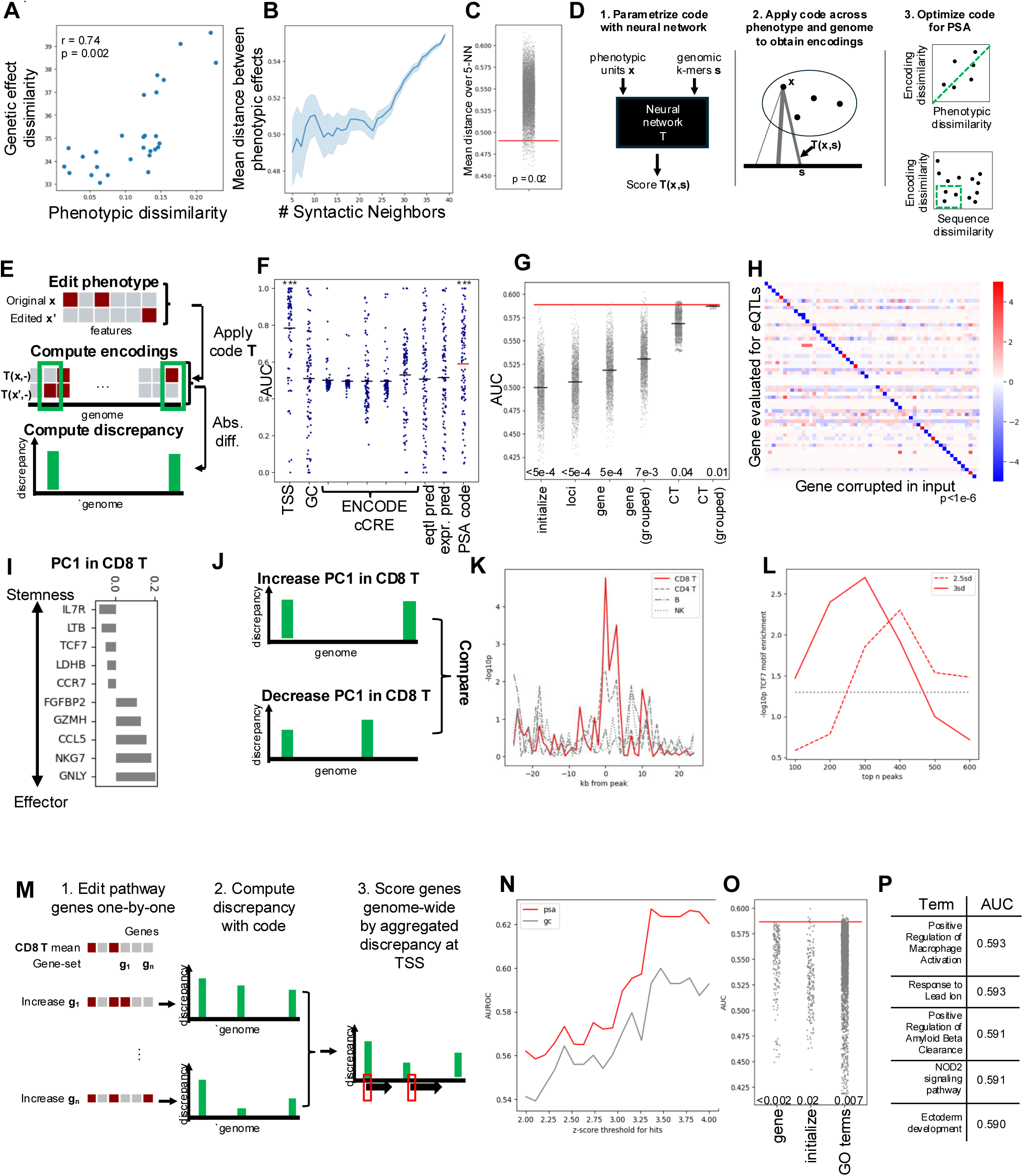
Benchmarking PSA-sufficiency of immune-cell transcriptional diversity. A. Correlation between phenotypic dissimilarity (cosine distance) between cell types and dissimilarity between eQTL effects (L1 distance) in Yazar et al. data. B. Average pairwise Jaccard distance between sets of significantly associated cell types (y-axis), averaged across k-nearest neighbors (x-axis) radii of SNPs by syntactic similarity measured with GENA language model. Range is standard deviation. C. Empirical null distribution (grey dots) for average pairwise Jaccard distance amongst 5-nearest syntactic neighbor SNPs when syntactic similarity features are permuted. Red line is test statistic: average pairwise distance with original graph. D. Schematic illustrating searching for codes exhibiting PSA with neural networks. 1. A neural network outputs the encoding **T(x,s**) between input phenotypic units **x** and genomic k-mers **s**. 2. Applying the code **T** across all units of phenotype and loci in genome produces a putative, real-valued encoding of the phenotype. 3. PSA is expressed with differentiable losses (top: the phenotypic dissimilarity between points should be approximately equal to the dissimilarity between their encodings, lower: syntactically similar loci should encode similar phenotypic units). E. Schematic illustrating QTL predictions with a code. A phenotypic unit is edited in silico by altering its features. The code **T** is applied to the original unit, and edited unit. QTL predictions are loci where the discrepancy between the encoding of the original unit and the encoding of the edited unit is high. F. Area under receiver operator curve (AUC) for eQTL prediction across genes; distance to TSS has the highest mean followed by the PSA code. G. Empirical null distributions for AUC from computing discrepancy for permuted edits. CT: cell type. Gray dots are AUC from null distribution samples, red line indicates test statistic (AUC obtained with PSA code). Numbers indicate p-values. Black lines indicate mean. H. Prediction AUC for eQTLs (rows) on models trained from corrupting each gene (columns) in input, z-normalized across rows; permutation p<1e-6 for diagonal. I. Highest magnitude loadings for first principal component of gene expression in CD8+ T cells from Hao et al. J. Schematic illustrating comparison of discrepancies from increasing cells by PC vs. decreasing cells by PC. K. Log10 t-test p-value for discrepancy difference for first PC in different cell types on memory vs effector CD8 T ATACseq peaks as a function of shifting peak location in genome. L. Log10 permutation p-value for colocalization of high-attribution regions on memory peaks with TCF7 motifs, with thresholds of 2sd and 2.5 sd per sequence as defining high-attribution regions, and across top N peaks (x-axis) by discrepancy difference. M. Schematic illustrating how pathway-level discrepancy scores for genes are obtained by scoring genes by discrepancy at TSS, aggregated across pathway genes increased one-by-one. N. Comparing CRISPR hit prediction AUC with interferon gamma geneset discrepancies from code and GC content as threshold for hit-calling is varied (x-axis). O. Null distributions for CRISPR hit prediction AUC, right indicates Type II Interferon gene set relative to other gene sets with at most 100 genes. P. Table of genesets with discrepancies best predicting CRISPR-screen hits.

**Figure 5.**
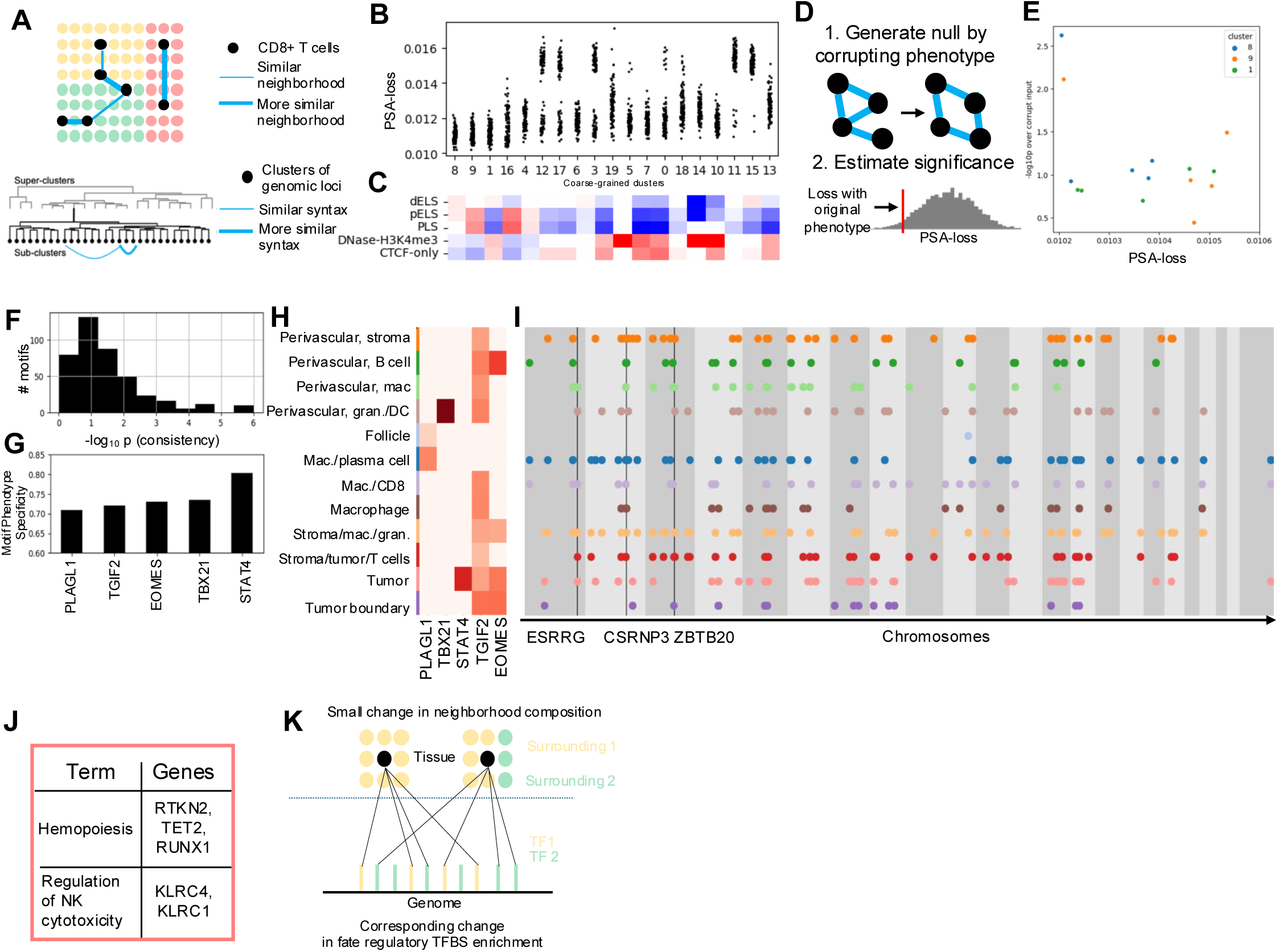
PSA-sufficiency of intratumoral CD8+ T cell neighborhoods reveals spatial gene regulatory syntax from images. A. Schematic illustrating de novo PSA for CD8+ T cell spatial gene expression. Upper: phenotypic units are CD8 T cells, considered similar when they have similar surrounding sets of cell types. Lower: loci from reference genome are clustered coarsely into twenty superclusters, and OT is used to search for a code within subclusters of each supercluster. B. PSA-loss for CD8+ T neighborhoods (y-axis) cell over sub-clustering initializations for each coarse-grained cluster (x-axis). C. Significant enrichment (FDR < 0.01) of ENCODE cCRE marks in each coarse-grained cluster, color indicates fold change. D. Schematic illustrating null distribution to evaluate phenotype specificity of PSA-loss by corrupting input phenotype similarity graph. E. PSA-loss and log10 p-value against specificity null for top three codes in top three coarse-grained clusters. F. Distribution of 417 HOMER inbuilt motifs by consistency of fold enrichment in clusters relative to permutations of the code. G. Top five consistent motifs ranked by specificity of consistency to input phenotype H. Heatmap of FDR < 0.0001 motif enrichment of specific motifs from (G) in assignments to each neighborhood cluster. I. Loci assigned to each neighborhood across chromosomes; genes with TSS within 30kb of more than four loci clusters in assignment labelled with black lines; only loci within 10kb of other loci clusters shown. J. Gene set enrichment analysis of TSS between 10kb and 30kb of loci cluster assigned to tumor-cell rich neighborhood. K. Schematic for spatial gene regulation encoding suggested by code. Upper: smooth changes in the neighborhood composition of a cell are accompanied by smooth changes in distribution of motifs of fate regulatory TFs.

We consider a genome as a set of loci (**Figure 1A, lower, points on black line**), with relationships between them **(Figure 1A, lower, colored edges between loci**). The relationships that we consider in the genome are proximity with respect to some notion of distance (not necessarily physical) between sequences. For more details, see **Supplementary Note 1.**

We define a genetic code to be an assignment of genomic loci to phenotypic units (**Figure 1B, dashed black lines**). We consider three types of code: single-locus codes (where each unit is assigned a single genomic locus), subset codes (where each unit is assigned a subset of loci) and real-valued codes (where each unit is assigned a weight for each locus in the genome). The locus (or set of loci or weights, for subset or real valued codes respectively) assigned to a phenotypic unit is its encoding. We first focus on single-locus codes.

Given a compositional phenotype, we can search through different codes and evaluate whether they preserve relationships (**Figure 1B).** We say that a code for a compositional phenotype exhibits Phenotype Sequence Alignment (PSA) when the relationships between its units are in correspondence with the relationships between their encodings (**Figure 1B, rightmost, dashed colored lines between edges of phenotype and genome)**. We refer to this task as PSA-search. The specific algorithm required to implement PSA-search depends on the structure of the phenotype and assumptions on the code.

Given any code for a compositional phenotype, we can evaluate its biological accuracy by comparing it to empirically established genotype-phenotype associations (**Figure 1C, right**). When the code produced by applying PSA-search to a compositional phenotype is biologically accurate, we say that the phenotype is PSA-sufficient. PSA-sufficiency is a property of a compositional phenotype.

Our central hypothesis is that for combinatorially complex compositional phenotypes, across scales, the only codes that respect PSA also agree with established genomic associations. We investigate this by formulating PSA-search algorithms, applying them to compositional phenotypes at molecular, cell and tissue scales and evaluating whether the output codes agree with measured biological associations.

### Rediscovering the genetic code from just one protein and genome sequence

The amino acid code is the prototypical genetic code. It specifies the genetic encodings of proteins. It was established by observing the RNA and protein output from cells as a function of input sequence (Crick et al., 1961). Given the right assumptions and computational capacity, can the amino acid code be found using just the genome sequence and protein sequence?

A protein sequence is a compositional phenotype, as follows. The units are residues (**Figure 1D, upper, black dots**), and the relationships are equality (when two residues have the same amino acid **Figure 1D, upper, blue lines between residues**) and adjacency (when two residues are adjacent, **Figure 1D, upper, red lines between residues**). We represent the genome as a set of nucleotide triplets (a DNA string of length 3n has 2(3n-2) accounting for 3 frames and 2 orientations). The relationships between loci are equality and adjacency as with the protein sequence (**Figure 1D, lower, colored lines**).

Given an amino acid sequence, we perform PSA-search as follows. Each amino acid in the protein determines a binary vector corresponding to the residues in the protein sequence where that amino acid is observed. Each of 64 nucleotide triplets determines six binary vectors corresponding to the locations it is observed in the genome, accounting for three frames and two orientations. We slide each amino-acid binary vector along each codon binary vector. The PSA-loss is the distance between the two binary vectors (**Figure 1E**). The output of the optimization problem is the position and amino-acid/triplet match with the smallest PSA-loss.

When PSA-search was applied to find codes for the recA protein in the *E. coli* genome, the best matching of binary vectors was uniquely obtained when I (isoleucine) is matched with the triplet ATC at the coding sequence of recA in the genome (**Figure 1F, vertical line indicates recA coding sequence**). The alignment between binary vectors can be observed visually (**Figure 1G**). After this best alignment has been found, adjacent nucleotide triplets in the genome are assigned to adjacent amino acids in the recA amino-acid sequence. In this way, PSA-search rediscovers the true genetic code from one protein and one genome. (**Figure 1H**).

The complexity of the protein sequence defines when PSA-search recovers the true coding sequence. We applied PSA-search to subsequences of the RecA protein in the *E. coli* genome. Only when the length of the subsequence is greater than 110 does the best matching location consistently yield the true coding sequence (**Figure 1I**).

Historically, establishing that the genetic code was a non-overlapping triplet code was an experimental discovery. A triplet code is the most parsimonious code (given 20 amino acids, there are only 16 nucleotide pairs). PSA-search shows quantitatively that a nonoverlapping code is the optimal such. We performed PSA-search but allowed overlapping spacing between triplets and computed the PSA-loss as above. Although allowing overlapping assignments improved the PSA-scores for RecA at each position in genome, it did not improve the PSA-score for the non-overlapping assignment at the RecA coding sequence (**Figure 1J**), indicating that the non-overlapping assignment is the optimally scoring one.

Thus, PSA-search with genomic 3-mers recovers the amino acid code from a single protein and genome sequence without genotype-phenotype covariation or prior mechanistic knowledge of coding sequences. This confirms that, in the context of proteins, structural correspondence is sufficient to identify causal genetic mappings: protein sequences are PSA-sufficient.

### PSA-sufficiency at the organ scale: similarity-preserving assignments of enhancers to brain regions

PSA-sufficiency held for proteins because the input amino acid sequence had a rich combinatorial structure that constrained assignments of genomic loci. We next asked whether PSA-sufficiency generalizes beyond the protein scale to complex organs, reasoning that phenotypes at the organ scale—such as the anatomical organization of the brain—possess a similarly rich structure.

Current understanding of gene regulation justifies PSA in this context. Anatomical regions that are phenotypically similar are expected to have similar active transcription factor circuits, so enhancers active in similar regions should, on average, possess more similar sequences than those active in distinct regions. The question then becomes PSA-sufficiency: does the correspondence between similarity structures provide sufficient constraint to coherently assign enhancers to brain regions without incorporating gene identity or covariation? We investigated this question in the context of brain regions in the Allen brain atlas (Hawrylycz et al., 2012) with enhancers from ENCODE encyclopedia (Moore et al., 2020).

The Allen brain atlas is a compositional phenotype. The units are brain regions (**Figure 2A**, **upper**), and the relationships are measured by gene-expression similarity. As a simplifying restriction to enable PSA-search, we sought a single-locus code (i.e., assigning one enhancer to each brain region). After preprocessing (we binned enhancers in ENCODE into loci of length 2000bp), there were 298,084 enhancer regions and 327 brain regions. The relationships between enhancers are sequence similarity (the choices of metric are discussed below).

We implemented PSA-search with an approach analogous to the brute-force one used for proteins (**Figure 2B)**. For computational feasibility of brute-force search, we selected three reference regions from the atlas and assigned enhancers to each reference region one-by-one as seeds for the code. Analogous to the amino-acid code example, a seed is an assignment of one enhancer to one region that can be extended to an assignment of enhancers to all regions (**Figure 2B, green boxes and edge indicate seed assignment)**. We computed, for each seed assignment, a PSA-loss (explained below) analogous to the number of mismatches when an amino acid in the protein sequence is assigned a particular genomic codon.

PSA-search requires choosing similarity metrics between phenotypic units and between sequences. The cosine distance between gene expression vectors (**Methods**) was the most conserved distance metric across the two donors with bilateral regional samples in the brain atlas (**Supplementary Figure 2A**). Spectral embedding captured evolutionary linkage between enhancers better than direct comparison of k-mer frequencies (**Supplementary Figure 2B, Methods**) and therefore was our choice of sequence similarity.

We score each seed assignment (i.e., an assignment of an enhancer to a reference brain region), with a PSA-loss. Here, the PSA-loss is the maximum mismatch when the vector of gene expression similarities of other brain regions to the reference region is inserted into the vector of sequence similarities of other enhancers to the seed enhancer. We allow for a global scaling and reordering of similarities (**Figure 2B, red line illustrates mismatch)**. Zero PSA-loss means that the set of distances from the assigned enhancer to other enhancers can be scaled to contain the set of distances of other brain regions to the reference region. We computed the PSA-loss when each of the 298,084 enhancer regions were assigned, one-by-one, to each of three reference regions as seeds: the left putamen, the left hypothalamus and left hippocampus. This process is analogous to sliding the binary vector for an amino acid along genomic loci in **Figure 1E**.

We investigated the seed assignments for the putamen, because their PSA-loss was consistent across the two individuals in the atlas. Specifically, the PSA-loss amongst the top seed enhancers for the putamen in Patient 2001 was significantly correlated with the PSA-loss measured for the putamen in Patient 2002 (**Figure 2C**). It was not significantly correlated with their PSA-loss when assigned to other brain regions (**Supplementary Figure 2C, center column)**. The PSA-losses for the top seeds for the hippocampus or the hypothalamus in Patient 2001 were not correlated with their PSA-losses when assigned to the same regions in Patient 2002 (**Supplementary Figure 2C outer columns).**

This indicates that the PSA-loss scores enhancers consistently for brain regions, but can also be driven by spurious, non-specific matches between enhancer and phenotype similarity distributions (**Supplementary Figure 2D)**. The fact that the PSA-loss for the enhancers assigned to the putamen is more conserved across biological replicates than for other regions is consistent with the putamen being a brain region with highly heritable anatomy (Zhao et al., 2019).

Two of the top twenty seed enhancers with lowest PSA-loss for the left putamen were proximal to GWAS hits for brain morphology (van der Meer et al., 2020) (**Figure 2D).** This enrichment (two hits out of twenty enhancers) was significant relative to background distributions: (i) relative to other enhancers across different hit thresholds (**Supplementary Figure 2E)** (ii) relative to null distributions sampling enhancers adjusting for GC content and sequence content (**Supplementary Figure 2F**), and (iii) sampling enhancers by Gene-Ontology annotations (**Supplementary Figure 2G)**. The Gene Ontology terms with respect to which the enrichment was not significant were terms plausibly involved in control of brain structure (**Supplementary Figure 2H**), which would be expected to enrich GWAS hits for brain morphology.

Each initial assignment extends to a code: enhancers are assigned to brain regions by matching similarities between sequences to similarities between regions (**Figure 2B, black dashed lines indicate best matches)**. Of the top five codes, the code with third lowest PSA-loss was a biologically coherent assignment of enhancers to brain regions, evaluated by region-specific expression and GWAS effects.

First, the expression of genes containing the enhancers in the code was on average higher in the brain regions to which they were assigned. We averaged the regional expression of genes that overlapped enhancer loci (183/327 regions were assigned enhancers that overlapped genes) (**Supplementary Figure 2I**). The expression along the diagonal was higher than off-diagonal – i.e., the expression in the regions of the genes containing the enhancer assigned to each region vs other regions (**Supplementary Figure 2J**). This was statistically significant relative to null distributions sampling assignments preserving sequence similarity to the seed enhancer (**Figure 2E**). This was also significant considering Bonferroni correction for testing the top codes.

Second, enhancers had on average higher GWAS effect sizes on the region-specific volumes of the brain regions to which they were assigned. We mapped regions from the brain atlas to regional volume GWAS from (Zhao et al., 2019) (**Supplementary Table 1)**. Enhancers assigned in the code to the triangular part of the inferior left frontal gyrus and left medial group of nuclei had absolute betas of 0.14 and 0.08 (with nominal GWAS p<0.001) on the volumes of the left pars triangularis and left thalamus proper GWAS (**Supplementary Figure 2K, blue boxes).** The code’s average region-specific effect size was significantly higher relative to the same null distributions as expression (**Figure 2F).**

Taken together, these data provide statistical evidence that the phenotypic similarity structure of the brain provides sufficient constraint to define codes that are consistent across individuals and biologically coherent. Thus, brain anatomy, relative to ENCODE enhancers, is partially PSA-sufficient.

### PSA-sufficiency of single-cell transcriptional diversity: assigning sets of enhancers to cells in the Tabula Sapiens atlas

We next investigated whether PSA-sufficiency held at the scale of cellular gene expression. In this context, phenotypically similar cell types typically share a developmental lineage and therefore the regulatory programs that drove that lineage. Therefore PSA is expected: the set of enhancers defining the developmental histories of similar cells should, on average, be more similar in sequence motif composition than the sets of enhancers defining the histories of distinct cells. The central question is therefore PSA-sufficiency: can PSA-search recover specific enhancers driving the developmental history of a cell type using only the similarity structure of a cell atlas?

We implemented PSA-search to assign sets of enhancers to cells in the Tabula Sapiens atlas (Consortium* et al., 2022), and evaluated whether the resulting code recapitulated enrichment of cell-type lineage-specific transcription factor binding. We first constructed similarity matrices between clusters of cells and clusters of enhancer loci, observing common structure to them.

We used the binned enhancer loci as in the brain analysis (a total of 298,084 enhancer regions) and grouped them into 1000 clusters with spectral clustering. We computed pairwise manifold distances (i.e., shortest paths in the nearest-neighbor graph) between clusters (**Figure 3A.1).** There was a core-periphery structure: a large central block of clusters with high average connectivity to other clusters (**Figure 3A.1, column colors indicate blocks and red row color indicates connectivity to other clusters),** as well as smaller blocks with limited connectivity with other non-central clusters.

We grouped cells from the Tabula Sapiens into 400 dense clusters using SEACells (Persad et al., 2023), a metacell approach for single-cell RNAseq, optimized to preserve the geometry of the cell distribution, and computed manifold similarities between clusters (**Figure 3A.2**). The overall structure reflected that of the enhancers: a large central block of clusters (containing stem cells and epithelial cell subsets), as well as smaller blocks of clusters with limited similarity to other non-central clusters (mainly immune cell subsets and fibroblasts) (**Supplementary Figure 3A, annotations).**

We performed PSA-search to find the optimal code: a map from cell clusters to enhancer clusters (**Figure 3A.3, right, upper)** that optimally preserves pairwise similarities (**Figure 3A.4**). The PSA-loss measures the discrepancy between the phenotypic similarity between cell clusters and the sequence similarity of assigned enhancer clusters. The matrix in **Figure 3A.4** comes from the extracting the entries of the sequence similarity in **Figure 3A.1** indicated by assignment between colors in **Figure 3A.3**. The fact that the two matrices (**Figure 3A.2** and **Figure 3A.4**) are similar indicate that PSA-search has found a code consistent with PSA.

The search is performed using a semi-relaxed Gromov-Wasserstein optimal transport (OT) algorithm (Vincent-Cuaz et al., 2022). The semi-relaxed OT variant is used because not all enhancer clusters should be assigned to a cell cluster. We matched manifold distance (path distance between symmetrized nearest neighbor graphs) in OT, since this is the best performing distance metric for multimodal single-cell feature alignment with OT (Demetci et al., 2022). The maximum distance between pairs of enhancer clusters and between pairs of cells was normalized to one. This normalization is justified because it corresponds to the assumption that the subsets of cells that are most different can be assigned to maximally dissimilar enhancer sequences, which is reasonable because Tabula Sapiens dataset contains diverse cell types. Applying this procedure with different random initializations of the clustering algorithm (k-means on spectral features) changes where boundaries are drawn between enhancer clusters and therefore yields codes differing in their PSA-loss.

We investigated the biological coherence of the lowest PSA-loss code. Given a code, we evaluate its biological coherence by comparing the gene-expression annotations of the cell clusters (**Figure 3B, phenotypic features**) with genomic annotations of the enhancer clusters to which they are mapped (**Figure 3B, lower**). Across random seeds, the code with the lowest PSA-loss on CD4+ T cells (excluding a code mapping CD4+ T cells to a cluster of only 19 enhancers) assigned CD4+ T cells to an enhancer cluster that was significantly enriched for hematopoietic lineage primed enhancers (**Figure 3C**), experimentally determined through mitochondrial lineage tracing (Weng et al., 2024). Thus, the code optimally preserving PSA may map cell types to genomic loci involved in their development.

We further evaluated the code’s biological coherence by evaluating enrichment of hematopoietic driver transcription factor binding measured with CHIP-seq, adjusting for GC% (**Supplementary Figure 3I** for unadjusted data, **Supplementary Table 2** for data sources) in all enhancer clusters comprising the code. There was significant enrichment (nominal p<0.05) of immune TF peaks in multiple enhancer clusters, and this enrichment was correct and specific to cell types they were assigned to, as we describe next (**Figure 3D**).

The binding of CEBPA, a key myeloid TF was enriched only in the enhancer cluster assigned to classical monocytes, erythroid/myeloid progenitors and a mixed subset immune/eye cells. The binding of MEIS1, a key erythroid TF, was enriched in the enhancer cluster assigned to erythrocytes and weakly in the enhancer cluster assigned to classical monocytes. Only T cell subsets (and one B cell cluster) were assigned enhancer clusters strongly enriched in the T cell master regulator TCF7. The hematopoietic lineage primed enhancers were enriched only in clusters assigned to immune cells. The binding of IRF8, an important dendritic cell and Th1/Treg transcription factor, was enriched in the enhancer clusters assigned to dendritic cells, Tregs, and the enhancer cluster assigned to a mixed monocyte/endothelial cluster.

The code also correctly assigned immune TF-enriching enhancer clusters to non-immune cell types. Specifically, mesenchymal stem cells were assigned to an enhancer cluster enriched for RUNX1 peaks and kidney epithelial cells were assigned to an enhancer cluster enriched for GATA3 peaks. GATA3 and RUNX1 play foundational roles in the development of kidney epithelial cells (Grote et al., 2006) and mesenchymal stem cells (Kim et al., 2014) respectively.

These data indicate that transcriptional diversity, at least in the context of immune cells, is PSA-sufficient: structural correspondence is sufficient to coherently assign enhancers to cell types that enriches for lineage-specific TF binding.

### Evolutionary conservation of the structural correspondence between T cells and TCF7 motifs

Is the code output by PSA-search capturing a fundamental structural correspondence? In the case of the amino acid code, the structural correspondence observed in **Figure 1G** between isoleucine in the recA protein and ATC in the *E. coli* genome, is universal across proteins and species. As potential evidence for its fundamental nature, we investigated the structural correspondences in the code.

The structural correspondence between TCF7 enhancer clusters and T cell clusters (i.e., the matching between their similarities to other enhancer and cell clusters to each other) was directly visible, mirroring that between isoleucine in recA and ATC in the *E. coli* genome in **Figure 1G**. T cell clusters consisted of two blocks (one enriching for CD4+ T cell clusters and one for CD8+ T cell clusters). These two blocks had high similarity amongst themselves, but differed in their average connectivity, and had similarity with a shared central cell clusters (corresponding to stem cells) as well as with low-connectivity blocks (**Figure 3E, upper)**. Likewise, TCF7 enriching enhancer clusters had two subsets with high similarity amongst themselves, differed in their average connectivity, and had similarity with both central and isolated blocks (**Figure 3E, lower**). We directly quantified this correspondence: amongst phenotypically similar blocks of cell clusters, the blocks of T cell clusters differed the most in their average connectivity (**Figure 3F)** and the same was true for TCF7 enhancer clusters (**Figure 3G**).

While the organization of T cells into subsets and its regulation by TCF7 is evolutionary conserved, we investigated whether their structural correspondence is as well. We searched for a code for Tabula Sapiens in mouse enhancers (**Figure 3H**).

The core structural correspondence between T cells and TCF7 is conserved: the cross-species code assigned T cells to enhancer clusters enriched for TCF7 peaks (**Figure 3I)**. Further assignments in the code were also correct. CEBPA peaks were enriched in the enhancer clusters assigned to dendritic cells (myeloid cells, albeit not monocytes). In addition, GATA3 peaks were enriched in the assignments to endothelial cell subsets (where it is shown to be a regulatory of Tie2 signaling, (Idowu et al., 2021)), and GATA2 peaks in the assignment to dendritic cells. However, the cross-species code also produced incorrect assignments: alveolar fibroblasts were mapped to an enhancer cluster enriched in GATA2 and keratinocytes were mapped to an enhancer cluster enriched in TCF7. Thus, while core correspondences are conserved, the code reflects divergences, which may arise from artefacts of the input data or genuine evolutionary differences.

Taken together these data show that structural correspondence between phenotype and enhancer sequences is sufficient to recover gene regulatory relationships across cell types, particularly in the context of immune cells. These data further highlight the possibility to deduce unexpected genetic regulators for cell types from PSA-search alongside functional genomic data in distal cell types or organisms. While previous studies have shown deep conservation of enhancer syntax, these data suggest further conservation of the structural correspondence between global enhancer sequence distribution and phenotypic structure. A key limitation of this analysis is the bias inherent in the cell types included in ENCODE and shared with Tabula Sapiens.

The data from the brain and Tabula Sapiens together indicate that PSA-sufficiency is a general principle underlying the gene regulatory encodings of phenotypes at the organ and single-cell scale.

### Benchmarking PSA-sufficiency without annotations: expression quantitative trait loci exhibit PSA

We showed above that PSA-search assigned enhancers with cell-type specificity in the context of immune cells, but this required pre-defining functionally relevant genomic loci (i.e., the ENCODE atlas). We next sought to remove this dependency and statistically benchmark whether the phenotypic diversity of immune cells is PSA-sufficient without prior annotations of the genome.

We reasoned that genetic regulation of cell type specific gene expression should exhibit PSA: phenotypically similar cell types should be regulated by similar genetic variants. We first evaluated this assumption using single-cell expression quantitative trait loci (eQTLs). eQTLs are loci where variant dosage is correlated with gene expression in a cell type. The effect sizes at eQTLs are not binary, but real-valued scores. We therefore define continuous codes to evaluate PSA in this context, extending the definitions from subset and single-locus codes.

A continuous code is a function **T(x,s)** producing a score representing the strength of the association of the locus **s** with phenotypic unit **x**. Unlike the single-locus or subset codes above, PSA for continuous codes has two distinct components:

- A code **T(x,s**) is phenotypically consistent if similar units **x, y** have similar vectors of encodings across loci **T(x,-), T(y,-)**.
- A code is syntactically consistent when genomic loci **s, t** that are syntactically similar have similar vectors of encodings **T(-, s), T(-,t).**

We say that a code exhibits PSA when it is both phenotypically and syntactically consistent. As with the single-locus and subset codes, defining PSA for continuous codes requires specifying phenotypic and syntactic similarity metrics. PSA-sufficiency for continuous codes is when these two conditions are sufficient to produce biologically coherent codes. PSA-sufficiency for continuous codes is formalized in **Supplementary Note 1**,

In the context of immune cell gene expression, eQTL effects are phenotypically consistent. In the dataset of (Yazar et al., 2022), cell types with similar gene expression (as determined in the single-cell atlas of (Hao et al., 2021)) had more similar eQTL effects than those with distinct gene expression (**Figure 4A**). This property did not depend on the choice of distance metric (**Supplementary Figure 4A-B).** This property also held on a gene-by-gene basis in the 2MB cis-windows around transcription start sites in which eQTLs were evaluated (**Supplementary Figure 4C)**.

Immune cell eQTLs are syntactically consistent. We computed syntactic similarity between 2000-mers as their similarity in the embedding space of the GENA human genome language model (Fishman et al., 2023). Loci that were neighbors in the similarity neighborhood graph contained eQTLs for similar sets of cell types and this was significant with respect to a null distribution permuting the similarity graph (**Figure 4B and Figure 4C, grey dots indicate null distribution, see Methods)**. Embeddings from a different language model with shorter context (500bp) were not significantly associated (**Supplementary Figure 4D**). The GENA language model was the only model pretrained exclusively on one human reference genome (T2T) with byte-pair encoding to enable long-context feature extraction.

These data indicate that immune cell eQTLs exhibit PSA. The question then becomes whether immune cell transcriptional diversity is PSA-sufficient, which we investigate by implementing PSA-search to produce continuous codes without prior genomic annotations, and evaluating whether these codes correctly predict cell-type specific eQTLs.

### Implementing a neural network PSA-search algorithm to find continuous codes

We developed a minimal neural network optimization strategy to perform PSA-search (termed neural PSA-search) to obtain continuous codes that (a) enable prediction of quantitative trait loci, and (b) do not require prior genome annotations.

We parameterize a code with a neural network **T (Figure 4D.1).** The input to the code **T** is a phenotypic unit **x** (for single-cell gene expression, a gene expression vector of a cell) and genomic k-mer **s** (we used 2000-mers). The output of the neural network **T** is a real number representing the encoding **T(x,s)** of **x** at **s.**

Parametrizing codes with neural networks enables finding codes that respect PSA via gradient descent. We compute encodings of all the units **x** of a compositional phenotype at all loci **s** in the genome with the output of the network **T(x,s**) (**Figure 4D.2**). Phenotypic consistency (**Figure 4D.3, upper**), syntactic consistency (**Figure 4D.3 lower**) and sparsity are expressed as differentiable loss functions, whose gradients are used to optimize the parameters of **T**. Sparsity, along with the neural network architecture of **T** constrains the parsimoniousness of the codes.

The neural network representation of a continuous code enables predicting genomic loci associated with in-silico alterations to a phenotype because it predicts association scores for phenotypic units not included during training. If the code is biologically coherent, then the encoding of a unit **x** at a locus **s**, given by T**(x,s)** should represent the extent to which the sequence at locus **s** contributes to properties of **x**. Therefore, the difference between the encodings of different phenotypic units **x** and **y** at a locus **s**, |**T(x,s)- T(y,s)|,** should be large at loci **s** that differentially contribute to **x** and **y**. If **x** and **y** are gene expression vectors that differ only by a small amount in one gene, the loci responsible for this difference should have high discrepancy.

Neural PSA-search therefore provides a quantitative way to benchmark PSA-sufficiency. If the input phenotype is PSA-sufficient, discrepancies from the code obtained by neural PSA-search should colocalize with eQTLs and causal loci.

### Statistical benchmarking of *de novo* codes for eQTL prediction

We quantitatively validated that the continuous codes from neural PSA-search correctly predicted eQTLs, validating that single-cell transcriptional diversity of immune cells is PSA-sufficient without prior genomic annotations.

Neural network models trained on covariation between sequence and epigenetic or expression measurements, such as Enformer (Avsec et al., 2021), effectively predict eQTLs with gene and cell type specificity. Since codes from PSA-search only use similarity, and not gene sequence, localization or functional measurements, our hypothesis is not that we obtain comparable results to sequence-to-function models that do use such information, but rather that we observe a statistically significant and specific enrichment.

We applied neural PSA-search (**see Methods**) to a single-cell gene expression atlas of PBMCs **(Hao et al., 2021)**. We grouped cells from three donors into 200 metacells and computed log1p normalized expression. We tiled the T2T reference genome with 2000-mers (with an overlap of 1000) and extracted embeddings from the GENA language model. The architecture of the neural network representing the code **T(x,s**) consists of two networks: sequence embedding and phenotype embedding. The sequence embedding network applies a linear projection, and the phenotype embedding network applies a linear projection followed by softmax normalization. The final score is the inner product of sequence and phenotype embeddings followed by tanh activation to normalize scores between −1 and 1. The softmax layer constrains the expressivity of the code.

Our hypothesis was that discrepancies from the PSA-search code correctly enrich for eQTLs in a gene and cell-type specific manner. We refer to QTL predictions from the code as quantitative-trait scores (QT-scores). QT-scores quantify the change in the code following in silico perturbation to the phenotype (**Figure 4E).** For a given **g** cell type **c** and locus **s,** the QT-score for **g** in cell type **c** at **s** is the change in the code when we shift the mean expression vector of the cell type **c** a small amount in the direction of gene **g.** Specifically, the QT-score is given by | **T(x_c_ + δg, s) – T(x_c_,s)**|, where **x_c_** is the cell type’s mean gene expression vector and **δg** indicates a small shift in the direction defined by gene **g.** Specificity means that the QT-scores for a different gene **g**’ at **c** and the QT-scores for **g** at a different cell type **c’** do not enrich for the eQTLs of gene **g** in cell type **c**.

We evaluate the code with eQTLs from (Yazar et al., 2022) as follows. We use QT-scores for each gene at each cell type to predict whether a given locus (a 2000-mer extracted as above) contains a significant eQTL (FDR < 0.1) for that gene in that cell type. Our results are obtained by downsampling the eQTL dataset to SNPs in the GWAS catalog, because these cluster less around transcription-start sites (TSS) and indicate functionally relevant regulatory variation. We include results on the full set of SNPs, which exhibit the same significance but lower effect sizes.

The QT-scores had higher area under the receiver-operator curve (AUC) averaged across genes and cell types (0.589) than baseline approaches to predict eQTLs. The baseline approaches included: GC-content, proximity to candidate cis-regulatory elements derived from functional measurements by ENCODE (the best performing annotation was 0.536 for loci containing distal enhancers), a supervised classifier predicting eQTLs with sequence features (ruling out trivial sequence biases in eQTL annotations), and a supervised linear predictor of gene expression from sequence embeddings (**Figure 4F** and **Supplementary Figure 5A**).

As expected, proximity to the gene’s transcription start site (TSS) outperformed the QT-scores, with an AUC of 0.78 (**Figure 4F)**. However, this baseline requires knowing a gene’s location in the genome, information not included in PSA-search. We note that sequence-to-function models like Enformer outperform TSS proximity, but are trained including gene positions and other functional annotations.

We evaluated the specificity and significance of these predictions with empirical null distributions. Each null distribution (**Figure 4G, gray dots in columns)** ablated a distinct aspect of the information used to compute QT-scores (**Figure 4G red line indicates test statistic**).

- We permuted the correspondence between loci and their genomic features (**Figure 4G, loci)**; the p-value relative to this distribution was <0.0005 indicating that prediction uses the information in the sequence features.
- We randomly initialized the weights of the neural network representing the code (**Figure 4G, initialize, see Methods**). The p-value with respect to this null distribution was less than 0.0005, so enrichment of eQTLs is not an artefact of the parametric format of the code or random projections of the syntactic features.
- We permuted genes, so that QT-scores for predicting random genes were used as predictions for a given gene (**Supplementary Figure 4E, blue arrow indicates a different gene edited in the correct cell type)**. The p-value with respect to this null (0.0005) indicates that correct enrichment of eQTLs requires the specific geometry of the genes (**Figure 4G, genes)**. The mean of the null distribution is above 0.5, so that non-gene specific sequence features partially enrich for eQTLs. This is consistent with the AUC obtained by the baseline eQTL predictor.
- We performed grouped permutation of genes, so that eQTLs for a given gene could only be predicted with QT-scores from a gene detectable in the same set of cell types. The predictions are significant with respect to this null (p = 0.007), but less so than for ungrouped gene permutations. Thus, indicates that predictions from the code require finer geometry of gene expression beyond cell type expression, but not the full resolution in the input phenotype.
- We permuted cell types (**Figure 4E, third row, blue arrow indicates correct gene edited in a different cell type)**, so that QT scores are computed by permuting the correct gene, but in a random cell type. The weaker significance and higher average from the null distribution indicates that the AUC requires the geometry of the cell types used to compute the QT-scores, but the code also enriches for eQTLs with gene and not cell-type specificity (**Figure 4G, CT)**.
- We permuted cell types within lineage groups (T cells, B cells and Monocytes). The weaker significance with respect to this null (p = 0.01) indicates that the QT scores are cell-type specific only up to the coarse geometry of the cell types into major subgroups, but not with respect to the finer geometry of cell types.

eQTL predictions were accurate to a resolution of approximately 10kb. We obtained empirical distributions by permuting sequence features across loci proximal in the genome at different distances. The observed AUC was significant with respect to these distributions only when loci were permuted at a length-scale of greater than 10kb (**Supplementary Figure 5B, null distributions increasing permutation length-scale,** see **Methods)**. This is the approximate length-scale of linkage disequilibrium, (Altshuler et al., 2005) so we cannot resolve whether this is a limitation of the code or the evaluation data.

We further confirmed that phenotype-specific structure is required for PSA-search to find a coherent code by corrupting the phenotype input to PSA-search. We applied PSA-search to phenotypes that had been corrupted by permuting the expression values of each gene one-by-one (see **Methods**) and computed the AUC for predicting each gene’s eQTLs (**Figure 4H, heatmap showing per gene AUCs with corrupted inputs**). The AUCs changed only for recovering eQTLs for the genes that were corrupted in the input (**Figure 4H**, permutation p < 1e-6 that the diagonal is lower). The decrease on the diagonal was correlated with significance in the gene permutations (**Supplementary Figure 5C),** corroborating that both metrics evaluate gene specificity. Thus, PSA-search leverages the geometry of distinct genes to correctly enrich for their regulatory loci.

A comparable AUC and significance with respect to the null distributions was obtained when codes were obtained by applying PSA with a different random seed to data of the same three individuals (**Supplementary Figure 5D**). A weaker significance was observed with a biological replicate dataset of three other individuals from the same study, but required increasing the expressivity of the code (**Supplementary Figures 5E-G**). In addition, ablating each of the terms in the loss led to a decrease in the AUC (**Supplementary Figure 5H**).

The code from PSA achieved an AUC of 0.523 on the data of all SNPs, without downsampling GWAS SNPs, and this AUC was significant with respect to the gene and grouped gene permutations with various thresholds for filtering. It was not significant with respect to cell type permutations, indicating enrichment of eQTLs is not cell type specific when averaged across all loci. (**Supplementary Figure 5I**). This significance was robust to filtering genes by number of evaluated loci (**Supplementary Figure 5K).**

Beyond binary classification, lead variants (defined as the top three loci by effect size) had on average higher QT-scores than other evaluated variants. This increase was specific: it was significant with respect to the gene, grouped gene and cell type empirical null distributions (**Supplementary Figure 5J**). Furthermore, these results were robust to filtering genes by number of evaluated loci (**Supplementary Figure 5M)**.

These data indicate that PSA-search finds a code – without annotations – for the PBMC atlas that significantly and specifically enriches for expression quantitative trait loci. Thus, immune cell transcriptional diversity is PSA-sufficient even without prior genomic annotations.

### Identifying drivers of of PSA-sufficiency: T cell transcriptional diversity is encoded by TCF7 motifs

The code correctly predicted eQTLs, suggesting that it learned aspects of transcriptional regulation. If the code is biologically meaningful, its predictions should be driven by TF motif composition. We investigated this with neural network attribution.

The first principal component (PC1) of gene expression in CD8+ T cells corresponded to a T cell differentiation axis from naive/stemness genes to effector genes (**Figure 4I)**. We hypothesized that QT-scores from in-silico perturbations of this axis would specifically distinguish regulatory elements in stem cell memory and effector T cells. We computed the difference between the QT-scores from increasing along this PC and the QT-scores from decreasing along this PC in CD8+ T cells (**Figure 4J)**.

The QT-score difference was higher on peaks with increased accessibility in stem cell memory CD8+ T cells than on peaks with increased accessibility in effector memory CD8+ T cells. This effect was abrogated when peaks were shifted by more than 3kb (**Figure 3K)** and is therefore specific to the positioning of the peak-sets in the genome. This effect was also cell type specific: the discrepancy difference specific to the peak set on the first PC (computed in each cell type) was observed to a lesser extent in CD4+ T cells, and not in B cells or NK cells. Thus, the QT-score difference between stem and effector skewed CD8+ T cells captures differences in regulatory sequence features between stem and effector CD8+ T cells.

The sequence features distinguishing peak sets for the QT-score differences are not due to confounding with GC content or nonspecific sequence features. The stemness peaks were enriched in GC rich regions, but a significant GC% difference across peak sets was observed even with shifts of up to 30kb (**Supplementary Figure 6D**) and so this predictor does not exhibit the specificity of the QT-scores. In addition, there were QT-score differences when considering the first PCs for myeloid cell types, but these were also not specific to the peak set.

Given the structural correspondence we had observed in the ENCODE analysis (**Figure 3)**, and the fact that TCF7 is a fundamental transcription factor determining T cell memory/effector differentiation, we investigated whether the QT-score differences were driven by TCF7 motifs.

The QT-score differences were driven by sequence features colocalizing with TCF7 binding sites. For each memory peak, we corrupted its sequence by shuffling nucleotides in 50bp blocks (**Supplementary Figure 6E**) and computed the change in the QT-score difference for each sequence with the code. We permuted the 50bp blocks and observed that high-attribution regions amongst the top peaks significantly colocalized with 50bp blocks containing TCF7 motifs (**Figure 4L**). This was consistent across different thresholds for defining regions as high-attribution as well as different thresholds for selecting the top peaks. Furthermore, this effect held when only considering sequences with unique high-attribution region (**Supplementary Figure 6F**) and was therefore not driven by sequences with multiple binding sites.

Thus, neural PSA-search learns a code that translates local axes of phenotypic variation to axes of sequence feature variation determined by TF motifs of the regulators of that phenotypic variation.

These results indicate that the PSA-sufficiency of immune cell gene expression is driven by a structural correspondence between transcriptional diversity and motif distribution. Furthermore, these results corroborate that the specific structural correspondence between TCF7 motifs and T cell diversity is a fundamental aspect of how the genome encodes gene expression.

### *De novo* prediction of causal regulators from sequence with neural PSA codes

The PSA-sufficiency of amino-acid sequences resolved the loci that were causal without covariation. Is this also the case for the continuous code for PBMC single-cell diversity? Since the code correctly resolved transcriptional regulators of T cell effector differentiation, we hypothesized it could also discover genes with causal effects on effector gene expression from their sequence. We therefore evaluated whether QT-scores from the PBMC code could predict CRISPR screen hits for IFNG expression in primary CD8+ T cells.

Given a gene pathway, a code predicts which genes, genome-wide, are responsible for altering it (**Figure 4M**). For each gene in the pathway, we compute the QT-score when increasing it in CD8+ T cells at the TSS of every gene in the genome (allowing 2.5kb either side). This provides, for every gene in the pathway, a ranking of all genes in the genome. We average the rankings for each perturbed gene in the pathway to obtain an overall ranking of genes, termed the pathway QT-ranking (**Figure 4M, right)**. A gene’s resultant pathway QT-rank depends only the sequence near its TSS and not on whether it belongs to the pathway being perturbed.

The pathway QT-rankings for the *Regulation of Type II Interferon Signaling Pathway* had an AUC for predicting screening hits of 0.6. This was higher than the AUC obtained by GC content across a range of thresholds (**Figure 4N**). Furthermore, the gene-ranking had a significant p-value when included in a logistic regression model with GC content as a predictor (**Supplementary Figure 6B**), consistent across thresholds, indicating that it provides distinct predictive information.

The high AUC was specific and statistically significant. QT-rankings from perturbing random subsets of genes with the same size as the Type II Interferon Signaling term, as well as from random initializations of the code, and pathway QT-rankings from other GO terms had lower AUC (permutation p <0.002, initialization p = 0.02, p = 0.007 respectively, **Figure 4O**). This effect was not trivially due to expression: GO terms more highly expressed in T cells tended to have higher AUCs, but GO terms with highest AUC were not exceptionally highly or weakly expressed, indicating that this specificity cannot be explained by expression (**Supplementary Figure 6C)**. Taken together, these findings indicate the code has learned specific correspondences between sequence features of causal genes and their phenotypic effects.

We examined the GO terms whose pathway QT-scores had highest AUCs for predicting hits in the IFNG screen, hypothesizing that these might reveal unexpected pathways that are downstream of IFNG. The top pathways were: Positive Regulation of Macrophage Activation, Response to Lead Ion, Positive Regulation of Amyloid Beta Clearance, NOD2 Signaling pathway, and Ectoderm development (**Figure 4P**). IFNG has an established role upstream of Macrophage Activation (Hu et al., 2008), NOD2 Signaling (Rosenstiel et al., 2003) and Lead response (Gao et al., 2006). Furthermore, these provide a mechanistic hypothesis that IFNG plays an upstream role in Amyloid Beta clearance, a finding substantiated in mice in (He et al., 2020).

As a complementary way of evaluating whether the codes from neural PSA-search captured causal loci, we tested whether PSA enriched for fine-mapped eQTLs. We obtained a neural PSA code for the Tabula Sapiens cell atlas (Consortium* et al., 2022), and evaluated whether it specifically enriched for fine-mapped eQTLs with a high causal probability (PIP > 0.5) from GTEx (GTEx Consortium, 2020). Using the same methodology as for the PBMC code, the QT-scores for PSA obtained an AUC of 0.57 (**Supplementary Figure 6A). This** was significant with respect to gene, grouped-gene, loci and initialization empirical null distributions, but not with respect to the tissue permutations (**Supplementary Figure 6I**). Thus, the code identifies putatively causal gene expression regulators with gene specificity but limited tissue specificity.

Taken together, these results highlight the PSA-sufficiency of immune cell gene expression: neural PSA-search codes correctly enrich regulatory loci, without prior genomic annotations, and are driven by TF syntax which in turn enables causal predictions from sequence. Furthermore, these results provide an explanation for PSA-sufficiency: transcriptional diversity of cells is encoded through structural correspondence with regulatory motif diversity in the genome.

### PSA-search on colorectal cancer T cell neighborhood similarity reveals sequence regulators of spatial gene expression

The analyses above associated T cells with their regulatory loci using two different formulations of PSA-search and input data. However, the input phenotypes both (a) defined similarity with gene expression, a measurement closely linked with the genome, and (b) were from homeostatic contexts. We therefore investigated whether the spatial localization for CD8+ T cells in the tumor microenvironment was PSA-sufficient. Can we discover the genetics of a clinically significant phenotype directly from images, without genetic data, from a small number of individuals?

We used OT-based PSA search to find codes for CD8+ T cells in the colorectal cancer tumor microenvironment based on their spatial neighborhoods. The scalability of the OT PSA-search allowed us to generate enough codes to statistically evaluate optimality. From a multiplexed immunofluorescence dataset (Schürch et al., 2020), we densely clustered CD8+ T cells by the cell type composition of their 20 nearest neighbors as input for PSA-search and computed pairwise manifold similarities between clusters (**Figure 5A, upper)**.

Unlike the code for Tabula Sapiens in enhancers, we expected the code to be constrained within a smaller portion of the sequence space of genomic loci – the maximum distance between T cells. We therefore coarsely clustered the GENA language model embeddings for 2000-mers in the genome into twenty super-clusters. Within each super-cluster, we sub-clustered loci into 500 clusters and performed PSA-search with OT, assuming that the maximum difference between T cell clusters was equal to the maximum distance between sub-clusters of loci (**Figure 5A, lower**). We repeated the sub-clustering 100 times for each super-cluster with different initializations.

Super-clusters differed by their ability to encode T cell localization as measured by PSA-loss (**Figure 5B**). Super-cluster 8 had the lowest average PSA-loss and was the only cluster that significantly enriched enhancer sequences (distal and proximal), but not promoter-like sequences. **(Figure 5C).**

Low PSA-loss might reflect clustering of loci with non-specific sequence properties and not phenotype specific coherence. We statistically evaluated phenotype specificity as follows. We perturbed the edges in the input T cell neighborhood graph (**Methods**), performed PSA-search with the corrupted graph in the same sequence clusters as the original code, and compared the PSA-loss of this code with that of original. Although the top codes differed a small amount in their PSA-loss, they differed substantially in their phenotype specificity (**Figure 4E**). The code with lowest PSA-loss was also highly phenotype specific (p<0.001 with respect to corrupted inputs).

We hypothesized that (TF) motif enrichment drives the code, given the enrichment of enhancer sequences in cluster 8 and the neural PSA-search results in PBMCs. We developed an approach that attributes the code to driver TFs by evaluating whether the patterns in enrichment of their TF motifs was specific to the phenotype (**Supplementary Figure 7A**).

We first compute TF motif enrichment with HOMER, which controls GC/2-mer and 3-mer frequencies for each locus cluster (Heinz et al., 2010). Next, we identify TFs whose motif enrichment is phenotypically consistent (i.e., locus clusters assigned to similar neighborhoods have similar fold enrichment of TF motifs). For 50/417 TFs, the enrichment of their motifs in the code was phenotypically consistent: similar neighborhoods were assigned genomic loci clusters with similar fold enrichment of TF motifs (**Figure 5F, histogram of consistency p-values**). This is expected for any code with low PSA-loss, since similar loci clusters have similar sequence features, and sequence features should capture sequence motif composition. Next, we define driver TFs as those whose consistency is phenotype-specific. We quantify this by corrupting the phenotype graph and measuring the change in consistency; a large decrease in the consistency of TF motif enrichment with a corrupted input a TF motif globally driving coherence of code with the input phenotype.

The optimal code was driven by TFs fundamental to CD8+ T cell fate in tumors (**Figure 5G, top smoothness specificity)**. The motifs whose consistency decreased the most with a corrupted phenotype were TBX21 (the gene encoding TBET, driver of effector cell differentiation), PLAGL1 (a follicular helper like and stem cell marker), EOMES (a key regulator of memory cell differentiation), STAT4 (a key signaling driver of T cell differentiation), and TGIF2 (an effector of the TGF-beta signaling pathway fundamental in tumors).

We identified the T cell neighborhood clusters that had been assigned loci clusters that strongly enriched these driver TF motifs (FDR < 1e-5) and annotated them with cell type composition. (**Figure 5H and Supplementary Figure 5A)**. TGIF2 motifs were strongly enriched ubiquitously, except in neighborhoods containing B cells and plasma cells. PLAGL1 – typically associated with T cells near B cell follicles (Araki et al., 2009), was strongly enriched in loci clusters assigned to neighborhoods proximal to B cells and plasma cells. TBET (encoded by the TBX21 gene) motifs were enriched most strongly in loci clusters assigned to perivascular, granulocyte and dendritic cell containing neighborhoods, suggesting, consistent with previous studies (Tooley et al., 2022), that these could be genetically encoded sites of effector cell differentiation. Loci clusters strongly enriched for STAT4 motifs were assigned to neighborhoods containing tumor cells.

We investigated colocalization of TSS with loci clusters in the code, which highlighted a single gene. There were only three genes that had a TSS within 50kb of at least four of the ten loci clusters enriching these smooth motifs, and only ZBTB20 had a TSS within 50kb of a locus in five or more clusters (**Figure 5I)**. Our data therefore suggest that spatial gene regulation in in colorectal cancer CD8+ T cells might converge upon ZBTB20. Consistent with this hypothesis, ZBTB20 was shown to be a pivotal regulator of CD8+ T cell metabolism and effector differentiation in cancers (Sun et al., 2020).

We further investigated the loci cluster assigned to tumor cell neighborhoods, which was the only cluster strongly enriched in STAT4 motifs and was also enriched in TGIF2 motifs. We found genes with TSS within 10-30kb of the loci in this cluster. These are therefore genes putatively regulated by TGF-beta and STAT4 in CD8+ T cells, specifically in CD8+ T cells within tumor parenchyma. Performing gene set enrichment analyses, these twenty genes significantly enriched for hematopoietic genes including the core CD8+ T cell transcription factor RUNX1, but also NK-cytotoxicity receptors KLRC1 and KLRC4 (**Figure 5J**). A recent study confirmed that IL-12 – activator of STAT4 signaling - and TGF beta in combination induce KLRC1 expression in colorectal cancer CD8+ T cells (Fesneau et al., 2024) within tumor parenchyma, validating this de novo prediction.

Taken together, these data indicate that the microenvironmental gene regulation of CD8+ T cells is PSA-sufficient, and enable discovering genetic regulators of spatial gene expression from a small number of samples without genotype-phenotype covariation or gene expression data.

The analysis of driver TFs further suggests a model linking the sequence patterns of the genome to T cell localization in the tumor microenvironment. We propose that small, continuous changes in the neighborhood composition of T cells (**Figure 5K, top, colored dots surrounding black cell in center**) are genetically specified **(Figure 5K, black lines)** by distinct subsets of enhancers. These subsets are in turn defined by correspondingly smooth changes in the binding of fate-determining TFs. (**Figure 5K, lower, colored lines indicating TF binding locations)**.

## Discussion

In this work, we introduced a principle to formally relate the global structure of phenotypes and the sequences in the genome that define them: Phenotype Sequence Alignment Sufficiency. A single approach – searching for structure preserving maps between phenotypes and the genome sequence – recovered established genotype-phenotype associations across scales without genotype-phenotype covariation. This framework rediscovered the genetic code for proteins, revealed an evolutionarily conserved structural correspondence between T cell transcriptional diversity and TCF7 motif diversity, correctly enriched for population and functional genomics associations in the context of immune cells and brain anatomy, and revealed sequence determinants of spatial gene regulation in cancer. Thus, structural correspondence is a fundamental feature of how phenotypic structure is shaped by the genome sequence.

Our framework generates mechanistic hypotheses in experimentally challenging contexts by using phenotypic complexity to constrain genetic associations. PSA-search predicts genomic loci associated with in-silico alterations of phenotypic units, from even a single patient’s data. By intersecting these assignments with genomic annotations (TF binding, GWAS summary statistics, perturbations), the output of PSA-search provides hypotheses for which motifs, loci, and genes influence a specific phenotypic unit. Since PSA-search derives codes from the global distribution of motifs across the genome, it can suggest loci contributing to a phenotype even when genomic data is not observed. For example, the codes we found predicted transcription factors unexpectedly regulating cell types, a role for interferon gamma upstream of amyloid beta plaque clearance, and targets of spatial gene regulation in colorectal cancer.

Our results suggest an explanation for PSA-sufficiency. The codes were consistently driven by sequence features corresponding to transcription factor binding sites, suggesting a property of the regulatory genome. We hypothesize that the distribution of TF motifs in the genome is sufficiently expressive in that it contains an “isomorphic” copy of cellular phenotypes within it. However, that the codes were specific (i.e., corrupted phenotypes had worse PSA-loss) suggests that this distribution is not so expressive that it can represent arbitrary phenotypes. We propose that this constrained expressivity underlies PSA-sufficiency.

Our hypothesized mechanism for PSA-sufficiency suggests biological contexts where PSA-search may produce meaningful genetic hypotheses. Specifically, our results suggest PSA-sufficiency holds when fine-grained variation in cellular context is genetically encoded as fine-grained variation in motif composition of regulatory sequences. Such genetic encoding would in turn be expected when fine-grained diversity of cellular context (which could include extracellular surrounding, developmental history or, state) translates to fine-grained, evolutionarily selectable, diversity of cellular output (for example, signaling or differentiation output). Immune cells are known to exhibit this continuous diversity of function (Nathan et al., 2022), which is consistent with our positive results in immune cells. If, instead, phenotypic data consists of a large number of highly separated clusters, their phenotypic similarities between them might not be genetically encoded in this way.

PSA-sufficiency is a framework in which to establish the experimental and computational priors required to recover causal associations across phenotypes without covariation. The parameterization of PSA-search specifies a prior on the structure of the code, analogous to the choice of nonoverlapping triplets in the context of proteins. Assuming the essential structure of codes does not change, applying PSA-search as established in homeostatic contexts (where data is more abundant) provides a strategy to discovery of genetic mechanisms in disease contexts (where data is more sparse).

Establishing the right priors for PSA-search requires ground-truth data. This includes mapping of genetic effects on tissue and cellular structures, as well as causal effects of genetic perturbations. One source of such data is through genetically diverse model organism populations such as in (Svenson et al., 2012)). Our results on the conservation of structural correspondences suggests that priors established here might be applicable to the human context. In addition, perturbation studies of non-coding loci to measure emergent effects on cellular diversity (such as in (Baglaenko et al., 2025)) will provide benchmark datasets with causal information. The concrete task is to find appropriate similarity metrics, code-formulation, optimization strategy, and biological interpretation that the codes found by PSA-search recapitulate these ground truth data.

PSA-sufficiency advances, and provides an application for, current efforts towards virtual cells, tissues and organisms (Rood et al., 2024). As an “upstream” ingredient of such efforts, genomic predictions from PSA can be used directly to predict perturbation effects (as we validated with CRISPR screen hits), or fused with other prediction models. In this context, PSA is an inductive bias on how models should integrate sequence, which can be explicitly incorporated in loss functions to enable generalizing across systems and scales. Downstream of virtual cell efforts, PSA-sufficiency provides a starting point for consolidating genetic prediction models into broad and interpretable theoretical biological principles.

Our results suggest an additional, systems-level mechanism for the conservation of T-cell transcriptional diversity across species. Assuming the global statistical distribution of TCF7 motifs is slow to change over evolutionary time, and because structural correspondence between its shape and T-cell diversity is conserved, T cell diversity is itself buffered against rapid evolutionary change.

Taken together, our results establish a formal principle connecting genotype and phenotype that enables mechanistic discovery for phenotypes across scales, as well as provides a paradigm towards understanding how the genome sequence delimits the phenotypes an organism can produce.

## Acknowledgements

We would like to acknowledge Aviv Regev and Adam Rubin for helpful comments. Funding for this work was in part by the Eric and Wendy Schmidt Center at the Broad Institute.

## Methods

### PSA search with proteins

The E.coli K12 genome was downloaded from the UCSC genome browser. The recA sequence was obtained from UniProt.

Six frames of codons were constructed (starting at positions 0,1 and 2 in both forward and reverse directions). For each frame, and each of the 64 possible nucleotide triplets, a binary vector of positions whose codon beginning at that position was equal to that nucleotide triplet was constructed. For each amino acid in recA, a binary vector of positions that were equal to that amino acid was constructed.

We convolved the binary amino acid vector with each frame and nucleotide triplet’s binary vector, and selected the amino acid, nucleotide triplet and starting position in the genome with the highest match between the amino acid and nucleotide triplet binary vector. We ranked amino acid and nucleotide triplet and position tuples by their match, i.e. alignment quality. We selected the amino acid, nucleotide and position tuple with the optimal alignment quality, and evaluated whether the identified position corresponded to the correct coding sequence of recA in the genome.

We repeated this process for all substrings of recA varying the length, and computed for which proportion of such substrings this optimal code resolved the coding sequence of recA.

Overlapping codes were evaluated as follows. We assigned each of 64 nucleotide triplets to isoleucine, and then counted the matches between the isoleucine binary vector and the respective nucleotide triplet at each position in the *E. coli* genome. Instead of the approach above, we allowed for matches occurring in any of the three frames.

### Sequence features for enhancer sequences [hg38/mm10 ENCODE]

The hg38 and mm10 ENCODE cCRE datasets were downloaded from the UCSC genome browser, encodeCcreCombined.bb, and filetered to include cCREs with proximal and distal enhancer-like signatures were selected. Spectral features for the ENCODE enhancers were obtained as follows.

First, enhancers were binned into genomic blocks, as follows. A graph was constructed, with an edge between two enhancers if the end of one was within 2000bp of the start of another. Connected components of this graph were extracted, and enhancer blocks were obtained as the start and end points of each connected component.

Next, the 1000bp sequence centered at the mean point of each block was extracted, and counts of 1,2,3,4,5 and 6-mers were computed. k-mers containing Ns were removed and counts were z-normalized. PCA was performed and 3050 components were selected as the elbow point.

Spectral embeddings were computed as follows. A 10-nearest-neighbor graph was computed using the cosine metric, with weights given by distances. A symmetrized, unnormalized graph Laplacian was computed and eigenvectors of this graph were obtained; these are the spectral embeddings.

### Evaluating features for enhancer sequences with evolutionary linkage

The 470-species multiz alignment scores for hg38 were downloaded. For each enhancer bin as above, its alignment score with a given species was set as the maximum alignment score with the alignment blocks intersecting it. Pairwise euclidean distances between alignment scores relative to species (with duplicate assemblies removed) were computed between 100 randomly sampled enhancers; this process was repeated 100 times. The correlation between evolutionary pairwise distances and pairwise Euclidean distances between the spectral features, as well as between Euclidean distances between k-mer frequencies, were computed.

### PSA search with Allen brain atlas

#### Preparing Allen atlas data

The Allen human brain microarray atlas was downloaded from (Hawrylycz et al., 2012). Expression matrices were averaged per region over multiple the multiple samples for each patient’s regions. Pairwise distances were computed and the cosine distance, as the metric with the highest correlation between shared regions across individuals, was selected for further analysis. Patient 2001 was selected as the reference patient.

#### Solving PSA with brute-force on Allen atlas

We describe the approach for computing the PSA-loss given a reference region and a seed enhancer assigned to that region. First, a vector of cosine distances in gene expression space from that region to other brain regions is computed. Next, a vector of sequence distances from the enhancer to other enhancers is constructed. This vector is multiplied by scale factors between 1 and 10 with a spacing of 0.5. For each such scale factor, the best matching between the vector phenotypic distances within the scaled vector of sequence distances is computed; the maximum difference across regions between the phenotypic distance from the reference region and the matched sequence distance is computed. The PSA-loss for that enhancer/reference region pair is the lowest maximum loss across scale factors.

We repeated this optimization to compute alignment loss relative to the three reference regions. These were the scores used in the downstream evaluation.

#### Evaluating Allen atlas codes

The twenty enhancers with lowest alignment loss for each of the reference regions were selected. For each region in the other patient with measurements from both hemispheres (Patient 2002), we computed the alignment loss at each of these top enhancers. We computed the correlation between the alignment loss for each region and between the alignment loss for the reference regions in Patient 2001. The alignment loss at each of these enhancers was computed by averaging the discrepancy over regions, instead of taking the maximum since the exact collection of regions is different between patients.

Summary statistics for the overall brain-morphology GWAS hits were obtained from (van der Meer et al., 2020). Enhancer positions were lifted to hg19 using the liftOver python package. For each enhancer, the estimated p-value was given by the minimum p-value of the two closest SNPs with estimated effects in the summary statistics.

Binomial tests were conducted at different GWAS p-value thresholds for obtaining two hits amongst the top twenty putamen enhancers relative to the whole set of enhancers. Enhancers were mapped to genes with gProfiler (Raudvere et al., 2019). p-values relative to Gene Ontology terms were computed using the frequency of enhancers mapped to genes contained in each term that have a GWAS p value < 1e-12 as hits. Only GO terms containing genes to which a top twenty enhancer were evaluated.

Empirical null distributions controlling for marginal sequence distributions were computed by clustering enhancers with minibatch k-means with 10, 100, 500, 1000 and 5000 clusters, and sampling twenty enhancers from these clusters while preserving the marginal distribution across clusters observed in the top twenty hits.

Empirical null distributions controlling for GC content were computed by binning enhancers and sampling random loci of the same size as each bin, using 1, 5, 10, 15 and 20 bins. p-values were obtained as the frequency 2 or more enhancers with GWAS p-values < 1e-10 were observed.

Empirical null distributions controlling for sequence similarity to the seed enhancer were computed by binning enhancers by sequence similarity to the seed enhancer and sampling random loci of the same size as each bin, varying the number of bins. p-values were obtained as the frequency 2 or more enhancers with GWAS p-values < 1e-10 were observed.

Gene expression was averaged over microarray probes per gene, and z-normalized across regions. For each enhancer in the alignment, z-normalized expression was averaged across the genes to which that enhancer was mapped (by gProfiler, as above), yielding a region x region matrix, where rows correspond to regions assigned to enhancers and columns correspond to regions measured for gene expression. The median along the diagonal was computed, and the average difference was compared to null alignments obtained by sampling according to the binned sequence distances from the hit enhancer, binning at different resolutions; bin 1 is a fully random assignment.

Brain regions from the biobank MRI volume GWAS of (Zhao et al., 2019) were mapped to brain regions in the Allen atlas (**Supplementary Table 1**). 1000bp bins centered at each enhancer center were extracted, the maximum absolute beta for each bin was computed, restricting to SNPs with nominally significant p-values < 0.001. Null distributions were computed as above.

#### Preparing single-cell gene expression data for PSA search

For the PBMC and Tabula Sapiens datasets, the SEACells package (Persad et al., 2023) was utilized to compute metacells.

#### Tabula Sapiens

The Tabula Sapiens dataset was downloaded from cellxgene (CZI Cell Science Program et al., 2025). The log1p transformed gene expression was divided by 1000 and 100000 randomly selected cells generated by “10X” method were selected. 400 metacells (with 15 waypoint eigenvalues) were obtained using 200 principal components using SEACells with default parameters, and the gene expression vectors for each metacell were obtained by averaging cells by their metacell assignments.

Datasets for neural PSA input were obtained as follows. 50 metacells were randomly selected as validation data. The input dataset for PSA consisted of gene expression vectors averaged over cells assigned to each metacells, whose 3 nearest neighbors did not intersect the validation metacells.

#### PBMC atlas

Log1p transformed gene expression vectors from cells at time 0 from donors 1,3, and 5 were obtained from the dataset of (Hao et al., 2021), and divided by 1000. Gene expression values were projected to 200 principal components. The SEACells algorithm was applied with 15 waypoint eigenvalues to obtain 200 metacells. The log transformed gene expression vectors for the original cells were grouped by metacell assignments to obtain the gene expression vectors for each metacell.

Datasets for input to neural PSA search were obtained as follows. First, 20 metacells were randomly selected as validation cells. Next, metacells whose 3 nearest neighbors did not include any of the validation cells were selected as training cells. The input to PSA was the training data matrix, consisting of the gene expression vectors for each training metacell.

The same procedure (metacell construction, aggregation, training data creation) was applied to the cells of donors 2,4 and 7 to produce a biological replicate input for neural PSA.

### Optimal transport PSA codes for Tabula Sapiens

#### Computing PSA codes

Spectral embeddings (with 300 eigenvectors) of ENCODE clusters as described above were clustered twenty times into 1000 clusters with the minibatch k-means algorithm. For each of these cluster sets, the manifold distance was computed (shortest path between cluster centroids with respect to a 5-nearest neighbor graph). The phenotype distance between two metacells was the manifold distance (shortest path between two metacells using a 20-nearest neighbor graph computed from cosine distance).

Semirelaxed Gromov-Wasserstein was computed using the POT python package with respect to these distance matrices. The default entropic regularization factor 0.001 was used, and varying this between 0.0001 and 0.1 did not change assignments.

For each cell type and metacell, the fraction of cells of that type assigned to that metacell was computed. The enhancer cluster for each cell type was the enhancer cluster assigned to the metacell with the highest fraction of cells of that type assigned to it.

For each metacell, the squared distances between its distances and the distances between the enhancer clusters assigned in the coupling was its loss; for each cell type the loss for the metacell to which it was assigned was considered its loss.

#### Chip-seq enrichment analysis

Chip-seq peaks were downloaded according to **Supplementary Table 2** and intersected with the enhancers. The set of significant lineage primed enhancers was obtained from (Weng et al., 2024) and evaluated similarly.

For each enhancer cluster, Fisher exact tests were conducted to evaluate the enrichment of enhancers intersecting CHIPseq peaks relative to the overall set of ENCODE enhancers.

A GC-matched enrichment p-value was obtained by taking 2500 random samples with replacement from the overall set of enhancers with the same size and GC content distribution (binned into 5 bins) as each enhancer cluster, and computing the proportion of times a greater or equal number of enhancers intersected the target peak set than the locus cluster.

#### Evaluating PSA for eQTL effects

The log transformed PBMC single-cell gene expression data was obtained from the Azimuth portal, and converted to AnnData using seuratdisk package. The mean gene expression vectors were computed for each of the “celltype.l2” cell types. The single-cell PBMC eQTL data was obtained from (Yazar et al., 2022)

The cell types from the gene expression dataset were mapped to the cell types from the eQTL dataset as follows:

- ‘Classic Monocyte’: ‘CD14 Mono’,
- ‘Non-classic Monocyte’: ‘CD16 Mono’,
- ‘Naïve/Immature B Cell’:‘B naive’,
- ‘Memory B Cell’:‘B memory’,
- ‘Plasma Cell’:‘Plasmablast’,
- ‘CD8 Effector memory’:‘CD8 TEM’,
- ‘CD4 Effector memory/TEMRA’:‘CD4 TEM’,
- ‘Natural Killer Cell’:‘NK’

5000 random gene-RSID pairs were extracted, and those that had reported Spearman correlations with all eight of the above cell types were selected. This produced, for each cell type, a vector of Spearman correlations. The genetic association distance between each pair of cell types was the L1 distance between their vectors of Spearman correlations. The same analysis was repeated within 2MB of each TSS grouped by the number of cell types with significant eQTLs.

The phenotypic distance between each pair of cell types was the cosine distance between their mean gene expression vectors.

A permutation test was conducted as follows.

- The test statistic was the average Spearman correlation between the pairwise distances. Each pair of cell types was counted only once, and the zero distances corresponding to a pair of identical cell types were not included.
- The null distribution for the test statistic was generated by permuting the matching between gene expression vectors and correlation vectors. This null distribution accounts for the presence of a given cell type in multiple pairs. 10000 permutations were utilized.
- The p-value was computed as 1- P, where P is the proportion of times that the null distribution value was strictly less than the test statistic. This is a one-sided test.

Similar tests were performed for the other distance metrics reported in **Supplementary Figure 2A.**

The database of EMBL GWAS Catalog RSIDs for the CHM13v2.0 assembly were downloaded from the UCSC genome browser (’chm13v2.0_GWASv1.0rsids_e100_r2022-03-08.vcf’).

The gene-RSID pairs were selected whose:

- RSID was in the GWAS database
- Had reported FDRs for association all eight of the above cell type
- Were strongly associated (FDR< 0.01) with at least three of cell types.

After these filtering steps, there were 46 RSIDs, and none corresponded to different variants at the same genomic position.

The phenotypic association for an RSID was the set of cell types to which it was significantly associated. The phenotypic association distance between a pair of RSIDs was the Jaccard distance between their sets of associated cell types, i.e. the proportion of cell types to which one was associated and other not.

The syntactic embedding for an RSID was considered to be the embedding of the 2000-mer starting closest to it and contains it. That is, the syntactic embedding of position i is the language model embedding of the 2000-mer beginning at position 1000*floor(i/1000). The syntactic distance between a pair of RSIDs was the cosine distance between their embeddings.

The phenotypic association distance between a pair of RSIDs was the Jaccard distance between their phenotypic associations (i.e., the number of cell types where one was significantly associated but the other not).

A k-nearest neighbor graph between RSIDs was computed using the scikit-learn package at each value of k between 5 and 40 using the syntactic distances. For each RSID, the average pairwise genetic association distance between their k neighbors as defined by this graph were computed, and this is the data of **Figure 3B**.

A permutation test was performed as follows:

- The test statistic was the mean across RSIDs of the average pairwise genetic associations amongst their 5 nearest neighbors.
- The null distribution was generated by permuting the assignments of RSIDs to vertices in the 5 nearest neighbor graph (generating a random collection of not necessarily syntactically near neighbors for each vertex). 10000 samples from this null distribution were generated.
- The p-value was computed as 1-P where P is the proportion of null values that were strictly greater than the observed test statistic. This is a one-sided test.

A similar analysis was performed, except removing RSIDs that were within a genomic distance of 5000 of each other; this is the permutation test in **Supplementary Figure 1C**.

The test in Supplementary Figure 2B is identical, except it uses syntactic embeddings of 512mers from the DNABERT pretrained model (Huix, 2022).

### Quantitative evaluation of neural PSA search codes

#### PSA-search with neural networks

Neural PSA was applied as described in **Supplementary Note 1**.

The precomputed syntactic embeddings, described in the sequence data preparation section of **Methods**, were divided by 50 to normalize their scale. The GPU-FAISS package was utilized to compute a 20-nearest neighbor graph with cosine distances between the 3.5M syntactic embeddings for computing the syntactic consistency loss.

All models were implemented in PyTorch; the Adam optimizer with learning rate 0.0001 was utilized. For all models, the weights in the loss term for the consistency with phenotypic geometry, consistency with genomic syntax and sparsity were 1,1 and 0.1 respectively.

We provide here the dataset specific details for the implementation.

#### Neural PSA search on PBMC dataset

For the dataset corresponding to cells from patients 1,3 and 5:

- The phenotype embedding and syntax embedding networks were linear maps with latent dimension 40. Both networks were initialized with PCA.
- The sequence batch size was 100000; all metacells were considered simultaneously.

The model was selected after 76 epochs through the genome. This was the point when the training phenotype consistency loss started increasing and the sparsity and syntax loss decreased sharply (**Figure S5**, **upper row**).

For the dataset corresponding to cells from patients 2,4 and 7, we applied PSA with two conditions. The first was the same conditions as for patients 1,3, and 5 described above and the model was selected after 124 epochs as per the stopping criteria; these parameters resulted in a code that did not enrich for eQTLs.

We trained another model increasing the dimension of the latent space:

- The phenotype embedding and syntax embedding networks were linear maps with latent dimension 80. Both networks were initialized with PCA.
- The sequence batch size was 60000. This was the largest batch size that fit on a single GPU with latent dimension size 80 without batching metacells.

The model was selected after 83 epochs through the genome. This was the point when the training phenotype consistency loss started increasing and the sparsity and syntax loss decreased sharply (**Figure S5**, **second row from top**).

#### Neural PSA search on Tabula Sapiens dataset

For the metacells from Tabula Sapiens dataset constructed as detailed above:

- The phenotype embedding and syntax embedding networks were linear maps with latent dimension 40. Both networks were initialized with PCA.
- The sequence batch size was 100000; all metacells were considered simultaneously.

The model was selected after 24 epochs through the genome. This was the point when the validation phenotype consistency loss started increasing and the sparsity and syntax loss decreased sharply (**Figure S5**, **third row from top**).

### Evaluation of cis-eQTL prediction

#### Data preparation

##### Classification task, GWAS SNPs

RSIDs in the eQTL dataset that were also in the T2T-GWAS SNPs were selected. The genome was binned into 1000 base pair loci. A locus was determined to be significantly associated with expression in a cell type if it contained a GWAS SNP within it that was significantly associated (FDR < 0.1 in the original eQTL dataset) with expression in that cell type. Only loci containing GWAS RSIDs evaluated in the eQTL dataset were considered.

The gene-cell type pairs were selected that had at least one significantly associated locus and one not significant locus. In addition, genes with zero expression in all of the cell types in the PBMC dataset were removed.

This resulted in a total of 7836 unique loci, 149 cell-type gene pairs across 95 unique genes.

Syntactic embeddings were extracted for each locus, as the embedding of the 2000-mer beginning at the same genomic position.

##### Classification task, all SNPs

RSIDs in the eQTL dataset that were also in the T2T dbSNP database were selected. The genome was binned into 1000 base pair loci. A locus was determined to be significantly associated with expression in a cell type if it contained any SNP within it that was significantly associated (FDR < 0.1 in the original eQTL dataset) with expression of the given gene in the given cell type.

The gene-cell type pairs were selected that had at least one significantly associated locus and one not significant locus. In addition, genes with zero expression in all of the cell types in the PBMC dataset were removed. We evaluated significance varying the minimal number of loci with available statistics (**Supplementary Figure 3H).**

Syntactic embeddings were extracted for each locus, as the embedding of the 2000-mer beginning at the same genomic position.

##### Effect size task, all SNPs

RSIDs in the eQTL dataset that were also in the T2T dbSNP databse were selected. The genome was binned into 1000 base pair loci. The maximum absolute effect size for SNPs within a locus was taken to be the effect size for that locus.

The gene-cell type pairs were selected that had at least one significantly associated locus and one not significant locus. In addition, genes with zero expression in all of the cell types in the PBMC dataset were removed. We evaluated significance varying the minimal number of loci with available statistics (**Supplementary Figure 3J-K).**

Syntactic embeddings were extracted for each locus, as the embedding of the 2000-mer beginning at the same genomic position.

#### Computation of QT-scores from PSA

The genetic code T(x,s) learned by PSA was used to compute absolute QT scores for each cell-type c, gene g and locus s. Specifically, the QT-score was |T(**x_c_**+ **δ_g_**,**s**) – T(**x_c_**,**s**)|, where:

- **x_c_** is the mean gene expression vector for the cell type,
- **δ_g_** is a shift of 0.0001 in the dimension corresponding to the gene g
- **s** is the syntactic embedding of the locus **s**

These QT-scores were computed only for locus evaluated for eQTLs as defined above. These QT-scores thus provided a score for each cell-type, gene and locus.

#### Evaluation metrics

##### Classification task

For each cell-type gene pair, the AUC for using the QT-scores for predicting loci significantly associated with gene expression for that cell type were computed using the scikit-learn function roc_auc_score for each cell-type gene pair.

##### Effect sizes

The average z-score of the QT-scores (relative to other loci for each gene/cell type pair) was computed for the 3 loci with highest effect size.

#### Permutation tests for GWAS SNPs

For each of the following hypothesis tests, the test statistic was computed as the mean (across genes) of the mean AUC (across cell types) at predicting eQTLs for that gene. The null distributions for the test were generated using the following alternatives to the QT-score as below (see above for notation).

1. The gene permutation null was generated by randomly sampling permutations q(g) (assigning each of the 95 genes to another not necessarily distinct one of the 95 genes), and computing QT-scores as |T(x_c_+ δ_q(g)_,s) – T(x_c_,s)|
2. The cell type permutation null was generated by randomly sampling permutations q(c) (assigning each of 8 cell types to another not necessarily distinct cell type), and computing QT-scores as |T(x_q(c)_+ δ_g_,s) – T(x_q(c)_,s)|
3. The locus permutation null was generated by randomly sampling permutations q(s) (assigning each of the 7836 unique loci to another not necessarily distinct locus). and computing QT-scores as |T(x_c_+ δ_g_,q(s)) – T(x_c_,q(s))|
4. The cell-type gene expression preserving permutation null was generated by first grouping the 95 genes by the set of cell types in which their average log expression was greater than 0.02, and only sampling permutations q(g) mapping genes to others expressed in the same set of cell types.
5. The random initialization null was obtained by randomly initializing the weights of PSA and computing the QT-scores as above.
6. A total of 72 grouped cell type permutation was obtained by restricting permutations to preserve monocytes (2 cell types), B cells (3 cell types) and T cells (2 cell types). NK cells were not permuted.
7. The resolution investigation null distributions were obtained as follows.

a. For each resolution distance, a graph was constructed in which two loci were connected by an edge if they were on the same resolution and within the same linear distance.
b. This graph was partitioned into connected components.
c. Permutations q(s) were sampled in which a locus could only be permuted with one in the same connected components.

For each of these null distributions, the p-value was 1-P, where P is the proportion of times that the null AUC was lower than the test statistic. 2000 sampled permutations were utilized in each case.

##### Permutation tests for all loci

Permutations were carried out in the same way as above. However, the groups for the gene permutations were determined by constructing a 1-nearest neighbor graph of genes by their correlation across expression and selecting the connected components of this graph.

#### Baseline predictors for eQTL classification task

##### GC content and distance to TSS

For each locus, the average proportion of bases in the T2T assembly that were G or C was computed. For each gene and locus, the minimum distance to a transcription start site in the RefSeq database was computed.

##### ENCODE cCREs

The hg38 ENCODE cCRE annotations were obtained from the UCSC genome browser. For each locus in the dataset, it was considered proximal to a given CRE mark at a given distance if it contained an RSID that was within that distance to a CRE mark from the ENCODE annotations in hg38. cCRE proximity was computed at distances of 1-11 kb in 1kb intervals and used as the predictor. The data in Figure 2B uses the best performing distance for each cCRE mark and **Supplementary Figure 2A** contains all the predictors.

##### Gene expression classifier

For each gene, the syntactic features of the 50 loci either side of the TSS was extracted. This covered, due to the spacing 1000bp apart as described above a total length of 50kb either side of the gene. A linear model was trained to predict the z-normalized expression (across the eight cell types) for the gene, using these features as input (averaging predictions across in the 100kb window). The model achieved an R^2 of 0.02 across validation set genes. For each locus and cell type, the predicted expression contribution of that locus’ features by the trained model was used as a predictor for effects.

##### eQTL classifier

For all loci containing GWAS SNPs, a classifier was trained to predict whether it contained an eQTL for any gene (FDR < 0.1), holding out all loci evaluated for eQTLs for each gene, one at a time. The prediction of this classifier evaluated for its ability to find eQTLs in the held out gene.

#### Generation and analysis of corrupted PSA input datasets

95 corrupted phenotypic datasets were generated by randomly permuting the entries of the column corresponding to each of the genes evaluated in the GWAS SNP tasks (one at a time).

PSA was applied to each of these with the same initialization and hyperparameters as the original dataset. The QT-scores were computed as above, using the code for each corrupted atlas.

The heatmap in Figure 2E was produced by taking the z-score across corrupted input datasets for eQTL prediction for each gene. The average AUC decrease on the diagonal was statistically compared to off diagonal terms with null distribution obtained by permuting rows.

#### Evaluation of fine-mapped eQTL identification

The dataset of causal probabilities for lead eQTLs (identified using CaVEMAN) was downloaded from the GTEx portal (“GTEx_v8_finemapping_CaVEMaN.txt”). Genomic coordinates were lifted from hg38 to CHM13v2.0 using the liftover python package.

Each locus s was assigned the maximum causal probability (over QTLs contained within it), for each gene and tissue. The following tissues were common between the Tabula Sapiens dataset and GTEx database (excluding blood): ‘Liver’, ‘Lung’, ‘Pancreas’, ‘Prostate’, ‘Spleen’, ‘Uterus’. We selected gene-tissue pairs which had at least three loci with reported causal probabilities, and at least one with causal probability 0.9 or greater and one without. This left a total of 853 unique loci, 119 unique genes across the tissues.

For each of these tissues c, an average gene expression vector x_c_ was obtained as follows. First, single cells were aggregated by their tissue and metacell assignments, yielding a weight for each tissue for each metacell. Then, the weighted means of each metacell’s gene expression were taken as the average tissue gene expression.

The QT-scores for a gene g, tissue c at a locus s were computed from the genetic architecture output from PSA applied to the Tabula Sapiens dataset T(x,s) as |T(x_c_ +δ_g_, s) – T(x_c_,s)| where δ_g_ indicates a shift of 0.0001 in the dimension corresponding to the gene g.

For each gene, tissue pair, the AUC for using the QT-scores to successfully predict loci with causal probability > 0.9 was computed. Note that in this analysis, as with the cis-eQTLs, we only evaluate the loci that are evaluated with each gene.

The AUC was averaged across tissues and genes to compute the test statistic. Tissue, gene, tissue-specific gene, loci and randomly initialized null distributions were generated as with the analysis of PBMC cis-eQTLs, using 2000 permutation samples each.

The p-values were computed as 1-P, where P is the proportion of time that the AUC exceeded that of the null values.

#### CRISPR gene hit prediction

The gene hits were those in the CRISPR activation screen from (Schmidt et al., 2022) for interferon gamma expression in CD8 T cells that had a computed z-score of greater than or equal to the thresholds indicated.

The QT-scores for each gene in CD8+ T cells were computed as |T(x_CD8_ +δ_g_, s) – T(x_CD8_,s)|, where δ_g_ indicates a shift of 0.0001 in the dimension corresponding to the gene g. The QT scores were averaged over the 5 loci centered at each transcription start site in the NCBI RefSeq database for CHM13v2.0 to produce a gene-level QT score.

Gene-level QT-scores for Gene Ontology modules were computed by averaging the rankings of the gene-level QT scores for each module.

The AUC for using the gene-level QT scores to predict the gene hits for IFNG was computed. The GC baseline was computed using the GC content within 1kb of the transcription start site. Logistic regression models with log GC content and QT-scores as predictors were fit using the statsmodels python package and the p-value for the QT-score is displayed.

The GO-term p-value was computed by evaluating the AUC for all the gene-level QT scores for GO terms with at most 100 genes and computing the number of such gene sets with a greater or equal AUC than the interferon gene set.

#### Discrepancy difference on memory vs. effector ATAC-peaks

For each cell type, the metacells with 50% or more cells belonging to that cell type were selected and PCA was performed. Next, the mean for each cell type was increased and decreased by 0.0001 times the first PC, and the discrepancies along genomic 2000-mers were computed as for above analyses.

The dataset of ATACseq peaks was obtained from (Giles et al., 2022) and lifted to T2T with the liftover package. Peaks were assigned to the 2000-mer bins their start was contained in. Memory peaks were those more accessible in in the stem-cell memory CD8+ T cell population relative to CD8+ Effector cells (FDR < 0.0001); effector peaks were those less accessible in this comparison.

A discrepancy difference was computed on each peak by subtracting the discrepancy for decreasing the cell type mean vector by PC1 from the discrepancy for increasing by PC1. A t-test was performed to test differences in discrepancy difference on memory vs. effector peaks. The same analysis was performed but with a shifted peak set obtained by shifting all peaks, this is the data in Figure 4P.

#### Attribution analysis

For each memory peak and 2000-mer used in the computation of discrepancy differences, we generated a corrupted 2000-mer by permuting the nucleotides within sliding 50-mer windows with an overlap of 25 base pairs, and computed the discrepancy difference on the corrupted sequence.

We computed the change in discrepancy difference and performed z-normalization across bins. High attribution bins were determined to be those whose corrupted sequence had a z-score of less than −2.5 and −3 (i.e., the discrepancy difference decreased by more than 2.5/3sd respectively).

For each bin, we computed whether it contained a TCF7 motif. Specifically, we computed the maximum PSSM for the TCF7 motif at positions within that bin using the Biopython motif calculate function and the PWM from JASPAR. A match was one with log odds > 5 relative to uniform.

We computed the number of high-attribution bins that overlapped a TCF7 motif, and permuted the attribution scores across bins to obtain a null distribution to obtain a p-value. We computed a p-value using different numbers of top peaks (by discrepancy difference), and performed computed the same p-value when restricting to sequences with a unique high-attribution region (to rule out biases from motifs being driven by a small number of sequences).

### De novo PSA on colorectal cancer T cell neighborhoods

#### Preprocessing CD8+ T cell neighborhoods

Multiplexed immunofluorescence data was downloaded from (Schürch et al., 2020). For each cell, the neighborhood counts were computed as the counts of each of the 31 cell types within the 20 nearest neighbors to that cell were computed. CD8+ T cells were selected, and their neighborhood counts were clustered using minibatch k-means with 100 clusters. The phenotypic distance was the normalized path distance with respect to the 10-nearest neighbor graph (by cosine distance).

#### De novo PSA-search with optimal transport

Loci from the T2T reference, with GENA features as defined above, were clustered into twenty clusters using minibatch k-means. Each of these twenty clusters were further clustered into 500 subclusters with minibatch k-means, one hundred times with different initial seeds. For each of these cluster sets, the manifold distance between pairs of clusters was used to compute the semirelaxed optimal transport coupling.

#### Statistically testing input phenotype specificity with optimal transport

A corrupted distribution over input phenotypic distance matrices was obtained by randomly selecting a pair of clusters i and j, with centers z_i and z_j respectively, setting the corrupted center of z_i_ as (1-t) * z_i + 0.25tz_j_ for t~ U[0,1], and recomputing the manifold distance. Samples where the distance matrix did not change were discarded. The p-value with respect to this distribution was the fraction of times that these corrupted distance matrices had a lower alignment loss. The optimal coupling was selected as shown in **Figure 4B**.

#### TF binding site enrichment with HOMER

Each locus was mapped to hg38 using the python liftOver package. Clusters with fewer than 100 loci mapped to hg38 were discarded. HOMER findMotifsGenome (Heinz et al., 2010) was used to perform enrichment analyses adjusting for GC and 1/2/3-mer frequencies, with a window size of 1000. The HOMER p-values for each cluster were concatenated and an overall FDR was computed.

#### Motif smoothness computation

The smoothness was measured with the phenotype graph Laplacian. First, the HOMER log motif enrichment for each loci cluster was computed. Each phenotype cluster was assigned the motif enrichment of the loci cluster to which it was assigned in the optimal coupling, giving a vector of enrichments **v.** Next, the Laplacian of the phenotype 10-nearest neighbor graph was computed **L**; **v**^t^**Lv** is the sum of the differences across edges in the graph of the signal **v** and is hence its smoothness. Null distributions were defined by (a) randomly permuting the entries of **v** and (b) randomly permuting the entries of **v**, but ensuring that phenotype clusters assigned to the same loci cluster were consistently permuted. Smooth motifs were considered as those with p<0.99 with respect to both of these null distributions.

A phenotype specific smoothness score was computed by corrupting the input phenotype by generating corrupted input phenotype k-NN graphs as described above, and recomputing the smoothness of **v** with respect to the Laplacian of the corrupted graph.

The loci clusters in the assignment with HOMER FDR<0.0001 for one or more of the top five most specific smooth motifs were selected for further analysis. For each loci cluster, the average cell type count in the phenotype clusters was used to annotate it.

#### Colocalization analyses

For each chromosome, a sliding window of size 50-loci (equivalent to 50kb sequence distance) was used to evaluate colocalization of loci clusters. Genes with a TSS within 50kb of more than five distinct loci clusters were selected.

To find putative genes distally regulated by loci clusters, genes with TSS between 10 and 30kb of the loci cluster assigned to the tumor T cell neighborhood were selected. Geneset enrichment analysis was performed with Enrichr (Kuleshov et al., 2016).

#### Declaration of generative AI and AI-assisted technologies in the manuscript preparation process

During the preparation of this work the authors used Gemini, Claude and ChatGPT LLMs to evaluate text for clarity. The authors reviewed and edited the content as needed and take full responsibility for the content of the published article.

## Supplementary Figure legends

**Supplementary Figure 1**

A. Match proportion with at each position with isoleucine allowing for frame-shifts

B. Best match with isoleucine assignment to each nucleotide triplet across the genome

**Supplementary Figure 2**

A. Log10 spearman rank positive correlation p-value between distances to other regions in across Patient 2001 and Patient 2002 in Allen atlas, averaged across regions for different choices of distance metric.

B. Pearson correlation between feature distance (spectral and kmer) and evolutionary distance for 50 random, nonoverlapping samples of 200 enhancer regions. Evolutionary distance computed as Euclidean distance between alignment scores with 400 other species as in Multiz-470 way alignment with duplicate species removed. p=0.02 with both relative Student’s t and Mann Whitney U tests.

C. Specificity of top putamen seed assignments: Spearman correlation (y-axis) between PSA-loss on top twenty enhancers assigned to each reference region in Patient 2001 (x-axis) and PSA-loss for those same enhancers assigned to other regions in Patient 2002, Labels indicate regions with PSA-loss correlation p<0.05.

D. Each point is an enhancer, values indicate PSA-loss for code when that enhancer is assigned as seed to the reference region indicated on axes.

E. Binomial p-value for enriching GWAS hits with varying p-value hit-thresholds (y-axis) across top N (x-axis) codes

F. −log10 p-value for enrichment of two GWAS hits amongst twenty randomly sampled enhancers controlling for sequence similarity by k-means clustering with 10,100,500,1000 and 5000 clusters (dotted line) and for GC content with 1,5,10,15,20 bins (dashed line).

G. Binomial p-value for enriching GWAS hits amongst enhancers contained in genes contained in GO terms containing genes containing one of the top 20 enhancers.

H. GO terms with binomial p-value > 0.05 from G.

I. Average normalized expression of genes containing enhancers for each region in code (rows) across brain regions (columns); row/column boxes indicate difference between mean and diagonal value in row/column respectively

J. Distribution comparing values on diagonal of Supplementary Figure 1I vs off-diagonal values (i.e. normalized expression of gene containing enhancer assigned to a region within that region vs in other regions)

K. Effect sizes for enhancers assigned to Allen brain region (row) on volumes from Zhang et al Nat Gen. (columns)

L. Enrichment of TF chipseq peaks (columns) in enhancer sets assigned to cell types (rows) with Fisher exact p-values <0.05 (no correction for GC bias).

**Supplementary Figure 3**

A. Frequency of Tabula Sapiens cell type assignments (rows) in each metacell (columns). Only cell types comprising >5% of at least one metacell are shown. Column ordering as in Figure 3A.

B. Enrichment of TF chipseq peaks (columns) in human enhancer sets assigned to cell types (rows) with Fisher exact p-values <0.05 (no correction for GC bias).

C. Enrichment of TF chipseq peaks (columns) in mouse enhancer sets assigned to cell types (rows) with Fisher exact p-values <0.05 (no correction for GC bias).

**Supplementary Figure 4**

A. Euclidean distance between gene expression vectors and L1 (cityblock) distance between vectors of variant effect (correlation) for each pair of cell types

B. Cosine distance between gene expression vectors and cosine distance between vectors of variant effect (correlation) for each pair of cell types

C. Correlation across pairs of cell types between gene expression vectors and distance between variant effects restricted within 2MB cis windows for each gene, grouped by the number of cell types with significant eQTLs reported per gene.

D. Empirical null distribution for average Jaccard similarity between sets of cell types associated with 20-neighbor windows obtained by random permutations of DNABERT syntax feature graphs.

E. Schematic illustrating construction of null distributions evaluating gene and cell type specificity of QT-scores by editing random genes in the correct cell type and correct genes in random cell types respectively

**Supplementary Figure 5**

A. Gene-level AUC with predictors, including proximity to indicated ENCODE cCRE marks at different distances.

B. Establishing kilobase resolution of code’s eQTL prediction with null distributions permuting loci at varying length-scale (2000 samples each). Black dotted lines indicate 95% percentile.

C. Correlation between gene-permutation p-value and z-score for AUC when predicting a gene’s eQTLs with code from phenotype where that gene has been corrupted vs. phenotypes in which other genes have been corrupted.

D. Empirical null distributions for neural PSA code when obtained with different random seed

E. Distribution of gene-level eQTL prediction AUCs with neural PSA code from three different biological donors.

F. Distribution of gene-level eQTL prediction AUCs with neural PSA code from three different biological donors when latent dimension of embedding networks increased to 80.

G. Empirical null distributions for code with biological replicates

H. Ablations of loss terms when performing neural PSA-search.

I. Empirical null distributions for AUCs including all SNPs

J. Empirical null p-values for AUCs as the threshold of loci with statistics for each cell type/ gene is filtered, removing celltype/gene pairs with no significant or all significant loci

K. Number of genes with each threshold in J

L. Empirical null distributions for increase in z-score of SNPs on top three variants with highest effect size

M. Empirical null p-values for AUCs as the threshold for number of evaluated loci with effect sizes reported

N. Number of genes for each threshold in M.

O. Size of gene groups for grouped gene permutations

**Supplementary Figure 6**

A. Log10 t-test p-value discrepancy difference between increasing and decreasing first PC on memory and effector CD8+ peaks as a function of shift per cell type, GC indicates difference for GC% for peaks.

B. Schematic for obtaining attribution scores by permuting in overlapping bins.

C. Log10 permutation p-value for colocalization of high-attribution regions on memory peaks with TCF7 motifs, with thresholds of 2sd and 2.5 sd per sequence as defining high-attribution regions, and across top N peaks (x-axis) by discrepancy difference; restricting to sequences with a unique high-attribution region.

D. Multivariate logistic regression p-values for coefficient of gene-level score from PSA alongside GC content at TSS in predicting gene hits

E. Expression (x-axis) and hit-prediction of AUC for each gene set (restricting to gene sets with at most 100 genes).

F. Distribution of gene level AUCs (averaged over tissues) for predicting fine-mapped QTLs (PIP > 0.5) from other lead eQTLs

G. Empirical null distributions for predicting fine-mapped GTEX eQTLs (PIP >0.5) with PSA-code on Tabula Sapiens input.

**Supplementary Figure 7**

A. Annotations of loci clusters by the average composition of the CD8+ T cell neighborhoods assigned to them by optimal code. Colors match those in Figure 4H-I.

**Supplementary Figure 8**

A. Training curve and early stopping checkpoint selection for code on PBMC dataset

B. Training curve and early stopping checkpoint selection for code on biological replicate PBMC dataset with latent-dimension 40

C. Training curve and early stopping checkpoint selection for code on biological replicate PBMC dataset with latent-dimension 80

D. Training curve and early stopping checkpoint selection for code on Tabula Sapiens

