## Supplementary Figures for "Deriving genetic codes for molecular phenotypes from first principles"

Supplementary Figure 1

**A**

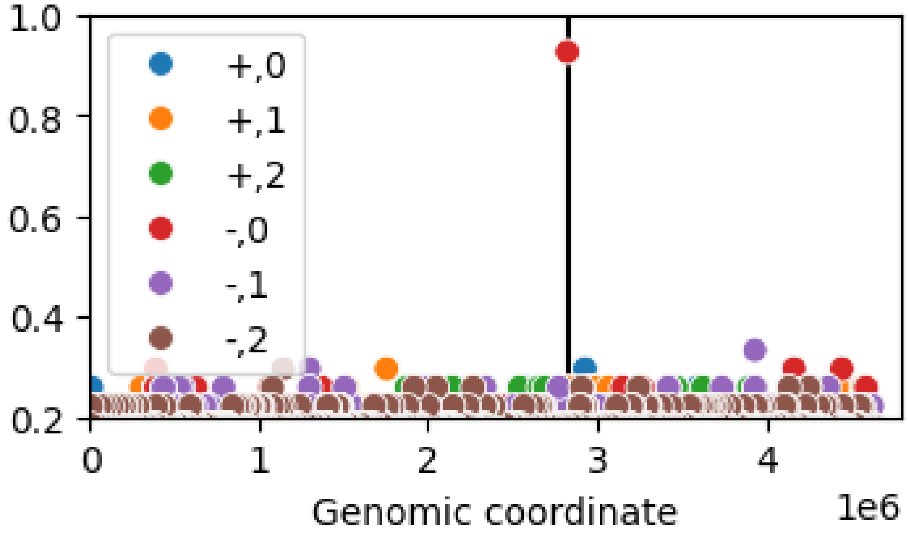


**B**

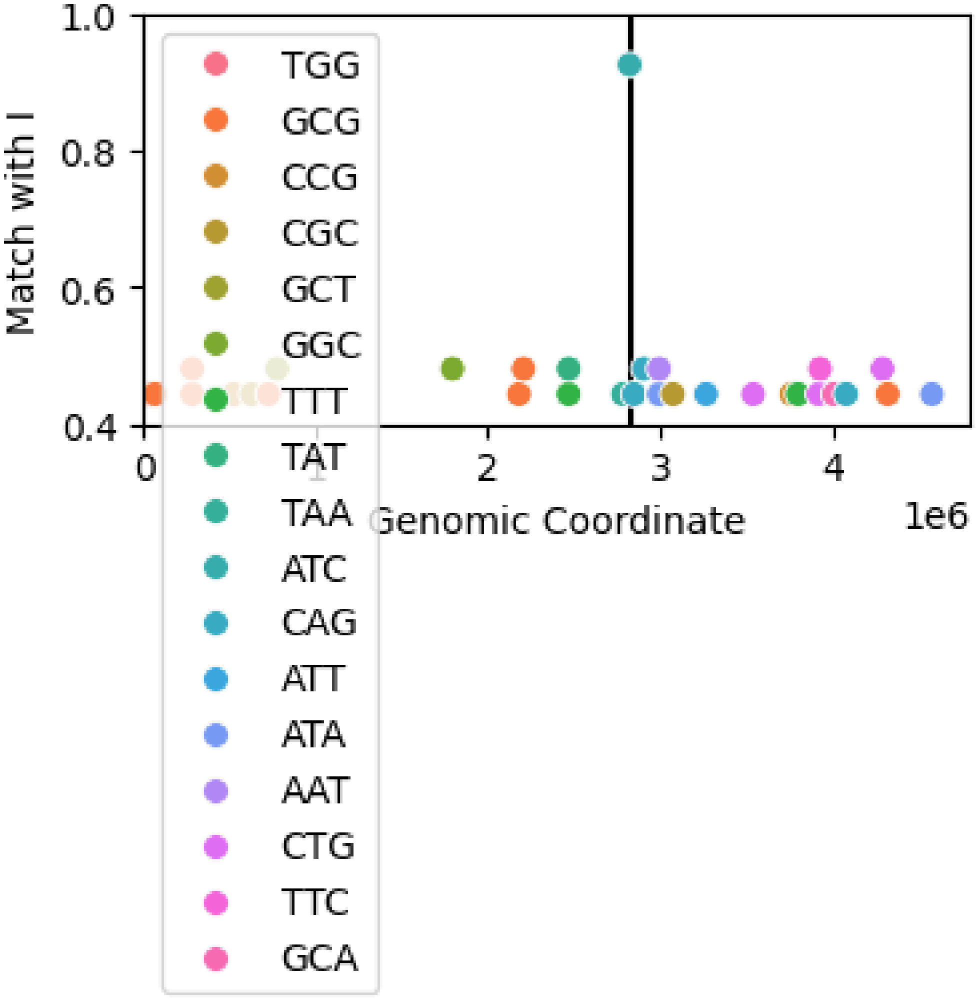

### Supplementary Figure 2

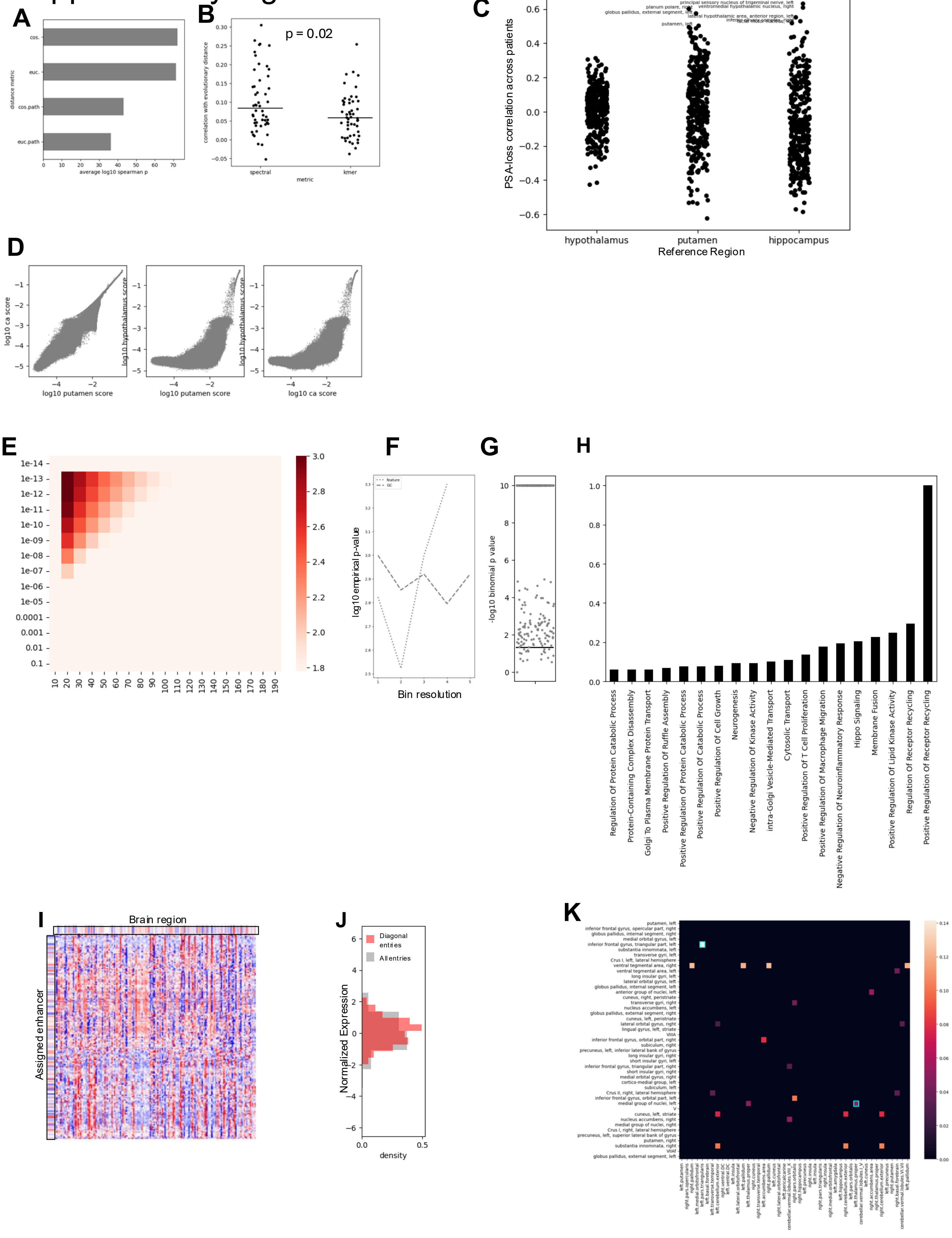

### Supplementary Figure 3

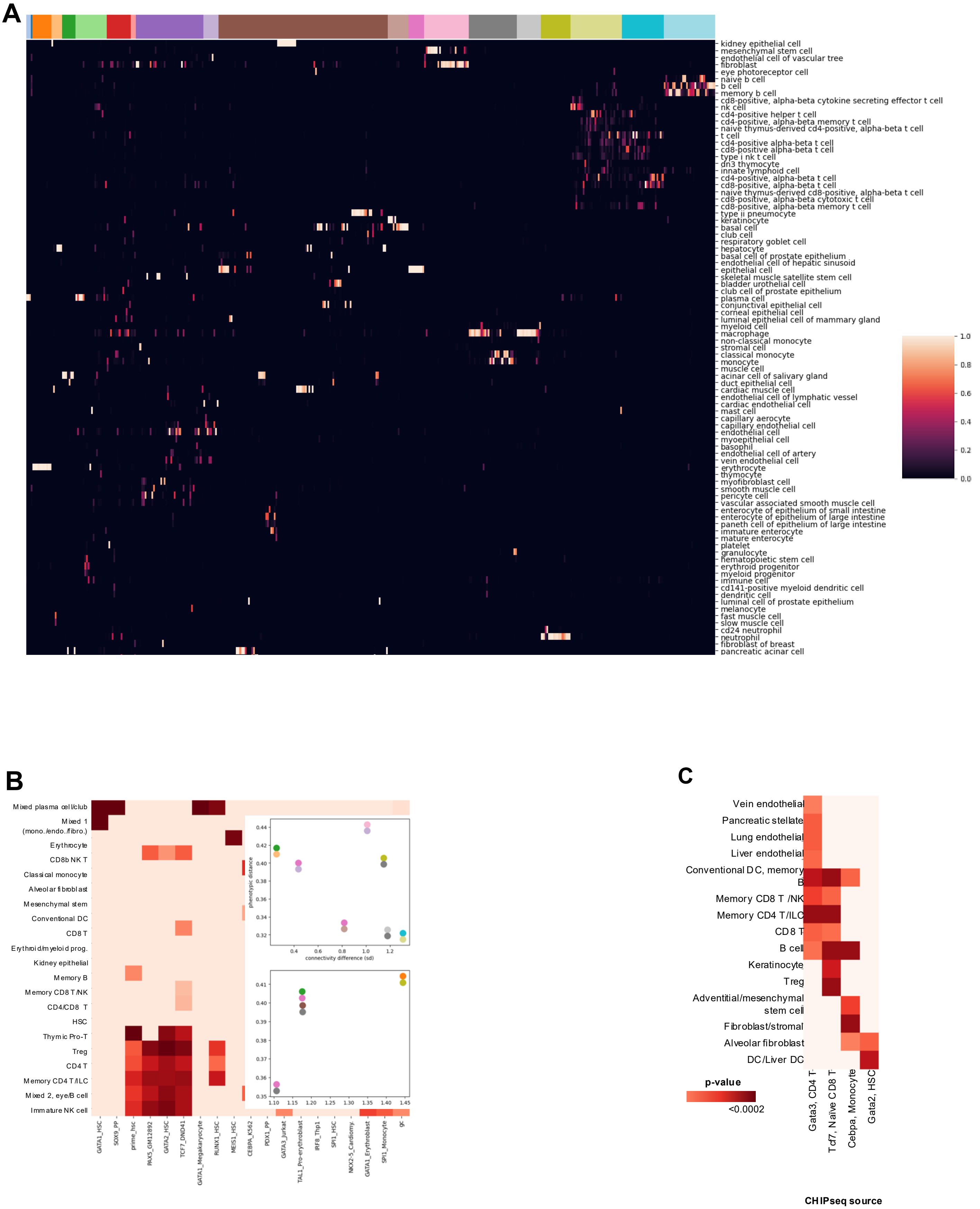

### Supplementary Figure 4

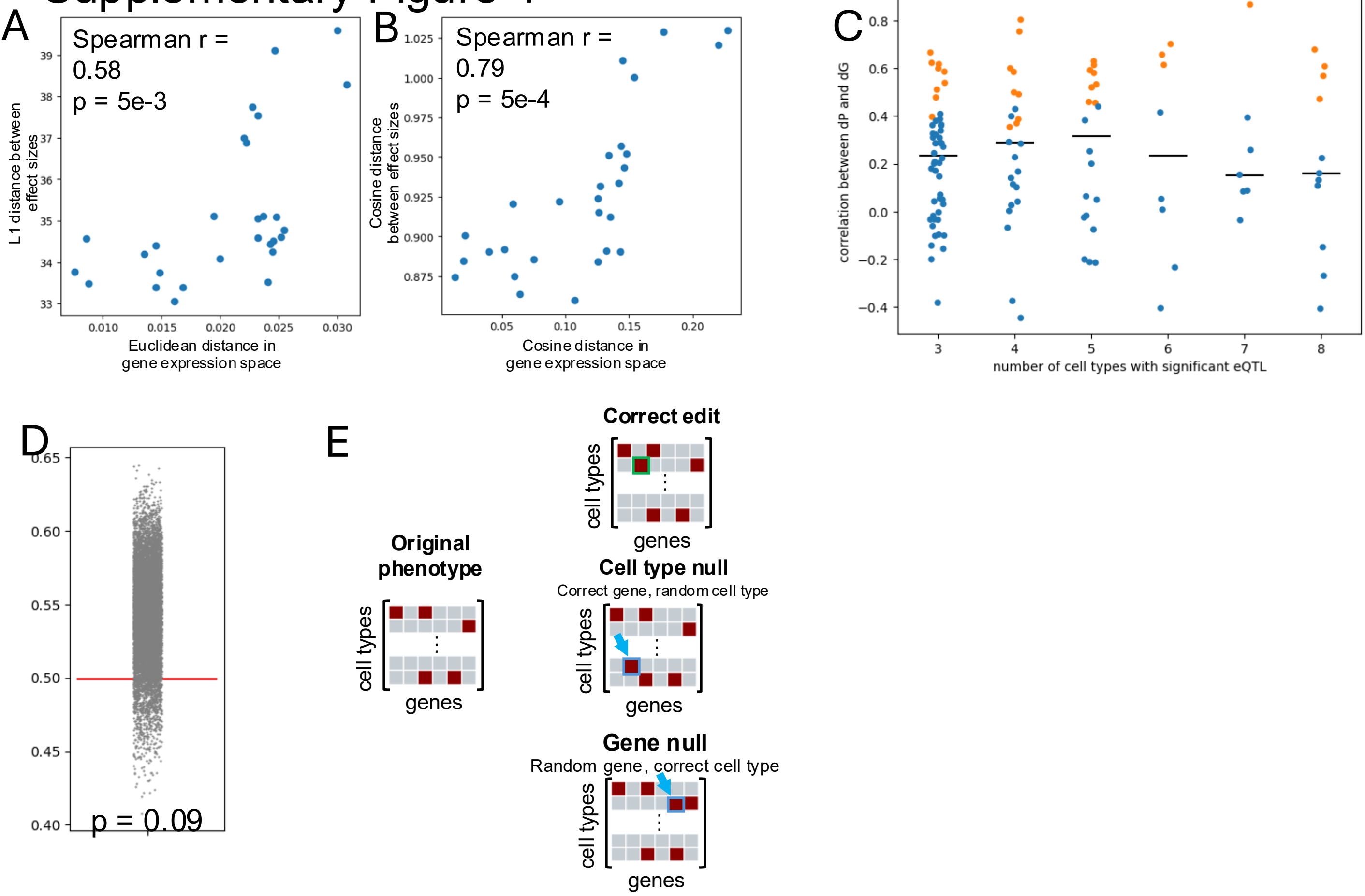

### Supplementary Figure 5

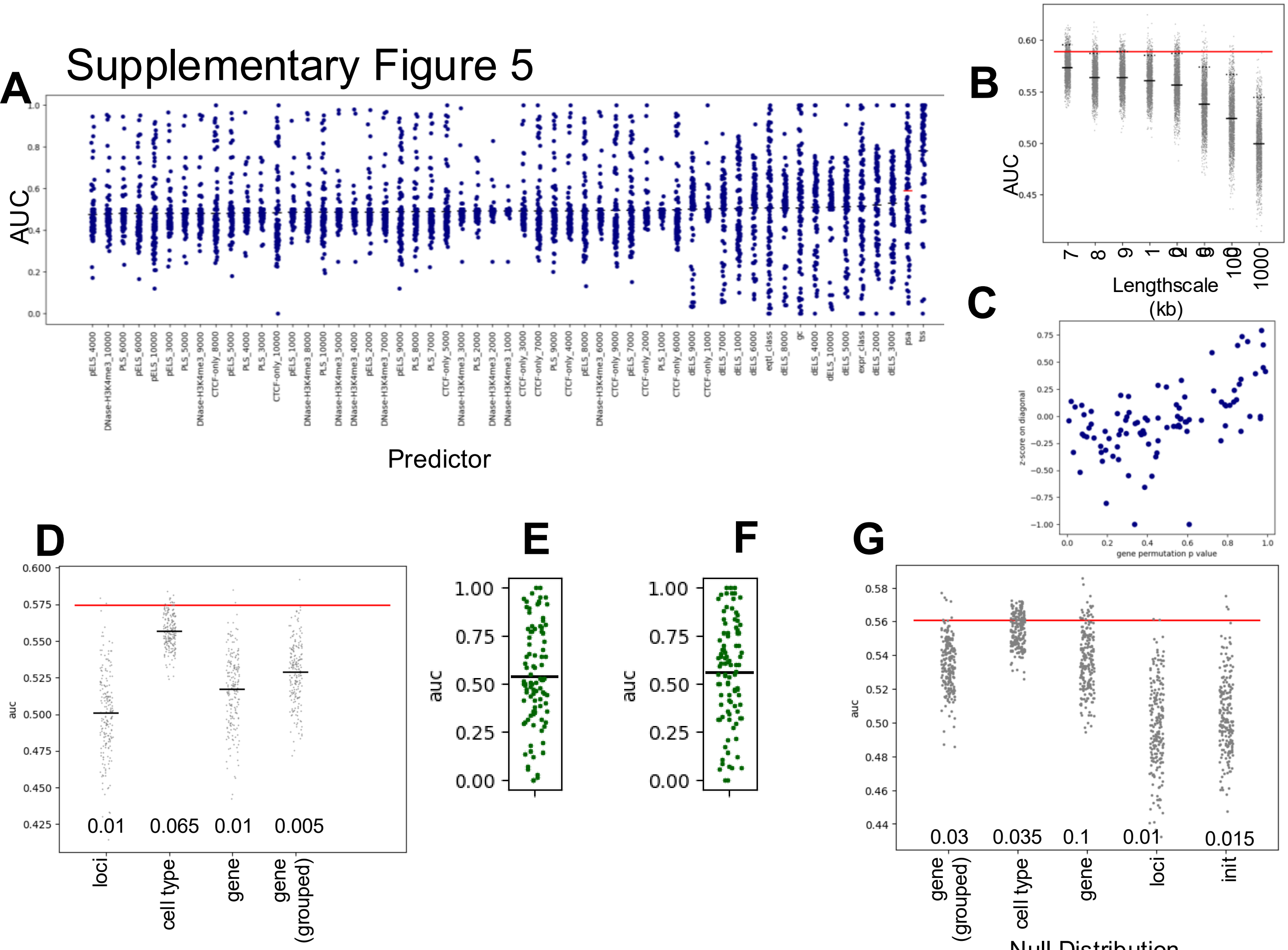


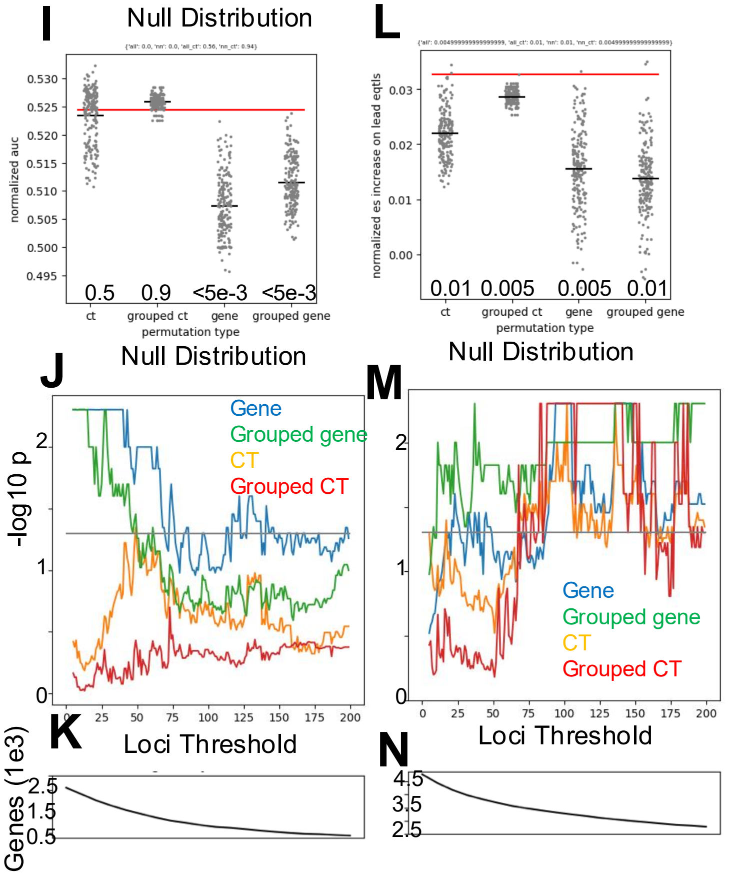


H

| Ablation | AUC |
|---|---|
| Original | 0.589 |
| No syntax consistency | 0.569 |
| Weak phenotype consistency | 0.548 |
| No phenotype consistency | 0.560 |

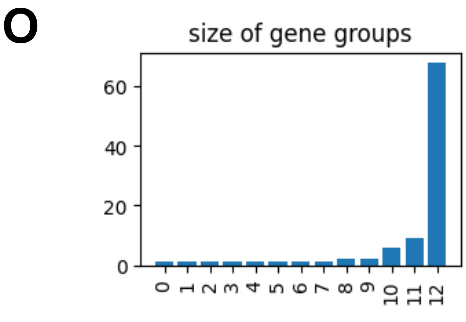

### Supplementary Figure 6

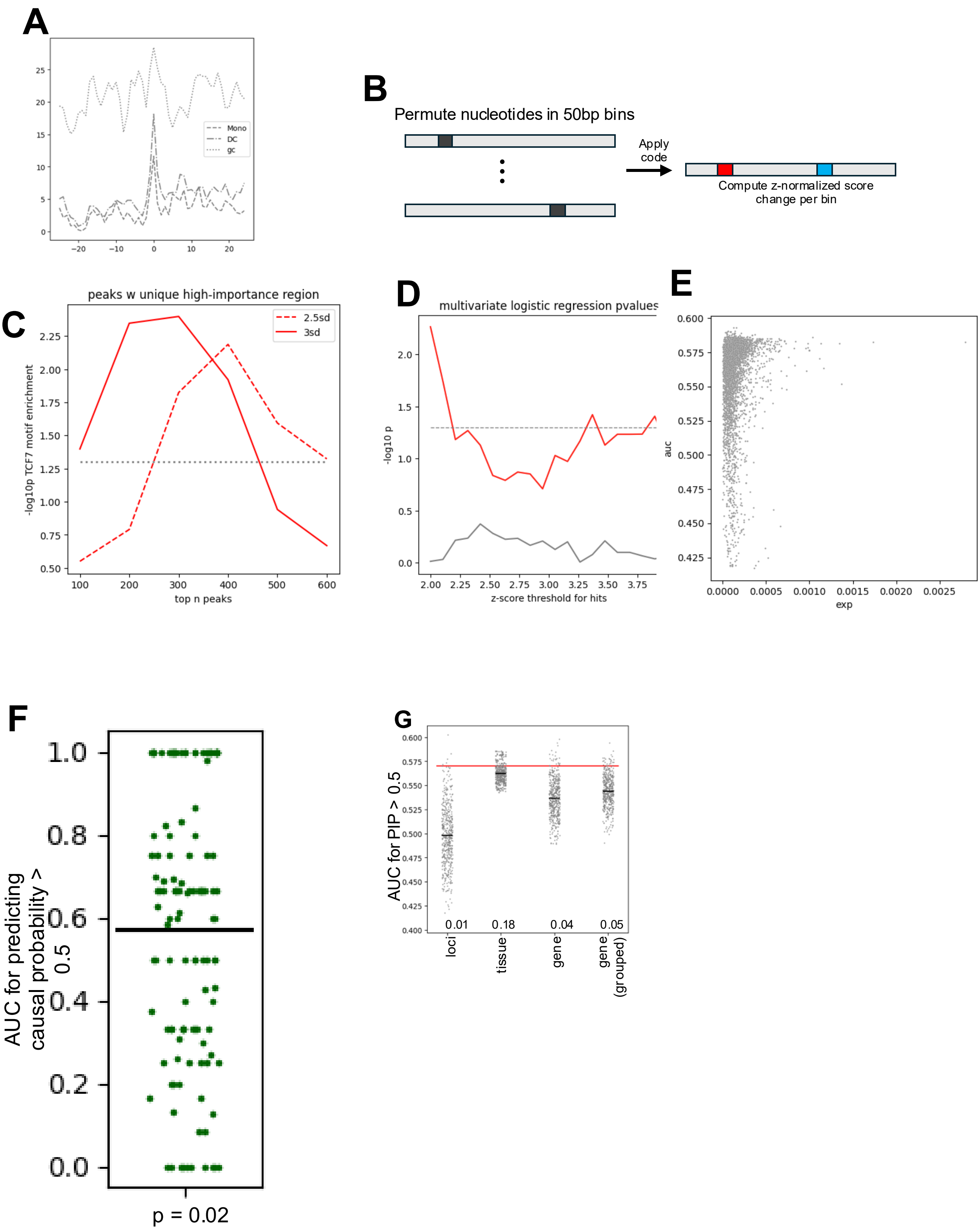

### Supplementary Figure 7

**A**

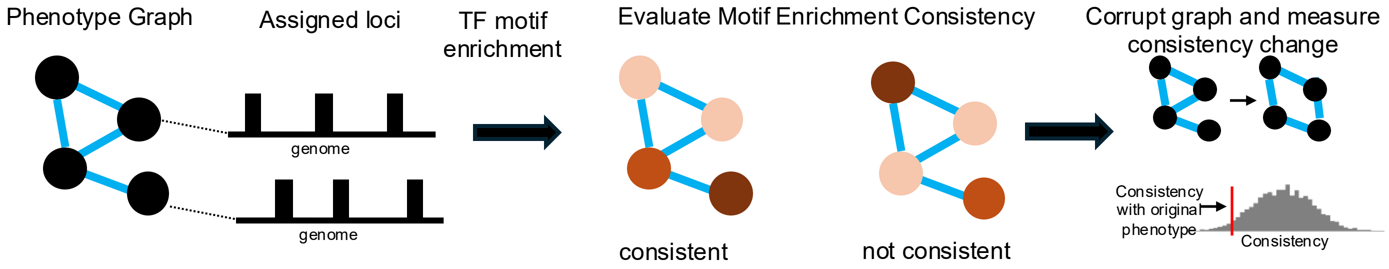


**B**

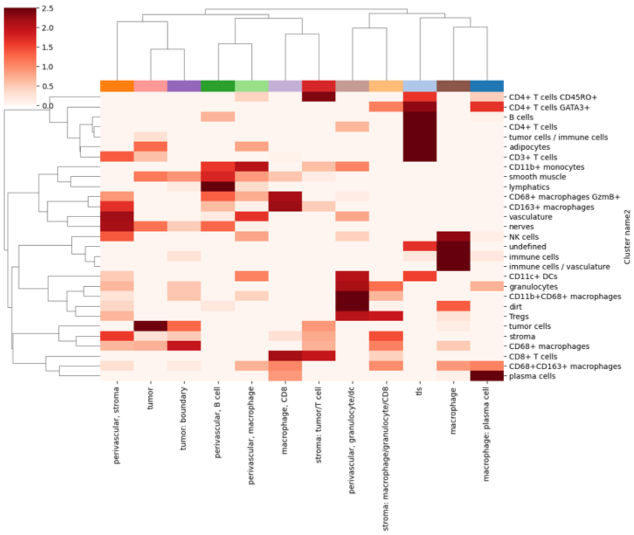

### Supplementary Figure 8

PBMC atlas, replicate 1

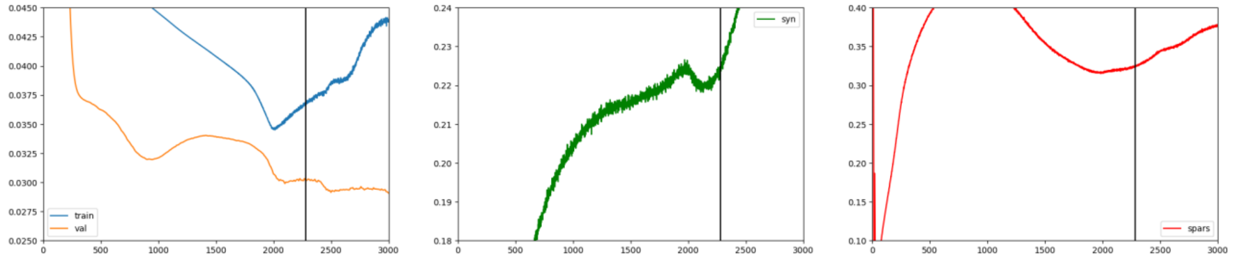


PBMC atlas, replicate 2, latent dim 40

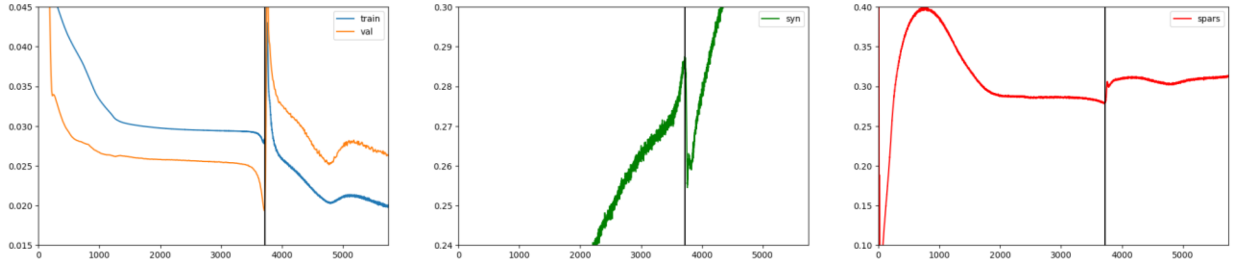


PBMC atlas, replicate 2, latent dim 80

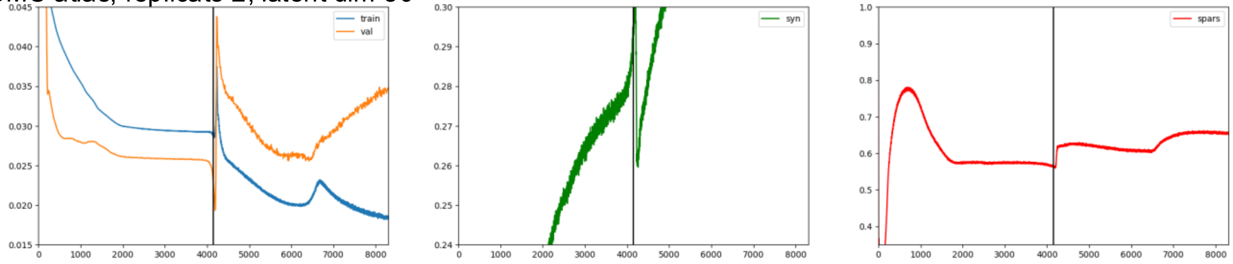


Tabula Sapiens

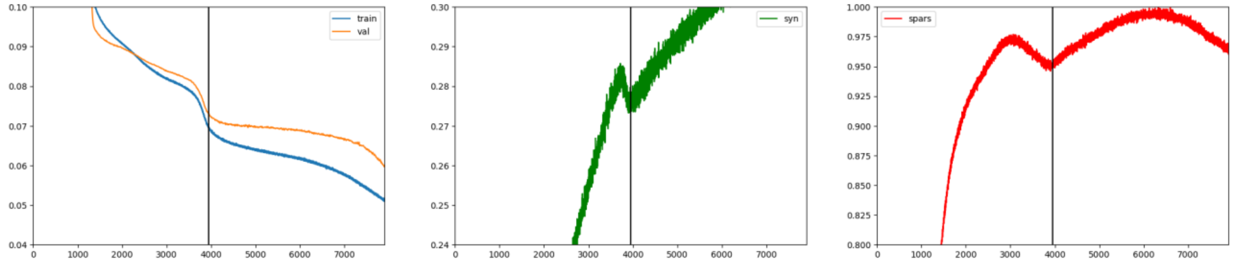
