## Supplementary Note 1 for "Deriving genetic codes for molecular phenotypes from first principles"

July 2025

**Abstract**

There are three notions of genetic code used in the manuscript: single-locus codes, subset encodings, and real-valued codes. They have corresponding PSA algorithms: brute-force PSA-search, optimal transport PSA-search, and neural PSA-search, each catering to the type of code at hand. Here we explain these notions and present a mathematical formalism to highlight their common structure.

#### 1 Introduction

A compositional phenotype is a phenotype with a collection of units with relationships between them. The relationship between units is a notion of proximity. A genetic code is an assignment of genomic loci to phenotypic units. We consider three types of code:

1. Single-locus codes: each unit is assigned a single genomic locus;
2. Subset codes: each unit is assigned a subset of loci; and
3. Real-valued codes: each unit is assigned a real weight for each locus in the genome.

The encoding of a phenotypic unit is then a locus assigned to it (in the single-locus case), a subset of loci assigned to it (in the subset encoding case) or a real-valued functional across loci (in the real-valued case). In this note we explain and relate these three settings.

#### 2 Single-locus codes and brute-force PSA-search

In a single-locus code, each unit of the phenotype is assigned a single locus in the genome. We represent a single-locus code as a function

$$E : X \to \mathbb{G}$$

where $\mathbb{G}$ is a set of genomic loci, and $X$ is the set of units of the phenotype. For $x \in X$ its encoding is $E(x) \in \mathbb{G}$. PSA-search defines relations on the phenotype and on the genomic loci. To do this, it assigns units of the phenotype $x \in X$, and genomic loci $s \in \mathbb{G}$ to measurements given by maps $P : X \to V$ and $Q : \mathbb{G} \to W$.

The two examples of single-locus codes in the manuscript are the protein sequence example and the Allen brain atlas example. We explain each of these in turn.

#### 2.1 Protein sequence example

Each unit $x \in X$ is a protein residue. The set $\mathbb{G}$ is the set of loci (accounting for two orientations and three frame shifts). We seek a map $E$ that sends each residue to the codon that it is encoded by in the genome.

For a protein sequence of length $m$, the measurements of the phenotype units lie in the set $V = \{\text{amino acids}\} \times \{1, \ldots, m\}$. The map $P : X \to V$ assigns each residue to the amino acid and its relative position in the protein sequence. The measurements of the genomic loci lie in the set $W = \{+, -\} \times \{0, 1, 2\} \times \{1, \ldots, n\} \times \{ATG, \ldots, GGG\}$, where the two orientations are $\{+, -\}$ and the three frames are $\{0, 1, 2\}$. The map $Q : \mathbb{G} \to W$ assigns each locus to its orientation, frame, position, and nucleotide triplet.

Put differently, we define the relations eq and adj on the protein sequence with $(a, a') \in \text{eq} \Leftrightarrow a$ and $a'$ have the same amino acid, and $(a, a') \in \text{adj} \Leftrightarrow a$ and $a'$ are adjacent. We analogously define these relations for the genomic triplets. A code $E : X \to \mathbb{G}$ preserves these relations if $(a, a') \in \text{eq} \Leftrightarrow (E(a), E(a')) \in \text{eq}$, and likewise for adjacency.

We search through assignments of codons and compute the number of mismatches when adjacent residues are assigned to adjacent codons. The output of PSA-search is the set of best assignments (they are unique, in practice, for sufficiently long sequences). The protein sequence example can be explained using binary vectors, as we now explain.

##### 2.1.1 Protein sequence encoding via binary vectors

We implement PSA-search by using the relations of equality and adjacency to turn a protein sequence into a set of binary vectors, and then solve a binary vector matching problem.

**Example 2.1.** Amino acid sequence $MKVKV$ is encoded by three binary vectors: the $M$ vector is $(1, 0, 0, 0)$, the $K$ vector is $(0, 1, 0, 1, 0)$ and the $V$ vector is $(0, 0, 1, 0, 1)$. The sequence $VKV$ is encoded by two binary vectors, $(1, 0, 1)$ and $(0, 1, 0)$.

Similarly, we turn a genome sequence into a set of binary vectors.

**Example 2.2.** Consider the genome sequence

$$ATGAAAAAAAGC.$$

It splits into triplets in one of six ways, accounting for three frames and two orientations:

$$ATG|AAA|AAA|AGC \quad A|TGA|AAA|AAA|GC \quad AT|GAA|AAA|AAG|C$$

$$CGA|AAA|AAA|GTA \quad C|GAA|AAA|AAG|TA \quad CG|AAA|AAA|AGT|A$$

There are nine triplets that appear across these:

$$ATG, AAA, AGC, TGA, GAA, AAG, CGA, GTA, AGT$$

Thus the 3-mers in the string are captured via adjacency and equality as nine sets of six binary strings. For example, the triplet $AAA$ therefore has as its six vector encodings

$$(0, 1, 1, 0), \quad (0, 1, 1), \quad (0, 1, 0), \quad (0, 1, 1, 0), \quad (0, 1, 0), \quad (1, 1, 0),$$

according to where it appears in each of the six binary strings.

Given a protein sequence and genome sequence, both encoded as binary vectors, we perform brute-force PSA search, as follows. We slide each amino-acid binary vector along each codon binary vector. The PSA-loss is the proportion of mismatches. We select the position and the resulting amino-acid/codon match with the highest proportion of matches.

**Example 2.3.** We apply brute-force PSA-search on protein sequence $VKV$ with the 3-mers of the genome sequence from Example 2.2. Its two binary strings are $(0, 1, 0)$ (from $K$) and $(1, 0, 1)$ (from $V$). Comparing against all $9 \times 6$ binary strings that come from the triplets in the genome sequence, we see that the string $(1, 0, 1)$ never appears, and $(0, 1, 0)$ appears twice: both times from the presence of $AAA$ in the string $GAA|AAA|AAG$. Hence the brute force search returns as its best alignment that $K$ is $AAA$ and that $V$ is either $AAG$ or $GAA$ (both of the two possibilities are equidistant).

As the sequence lengths grow, the approach is able to recover the true alignment. We consider the computational complexity of the approach. Let $a$ denote the number of unique amino acids in the protein sequence and let $m$ be the length of the sequence. The protein sequence is encoded by $a$ binary strings of length $m$.

Consider a genome sequence of length $kn$ with $h$ distinct $k$-mers. There are $2k$ different ways to split the genome sequence into $k$-mers, accounting for the $k$ frames and two orientations. Considering the ways to split a vector of length $kn$ into pieces of length $k$, we see that two splits have length $n$ and the remaining $2(k-1)$ have length $n-1$, cf. Example 2.2. So the genome sequence is encoded by $2h$ binary vectors of length $n$ and $2h(k-1)$ of length $n-1$.

We slide the protein sequence vectors, along the genome sequence binary vectors. In each genome sequence binary vector of length $n$ there are $n-m+1$ positions and in the vectors of length $n-1$ there are $n-m$ positions. This gives $2h(k(n-m)+1)$ binary strings of length $m$.

There are therefore $2ah(k(n-m)+1)$ binary vectors to compute (subtracting the protein vectors from the corresponding genome vector at each position). The output of PSA-search is the set of sparsest such difference vectors (i.e., those with fewest mismatches).

**Example 2.4.** In Example 2.3, the protein sequence has length $m = 3$ with two distinct amino acids appearing (so $a = 2$). We are considering 3-mers (so $k = 3$) of which $h = 9$ are distinct in the genome sequence. The genome sequence has length $3n = 12$ so $n = 4$. Thus the brute force search amounts to looking at 144 binary strings and seeing is sparsest.

#### 2.2 The brain atlas example

Each unit $x \in X$ is a brain region, the set $\mathbb{G} =$ ENCODE enhancers. We seek a map $E$ that sends each brain region to an enhancer that can causally alter cellular phenotypes in the brain region. The measurement maps are $P : X \to \mathbb{R}^{20000}$, which sends each brain region to its gene expression, and $Q : \mathbb{G} \to \mathbb{R}^{2000}$, which sends each enhancer to its sequence features. These maps define similarity between gene expression and sequence features.

An encoding preserves these relationships if the phenotypic distance between brain regions is the same as the feature distance between enhancers. Formally, we define the relations

$d_r$ for $r \in \mathbb{R}^+ \cup \{0\}$ where $(a, a') \in d_r \Leftrightarrow d(a, a') = r$; where $d$ is the cosine distance between gene expression vectors for the brain atlas.

We search through seed assignments of each enhancer to a fixed reference brain region region and compute the maximum difference between the sequence distances between enhancers from the seed enhancer to other enhancers and the reference region to other regions. We select the top twenty assignments.

### 3 Subset encodings and optimal transport PSA-search

Subset encodings assign phenotypic units to a set of genomic loci. We define a subset code to be a function $E : X \to \mathcal{P}(\mathbb{G})$ where $\mathcal{P}(\mathbb{G})$ is the set of subsets of $\mathbb{G}$. The subset codes in the manuscript are the encodings of the Tabula Sapiens atlas in ENCODE enhancers and the code for the CD8+ T cell neighborhoods in T2T.

Evaluating PSA for subset encodings requires defining relations on the phenotype and on subsets of the genomic loci.

- $E$ sends units to subsets of genomic loci that are either equal or nonoverlapping; i.e. $E$ first sends a unit to a cluster of phenotypic units and then assigns all units in that cluster to a cluster of loci.
- We define the distance between subsets of genomic loci to be the distance between their centroids.
- The encoding exhibits PSA if the centroid distance between the subsets assigned to each phenotypic unit are equal to distance between the units.

In the manuscript, we perform semi-relaxed Gromov-Wasserstein optimal transport to find such a subset encoding. We infer the codes from the encoding by performing motif enrichment analysis. We show that the code that generates the subset encodings we find corresponds to transcription factor motif enrichment.

### 4 Real-valued codes and neural PSA-search

Real valued codes have as encoding of a phenotypic unit a weight assigned to each genomic locus. A code here is a map $E : X \to (\mathbb{G} \to \mathbb{R})$. We assume measurements $P : X \to V$ and $Q : \mathbb{G} \to W$ where $V$ and $W$ are real vector spaces. The code is now no longer parameterized as a discrete correspondence. Instead, we use neural networks, as follows.

- In neural PSA-search, the encoding $E(x)$ is represented by a neural network $T(p, q)$, where $E(x)(s) = T(P(x), Q(s))$. The function $T$ computes the encoding of $x$ at $s \in \mathbb{G}$.
- We seek a code $T$ that represents how much a locus with features $q$ contributes to the characteristics of units with features $p$. The difference $|T(p, q) - T(p', q)|$ is high at loci with features $q$ that contribute differentially to characteristics of units with features $p$ and $p'$. Hence when $p$ and $p'$ are gene expression vectors differing by a small amount in the expression of one gene, we interpret $|T(x, q) - T(x, q')|$ as a prediction score for expression quantitative trait loci for that gene.

We define two loss functions that measures the extent to which PSA holds for a real-valued code $T$.

- Phenotypic consistency measures the extent to which the similarities between the encodings of phenotypic units deviate from their similarities with respect to measurements. It is captured by the loss function:

$$C_P(T) := \sum_{x,y \in X} \left[ d(p(x), p(y)) - \frac{1}{N} \sum_{i \in \mathbb{G}} |T(p(x), q(i)) - T(p(y), q(i))| \right]$$

- Syntactic consistency measures the extent to which loci with similar measurements encode phenotypic units similarly. It is given by loss function

$$C_G(T) := \sum_{x} \left[ \frac{(T(p(x), q(s)) - \frac{1}{K} \sum_{t \in N(s)} T(p(x), q(t)))^2}{\sum_{x \in X} T(p(x), q(s))^2} \right],$$

  where $N(s)$ encodes the $K$-nearest neighbors of $s$ with respect to features $q$. This term is small when variation of the phenotypic associations across similar loci is small; the denominator normalizes the loss function.

#### 4.1 Neural PSA-search

Neural PSA-search is a minimal algorithm to search for parsimonious codes that optimize for phenotypic consistency, syntactic consistency and sparsity. We outline the computational implementation. The code is represented by a neural network whose parameters are optimized by gradient descent in the following way:

1. A batch of $k$-mers is sampled. If using a large input phenotype, a batch of phenotype points must also sampled.
2. The value of $T(x, s)$ for the phenotype points and sampled $k$-mers is evaluated.
3. The loss terms $C_P(T)$ and $C_G(T)$ are computed.
4. A gradient step is conducted on the weights of $T(x, s)$ using the loss function:

$$\mathcal{L}(T) = C_P + C_G + \lambda C_S,$$

   where $C_S$ is a sparsity regularizer for the code $T(x, s)$.

We now detail each of these steps.

#### 4.2 Neural network architecture of T

The code $T(x, s)$ is represented using neural networks as follows.

- A phenotype embedding network $F : \mathbb{R}^p \to \mathbb{R}^q$ extracts a latent representation of a point in phenotype space.

- A syntax embedding network $G : \{A, T, C, G\}^k \to \mathbb{R}^q$ extracts a latent representation of a $k$-mer from the genome.
- The encodings are obtained from the $F$ and $G$ as:

$$T(x, s) = \tanh\left(\frac{\langle \exp(F(x)), G(s)\rangle}{\sum_j \exp(F(x)_j)}\right)$$

In the manuscript, the phenotype embedding network is a linear map and the syntax embedding network is a linear map applied to features extracted from a pretrained language model, whose weights are frozen.

#### 4.3 Training

We now discuss how the loss functions are computed using batch-wise training.

##### 4.3.1 Syntactic consistency

Syntactic consistency utilizes the following precomputation, prior to training:

- The embeddings from a language model are extracted for 2000mers in the genome beginning every 1000 nucleotides, yielding a collection of approximately 3.5M vectors of syntactic embeddings.
- A 20-nearest neighbor graph using the cosine similarity of embeddings is computed, yielding for each syntactic embedding vector $s$ a set of syntactically 20 similar embeddings $N(s)$

At training time, a batch of sequences $\mathcal{B}$ (of size $10^5$) is a sampled from the syntactic embedding vectors the set of 3.5M vectors. No other sequence information is used in the training procedure. The loss function for syntactic consistency $C_G(T)$ is:

$$C_G(T) := \sum_{s\in\mathcal{B}} \sum_{x\in X} \left[\frac{(T(x,s) - \frac{1}{K}\sum_{t\in N(s)} T(x,t))^2}{\sum_{x\in X} T(x,s)^2}\right].$$

##### 4.3.2 Phenotypic consistency

The loss function for consistency with phenotypic geometry $C_P(T)$ is computed on the batch $\mathcal{B}_S$ of sampled sequences:

$$C_P(T) = \sum_{i>j} \left| 1 - \frac{|\langle x_i, x_j\rangle|}{||x_i||\,||x_j||} - \frac{1}{|\mathcal{B}|}\sum_{s\in\mathcal{B}} |T(x_i, s) - T(x_j, s)| \right|.$$

##### 4.3.3 Sparsity

The sparsity constraint is:

$$C_S(T) := \sum_x \sum_s |T(x, s)|.$$

###### 4.3.4 Loss function

The final form of the loss function was

$$L(T) = C_P(T) + \alpha_G C_G(T) + \alpha_S C_S(T).$$

The weights for each of the loss terms were $\alpha_G = 1$ and $\alpha_S = 0.1$.

##### 4.4 Model selection

We now details how the model used in downstream evaluation is obtained.

###### 4.4.1 Initialization

We initialize the phenotype embedding and syntax embedding networks with PCA applied to the phenotype features or the syntactic features respectively. This eliminates randomness from initialization.

###### 4.4.2 Early stopping

We observed in training that the loss is dominated by the loss for phenotypic consistency. In addition, we observed that PSA starts overfitting to the input data (measured by evaluating consistency on a held out set of phenotype datapoints). We therefore implemented early stopping of the model at the epoch where: the training or validation loss for consistency with phenotypic geometry starts increasing and we observe a trough in the sparsity and syntactic consistency loss. The specific stopping points for the codes in the manuscript, along with their loss curves, are detailed in the Methods section.
