## Supplementary material for "Deriving genetic codes for molecular phenotypes from first principles": Code: Fig1_AA_codes.ipynb

```
# %% [code]
import pandas as pd
import numpy as np
import itertools
from tqdm.notebook import tqdm
import torch
torch.cuda.set_device(2)

# %% [code]
with open('../data/seq/ecoli') as f:
    w = f.read()

# %% [code]
comp = {'A':'T','T':'A','G':'C','C':'G'}

# %% [code]
seq = ''.join(w.split('\n')[1:])
rev = ''.join([comp[a] for a in seq[::-1]])

# %% [code]
overlap_f1 = [[seq[i:i+3], seq[i+1:i+4], seq[i+2:i+5]] for i in range(0, len(seq), 3)][:-2]

# %% [code]
overlap_f1[-2:]

# %% [code]
f1 = [seq[i:i+3] for i in range(0, len(seq),3)][:-2]
f2 = [seq[i:i+3] for i in range(1, len(seq),3)][:-1]
f3 = [seq[i:i+3] for i in range(2, len(seq),3)][:-1]

# %% [code]
rf1 = [rev[i:i+3] for i in range(0, len(rev),3)][:-2]
rf2 = [rev[i:i+3] for i in range(1, len(rev),3)][:-1]
rf3 = [rev[i:i+3] for i in range(2, len(rev),3)][:-1]

# %% [code]
vocab = {}
for k, tup in enumerate(itertools.product(['A','T','C','G'],['A','T','C','G'],['A','T','C','G'])):
    vocab[''.join(tup)] = k
vocab =pd.Series(vocab)
inv_vocab = {v:k for k,v in vocab.to_dict().items()}
cols = [inv_vocab[k] for k in range(len(inv_vocab))]

# %% [code]
vocab

# %% [code]
f1_t = vocab.loc[f1].values.astype(np.uint8)
f2_t = vocab.loc[f2].values.astype(np.uint8)
f3_t = vocab.loc[f3].values.astype(np.uint8)

rf1_t = vocab.loc[rf1].values.astype(np.uint8)
rf2_t = vocab.loc[rf2].values.astype(np.uint8)
rf3_t = vocab.loc[rf3].values.astype(np.uint8)

# %% [code]
y.shape

# %% [code]
neye =np.eye(64)
y = torch.from_numpy(np.stack([neye[f1_t],neye[f2_t],neye[f3_t],neye[rf1_t],neye[rf2_t],neye[rf3_t]])).long()
y = y.permute(0,2,1).cuda()

# %% [code]
with torch.no_grad():
    y_ind = y.argmax(axis = 1)

# %% [code]
recA = 'MAIDENKQKALAAALGQIEKQFGKGSIMRLGEDRSMDVETISTGSLSLDIALGAGGLPMGRIVEIYGPESSGKTTLTLQVIAAAQREGKTCAFIDAEHALDPIYARKLGVDIDNLLCSQPDTGEQALEICDALARSGAVDVIVVDSVAALTPKAEIEGEIGDSHMGLAARMMSQAMRKLAGNLKQSNTLLIFINQIRMKIGVMFGNPETTTGGNALKFYASVRLDIRRIGAVKEGENVVGSETRVKVVKNKIAAPFKQAEFQILYGEGINFYGELVDLGVKEKLIEKAGAWYSYKGEKIGQGKANATAWLKDNPETAKEIEKKVRELLLSNPNSTPDFSVDDSEGVAETNEDF'


# %% [code]


# %% [code]
len(input_seq)

# %% [code]
from torch.nn.functional import conv1d as conv

# %% [code]
corrects = {}
# build seq
k =0
input_seq = recA
np.random.seed(123)
detection = {}
pos_samp = [0]
for position in tqdm(pos_samp):
    coded = []
    counts = {}
    aa_vocab = {}

    for idx,aa in enumerate(input_seq[position:(position+length)]):
        if aa not in aa_vocab:
            aa_vocab[aa] = k
            counts[aa_vocab[aa]] = np.zeros(len(input_seq),dtype = int)
            k+=1

        coded.append(aa_vocab[aa])

        counts[aa_vocab[aa]][idx]= 1

    coded = np.array(coded)
    inv_aa_vocab = {v:k for k,v in aa_vocab.items()}

    sorted_counts = pd.Series({k:v.sum() for k,v in counts.items()}).sort_values(ascending = False)
    order = sorted_counts.index.values

    #see if correctly localized
    res = {}
    conv_out = {}
    pos = {}
    for codon in range(64):
        codq = (y_ind==codon).half()
        for aa in order[:5]:
            x = torch.from_numpy(counts[aa]).half().cuda()
            r = conv(codq[:,None,:], x[None,None])/x.sum().item()
            conv_out[codon,aa] = (r[:3,0].amax(axis=0).cpu().numpy(), r[-3:,0].amax(axis=0).cpu().numpy()[::-1])
            R, p = r[:,0].amax(axis = 0).max(axis = 0) # says where in the linear genome there is best match
            res[codon,aa] = R.item()/x.sum().item() # % equalities correct
            pos[codon,aa] = (p.item(),r[:,0,p].argmax(axis = 0).item())

    df = pd.melt(pd.Series(res).unstack().reset_index(),id_vars = ['index']).sort_values('value',ascending = False)
    ranks = []
    for codon,aa in zip(df['index'].iloc[:20], df['variable'].iloc[:20]):
        a, f = pos[codon,aa]
        if f>2:
            pp = 3*((y.shape[-1])-a) # reverse orientation to find position - the reverse ones are concatenated with forward ones so positions do not correspond
        else:
            pp = 3*a # forward orientation
        ranks.append(pp)
    ranks = np.array(ranks)    
    c = np.where(ranks==2823765)[0]
    if len(c)==0:
        s = -1
    else:
        s = np.min(c)
    detection[length,position] = s

# %% [code]
A = np.stack([v for k,v in conv_out.items() if k[1]==2])
fcodons = A[:,0].argmax(axis = 0)
rcodons = A[:,1].argmax(axis = 0)
B = A.max(axis = 0)
scores = pd.DataFrame({'score': B.max(axis = 0), 'strand': B.argmax(axis = 0), 'fcodon':fcodons,'rcodons':rcodons})
scores['pos'] = 3*scores.index.values #+ 3* (len(scores) - scores.index.values + y.shape[-1]- len(scores)) *(scores['strand']==1)
scores['codon'] = [inv_vocab[a] if strand==0 else inv_vocab[b] for a,b, strand in zip(scores['fcodon'],scores['rcodons'], scores['strand'])]
plt.figure(figsize = (4,2))
plt.vlines(2823769, 0.15,1, color = 'black',zorder = -50)

sns.scatterplot(data = scores[scores['score']>0.4], hue = 'codon', y = 'score', x = 'pos',s = 50)
plt.legend(loc = 'upper left')
plt.xlabel('Genomic Coordinate')
plt.ylabel('Match with I')
plt.ylim(0.2,1)
plt.xlim(0,4.8e6)

# %% [code]
meltR2[meltR2['value']>0.2]
meltR2['codon'] = meltR2['variable'].map(inv_vocab)
meltR2


# %% [code]
codon = 6
codq = (y_ind==codon).half()

aa = 2
x = torch.from_numpy(counts[aa]).half().cuda()
r = conv(codq[:,None,:], x[None,None])
R, p = r[:,0].amax(axis = 0).max(axis = 0) # says where in the linear genome there is best match
fw = (r[:3,0].amax(axis = 0)/x.sum().item()).cpu().numpy()
rev = (r[-3:,0].amax(axis = 0)/x.sum().item()).cpu().numpy()

R.item()/x.sum().item()
R = pd.DataFrame(r[:,0].cpu().numpy()/x.sum().item()).T.melt(ignore_index = False)
R['frame'] = R['variable'].map({0:'+,0', 1:'+,1',2:'+,2',3:'-,0',4:'-,1',5:'-,2'})
R['pos'] = 3*R.index.values * (R['variable']<3) + 3*(r.shape[-1]  - R.index.values + y.shape[-1]- r.shape[-1])* (R['variable']>=3) 

plt.figure(figsize = (4,2))
sns.scatterplot(data = R[R['value']>0.15], y = 'value', hue = 'frame', x = 'pos', s =50)
plt.legend(loc = 'upper left')
plt.vlines(2823769, 0.15,1, color = 'black',zorder = -50)
plt.ylim(0.2,1)
plt.xlabel('Genomic coordinate')
plt.ylabel('Score I==ATC')
# plt.scatter(np.arange(len(fw)),rev[::-1], c= 'navy')
plt.ylim(0.2,1)
plt.xlim(0,4.8e6)


# %% [code]
# codon = 6
# aa = 2
all_scores = []
all_aa = []
for codon,aa in pos.keys():
    codon_where = (y_ind==codon).half()
    
    position,frame = pos[codon,aa]
    
    
    N = 300
    codon_frame = codon_where[frame,position:position+N]
    aa_where = torch.from_numpy(counts[aa]).half().cuda()
    all_scores.append(((codon_frame==aa_where[:N]).float().mean()).item())
    all_aa.append((codon,aa))
all_scores = np.array(all_scores)
all_aa = np.array(all_aa)

# sanity check:
all_aa[all_scores.argmax()]

# %% [code]
codon = 6
aa = 2

codon_where = (y_ind==codon).half()

position,frame = pos[codon,aa]


N = len(counts[aa])
codon_frame = codon_where[frame,position-N:position+N].cpu().numpy()
aa_where = torch.from_numpy(counts[aa]).half().cuda().cpu().numpy()
aa_where = torch.from_numpy(counts[aa]).half().cuda().cpu().numpy()

plt.figure(figsize = (20,2))
sns.heatmap(codon_frame[None],cmap = 'PuOr_r',cbar = False,xticklabels=False,yticklabels=False)


# %% [code]
plt.figure(figsize = (10,2))
sns.heatmap(aa_where[None],cmap = 'PuOr_r',cbar = False,xticklabels=False,yticklabels=False)


# %% [code]

from matplotlib import pyplot as plt

# %% [code]
code = {}
for aa, codon in zip(recA,[inv_vocab[a.item()] for a in  y_ind[frame,(position):position+500]]):
    code.setdefault(aa,[]).append(codon)

# %% [code]
B = pd.concat({k: pd.Series(v).value_counts(normalize = True) for k,v in code.items()})

A = B[B>0.4].unstack().fillna(0)
A = A[A.sum(axis = 1)>0]

f = sns.clustermap(A.sort_index(),figsize = (5,5),xticklabels = True, yticklabels = True,vmin = 0.4)
f.ax_col_dendrogram.set_visible(False)
f.ax_cbar.set_visible(False)
f.ax_row_dendrogram.set_visible(False)

# %% [markdown]
### Compare overlapping and non-overlapping

# %% [code]
import seaborn as sns

# %% [code]
conv_out2 = {}
res2 = {}
pos2 = {}
for codon in range(64):
    codq_non = (y_ind==codon).half()
    codq = torch.stack([codq_non[:3].amax(axis =0),codq_non[-3:].amax(axis =0)])
    for aa in order[:5]:
        x = torch.from_numpy(counts[aa]).half().cuda()
        r = conv(codq[:,None,:], x[None,None])/x.sum().item()
        conv_out2[codon,aa] = (r[0,0].cpu().numpy(), r[1,0].cpu().numpy()[::-1])
        R, p = r[:,0].amax(axis = 0).max(axis = 0) # says where in the linear genome there is best match
        res2[codon,aa] = R.item()/x.sum().item() # % equalities correct
        pos2[codon,aa] = (p.item(),r[:,0,p].argmax(axis = 0).item())

# %% [code]
codq_non[:3].amax(axis = 0)

# %% [code]
scores_nonov = scores.copy()

# %% [code]
scores[scores['score']>0.8]

# %% [code]
plt.figure(figsize = (5,5))
plt.scatter(scores_nonov['score'],scores['score'], s = 25, c = 'black')
plt.plot([0,1],[0,1],zorder = -1, color = 'black',linewidth = 1)
plt.xlabel('score, strictly non-overlapping')
plt.ylabel('score, overlaps permitted')

# %% [code]

A = np.stack([v for k,v in conv_out2.items() if k[1]==2])
fcodons = A[:,0].argmax(axis = 0)
rcodons = A[:,1].argmax(axis = 0)
B = A.max(axis = 0)
scores = pd.DataFrame({'score': B.max(axis = 0), 'strand': B.argmax(axis = 0), 'fcodon':fcodons,'rcodons':rcodons})
scores['pos'] = 3*scores.index.values #+ 3* (len(scores) - scores.index.values + y.shape[-1]- len(scores)) *(scores['strand']==1)
scores['codon'] = [inv_vocab[a] if strand==0 else inv_vocab[b] for a,b, strand in zip(scores['fcodon'],scores['rcodons'], scores['strand'])]
plt.figure(figsize = (4,2))
plt.vlines(2823769, 0.15,1, color = 'black',zorder = -50)

sns.scatterplot(data = scores[scores['score']>0.42], hue = 'codon', y = 'score', x = 'pos',s = 50)
plt.legend(loc = 'upper left')
plt.xlabel('Genomic Coordinate')
plt.ylabel('Match with I')
plt.ylim(0.4,1)
plt.xlim(0,4.8e6)

# %% [code]
scores

# %% [code]


# %% [code]
codon = 6
aa = 2

codq_non = (y_ind==codon).half()
codon_where = torch.stack([codq_non[:3].amax(axis =0),codq_non[-3:].amax(axis =0)])

position,frame = pos2[codon,aa]


N = len(counts[aa])
codon_frame = codon_where[frame,position-N:position+N].cpu().numpy()
aa_where = torch.from_numpy(counts[aa]).half().cuda().cpu().numpy()
aa_where = torch.from_numpy(counts[aa]).half().cuda().cpu().numpy()

plt.figure(figsize = (20,2))
sns.heatmap(codon_frame[None],cmap = 'PuOr_r',cbar = False,xticklabels=False,yticklabels=False)


# %% [code]
codon = 6
aa = 2

codq_non = (y_ind==codon).half()
codon_where = torch.stack([codq_non[:3].amax(axis =0),codq_non[-3:].amax(axis =0)])

position,frame = pos[codon,aa]


N = len(counts[aa])
codon_frame = codon_where[frame,position-N:position+N].cpu().numpy()
aa_where = torch.from_numpy(counts[aa]).half().cuda().cpu().numpy()
aa_where = torch.from_numpy(counts[aa]).half().cuda().cpu().numpy()

plt.figure(figsize = (20,2))
sns.heatmap(codon_frame[None],cmap = 'PuOr_r',cbar = False,xticklabels=False,yticklabels=False)


# %% [code]


# %% [code]
pos[6,2]

# %% [markdown]
### Subsequence analysis

# %% [code]

# build seq
k =0

np.random.seed(123)
detection = {}
for length in tqdm(range(50,150,10)):
    valid_pos = np.arange(len(recA)-length)
    pos_samp = sorted(np.random.choice(valid_pos, size = 20, replace = False))
    for position in pos_samp:
        coded = []
        counts = {}
        aa_vocab = {}
        input_seq = recA[position:(position+length)]
        for idx,aa in enumerate(input_seq):
            if aa not in aa_vocab:
                aa_vocab[aa] = k
                counts[aa_vocab[aa]] = np.zeros(len(input_seq),dtype = int)
                k+=1

            coded.append(aa_vocab[aa])

            counts[aa_vocab[aa]][idx]= 1

        coded = np.array(coded)
        inv_aa_vocab = {v:k for k,v in aa_vocab.items()}

        sorted_counts = pd.Series({k:v.sum() for k,v in counts.items()}).sort_values(ascending = False)
        order = sorted_counts.index.values

        #see if correctly localized
        res = {}
        pos = {}
        with torch.no_grad():
            for codon in range(64):
                codq = (y_ind==codon).half()
                for aa in order[:5]:
                        x = torch.from_numpy(counts[aa]).half().cuda()
                        r = conv(codq[:,None,:], x[None,None])
                        R, p = r[:,0].amax(axis = 0).max(axis = 0)
                        res[codon,aa] = R.item()/x.sum().item() # % equalities correct
                        pos[codon,aa] = (p.item(),r[:,0,p].argmax(axis = 0).item())

        df = pd.melt(pd.Series(res).unstack().reset_index(),id_vars = ['index']).sort_values('value',ascending = False)
        ranks = []
        for codon,aa in zip(df['index'].iloc[:20], df['variable'].iloc[:20]):
            a, f = pos[codon,aa]
            if f>2:
                pp = 3*((y.shape[-1])-a)
            else:
                pp = 3*a
            ranks.append(pp)
        ranks = np.array(ranks)    
        c = np.where(np.abs(ranks//100-28237)<12)[0] #correct position
        if len(c)==0:
            s = -1
        else:
            s = np.min(c)
        detection[length,position] = s

# %% [code]
results = pd.Series(detection).reset_index()
results[0][results[0]==-1] = 10
results['found'] = results[0]<=5
results[0][~results['found']] = 6

# %% [code]
results.groupby(['level_0'])['found'].mean().plot()
plt.xlabel('Length of recA AA substring')
plt.ylabel('Fraction sufficent to locate')

# %% [code]
```
