## Supplementary material for "Deriving genetic codes for molecular phenotypes from first principles": Code: Fig2_evaluating_Allen_codes.ipynb

```
# %% [code]
import pandas as pd
import numpy as np
import seaborn as sns
from scipy.spatial.distance import cdist
import os
from numpy.lib.stride_tricks import sliding_window_view
from matplotlib import pyplot as plt

# %% [code]
def get_dataframe(dir):
    x2 = pd.read_csv('{}/MicroarrayExpression.csv'.format(dir),header=None,index_col = 0).T
    var2 = pd.read_csv('{}/Probes.csv'.format(dir)).set_index('probe_id')
    obs2 = pd.read_csv('{}/SampleAnnot.csv'.format(dir))
    region_exp = x2.groupby(obs2['structure_name']).mean()
    
    distance_matrix2 = cdist(region_exp,region_exp,metric = 'cosine')
    distance_matrix2/=distance_matrix2.max()
    distance_matrix2 = pd.DataFrame(distance_matrix2, index = region_exp.index)
    return region_exp, distance_matrix2
#read in atlas for all samples
dirs = [a for a in os.listdir('./') if 'h0' in a and '.z' not in a]
region_exps = {}
dists  = {}
for dir in dirs:
    region_exps[dir], dists[dir] = get_dataframe(dir)
    


# %% [markdown]
### GWAS hit enrichment

# %% [code]
gwas3 = pd.read_table('gwas/ukb_roi_volume_may12_2019_phase1and2_pheno81_allchr_withA2.txt',delim_whitespace=True)
gwas3_chrom_groups = gwas3.groupby('CHR').groups
gwas = pd.read_table('gwas/ukb_roi_volume_may12_2019_phase1and2_pheno79_allchr_withA2.txt',delim_whitespace=True)
gwas_chrom_groups = gwas.groupby('CHR').groups
enh_pos = pd.read_pickle('../enh_pos')
gwas2 = pd.read_table('aseg_vol_most.sumstats',delim_whitespace=True)
gwas2_chrom_groups = gwas2.groupby('CHR').groups


# %% [code]

from liftover import get_lifter
converter = get_lifter('hg38', 'hg19')
chrom = '1'
pos = 103786442
converter[chrom][pos]

# %% [code]
lp_ranks = np.load('left_putamen.npy').max(axis = -1).min(axis=0)
lca_ranks = np.load('left_ca1.npy').max(axis = -1).min(axis=0)
llh_ranks = np.load('left_lat_hypoth.npy').max(axis = -1).min(axis=0)
# lp_ranks = np.load('left_putamen_300.npy').max(axis = -1).min(axis=0)
# lca_ranks = np.load('left_ca1_300.npy').max(axis = -1).min(axis=0)
# llh_ranks = np.load('left_lat_hypoth_300.npy').max(axis = -1).min(axis=0)


enh_pos['lp_score'] = lp_ranks
enh_pos['lca_score'] = lca_ranks
enh_pos['llh_score'] = llh_ranks
out = enh_pos.sort_values('lp_score').copy()
auto = out[(out[0]!='chrX') & (out[0]!='chrY')]
top_hits = auto.copy()

hg19_chroms = []
hg19_positions = []
mapped_inds = []
for ind, chrom, pos in zip(top_hits.index, top_hits[0], top_hits['mean']):
    ch = converter[chrom][pos]
    if len(ch)==1:
        if ch[0][0] not in ['chr{}'.format(i) for i in range(1,23)]:
            continue
        hg19_chrom, hg19_pos, _  = ch[0]
        hg19_chroms.append(hg19_chrom)
        hg19_positions.append(hg19_pos)
        mapped_inds.append(ind)

top_hits['hg19_chrom'] = "NA"
top_hits['hg19_pos'] = -1

top_hits['hg19_chrom'].loc[mapped_inds] = hg19_chroms
top_hits['hg19_pos'].loc[mapped_inds] = hg19_positions

top_hits = top_hits[top_hits['hg19_pos']!=-1]


oo = []
for chm, inds in top_hits.groupby(0).groups.items():
    ch = int(chm.split('chr')[-1])
    chr_stats = gwas2.loc[gwas2_chrom_groups[ch]].sort_values("BP")
    
    hitpos = top_hits.loc[inds]['hg19_pos'].sort_values()
    hitidx = hitpos.index.values
    sliding = sliding_window_view(chr_stats['PVAL'],4)
    idx = (np.searchsorted(chr_stats['BP'],hitpos)-2)
    nonzero = np.where((idx>=0) & (idx< sliding.shape[0]))
    hitidx2 = hitidx[nonzero]
    sliding_min = sliding.min(axis = 1)
    minp = sliding_min[idx[nonzero]]
    ser = pd.Series(minp, index = hitidx2)

    chr_stats_puta = gwas.loc[gwas_chrom_groups[ch]].sort_values("BP")
    sliding = sliding_window_view(chr_stats_puta['P'],4)
    idx = (np.searchsorted(chr_stats_puta['BP'],hitpos)-2)
    nonzero = np.where((idx>=0) & (idx< sliding.shape[0]))
    hitidx2 = hitidx[nonzero]
    sliding_min = sliding.min(axis = 1)
    puta_minp = sliding_min[idx[nonzero]]
    puta_ser =pd.Series(puta_minp,index = hitidx2)

    chr_stats_puta_effect = gwas.loc[gwas_chrom_groups[ch]].sort_values("BP")
    sliding = sliding_window_view(chr_stats_puta['BETA'],4)
    idx = (np.searchsorted(chr_stats_puta['BP'],hitpos)-2)
    nonzero = np.where((idx>=0) & (idx< sliding.shape[0]))
    hitidx2 = hitidx[nonzero]
    sliding_min = sliding.mean(axis = 1)
    puta_beta_minp = sliding_min[idx[nonzero]]
    puta_beta_ser =pd.Series(puta_beta_minp,index = hitidx2)

    chr_stats_hip = gwas3.loc[gwas3_chrom_groups[ch]].sort_values("BP")
    sliding = sliding_window_view(chr_stats_hip['P'],4)
    idx = (np.searchsorted(chr_stats_hip['BP'],hitpos)-2)
    nonzero = np.where((idx>=0) & (idx< sliding.shape[0]))
    hitidx2 = hitidx[nonzero]
    sliding_min = sliding.min(axis = 1)
    hip_minp = sliding_min[idx[nonzero]]
    hip_ser =pd.Series(hip_minp,index = hitidx2)

    o = pd.concat([top_hits,ser,puta_ser,puta_beta_ser],axis=1).dropna()
    
    oo.append(pd.concat([o, hip_ser],axis = 1).dropna())
    
res = pd.concat(oo).sort_values('lp_score')
res.columns = ['chr','start','stop',    'mean',   'lp_score','lca_score',  'llh_score', 'hg19_chrom',   'hg19_pos',            'minp','puta_minp','puta_beta', 'hip_minp']
res['pval'] = (res['minp']).argsort().argsort()/res.shape[0]
res['puta_pval'] = (res['puta_minp']).argsort().argsort()/res.shape[0]
# res['mean_score']  = left_putamen_results.mean(axis = -1).min(axis=0)[res.index.values]

# %% [code]
plt.figure(figsize = (9,3))
plt.subplot(1,3,1)
plt.scatter(np.log10(res['lp_score']), np.log10(res['lca_score']), s = 0.1, color = 'grey')
plt.xlabel('log10 putamen score')
plt.ylabel('log10 ca score')
plt.subplot(1,3,2)
plt.scatter(np.log10(res['lp_score']), np.log10(res['llh_score']), s = 0.1, color = 'grey')
plt.xlabel('log10 putamen score')
plt.ylabel('log10 hypothalamus score')
plt.subplot(1,3,3)
plt.scatter(np.log10(res['lca_score']), np.log10(res['llh_score']), s = 0.1, color = 'grey')
plt.xlabel('log10 ca score')
plt.ylabel('log10 hypothalamus score')

# %% [code]
from scipy.stats import binomtest
pvs = {}
#res = res_2000.sort_values('lp_score')
for n in range(10,200, 10):
    for thresh in 10.0**(-np.arange(1,15)):
        col = 'minp'
        k = (res[col].iloc[:n]<thresh).sum()
        pvs[n,thresh] = binomtest(k = k, n = n, p = (res[col]<thresh).mean()).pvalue
        # print(n,thresh,k)
sns.heatmap(-np.log10(pd.Series(pvs).unstack(0)),vmin = -np.log10(0.05/3),vmax = 3, cmap = 'Reds')

# %% [markdown]
#### check enrichment relative to different GO categories

# %% [code]
dd = enh_pos
reformat = pd.Series(['{}:{}:{}'.format(a[3:],b,c) for a,b,c in zip(dd[0],dd[1],dd[2])],index = enh_pos.index.values)
enh_pos['reformat'] = ['{}:{}:{}'.format(a[3:],b,c) for a,b,c in zip(dd[0],dd[1],dd[2])]

# %% [code]
converted = pd.read_pickle('gprof_converted')

# %% [code]
converted[converted['name']!='None'].sample(20)

# %% [code]
len(np.unique(converted[converted['name']!='None'].index.values))

# %% [code]
import blitzgsea as blitz
lib = blitz.enrichr.get_library('GO_Biological_Process_2023')
res['reformat'] = enh_pos['reformat'].loc[res.index.values].values.copy()
converted = pd.read_pickle('gprof_converted')
tops = set([a for a in converted.loc[res['reformat'].iloc[:20]]['name'].values if a !='None'])
covered_genesets = [k for k,v in lib.items() if len(set(v).intersection(tops))>0 ]
covered_geneset_frequencies = {k:(res['minp'][res['reformat'].isin(converted.index[converted['name'].isin(lib[k])].values)]<1e-12).mean() for k in covered_genesets}

# %% [code]


# %% [code]


# %% [code]
plt.figure(figsize = (1,5))
go_binomp = pd.Series({k.split('(')[0]: binomtest(n = 20, k= 2, p = v).pvalue for k,v in covered_geneset_frequencies.items()}).sort_values()
sns.stripplot(-np.log10(1e-10+ go_binomp ), s = 3, c = 'grey')
plt.hlines(-np.log10(0.05),-0.1,0.1, colors = 'black')
plt.ylabel('-log10 binomial p value')


# %% [code]
plt.figure(figsize = (8,5))
go_binomp[go_binomp>0.05].plot.bar(color = 'black')

# %% [code]

import torch
from sklearn.cluster import MiniBatchKMeans

# %% [code]
import torch
from tqdm import tqdm
from matplotlib import pyplot as plt
from numpy.lib.stride_tricks import sliding_window_view
torch.cuda.set_device(1)
seq = torch.from_numpy(np.load('../hg38_cre_evs.npy')).float().cuda()[:,1:]
scales = torch.linspace(1,10,21,device = 'cuda')[:,None,None]

def get_match(inds, targv,num_dim=300):
    origr = []
    origr_i = []
    
    
    with torch.no_grad():
        for i in inds:
            di = torch.cdist(seq[i][None][:,:num_dim],seq[:,:num_dim], compute_mode = 'donot_use_mm_for_euclid_dist')
            di_scaled = (di[None]*scales)
            minv, mink = (di_scaled[:,:,:,None]-targv[None,None,None]).abs().min(axis = 2)
            origr.append(minv.cpu().numpy())
            origr_i.append(mink.cpu().numpy())
            
    return origr, origr_i

# %% [markdown]
#### Empirical nulls

# %% [code]
GC: [1,5,10,15,20]
km: [10,100,500,1000,5000]

# %% [code]
### perm test controlling for cluster similarity
cluster_seq_stats = {}
for n_cluster in [10,100,500,1000,5000]:
    mbk = MiniBatchKMeans(n_clusters = n_cluster,random_state = 4)
    mbk.fit(seq.cpu().numpy()[res.index])
    stat = []
    c_counts = pd.Series(mbk.labels_[:20]).value_counts()
    wheres = [np.where(mbk.labels_==i)[0].copy() for i in c_counts.index]
    for perm in tqdm(range(2000)):
        if perm>0:
            for arr in wheres:
                np.random.shuffle(arr)
        stat.append(((res['minp']<1e-10).values[np.concatenate([arr[:cc] for arr,cc in zip(wheres,c_counts.values)])]).sum())
    cluster_seq_stats[n_cluster] = np.array(stat)

# %% [code]
#emp null controlling for GC content
kmer_counts = pd.read_pickle('../hg38_enh_kmer_counts')
gc = pd.DataFrame({'gc':(kmer_counts.loc['G'] + kmer_counts.loc['C'])/(kmer_counts.loc['A'] + kmer_counts.loc['T'] + (kmer_counts.loc['G'] + kmer_counts.loc['C']))}).reset_index()['gc']
gc_bin_stats_topp = {}
sd = gc.values
for nbins in [1,5,10,15,20]:
    counts,bins,_ = plt.hist(sd[res.index[:20]],bins = nbins);
    sel_inds_orig = res.index[:20]
    intervals = []
    for start, stop in zip(bins, bins[1:]):
        intervals.append(np.array(list(set(np.where((sd>=start) & (sd<stop))[0]).intersection(res.index))))
    stats = []
    for perm in tqdm(range(5000)):
        if perm>0:
            for arr in intervals:
                np.random.shuffle(arr)
            
            sel_inds = np.concatenate([binz[:int(count)] for binz,count in zip(intervals,counts)])
        else:
            sel_inds=sel_inds_orig

        stats.append((res['minp'].loc[sel_inds]<1e-10).sum())
    
    
    gc_bin_stats_topp[nbins] = np.array(stats).copy()

# %% [code]
(-np.log10(pd.Series({k : 1-(v[0]>v[1:]).mean() for k,v in gc_bin_stats_topp.items()}))).plot(label = 'GC',color = 'grey',linestyle = '--')

# %% [code]
plt.figure(figsize = (5,10))
(-np.log10(pd.Series({i:(v[1:]>=v[0]).mean() for i,(k,v) in enumerate(cluster_seq_stats.items())}))).plot(label = 'feature',color = 'grey',linestyle = ':',linewidth = 3)
(-np.log10(pd.Series({i:(v[1:]>=v[0]).mean() for i,(k,v) in enumerate(gc_bin_stats_topp.items())}))).plot(label = 'GC',color = 'grey',linestyle = '--',linewidth = 3)
plt.legend()
plt.xticks(range(0,5),range(1,6))
plt.xlabel('bin resolution')
# (-np.log10(pd.Series({k : 1-(v[0]>v[1:]).mean() for k,v in gc_bin_stats_topp.items()}))).plot(label = 'GC',color = 'grey',linestyle = '--')

# %% [code]
sf_bin_stats_topp2 = {}
sd = torch.cdist(seq[res.index[2]][None],seq, compute_mode = 'donot_use_mm_for_euclid_dist').flatten().cpu().numpy()
for nbins in [1,5,10,15,20]:
    counts,bins,_ = plt.hist(sd[res.index[:20]],bins = nbins);
    sel_inds_orig = res.index[:20]
    intervals = []
    for start, stop in zip(bins, bins[1:]):
        intervals.append(np.array(list(set(np.where((sd>=start) & (sd<stop))[0]).intersection(res.index))))
    stats = []
    for perm in tqdm(range(5000)):
        if perm>0:
            for arr in intervals:
                np.random.shuffle(arr)
            
            sel_inds = np.concatenate([binz[:int(count)] for binz,count in zip(intervals,counts)])
        else:
            sel_inds=sel_inds_orig

        stats.append((res['minp'].loc[sel_inds]<1e-10).sum())
    
    
    sf_bin_stats_topp2[nbins] = np.array(stats).copy()

# %% [code]
sf_bin_stats_topp1 = {}
sd = torch.cdist(seq[res.index[1]][None],seq, compute_mode = 'donot_use_mm_for_euclid_dist').flatten().cpu().numpy()
for nbins in [1,5,10,15,20]:
    counts,bins,_ = plt.hist(sd[res.index[:20]],bins = nbins);
    sel_inds_orig = res.index[:20]
    intervals = []
    for start, stop in zip(bins, bins[1:]):
        intervals.append(np.array(list(set(np.where((sd>=start) & (sd<stop))[0]).intersection(res.index))))
    stats = []
    for perm in tqdm(range(5000)):
        if perm>0:
            for arr in intervals:
                np.random.shuffle(arr)
            
            sel_inds = np.concatenate([binz[:int(count)] for binz,count in zip(intervals,counts)])
        else:
            sel_inds=sel_inds_orig

        stats.append((res['minp'].loc[sel_inds]<1e-10).sum())
    
    
    sf_bin_stats_topp1[nbins] = np.array(stats).copy()

# %% [code]
sf_bin_stats_topp = {}
sd = torch.cdist(seq[res.index[0]][None],seq, compute_mode = 'donot_use_mm_for_euclid_dist').flatten().cpu().numpy()
for nbins in [1,5,10,15,20]:
    counts,bins,_ = plt.hist(sd[res.index[:20]],bins = nbins);
    sel_inds_orig = res.index[:20]
    intervals = []
    for start, stop in zip(bins, bins[1:]):
        intervals.append(np.array(list(set(np.where((sd>=start) & (sd<stop))[0]).intersection(res.index))))
    stats = []
    for perm in tqdm(range(5000)):
        if perm>0:
            for arr in intervals:
                np.random.shuffle(arr)
            
            sel_inds = np.concatenate([binz[:int(count)] for binz,count in zip(intervals,counts)])
        else:
            sel_inds=sel_inds_orig

        stats.append((res['minp'].loc[sel_inds]<1e-10).sum())
    
    
    sf_bin_stats_topp[nbins] = np.array(stats).copy()

# %% [code]


# %% [code]


# %% [code]
(-np.log10(pd.Series({k : 1-(v[0]>v[1:]).mean() for k,v in sf_bin_stats_topp.items()}))).plot(label = 'features',linestyle = ':', color = 'grey')
(-np.log10(pd.Series({k : 1-(v[0]>v[1:]).mean() for k,v in sf_bin_stats_topp1.items()}))).plot(label = 'features',linestyle = ':', color = 'grey')
(-np.log10(pd.Series({k : 1-(v[0]>v[1:]).mean() for k,v in sf_bin_stats_topp2.items()}))).plot(label = 'features',linestyle = ':', color = 'grey')
plt.hlines(1.3, 0, 20)

# %% [markdown]
### Alignment loss per putamen locus correlation across individuals

# %% [code]


# %% [code]

from tqdm.notebook import tqdm
import torch

# %% [code]

import torch
from tqdm import tqdm
from matplotlib import pyplot as plt
from numpy.lib.stride_tricks import sliding_window_view
torch.cuda.set_device(1)
seq = torch.from_numpy(np.load('../hg38_cre_evs.npy')).float().cuda()[:,1:]
scales = torch.linspace(1,10,21,device = 'cuda')[:,None,None]

def get_match(inds, targv,num_dim=300):
    origr = []
    origr_i = []
    
    
    with torch.no_grad():
        for i in inds:
            di = torch.cdist(seq[i][None][:,:num_dim],seq[:,:num_dim], compute_mode = 'donot_use_mm_for_euclid_dist')
            di_scaled = (di[None]*scales)
            minv, mink = (di_scaled[:,:,:,None]-targv[None,None,None]).abs().min(axis = 2)
            origr.append(minv.cpu().numpy())
            origr_i.append(mink.cpu().numpy())
            
    return origr, origr_i

# %% [code]
def get(inds, targv,num_dim=300):
    origr = []
    origr_i = []
    
    
    with torch.no_grad():
        for i in inds:
            di = torch.cdist(seq[i][None][:,:num_dim],seq[:,:num_dim], compute_mode = 'donot_use_mm_for_euclid_dist')
            di_scaled = (di[None]*scales)
            minv, mink = (di_scaled[:,:,:,None]-targv[None,None,None]).abs().min(axis = 2)
            origr.append(minv.cpu().numpy())
            origr_i.append(mink.cpu().numpy())
            
    return origr, origr_i


topinds = []
cols = ['llh_score','lca_score', 'lp_score']
topindcols = []
for col in cols:
    topinds = topinds+res.sort_values(col).iloc[:20].index.values.tolist()
    topindcols += [col]*20
    
inds = topinds
calc = {}
for samp, distance_matrix in dists.items():
    if '200' not in samp:
        continue
    for region in tqdm(distance_matrix.index):
        if '2001' in samp:
            if ('CA1 field, left' not in region) & ('lateral hypothalamic area, mammillary region, left' not in region) & ('putamen, left' not in region):
                continue
        else:
            if ('CA1 field' not in region) & ('lateral hypothalamic area, mammillary region' not in region) & ('putamen' not in region):
                continue
        val_targ,val_targ_i = get(inds, torch.from_numpy(distance_matrix.loc[region].values).float().cuda(),num_dim= 2000)
        calc[samp, region] = np.concatenate(val_targ,1).mean(axis = -1).min(axis = 0)


# %% [code]
from scipy.stats import pearsonr,spearmanr

# %% [code]
c2 = {}
for k,v in calc.items():
    samp, targ = k
    c2[samp,targ,cols[0]] = v[:20]
    c2[samp,targ,cols[1]] = v[20:40]
    c2[samp,targ,cols[2]] = v[-20:]

cors = {}
for k,v in tqdm(c2.items()):
    if '2001' not in k[0]:
        continue
    for k2,v2 in c2.items():
        cors[k,k2] = spearmanr(v,v2)

# %% [code]
comb_cor = pd.DataFrame(cors).T

# comb_cor.to_pickle('hit_correlations')

# %% [code]
comb_cor = pd.read_pickle('hit_correlations')

# %% [code]
sub = []
for i,(a,b) in enumerate(comb_cor.index):
    if '2001' in b[0]:
        continue
    if ('CA1' in a[1]) & ('lca' not in a[2]):
        continue
    if ('hypo' in a[1]) & ('llh' not in a[2]):
        continue

    if ('putamen' in a[1]) & ('lp' not in a[2]):
        continue

    if a[2] != b[2]:
        continue
    sub.append(i)

# %% [code]
sub = comb_cor.iloc[sub]

# %% [code]
sub[(sub[1]<0.01) & (sub[0]>0)]

# %% [code]

plt.figure(figsize = (5,5))
plt.scatter(c2[('h0351-2001',
  'putamen, left',
  'lp_score')],c2[('h0351-2002',
  'putamen, right','lp_score')],c = 'black')
plt.xlabel('Patient 1, left putamen alignment loss')
plt.ylabel('Patient 2, right putamen alignment loss')

# %% [code]

cor_df_mean = pd.read_pickle('results/mean_cors_tophits')
a1 = cor_df_mean.loc[('h0351-2001','lateral hypothalamic area, mammillary region, left', 'llh_score')].filter(regex = '2002',axis = 0).filter(regex = 'llh_score',axis=0)[0]
a2 = cor_df_mean.loc[('h0351-2001','putamen, left', 'lp_score')].filter(regex = '2002',axis = 0).filter(regex = 'lp_score',axis=0)[0]
a3 = cor_df_mean.loc[('h0351-2001','CA1 field, left', 'lca_score')].filter(regex = '2002',axis = 0).filter(regex = 'lca_score',axis=0)[0]
plt.figure(figsize = (6,5))
sns.stripplot(data = pd.concat({'hypothalamus':a1,'putamen':a2, 'hippocampus':a3}).reset_index(),
               x = 'level_0', y = 0,c = 'black')

for i, a in enumerate([a1,a2,a3]):
    for name,j in zip(a.sort_values().index[-5:], a.sort_values().iloc[-5:]):
        if j>0.5:
            plt.text(i-0.05,j,name[1], fontsize = 5,ha = 'right')

plt.ylabel('alignment correlation')
plt.xlabel('input region')

# %% [code]
plt.figure(figsize = (5,5))
plt.scatter(c2[('h0351-2001',
  'putamen, left',
  'lp_score')],c2[('h0351-2002',
  'putamen, right','lp_score')],c = 'black')
plt.xlabel('Patient 1, left putamen alignment loss')
plt.ylabel('Patient 2, right putamen alignment loss')

# %% [code]
cor_df_mean = pd.DataFrame(cors).T.sort_values(0)

# %% [code]
plt.figure(figsize = (15,2))
k = 0
for sc in ['llh_score','lca_score','lp_score']:
    k+=1
    plot_df = cor_df_mean[0].unstack()
    plot_df = plot_df [ [a for a in plot_df.columns if ('2002' in a[0]) and (a[2] == sc)]]
    plot_df = plot_df.iloc[[i for i,a in enumerate(plot_df.index) if a[2]==sc]]
    plot_df2 = cor_df_mean[1].unstack()
    plot_df2 = plot_df2 [ [a for a in plot_df.columns if ('2002' in a[0]) and (a[2] == sc)]]
    plot_df2  = plot_df2.iloc[[i for i,a in enumerate(plot_df2.index) if a[2]==sc]]
    plt.subplot(1,3,k)
    sns.heatmap(plot_df*(plot_df2<0.01) , cmap = 'bwr',vmin = -1, vmax = 1,xticklabels = True,yticklabels = False)

# %% [code]
res_2000['lmp'] = -np.log10(res_2000['minp'])
res_2000['lpp'] = -np.log10(res_2000['puta_minp'])


# %% [code]
-np.log10(0.05/(4*20))

# %% [code]
plt.figure(figsize = (5,5))
sns.scatterplot(data = res_2000.iloc[:20], x = 'lmp', y = 'lpp',c = 'black')
plt.xlabel('-log10 gwas p-value, brain structure')
plt.ylabel('-log10 gwas p-value, putamen volume')


# %% [code]

res

# %% [code]
res['lmp'] = -np.log10(res['minp'])
plt.figure(figsize = (3,5))
sns.scatterplot(data = res, x = 'lp_score', y = 'lmp',c = 'black')
plt.hlines(-np.log10(0.05/(4*20)),0, 1e-5 ,colors = 'black',linewidth = 0.5)
plt.xlim(5e-6, 6e-6)
plt.xlabel('alignment loss')
plt.ylabel('-log10 gwas p-value, brain structure')
# plt.ylim(0,25)


# %% [markdown]
### Evaluating regional gene expression in mapped enhancers

# %% [code]
import torch
from tqdm import tqdm
from matplotlib import pyplot as plt
from numpy.lib.stride_tricks import sliding_window_view
torch.cuda.set_device(1)
seq = torch.from_numpy(np.load('../hg38_cre_evs.npy')).float().cuda()[:,1:]
scales = torch.linspace(1,10,21,device = 'cuda')[:,None,None]

def get_match(inds, targv,num_dim=300):
    origr = []
    origr_i = []
    
    
    with torch.no_grad():
        for i in inds:
            di = torch.cdist(seq[i][None][:,:num_dim],seq[:,:num_dim], compute_mode = 'donot_use_mm_for_euclid_dist')
            di_scaled = (di[None]*scales)
            minv, mink = (di_scaled[:,:,:,None]-targv[None,None,None]).abs().min(axis = 2)
            origr.append(minv.cpu().numpy())
            origr_i.append(mink.cpu().numpy())
            
    return origr, origr_i

# %% [code]
lp = np.load('left_putamen.npy')


# %% [code]

origr,origr_i = get_match(res.sort_values('lp_score').iloc[:20].index.values,torch.from_numpy(dists['h0351-2001'].loc['putamen, left'].values).float().cuda(), num_dim = 2000 )
top_inds = res.sort_values('lp_score').index[:20]
top_scales = lp[:,top_inds].max(axis = -1).argmin(axis = 0)

dat = {i:pd.DataFrame({'loss':pd.Series(lp[top_scales[i]][top_inds[i]],index = dists['h0351-2001'].index).sort_values(),
 'ind':pd.Series(origr_i[i][top_scales[i],0],index = dists['h0351-2001'].index), 'd2p': pd.Series(dists['h0351-2001'].loc['putamen, left'].values, index = dists['h0351-2001'].index)}).sort_values('loss') for i in range(20)}

ddat = pd.concat(dat)

# %% [code]
for k,v in dat.items():
    v['pval'] = res.reindex(v['ind'])['pval'].values
    v['puta_pval'] = res.reindex(v['ind'])['puta_pval'].values
    # v = v.dropna()

# %% [code]

dd = enh_pos
reformat = pd.Series(['{}:{}:{}'.format(a[3:],b,c) for a,b,c in zip(dd[0],dd[1],dd[2])],index = enh_pos.index.values)
enh_pos['reformat'] = ['{}:{}:{}'.format(a[3:],b,c) for a,b,c in zip(dd[0],dd[1],dd[2])]
ddat['reformat'] = enh_pos['reformat'].loc[ddat['ind'].values].values

# %% [code]
# from gprofiler import GProfiler

# prof = GProfiler(return_dataframe=True)
# prof.profile(organism='hsapiens',
#             query=['NR1H4','TRIP12','UBC','FCRL3','PLXNA3','GDNF','VPS11'],sources=['GO:BP'])

# convs = []
# for idx in tqdm(np.array_split(np.arange(len(reformat)), 30)):
#     convs.append(prof.convert(organism='hsapiens',
#                 query=reformat.iloc[idx].values.tolist(),
#                 target_namespace='ENTREZGENE_ACC'))

# converted = pd.concat(convs).set_index('incoming')
# converted.to_pickle('gprof_converted')

# %% [code]
converted = pd.read_pickle('gprof_converted')

# %% [code]
def znorm(df):
    a = df.values
    return pd.DataFrame((a-a.mean(axis = 0,keepdims=True))/a.std(axis=0,keepdims=True), index = df.index, columns = df.columns)

re = region_exps['h0351-2001'].copy()
re.columns = pd.read_csv('h0351-2001/Probes.csv').set_index('probe_id')['gene_symbol'].loc[re.columns]
common_genes = sorted(set(re.columns).intersection(converted['name']))
re_avg = re[common_genes].melt(ignore_index = False).reset_index().groupby(['structure_name','gene_symbol']).mean()['value'].unstack()
zavg = znorm(re_avg)
p2e = pd.concat({k: converted['name'].map(zavg.loc[k]).dropna() for k in zavg.index}).reset_index()
p2e = p2e.groupby(['incoming','level_0'])['name'].mean()
p2e = p2e.unstack()

# %% [code]
hit = 2
gg2 = ddat.loc[hit].reset_index().set_index('reformat')['structure_name']
top_ind = ddat.loc[2].sort_values('d2p')['ind'].iloc[0]
sd = torch.cdist(seq[top_ind][None],seq, compute_mode = 'donot_use_mm_for_euclid_dist').flatten().cpu().numpy()
sel_inds_orig = ddat.loc[hit]['ind'].values
# gg2 = pd.DataFrame({'structure_name':gg2.values.flatten()}, index = sel_reformat)
        
comb = pd.merge(gg2,p2e,left_index = True, right_index=True).set_index('structure_name')
comb_orig = comb[comb.index]


# %% [code]
pd.merge(gg2,p2e,left_index = True, right_index=True) # so rows are ASSIGNED enhancer

# %% [code]
f = sns.clustermap(comb_orig,figsize = (10,10),cmap = 'bwr',vmin = -2,vmax = 2)

# %% [code]
plt.figure(figsize = (1.5,5))
plt.hist(comb_orig.values.flatten(),density=True,bins = 10,alpha = 0.5,orientation = 'horizontal',color = 'grey')
plt.hist(np.diag(comb_orig),density = True,bins = 10,alpha = 0.5,orientation = 'horizontal',color = 'red')
plt.xlabel('density')
plt.ylabel('z-score')


# %% [code]


# %% [code]
A = comb_orig.iloc[f.dendrogram_row.reordered_ind,:].iloc[:,f.dendrogram_row.reordered_ind].values
rowc = np.diag(A) - A.mean(axis=1)
colc = np.diag(A) - A.mean(axis=0)
plt.scatter(rowc,colc)
plt.show()

rowc = (rowc+2.5)/5
colc = (colc+2.5)/5
pal = np.array(sns.color_palette('bwr',21))
col_colors = pal[(20*colc).astype(int)]
row_colors = pal[(20*rowc).astype(int)]
sns.clustermap(comb_orig.iloc[f.dendrogram_row.reordered_ind,:].iloc[:,f.dendrogram_row.reordered_ind], cmap = 'bwr',vmin = -2, vmax = 2,row_cluster = False, col_cluster = False,yticklabels = False,figsize = (10,10),xticklabels=False,
              col_colors = col_colors, row_colors = row_colors)


# %% [code]
stat = []
orig = np.arange(len(A))
shuf = orig.copy()
for perm in range(20):
    if perm>0:
        np.random.shuffle(shuf)
    An = A[:,shuf]
    q = np.diag(A)-A.mean(axis = 1)
    stat.append(np.median(q))

stat = np.array(stat)


# %% [code]
stat

# %% [code]
mask = ~np.eye(comb_orig.shape[0], dtype=bool)
plt.figure(figsize = (10,10))
from matplotlib.patches import Rectangle
# sns.heatmap(comb_orig,mask = ~mask, cmap = 'bwr',vmin = -2, vmax = 2,xticklabels=False, yticklabels = False,alpha = 0.3,cbar = None)
ax =sns.heatmap(comb_orig.iloc[f.dendrogram_row.reordered_ind,:].iloc[:,f.dendrogram_row.reordered_ind], cmap = 'bwr',vmin = -2, vmax = 2,xticklabels=False, yticklabels = False,cbar = None)
# for i in range(len(comb_orig)):
#     ax.add_patch(Rectangle((i, i), 
#                           width=1, 
#                           height=1, 
#                           fill=False,    # No fill, only border
#                           edgecolor='black',
#                           linewidth=1))

# %% [code]
A = comb_orig.iloc[f.dendrogram_row.reordered_ind,:].iloc[:,f.dendrogram_row.reordered_ind].values
plt.figure(figsize = (5,1))
plt.scatter(range(len(A)),[0]*len(A),c = np.diag(A) - A.mean(axis=0),cmap = 'bwr',vmin = -2.5, vmax = 2.5,s = 10)
plt.axis('off')
plt.gcf().patch.set_alpha(0)

# %% [code]

plt.figure(figsize = (1,10))
A = comb_orig.iloc[f.dendrogram_row.reordered_ind,:].iloc[:,f.dendrogram_row.reordered_ind].values
rowc = np.diag(A) - A.mean(axis=1)
colc = np.diag(A) - A.mean(axis=0)

plt.barh(range(len(A)),color = 'grey')
plt.xlim(-2.5,2.5)

# %% [code]
mask = ~np.eye(comb_orig.shape[0], dtype=bool)
plt.figure(figsize = (10,10))
from matplotlib.patches import Rectangle
# sns.heatmap(comb_orig,mask = ~mask, cmap = 'bwr',vmin = -2, vmax = 2,xticklabels=False, yticklabels = False,alpha = 0.3,cbar = None)
ax =sns.heatmap(comb_orig[f.dendrogram_col.reordered_ind,:][:,f.dendrogram_col.reordered_ind], cmap = 'bwr',vmin = -2, vmax = 2,xticklabels=False, yticklabels = False,cbar = None)
# for i in range(len(comb_orig)):
#     ax.add_patch(Rectangle((i, i), 
#                           width=1, 
#                           height=1, 
#                           fill=False,    # No fill, only border
#                           edgecolor='black',
#                           linewidth=1))

# %% [code]
f = sns.clustermap(comb, cmap = 'bwr',vmin = -3, vmax = 3,xticklabels=False, yticklabels = False,figsize = (0.1,0.1))


# %% [code]
comb_orig.shape

# %% [code]


# %% [code]


# %% [code]
hit = 2
gg2 = ddat.loc[hit].reset_index().set_index('reformat')['structure_name']
top_ind = ddat.loc[2].sort_values('d2p')['ind'].iloc[0]
sd = torch.cdist(seq[top_ind][None],seq, compute_mode = 'donot_use_mm_for_euclid_dist').flatten().cpu().numpy()
sel_inds_orig = ddat.loc[hit]['ind'].values
# gg2 = pd.DataFrame({'structure_name':gg2.values.flatten()}, index = sel_reformat)
        
comb = pd.merge(gg2,p2e,left_index = True, right_index=True).set_index('structure_name')
comb_orig = comb[comb.index].values
bin_stats = {}
for nbins in [1,5,10,50]:
    counts,bins,_ = plt.hist(sd[ddat.loc[hit]['ind']],bins = nbins);
    stats = []
    intervals = []
    for start, stop in zip(bins, bins[1:]):
        intervals.append(np.array(list(set(np.where((sd>=start) & (sd<stop))[0]))))
    for perm in tqdm(range(2000)):
        if perm>0:
            for arr in intervals:
                np.random.shuffle(arr)
            
            sel_inds = np.concatenate([binz[:int(count)] for binz,count in zip(intervals,counts)])
            if len(sel_inds)<327:
                continue
        else:
            sel_inds = sel_inds_orig.copy()
        sel_reformat= enh_pos['reformat'].loc[sel_inds]
        gg2 = pd.DataFrame({'structure_name':gg2.values.flatten()}, index = sel_reformat)
        
        comb = pd.merge(gg2,p2e,left_index = True, right_index=True).set_index('structure_name')
        comb = comb[comb.index].values
        stats.append(np.median(np.diag(comb)))    
    
    
    bin_stats[nbins] = np.array(stats).copy()

# %% [code]
print({k : 1-(v[0]>v[1:]).mean() for k,v in bin_stats.items()})
plt.figure(figsize = (2,5))
sns.stripplot(data = pd.DataFrame(bin_stats).iloc[1:].melt(),x = 'variable',y='value',c = 'grey',s = 2)
plt.hlines(bin_stats[1][0],-.1,3.1,colors = 'red')
plt.xlabel('bin width')
plt.ylabel('median diagonal expression')

# %% [code]
hit = 2
gg2 = ddat.loc[hit].reset_index().set_index('reformat')['structure_name']
top_ind = ddat.loc[2].sort_values('d2p')['ind'].iloc[0]
stats = []
sel_inds = ddat.loc[hit]['ind'].values
for perm in tqdm(range(2000)):
    if perm>0:
        sel_inds = enh_pos.sample(len(sel_inds),replace=False).index.values.copy()
    sel_reformat= enh_pos['reformat'].loc[sel_inds]
    gg2 = pd.DataFrame({'structure_name':gg2.values.flatten()}, index = sel_reformat)
    
    comb = pd.merge(gg2,p2e,left_index = True, right_index=True).set_index('structure_name')
    comb = comb[comb.index].values
    stats.append(np.median(np.diag(comb)))    


stats = np.array(stats)

# %% [code]


# %% [markdown]
### Evaluating region specific volume gwas effect sizes

# %% [code]
pheno_dict = {}
for a in pd.read_table('gwas/zhao_101_regions.txt',header=None)[0]:
    phenocat = a.split('_')
    pheno = phenocat[0]
    pheno_dict['_'.join(phenocat[1:])] = int(pheno[5:])

lefts= set()
rights = set()
cols = region_exps['h0351-2001'].index
for i in cols:
    if 'left' in i:
        lefts.add(i)
    if 'right' in i:
        rights.add(i)
sim = {}
for i in pheno_dict:
    if 'left' in i:
        oth = lefts
    if 'right' in i:
        oth = rights
    for j in oth:
        
        sim[i,j]  = 0
        for w in i.split('.'):
            sim[i,j] = sim[i,j]+ (w.upper() in j.upper())
sim = pd.Series(sim)        

n2gmap =pd.read_table('nat_gen2allen.txt',header=None).set_index(1)[0].to_dict()

sim = {}
lens = {}
for name in cols:
    for key in n2gmap.keys():
        sim[name,key]=0
        lens[name,key] = 0
        for i in name.split(' '):
            for j in i.split(','):
                lens[name,key] = lens[name,key]+1
                if j in key:
                    sim[name,key] = sim[name,key]+ 1
sim =pd.Series(sim)    
lens = pd.Series(lens)
mapkey = {}
for k1, k2 in sim[sim==lens].index:
    if k1 not in ['V','VI']:
        mapkey[k1] = k2

mapkey['V'] = 'lobules I-V, cerebellum'
mapkey['VI'] = 'VI, paravermis/VIIAf/At'

def map_regions(cols):
    return cols.map(mapkey).map(n2gmap)

# %% [code]
gwas = pd.read_pickle('brain_regions_gwas')

# %% [code]
gwas_pos = gwas[['CHR','BP']].reset_index().rename({'CHR':'Chromosome','BP':'Start'},axis=1)
gwas_pos['End'] = gwas_pos['Start']+1

# %% [code]
import pyranges as pr
gwas_pos_bed = pr.PyRanges(gwas_pos)

# %% [code]
pheno_dict_inv = {}
for i,j in pheno_dict.items():
    pheno_dict_inv[j] = i

# %% [code]
lift_pos = top_hits.reset_index()[['index','hg19_chrom','hg19_pos']].rename({'hg19_chrom':'Chromosome'},axis=1)
lift_pos['Chromosome'] = [a[3:] for a in lift_pos['Chromosome']]

lift_pos = lift_pos[lift_pos['Chromosome'].isin(np.arange(1,23).astype(str))]
lift_pos['Chromosome'] = lift_pos['Chromosome'].astype(int)
lift_pos['Start'] = lift_pos['hg19_pos']-1000
lift_pos['End'] = lift_pos['hg19_pos']+1000
lift_pos_bed = pr.PyRanges(lift_pos[['Chromosome','Start','End','index']])

# %% [code]
combined = lift_pos_bed.join(gwas_pos_bed).as_df()

# %% [code]
enh_betas = gwas.iloc[:,2:].iloc[:,0:].iloc[:,::3].loc[combined['index_b'].values]
enh_ps = gwas.iloc[:,2:].iloc[:,2:].iloc[:,::3].loc[combined['index_b'].values]

# %% [code]
threshb = pd.DataFrame((enh_betas.abs() * (enh_ps.values<0.001)), index = enh_betas.index, columns = enh_betas.columns)
threshb['enh_index'] = combined['index'].values
max_beta =  threshb.groupby('enh_index').max()
max_beta.columns = [pheno_dict_inv[int(a[0].split('pheno')[-1])] for a in max_beta.columns]

# %% [code]
sel_inds[isna_gg].shape

# %% [code]
sel_inds[ismapped_gg]

# %% [code]


bin_stats_beta = {}
for nbins in [1,5,10,50]:
    counts,bins,_ = plt.hist(sd[ddat.loc[hit]['ind']],bins = nbins);
    gg = ddat.loc[2].copy()
    gg['gwas_region'] = map_regions(gg.index).values
    ismapped_gg = ~gg['gwas_region'].isna().values.copy()
    gg = gg.dropna()
    stats = []
    for perm in tqdm(range(2000)):
        if perm>0:
            
            intervals = []
            for start, stop in zip(bins, bins[1:]):
                intervals.append(np.array(list(set(np.where((sd>=start) & (sd<stop))[0]))))
        
            for arr in intervals:
                np.random.shuffle(arr)
            
            sel_inds = np.concatenate([binz[:int(count)] for binz,count in zip(intervals,counts)])
            if len(sel_inds)<327:
                continue
            gg['ind'] = sel_inds[ismapped_gg]
        
        
        mgg = pd.merge(gg,max_beta, left_on = 'ind', right_index = True).dropna()
        comb = mgg[mgg['gwas_region'].values].values
        if perm==0:
            beta_comb_orig = mgg[mgg['gwas_region'].values]
        
        
        stats.append(np.mean(np.diag(comb)))
        
    
    
    bin_stats_beta[nbins] = np.array(stats).copy()


# %% [code]
mgg[mgg['gwas_region'].values]

# %% [code]
[i for i in pheno_dict if 'thalam' in i]

# %% [code]
mgg[mgg['gwas_region'].str.contains('proper')]

# %% [code]
plt.figure(figsize = (12,10))
ax =sns.heatmap(beta_comb_orig,xticklabels = True,yticklabels = True)
for i in range(len(beta_comb_orig)):
    if beta_comb_orig.iloc[i,i]>0:
        print(i)
        ax.add_patch(Rectangle((i, i), 
                              width=1, 
                              height=1, 
                              fill=False,    # No fill, only border
                              edgecolor='cyan',
                              linewidth=2))

# %% [code]
{k : 1-(v[0]>v[1:]).mean() for k,v in bin_stats_beta.items()}

# %% [code]
plt.figure(figsize = (2,5))
sns.stripplot(data = pd.DataFrame(bin_stats_beta).iloc[1:].melt(),x = 'variable',y='value',c = 'grey',s = 2)
plt.hlines(bin_stats_beta[1][0],-.1,3.1,colors = 'red')
plt.xlabel('bin width')
plt.ylabel('mean absolute beta')

# %% [code]
enh_betas.iloc[:,:10].std(axis=0)

# %% [code]
for i in range(3):
    gg = ddat.loc[i]
    gg['gwas_region'] = map_regions(gg.index).values
    mgg = pd.merge(gg,max_beta, left_on = 'ind', right_index = True).dropna()
    comb = mgg[mgg['gwas_region'].values].values
    comb = comb/(1e-5+comb.max(axis=0))
    orig = np.arange(comb.shape[0])
    shuf = np.arange(comb.shape[1])
    stats = []
    for perm in range(1000):
        if perm >0:
            np.random.shuffle(shuf)
        stats.append(np.mean(np.diag(comb[orig,shuf])))
    stats = np.array(stats)
    print((stats[0]>stats[1:]).mean())

# %% [code]
(stats[0]>stats[1:]).mean()
```
