## Supplementary material for "Deriving genetic codes for molecular phenotypes from first principles": Code: Fig2_finding_Allen_codes.ipynb

```
# %% [code]
import pandas as pd
import numpy as np
import seaborn as sns
from scipy.spatial.distance import cdist

# %% [code]
x = pd.read_csv('MicroarrayExpression.csv',header=None,index_col = 0).T
var = pd.read_csv('Probes.csv').set_index('probe_id')
obs = pd.read_csv('SampleAnnot.csv')
region_exp = x.groupby(obs['structure_name']).mean()
distance_matrix = cdist(region_exp,region_exp,metric = 'cosine')
distance_matrix/=distance_matrix.max()
distance_matrix = pd.DataFrame(distance_matrix, index = region_exp.index)
#this is for sample 2001

# %% [code]
import torch
from tqdm import tqdm
from matplotlib import pyplot as plt
torch.cuda.set_device(3)

# %% [code]
seq = torch.from_numpy(np.load('../hg38_cre_evs.npy')).float().cuda()[:,1:301]


# %% [code]

splits = np.array_split(np.arange(seq.shape[0]),150000)
scales = torch.linspace(1,10,21,device = 'cuda')[:,None,None]
results1 = []
results2 = []
results3 = []
targ1 = torch.from_numpy(distance_matrix.loc['CA1 field, left'].values).float().cuda()
targ2 = torch.from_numpy(distance_matrix.loc['lateral hypothalamic area, mammillary region, left'].values).float().cuda()
targ3 = torch.from_numpy(distance_matrix.loc['putamen, left'].values).float().cuda()
with torch.no_grad():
    for i in tqdm(splits):
        di = torch.cdist(seq[i],seq, compute_mode = 'donot_use_mm_for_euclid_dist')
        di_scaled = (di[None]*scales)
        results1.append((di_scaled[:,:,:,None]-targ1[None,None,None]).abs().amin(axis = 2).cpu().numpy())
        results2.append((di_scaled[:,:,:,None]-targ2[None,None,None]).abs().amin(axis = 2).cpu().numpy())
        results3.append((di_scaled[:,:,:,None]-targ3[None,None,None]).abs().amin(axis = 2).cpu().numpy())
        


# %% [code]
np.save('left_ca1_300',np.concatenate(results1,axis=1))
np.save('left_lat_hypoth_300',np.concatenate(results2,axis=1))
np.save('left_putamen_300',np.concatenate(results3,axis=1))

# %% [code]
```
