## Supplementary material for "Deriving genetic codes for molecular phenotypes from first principles": Code: Fig3_evaluating_TabSap_codes.ipynb

```
# %% [code]
import pandas as pd
import numpy as np
import pyranges as pr
from scipy.spatial.distance import cdist
from scipy.sparse.csgraph import shortest_path
from sklearn.neighbors import kneighbors_graph
from sklearn.cluster import MiniBatchKMeans
import torch
torch.cuda.set_device(3)
# from ot.gromov import entropic_semirelaxed_gromov_wasserstein as gwd
from scipy.stats import fisher_exact, binomtest
import seaborn as sns

# %% [code]
ens = pd.read_pickle('../cre/enh_pos')
# ens.iloc[:,:3].reset_index().iloc[:,[1,2,3,0]].to_csv('enh_pos.bed', sep = '\t',index = None,header=None)
# enh_bed = pr.read_bed('../cre/brain/enh_bed500.bed')
enh_bed = pr.read_bed('enh_pos.bed')
sheets = pd.ExcelFile('41586_2024_7066_MOESM9_ESM.xlsx')
scores = pd.read_excel(sheets,'Young-1_ATAC_result.slope')
scores['chr'] = [a[0] for a in scores.iloc[:,0].str.split('-')]
scores['start'] = [int(a[1]) for a in scores.iloc[:,0].str.split('-')]
scores['end'] = [int(a[2]) for a in scores.iloc[:,0].str.split('-')]

hits = enh_bed.join(pr.PyRanges(scores.rename({'chr':'Chromosome', 'start':'Start', 'end':'End'},axis=1)))
hdf = hits.as_df()
hdf_gp = hdf.groupby('Name')[['Lym','Mye','MK','ME']].apply(lambda w:w.abs().max())

tabsap = pd.read_pickle('../data/tabsap_SEACELL_means')
tsd = shortest_path(kneighbors_graph(tabsap, metric = 'cosine',n_neighbors=20),directed=False)
tsd/=tsd.max()
tinfo = pd.read_pickle('../data/tabsap_SEACell_assignments')

# %% [code]
labels = {}
centers = {}
coup = {}
clusts = {}
loss = {}

for i in range(20):
    labels[i] = np.load('results/mbk{}_labels.npy'.format(i))
    centers[i] = np.load('results/mbk{}_centers.npy'.format(i))
    coup[i] = np.load('results/mbk{}_coupling.npy'.format(i))
    clusts[i] = shortest_path(kneighbors_graph(centers[i],5,mode = 'connectivity'),directed=False)
    clusts[i]/=clusts[i].max()
    assn = coup[i].argmax(axis = 1)
    loss[i] = ((clusts[i][assn,:][:,assn]-tsd)**2).sum(axis =1)
    

# %% [code]
def znorm(df,axis,eps = 1e-10):
    a = df.values
    return pd.DataFrame((a -a.mean(axis =axis,keepdims=True))/(eps + a.std(axis = axis,keepdims=True)),index = df.index,columns = df.columns)

# %% [markdown]
### Enrichment analysis with Chipseq

# %% [code]
chipseq_info = pd.read_table('chipseq.txt')
print('wget ' + ' '.join(chipseq_info['Donwload Link']))
chipseq_info['name'] = chipseq_info['TF'] + '_'+ chipseq_info['Cell Type']

beds = {}
for name,file in zip(chipseq_info['name'], chipseq_info['ID']):
    beds[name] = enh_bed.join(pr.read_bed(file)).as_df()
beds['prime_hsc'] = hdf.copy()

# %% [code]
from scipy.stats import chi2_contingency

# %% [code]
cts = tinfo[6].unique()
ct_mc = {ct:np.where(tabsap.index == tinfo[tinfo[6].str.contains(ct)][11].value_counts().index[0])[0][0] for ct in cts}
ct_mc_assn = {k:tinfo[6][tabsap.index[v]==tinfo[11]].value_counts().iloc[:3] for k,v in ct_mc.items()}
dat ={}
for ct in cts:
    card_endo = np.where(tabsap.index == tinfo[tinfo[6].str.contains(ct)][11].value_counts().index[0])[0][0]
    for i in labels:
        if i in dat:
            continue
        
        assn  = coup[i].argmax(axis = 1)
        labs = labels[i]
        
        
        card_assn= assn[card_endo]
        whichc = np.where(labs==card_assn)[0]
        for name,islet_df in beds.items():
            
            m = len(islet_df['Name'][islet_df['Name'].isin(whichc)].unique())
            
            n = len(islet_df['Name'].unique())
            p = len(enh_bed)-n
            
            # fe = fisher_exact([[m,len(whichc)-m], [n, len(labs)-n]])
            fe = fisher_exact([[m,len(whichc)-m], [n-m, p-len(whichc)]])
            dat[i,ct,name] = ([loss[i][card_endo],card_assn,len(whichc), fe.pvalue,np.isin(whichc, islet_df['Name']).mean()/np.isin(np.arange(ens.shape[0]), islet_df['Name']).mean(),card_endo])


# %% [code]
d = pd.DataFrame(dat).T

# %% [code]
d['lp'] = -np.log10(1e-20+d[3])

# %% [code]
d.loc[12]

# %% [code]
w = d.reorder_levels([1,0,2]).loc['cd4-positive, alpha-beta memory t cell'].reorder_levels([1,0],axis=0).loc['prime_hsc'].sort_values(0)
w = w[w[2]>20]

# %% [code]
from matplotlib import pyplot as plt

# %% [code]
plt.figure(figsize = (6,5))
sns.scatterplot(data = w,x = 0, y = 'lp',c = 'black')
plt.xlabel('PSA-loss, CD4 T cells')
plt.ylabel('Enrichment of immune lineage primed enhancers')

# %% [code]
ct_mc_assn = {k:tinfo[6][tabsap.index[v]==tinfo[11]].value_counts().iloc[:3] for k,v in ct_mc.items()}

# %% [code]
ct_mc_assn['mesenchymal stem cell']

# %% [code]
ct_mc_assn['alveolar fibroblast']

# %% [code]
ct_mc_assn['eye photoreceptor cell']

# %% [code]
ct_mc_assn['corneal keratocyte']

# %% [code]
ct_mc_assn['hillock-club cell of prostate epithelium']

# %% [code]
ct_mc_assn['keratinocyte']

# %% [code]

cg.dendrogram_row.reordered_ind

# %% [code]

import matplotlib.pyplot as plt

# %% [code]


# %% [code]
w = d.reset_index()

# %% [code]
w['fdr'] = fdrcorrection(w[3])[1]

# %% [code]
k = w[(w['fdr']<0.00001) & (w[4]>1)].groupby([5,'level_2']).size().sort_values()
# kk = znorm(k.unstack().fillna(0),axis=0)
# i,j = np.where(kk>3)
# list(zip(kk.index[i],kk.columns[j]))

# %% [code]
counts  = pd.read_pickle('../cre/hg38_enh_kmer_counts')

# %% [code]

ens['gc'] = (counts.loc[['G','C']].sum(axis=0)/counts.loc[['G','C','A','T']].sum(axis=0)).values

# %% [code]
ens

# %% [code]
from tqdm.notebook import tqdm

# %% [markdown]
#### GC corrected sampling

# %% [code]

lab = 12
labs = labels[lab]
all_dat = {}

for clust in tqdm(np.unique(labels[12])):
    whichc = np.where(labs==clust)[0]
    bg_size = 5*len(whichc)

    bins = np.histogram(ens['gc'].loc[whichc],bins=5)
    gc_matched_background =  np.concatenate([np.random.choice(np.where((ens['gc']>bin_start)&(ens['gc']<=bin_end))[0],size =int(bg_size*bin_size/bins[0].sum())) for bin_start,bin_end,bin_size in zip(bins[1],bins[1][1:], bins[0])])
    
    for name,islet_df in beds.items():
        
        m = len(islet_df['Name'][islet_df['Name'].isin(whichc)].unique())
        
        n = len(islet_df['Name'][islet_df['Name'].isin(gc_matched_background)].unique())
        p = len(gc_matched_background)-n
        
        fe = fisher_exact([[m,len(whichc)-m], [n, p]])
        # fe = fisher_exact([[m,len(whichc)-m], [n-m, p-len(whichc)]])
        all_dat[lab,clust,name] = ([len(whichc), fe.pvalue,fe.statistic])


# %% [code]
r = pd.DataFrame(all_dat).T.loc[12].filter(regex = 'TCF7',axis=0)
tcf7_clusts = np.where((r[1]<0.001)&(r[2]>1))[0]

# %% [code]
pd.DataFrame(perm_dat).T.sort_values(2).iloc[-10:]

# %% [code]
def map_color(cs, cmap, n = 10,range = None):
    
    if range == None:
         rat = ((n*(cs-cs.min())/(cs.max()-cs.min())).values).astype(int)
    else:
        rat = (n*np.clip((cs-range[0])/(range[1]-range[0]),0,1)).astype(int)
    pal = sns.color_palette(cmap,n+1)
    
    return np.array(pal)[rat]

# %% [code]
cd4_assns = d.loc[12].filter(regex = 'cd4',axis=0)[1].unique()

# %% [markdown]
### Non GC corrected plots

# %% [code]
gc_content = ens['gc'].groupby(labels[12]).mean()
names_dict = {}
w = d.loc[12].reset_index()
name_inv = {}
for clust in w[1].unique():
    names_dict[clust] = '; '.join(sorted(set(w['level_0'][w[1]==clust])))
    name_inv['; '.join(sorted(set(w['level_0'][w[1]==clust])))] = clust
temp= d[(d[3]<0.01) & (d[4]>1)].loc[12].reset_index()[[1,'level_1','ld']].drop_duplicates()
temp['name'] = temp[1].map(names_dict)
plot_df = temp.set_index(['name','level_1'])['ld'].unstack().fillna(0)
plot_df = plot_df.loc[plot_df.sum(axis=1)>0]
plot_df['gc'] = 10*(plot_df.index.map(name_inv).map(gc_content)-0.4)
cg = sns.clustermap(plot_df, cmap = 'Reds',vmax = 4,figsize = (0.1,0.1))
plt.figure(figsize = (10,10))
# .drop(['SOX9_PP','NKX2-5_Cardiomy.','PDX1_PP'],axis=1)
sns.heatmap(plot_df.iloc[cg.dendrogram_row.reordered_ind,:].iloc[:,cg.dendrogram_col.reordered_ind], cmap = 'Reds',vmax = 4,cbar = False)

# %% [code]
d.filter(regex = 't cell',axis=0).loc[12]

# %% [code]
names_dict = {}
w = d.loc[12].reset_index()
name_inv = {}
for clust in w[1].unique():
    names_dict[clust] = '; '.join(sorted(set(w['level_0'][w[1]==clust])))
    name_inv['; '.join(sorted(set(w['level_0'][w[1]==clust])))] = clust
temp= d[(d[3]<0.01) & (d[4]>1)].loc[12].reset_index()[[1,'level_1','ld']].drop_duplicates()
temp['name'] = temp[1].map(names_dict)
plot_df = temp.set_index(['name','level_1'])['ld'].unstack().fillna(0)
plot_df = plot_df.loc[plot_df.sum(axis=1)>0]
plot_df['gc'] = 10*(plot_df.index.map(name_inv).map(gc_content)-0.4)
cg = sns.clustermap(plot_df, cmap = 'Reds',vmax = 4,figsize = (0.1,0.1))
plt.figure(figsize = (10,10))
# .drop(['SOX9_PP','NKX2-5_Cardiomy.','PDX1_PP'],axis=1)
sns.heatmap(plot_df.iloc[cg.dendrogram_row.reordered_ind,:].iloc[:,cg.dendrogram_col.reordered_ind], cmap = 'Reds',vmax = 4,cbar = False)

# %% [markdown]
### GC Corrected Sampling

# %% [code]

lab = 12
labs = labels[lab]
perm_dat = {}
# np.unique(labels[12])

#temp has the assignments where there is overall enrichment
for clust in tqdm(temp[1].unique()):
    stats = {}
    whichc = np.where(labs==clust)[0]
    bg_size = len(whichc)

    bins = np.histogram(ens['gc'].loc[whichc],bins=10)
    

    for perm in tqdm(range(2500)):
        if perm==0:
            query_set = whichc
        else:
            query_set=  np.concatenate([np.random.choice(np.where((ens['gc']>bin_start)&(ens['gc']<=bin_end))[0],size =int(bg_size*bin_size/bins[0].sum())) for bin_start,bin_end,bin_size in zip(bins[1],bins[1][1:], bins[0])])
        for name,islet_df in beds.items():
            n = np.isin(query_set,islet_df['Name']).sum()
            stats.setdefault(name,[]).append(n)
            
    for name in beds:
        perm_dat[lab,clust,name] = [len(whichc), stats[name][0], (stats[name][0]> np.array(stats[name][1:])).mean(), np.mean(stats[name][1:]),np.max(stats[name][1:])]


# %% [code]
w = pd.DataFrame(perm_dat).T.loc[12].reset_index()
temp = w[(w[2]>0.95) & ((1.5)*w[3]<w[1])]
temp['ld'] = -np.log10(1/5000 + 1-w[2])
temp = temp[['level_0','level_1','ld']].drop_duplicates()
temp['name'] = temp['level_0'].map(names_dict)

plot_df = temp.set_index(['name','level_1'])['ld'].unstack().fillna(0).drop(['SOX9_PP','NKX2-5_Cardiomy.','PDX1_PP'],axis=1)
plot_df = plot_df.loc[plot_df.sum(axis=1)>0].loc[:,plot_df.sum(axis=0)>0]

cg = sns.clustermap(plot_df, cmap = 'Reds',vmax = 2,figsize = (0.1,0.1),metric = 'correlation')
plt.figure(figsize = (10,10))

sns.heatmap(plot_df.iloc[cg.dendrogram_row.reordered_ind,:].iloc[:,cg.dendrogram_col.reordered_ind], cmap = 'Reds',vmax = 3,cbar = False)

# %% [markdown]
### Visualize heatmaps

# %% [code]
g = sns.clustermap(clusts[12], col_colors = [['red' if a in tcf7_clusts else 'black' for a in range(1000)]],figsize = (0.01,0.01))
gmetaclust = fcluster(g.dendrogram_row.linkage, t = 10, criterion = 'maxclust')-1
gcol_pal = np.array(sns.color_palette('tab10',10))
gcols = gcol_pal[gmetaclust]
gcolor_map = {k:v for k,v in enumerate(gcols)}

v = clusts[12].mean(axis=1)
v-=v.mean()
v/=v.std()
v = np.maximum(np.minimum(v,3),-3)
v/=3

# v-=0.5
# v = np.maximum(np.minimum(v,0.25),-0.25)
# v/=0.25
v+=1
v/=2

gdeg_cols = np.array(sns.color_palette('bwr_r',101))[(100*(v)).astype(int)]
# gdeg_cols = np.array(sns.color_palette('bwr',101))[(100*((v-v.min())/(v.max()-v.min()))).astype(int)]
g = sns.clustermap(clusts[12],figsize = (10,10), col_colors = [gcols],row_colors = gdeg_cols,xticklabels=False, yticklabels=False,cmap= 'PuOr')
g.ax_col_dendrogram.set_visible(False)
g.ax_row_dendrogram.set_visible(False)

# %% [code]
f = sns.clustermap(tsd, col_colors = [['red' if a in Tcell_metacells else 'black' for a in range(400)]],figsize = (0.01,0.01))
metaclust = fcluster(f.dendrogram_row.linkage, t = 20, criterion = 'maxclust')-1
col_pal = np.array(sns.color_palette('tab20_r',20))
cols = col_pal[metaclust]
color_map = {k:v for k,v in zip(assn,cols)}
v = tsd.mean(axis=1)
v-=v.mean()
v/=v.std()
v = np.maximum(np.minimum(v,3),-3)
v/=3

# v-=0.5
# v = np.maximum(np.minimum(v,0.25),-0.25)
# v/=0.25
v+=1
v/=2
deg_cols = np.array(sns.color_palette('bwr_r',101))[(100*(v)).astype(int)]
tcell_cols = ['red' if a in Tcell_metacells else 'black' for a in range(400)]
f = sns.clustermap(tsd,figsize = (.01,.01), col_colors = [cols,deg_cols],row_colors = deg_cols)
ri = f.dendrogram_row.reordered_ind[::-1]

fp = sns.clustermap(tsd[ri,:][:,ri],figsize = (20,20), col_colors = [cols[ri]],row_colors = deg_cols[ri], col_cluster = False, row_cluster = False,xticklabels=False, yticklabels=False,cmap = 'PuOr')

fp.ax_col_dendrogram.set_visible(False)
fp.ax_row_dendrogram.set_visible(False)

# %% [code]
assn = coup[12].argmax(axis = 1)

# %% [code]
#fig
fp = sns.clustermap(clusts[12][assn[ri],:][:,assn[ri]],figsize = (20,20), col_colors = [cols[ri],gcols[assn[ri]]],row_colors = [deg_cols[ri],gdeg_cols[assn[ri]]],xticklabels=False, yticklabels=False,col_cluster = False, row_cluster = False,cmap = 'PuOr',vmax = 1)


# %% [markdown]
### Plot discovered code

# %% [code]
plt.figure(figsize = (20,10),facecolor = None)
alphas = [0.5 if i in assn else 0.05 for i in np.arange(1000)]
xcoord = np.arange(400)*1000/400
ri = np.array(f.dendrogram_row.reordered_ind[::-1])
plt.scatter(xcoord ,[0]*400, c = cols[ri],edgecolor = 'black',s=100,zorder =-20,marker = '|',linewidth = 15)

bot = -0.2
gi = np.array(g.dendrogram_row.reordered_ind)
plt.scatter( range(1000),[bot]*1000, c = gcols[gi],s = 100,zorder = -20,marker = '|')
a = np.where(metaclust[ri]==1)[0]
b = np.where(np.isin(gi, assn[np.where(metaclust==1)[0]]))[0]
# plt.scatter(xcoord[a], [0]*len(a), s = 100,zorder = 10)
# plt.scatter(b, [-1]*len(b), s = 100,zorder = 10, c = gcols[gi[b]])
# plt.scatter(np.where(np.isin(gi,np.unique(assn)))[0], [-1]*len(np.unique(assn)), s = 5, color = 'black')
for i,a in zip(ri,assn[ri]):
    # if metaclust[i]!=1:
    #     continue
    # print(a)
    plt.plot([xcoord[np.where(ri==i)[0][-1]],np.where(gi==a)[0][-1]], [0,bot],linewidth = 0.3,color = 'black', linestyle = '--',zorder = -2)
    # break

plt.axis("off")
plt.ylim(-1.2,0.3)

# %% [markdown]
### Inspection of TCF7/T cell correspondence

# %% [markdown]
#### Investigating T cell subsets

# %% [code]
which_T = np.where(np.isin(metaclust, [1,2]))[0]


# %% [code]
central_T = which_T[np.argmin(tsd.mean(axis =1)[which_T])]

# %% [code]
from scipy.stats import spearmanr,pearsonr

# %% [code]
grads  = np.abs(gradx[which_T]).max(axis = 1)

# %% [code]
spearmanr(grads,tabsap['IL7R'].iloc[which_T])

# %% [code]
topgrad = which_T[metaclust[which_T]==1][(np.abs(grads[metaclust[which_T]==1])>2e-6)]

# %% [code]
plt.scatter(grads,tabsap['CCR7'].iloc[which_T], c = cols[which_T])

# %% [code]
plt.scatter(tsd[which_T][:,~np.isin(range(400), which_T)].mean(axis=1),tabsap['CCR7'].iloc[which_T], c = cols[which_T])

# %% [code]
spearmanr(tsd[which_T][:,~np.isin(range(400), which_T)].mean(axis=1),tabsap['BCL2'].iloc[which_T])

# %% [code]
r[r.isin(which_T)]

# %% [code]
r = pd.Series(ct_mc)
metaclust[r[r.isin(which_T)].filter(regex = 'cd4').values]

# %% [code]
plt.scatter(tsd[which_T][:,~np.isin(range(400), which_T)].mean(axis=1),tabsap['CD8A'].iloc[which_T], c = cols[which_T])

# %% [code]
plt.scatter(tsd[which_T][:,~np.isin(range(400), which_T)].mean(axis=1),tabsap['IL7R'].iloc[which_T], c = cols[which_T])

# %% [code]
plt.scatter(tsd[which_T][:,~np.isin(range(400), which_T)].mean(axis=1),tabsap['FAS'].iloc[which_T], c = cols[which_T])

# %% [markdown]
#### Plot similarity heatmaps restricted to T cells/TCF7

# %% [code]
gi = np.array(g.dendrogram_row.reordered_ind)

# aa = np.where(~np.isin(gmetaclust,[0]))[0]
aa = tcf7_clusts#np.where(np.isin(gi,tcf7_clusts))[0]
# gi2 = np.concatenate([np.array(gi)[~np.isin(gi, tcf7_clusts)],np.array(gi)[np.isin(gi, tcf7_clusts)]])
gi2 = gi
tc = np.array(gi2)[np.isin(gi2, aa)]

# tc = np.concatenate([tc[~np.isin(tc,tcf7_clusts)],tc[np.isin(tc, tcf7_clusts)]])
tcf_cols = np.array(['black','green'])[np.isin(np.arange(1000), tcf7_clusts).astype(int)]
t_assn_cols = np.array(['black','red'])[np.isin(np.arange(1000), assn[clust12]).astype(int)]
h = sns.clustermap(clusts[12][tc][:,gi2], row_colors = [gcols[tc], gdeg_cols[tc]],figsize = (20,3),row_cluster = True,col_cluster = False, col_colors = [tcf_cols[gi2],tcf_cols[gi2],tcf_cols[gi2], t_assn_cols[gi2],t_assn_cols[gi2],t_assn_cols[gi2],gdeg_cols[gi2],gdeg_cols[gi2],gdeg_cols[gi2],gdeg_cols[gi2],gdeg_cols[gi2]],cmap = 'PuOr',yticklabels=False,xticklabels=False,vmin = 0.2, vmax = 1)
h.ax_row_dendrogram.set_visible(False)
h.ax_col_dendrogram.set_visible(False)


# tc2 = tc[h.dendrogram_row.reordered_ind[::-1]]
# gi2 = gi2[h.dendrogram_col.reordered_ind[::-1]]
# h = sns.clustermap(clusts[12][tc2][:,gi2], row_colors = [gcols[tc], gdeg_cols[tc]],figsize = (20,3), col_colors = [tcf_cols[gi2],tcf_cols[gi2],tcf_cols[gi2], t_assn_cols[gi2],t_assn_cols[gi2],t_assn_cols[gi2],gdeg_cols[gi2],gdeg_cols[gi2],gdeg_cols[gi2]],cmap = 'PuOr',yticklabels=False,xticklabels=False,vmin = 0.2, vmax =0.9,col_cluster = False, row_cluster = False)
# h.ax_row_dendrogram.set_visible(False)
# h.ax_col_dendrogram.set_visible(False)

# %% [code]
clust12 = np.where(np.isin(metaclust,[1,2]))[0]
aa = clust12

# aa = Tcell_metacells#np.where(np.isin(metaclust,[1,2]))[0]
# ri2 = np.concatenate([np.array(ri)[~np.isin(ri, aa)],np.array(ri)[np.isin(ri, aa)]])
ri2 = np.argsort(np.argsort(g.dendrogram_row.reordered_ind)[assn])

# ri2 = np.argsort(v)
tc = np.array(ri2)[np.isin(ri2, aa)]
t_cols = np.array(['black','red'])[np.isin(np.arange(400), aa).astype(int)]
tcf_assn_cols = np.array(['black','green'])[np.isin(assn, tcf7_clusts).astype(int)]

h = sns.clustermap(tsd[tc][:,ri2], row_colors = [cols[tc], deg_cols[tc]],figsize = (10,1.5),col_cluster = False,row_cluster = True, col_colors = [ tcf_assn_cols[ri2], tcf_assn_cols[ri2], tcf_assn_cols[ri2],t_cols[ri2],t_cols[ri2],t_cols[ri2],deg_cols[ri2],deg_cols[ri2],deg_cols[ri2],deg_cols[ri2],deg_cols[ri2]],cmap = 'PuOr',yticklabels=False,xticklabels=False,vmin = 0.2, vmax = 1)
h.ax_row_dendrogram.set_visible(False)
h.ax_col_dendrogram.set_visible(False)

# %% [code]
from scipy.spatial.distance import pdist

# %% [markdown]
####  Direct quantification

# %% [code]

def select_hits(x_dist, x_clust,x_color, diff = 0.001, thresh = 0.4):
    v = x_dist.mean(axis=1)
    v-= v.mean()
    z=v/v.std()
    zstd = pd.Series(z).groupby(x_clust).mean()
    v = np.maximum(np.minimum(z,3),-3)
    v/=3
    v+=1
    v/=2
    
    v = pd.Series(v).groupby(x_clust).max()
    subdeg_cols = np.array(sns.color_palette('bwr_r',101))[(100*(v)).astype(int)]
    av = pd.DataFrame(x_dist).groupby(x_clust).mean().T.groupby(x_clust).mean()
    
    
    for i,j in zip(*np.where(av<thresh)):
        if i>=j:
            continue
        
        # print(i,j)
        zi = zstd.loc[i]
        zj = zstd.loc[j]
        if min(zi,zj)<=-0.1:
            continue
        if np.abs(zi-zj)<1:
            continue

        print(i,j,zi,zj, np.abs(zi-zj))
    
    # sns.clustermap(av, row_colors = subdeg_cols,col_colors = subdeg_cols,figsize = (5,5),cmap = 'PuOr')
    P1 = np.abs(zstd.values[:,None]- zstd.values[None,:])
    P2 = av.values
    i,j = np.where(av < thresh)
    
    ii = i[i<j]
    jj = j[i<j]

    i = ii[(zstd.values[ii]>-0.1) & (zstd.values[jj]>-0.1)]
    j = jj[(zstd.values[ii]>-0.1) & (zstd.values[jj]>-0.1)]
    
    plt.style.use("default")      #
    plt.scatter(P1[(i,j)], P2[i,j], c = x_color[i],s = 150)
    plt.scatter(P1[(i,j)], P2[i,j]-diff, c = x_color[j],s = 150)
    

    


# %% [code]
# two phenotypically similar blocks that differ in connectivity and have a shared central component

# %% [code]
plt.figure(figsize = (5,10))
plt.subplot(2,1,2)
select_hits(clusts[12],gmetaclust,gcol_pal,diff = 0.0035,thresh = 0.45)

plt.subplot(2,1,1)
select_hits(tsd,metaclust,col_pal,diff = 0.0069,thresh = 0.45)
plt.xlabel("connectivity difference (sd)")
plt.ylabel('phenotypic distance')
# plt.ylabel('phenotypic distance')

# %% [code]


# %% [code]
sns.palplot(np.array(sns.color_palette('hls',10))[[1,8]])

# %% [code]

lab = 12
labs = labels[lab]
perm_dat = {}
# np.unique(labels[12])

#temp has the assignments where there is overall enrichment
for clust in tqdm(temp[1].unique()):
    stats = {}
    whichc = np.where(labs==clust)[0]
    bg_size = len(whichc)

    bins = np.histogram(ens['gc'].loc[whichc],bins=10)
    

    for perm in tqdm(range(2500)):
        if perm==0:
            query_set = whichc
        else:
            query_set=  np.concatenate([np.random.choice(np.where((ens['gc']>bin_start)&(ens['gc']<=bin_end))[0],size =int(bg_size*bin_size/bins[0].sum())) for bin_start,bin_end,bin_size in zip(bins[1],bins[1][1:], bins[0])])
        for name,islet_df in beds.items():
            n = np.isin(query_set,islet_df['Name']).sum()
            stats.setdefault(name,[]).append(n)
            
    for name in beds:
        perm_dat[lab,clust,name] = [len(whichc), stats[name][0], (stats[name][0]> np.array(stats[name][1:])).mean(), np.mean(stats[name][1:]),np.max(stats[name][1:])]


# %% [code]
w = pd.DataFrame(perm_dat).T.loc[12].reset_index()
temp = w[(w[2]>0.95) & ((1.5)*w[3]<w[1])]
temp['ld'] = -np.log10(1/5000 + 1-w[2])
temp = temp[['level_0','level_1','ld']].drop_duplicates()
temp['name'] = temp['level_0'].map(names_dict)

plot_df = temp.set_index(['name','level_1'])['ld'].unstack().fillna(0).drop(['SOX9_PP','NKX2-5_Cardiomy.','PDX1_PP'],axis=1)
plot_df = plot_df.loc[plot_df.sum(axis=1)>0].loc[:,plot_df.sum(axis=0)>0]

cg = sns.clustermap(plot_df, cmap = 'Reds',vmax = 2,figsize = (0.1,0.1),metric = 'correlation')
plt.figure(figsize = (10,10))

sns.heatmap(plot_df.iloc[cg.dendrogram_row.reordered_ind,:].iloc[:,cg.dendrogram_col.reordered_ind], cmap = 'Reds',vmax = 3,cbar = False)
```
