## Supplementary material for "Deriving genetic codes for molecular phenotypes from first principles": Code: Fig3_mm10_OT_analysis.ipynb

```
# %% [code]
import pandas as pd
import numpy as np
import pyranges as pr
from scipy.spatial.distance import cdist
from scipy.sparse.csgraph import shortest_path
from sklearn.neighbors import kneighbors_graph
from sklearn.cluster import MiniBatchKMeans
import torch
torch.cuda.set_device(3)
# from ot.gromov import entropic_semirelaxed_gromov_wasserstein as gwd
from scipy.stats import fisher_exact, binomtest

# %% [code]
ens = pd.read_pickle('mm10_enh_pos')


# %% [code]
tabsap = pd.read_pickle('../data/tabsap_SEACELL_means')
tsd = shortest_path(kneighbors_graph(tabsap, metric = 'cosine',n_neighbors=20),directed=False)
tsd/=tsd.max()
tinfo = pd.read_pickle('../data/tabsap_SEACell_assignments')

# %% [code]
seq_features = np.load('mm10_cre_evs.npy')

# %% [code]

from tqdm.notebook import tqdm

# %% [code]
mbk_coup = {}
for i in tqdm(range(20)):
    mbk_coup[i] = [MiniBatchKMeans(n_clusters = 1000)]
    mbk_coup[i][0].fit(seq_features[:,:300])
    clusts = shortest_path(kneighbors_graph(mbk_coup[i][0].cluster_centers_,5,mode = 'connectivity'),directed=False)
    assert np.max(clusts)<100
    # clusts = cdist(mbk.cluster_centers_,mbk.cluster_centers_)
    clusts/=clusts.max()
    
    a1 = torch.from_numpy(tsd).float().cuda()
    a2 = torch.from_numpy(clusts).float().cuda()
    mbk_coup[i].append(gwd(a1,a2,epsilon=0.1))
    

# %% [code]
for i in mbk_coup:
    mbk,coup = mbk_coup[i]
    np.save('results/mm10_mbk{}_labels'.format(i),mbk.labels_)
    np.save('results/mm10_mbk{}_centers'.format(i),mbk.cluster_centers_)
    np.save('results/mm10_mbk{}_coupling'.format(i),coup.detach().cpu().numpy())
    
    

# %% [code]
mm10_labels = {}
mm10_centers = {}
mm10_coup = {}
mm10_clusts = {}
mm10_loss = {}

for i in range(20):
    mm10_labels[i] = np.load('results/mm10_mbk{}_labels.npy'.format(i))
    mm10_centers[i] = np.load('results/mm10_mbk{}_centers.npy'.format(i))
    mm10_coup[i] = np.load('results/mm10_mbk{}_coupling.npy'.format(i))
    mm10_clusts[i] = shortest_path(kneighbors_graph(mm10_centers[i],5,mode = 'connectivity'),directed=False)
    mm10_clusts[i]/=mm10_clusts[i].max()
    assn = mm10_coup[i].argmax(axis = 1)
    mm10_loss[i] = ((mm10_clusts[i][assn,:][:,assn]-tsd)**2).sum(axis =1)
    

# %% [code]
from scipy.cluster.hierarchy import fcluster
g = sns.clustermap(mm10_clusts[12], figsize = (0.01,0.01))
gmetaclust = fcluster(g.dendrogram_row.linkage, t = 10, criterion = 'maxclust')-1
gcol_pal = np.array(sns.color_palette('tab10',10))
gcols = gcol_pal[gmetaclust]
gcolor_map = {k:v for k,v in enumerate(gcols)}

v = mm10_clusts[12].mean(axis=1)
v-=v.mean()
v/=v.std()
v = np.maximum(np.minimum(v,3),-3)
v/=3

# v-=0.5
# v = np.maximum(np.minimum(v,0.25),-0.25)
# v/=0.25
v+=1
v/=2

gdeg_cols = np.array(sns.color_palette('bwr_r',101))[(100*(v)).astype(int)]
# gdeg_cols = np.array(sns.color_palette('bwr',101))[(100*((v-v.min())/(v.max()-v.min()))).astype(int)]
g = sns.clustermap(mm10_clusts[12],figsize = (10,10), col_colors = [gcols],row_colors = gdeg_cols,xticklabels=False, yticklabels=False,cmap= 'PuOr')
g.ax_col_dendrogram.set_visible(False)
g.ax_row_dendrogram.set_visible(False)
g.ax_cbar.set_visible(False)

# %% [code]
frequent_cells = tinfo[6].value_counts(normalize = True).iloc[:100].index

# %% [code]
w = tinfo.groupby(11).apply(lambda w : w[6].value_counts(normalize = True)).unstack().fillna(0)
frequent_cells = w.columns[(w.max(axis =0)>0.2)]
ct_hm = sns.clustermap(w[frequent_cells].iloc[ri].T,yticklabels = True,col_colors  = [cols[ri]],col_cluster = False,metric = 'correlation',xticklabels=False, figsize = (20,15))
ct_hm.ax_row_dendrogram.set_visible(False)


# %% [code]
f = sns.clustermap(tsd, figsize = (0.01,0.01))
metaclust = fcluster(f.dendrogram_row.linkage, t = 20, criterion = 'maxclust')-1
col_pal = np.array(sns.color_palette('tab20_r',20))
cols = col_pal[metaclust]
color_map = {k:v for k,v in zip(assn,cols)}
v = tsd.mean(axis=1)
v-=v.mean()
v/=v.std()
v = np.maximum(np.minimum(v,3),-3)
v/=3

# v-=0.5
# v = np.maximum(np.minimum(v,0.25),-0.25)
# v/=0.25
v+=1
v/=2
deg_cols = np.array(sns.color_palette('bwr_r',101))[(100*(v)).astype(int)]

f = sns.clustermap(tsd,figsize = (.01,.01), col_colors = [cols,deg_cols],row_colors = deg_cols)
ri = f.dendrogram_row.reordered_ind[::-1]

fp = sns.clustermap(tsd[ri,:][:,ri],figsize = (20,20), col_colors = [cols[ri]],row_colors = deg_cols[ri], col_cluster = False, row_cluster = False,xticklabels=False, yticklabels=False,cmap = 'PuOr')

fp.ax_col_dendrogram.set_visible(False)
fp.ax_row_dendrogram.set_visible(False)

# %% [code]
from matplotlib import pyplot as plt
plt.figure(figsize = (20,10),facecolor = None)
alphas = [0.5 if i in assn else 0.05 for i in np.arange(1000)]
xcoord = np.arange(400)*1000/400
ri = np.array(f.dendrogram_row.reordered_ind[::-1])
plt.scatter(xcoord ,[0]*400, c = cols[ri],edgecolor = 'black',s=100,zorder =-20,marker = '|',linewidth = 15)

bot = -0.2
gi = np.array(g.dendrogram_row.reordered_ind)
plt.scatter( range(1000),[bot]*1000, c = gcols[gi],s = 100,zorder = -20,marker = '|')
a = np.where(metaclust[ri]==1)[0]
b = np.where(np.isin(gi, assn[np.where(metaclust==1)[0]]))[0]
# plt.scatter(xcoord[a], [0]*len(a), s = 100,zorder = 10)
# plt.scatter(b, [-1]*len(b), s = 100,zorder = 10, c = gcols[gi[b]])
# plt.scatter(np.where(np.isin(gi,np.unique(assn)))[0], [-1]*len(np.unique(assn)), s = 5, color = 'black')
for i,a in zip(ri,assn[ri]):
    # if metaclust[i]!=1:
    #     continue
    # print(a)
    plt.plot([xcoord[np.where(ri==i)[0][-1]],np.where(gi==a)[0][-1]], [0,bot],linewidth = 0.3,color = 'black', linestyle = '--',zorder = -2)
    # break

plt.axis("off")
plt.ylim(-1.2,0.3)

# %% [code]


# %% [code]
cd4_ind = np.where(tabsap.index==tinfo[11][tinfo[6]=='cd4-positive, alpha-beta memory t cell'].value_counts().index[0])[0][0]

# %% [code]
labels = {}
centers = {}
coup = {}
clusts = {}
loss = {}

for i in range(20):
    labels[i] = np.load('results/mbk{}_labels.npy'.format(i))
    centers[i] = np.load('results/mbk{}_centers.npy'.format(i))
    coup[i] = np.load('results/mbk{}_coupling.npy'.format(i))
    clusts[i] = shortest_path(kneighbors_graph(centers[i],5,mode = 'connectivity'),directed=False)
    clusts[i]/=clusts[i].max()
    assn = coup[i].argmax(axis = 1)
    loss[i] = ((clusts[i][assn,:][:,assn]-tsd)**2).sum(axis =1)
    

# %% [code]
i = 12
assn = coup[i].argmax(axis = 1)
((clusts[i][assn,:][:,assn]-tsd)**2).mean(axis =1)[cd4_ind]

# %% [code]


# %% [code]

import seaborn as sns

# %% [code]
hg_df = pd.DataFrame({'count':pd.Series({i:(labels[i]==coup[i].argmax(axis=1)[cd4_ind]).mean() for i in loss}),
'loss':pd.Series({k: v[cd4_ind] for k,v in loss.items()})})
sns.scatterplot(data = hg_df,x = 'count',y = 'loss')

# %% [code]
mm10_df = pd.DataFrame({'count':pd.Series({i:(mm10_labels[i]==mm10_coup[i].argmax(axis=1)[cd4_ind]).mean() for i in mm10_loss}),
'loss':pd.Series({k: v[cd4_ind] for k,v in mm10_loss.items()})})
sns.scatterplot(data = mm10_df,x = 'count',y = 'loss')

# %% [code]

k = np.arange(24).reshape(4,6)

# %% [code]
np.random.shuffle(k[:,3])

# %% [code]
tabsap.mean(axis = 0).sort_values()

# %% [code]


hg_clust = torch.from_numpy(clusts[12]).float().cuda()
mm_clust = torch.from_numpy(mm10_clusts[1]).float().cuda()
perm_hg = []
perm_mm = []


with torch.no_grad():
    for perm in tqdm(range(200)):
        mat = tabsap.values.copy()
        if (len(perm_hg)>0):
            sel = np.random.choice(tabsap.shape[1],50)
            for j in sel:
                np.random.shuffle(mat[:,j])
            
        tsd = shortest_path(kneighbors_graph(mat, metric = 'cosine',n_neighbors=20),directed=False)
        tsd/=tsd.max()
        a1 = torch.from_numpy(tsd).float().cuda()
    
        hg_assn = gwd(a1,hg_clust,epsilon=0.1).argmax(axis=1)
        mm_assn = gwd(a1,mm_clust,epsilon=0.1).argmax(axis=1)
        
        
        perm_hg.append(((hg_clust[hg_assn[cd4_ind],:][hg_assn].cpu().numpy() - tsd[cd4_ind,:])**2).mean())
        perm_mm.append(((mm_clust[mm_assn[cd4_ind],:][mm_assn].cpu().numpy() - tsd[cd4_ind,:])**2).mean())


# %% [code]
(np.array(perm_mm)[1:]<perm_mm[0]).mean()

# %% [code]
(np.array(perm_hg)[1:]<perm_hg[0]).mean()

# %% [code]
hg_assn[cd4_ind]

# %% [code]
a = np.array(perm_hg)
b =np.array(perm_mm)

# %% [code]

a

# %% [code]
b

# %% [code]
a[0]*len(hg_assn)

# %% [code]
(b[1:]<b[0]).mean()

# %% [code]
!wget https://chip-atlas.dbcls.jp/data/mm10/eachData/bed05/SRX035992.05.bed

# %% [code]
import pyranges as pr

# %% [code]
ens = pd.read_pickle('mm10_enh_pos')
ens.iloc[:,:3].reset_index().iloc[:,[1,2,3,0]].to_csv('mm10_enh_pos.bed', sep = '\t',index = None,header=None)
enh_bed = pr.read_bed('mm10_enh_pos.bed')


# %% [code]
from scipy.stats import fisher_exact

# %% [code]
!wget https://chip-atlas.dbcls.jp/data/mm10/eachData/bed05/SRX2341960.05.bed

# %% [code]
!wget https://chip-atlas.dbcls.jp/data/mm10/eachData/bed05/SRX1144904.05.bed

# %% [code]
!wget https://chip-atlas.dbcls.jp/data/mm10/eachData/bed05/SRX9269217.05.bed

# %% [code]
!wget https://chip-atlas.dbcls.jp/data/mm10/eachData/bed05/SRX9833106.05.bed

# %% [code]
tcf7 = (pr.read_bed('SRX9833106.05.bed').join(enh_bed).as_df())

# %% [code]
cebpa_mono = (pr.read_bed('SRX9269217.05.bed').join(enh_bed).as_df())

# %% [code]
beds['tcf7'] = tcf7

# %% [code]
beds['cebpa_mono']  = cebpa_mono

# %% [code]
gata_bed = pr.read_bed('SRX035992.05.bed')
cebpa_bed = pr.read_bed('SRX2341960.05.bed')
gata3_bed = pr.read_bed('SRX1144904.05.bed')
gata3_ints = sorted(set(gata3_bed.join(enh_bed).Name_b))
gata2_ints = sorted(set(gata_bed.join(enh_bed).Name_b))
cebpa_ints = sorted(set(cebpa_bed.join(enh_bed).Name_b))

# %% [code]
mods = {}
for i in range(20):
    cd4_enh_clust = mm10_coup[i].argmax(axis=1)[cd4_ind]
    M = np.where(mm10_labels[i]==cd4_enh_clust)[0]
    m =np.isin(M,gata2_ints).sum()
    N = np.arange(len(mm10_labels[i]))
    n = len(gata2_ints)
    fe = fisher_exact([[m,len(M)-m],[n,len(N)-n]])
    stat,p = fe.statistic, fe.pvalue
    
    print(i,stat,p,mm10_df['loss'].loc[i])
    mods[i] = (-np.log10(p))*(stat>1)

# %% [code]
plt.figure(figsize = (2,5))
sns.scatterplot(data = pd.concat([mm10_df,pd.Series(mods)],axis=1), x = 'loss',y = 0,c = 'black')
plt.xlabel( 'alignment loss')
plt.ylabel( 'gata2 enrichment')

# %% [code]
cts = tinfo[6].unique()
dat ={}
i = 1
for ct in cts:
    card_endo = np.where(tabsap.index == tinfo[tinfo[6].str.contains(ct)][11].value_counts().index[0])[0][0]

    
    assn  = mm10_coup[i].argmax(axis = 1)
    labs = mm10_labels[i]
    
    
    card_assn= assn[card_endo]
    whichc = np.where(labs==card_assn)[0]

    
    M = np.where(mm10_labels[i]==card_assn)[0]
    N = np.arange(len(mm10_labels[i]))
    for name,ints_set in zip(['gata2','gata3','cebpa'], [gata2_ints,gata3_ints,cebpa_ints]):
        m =np.isin(M,ints_set).sum()
        n = len(ints_set)
        fe = fisher_exact([[m,len(M)-m],[n,len(N)-n]])
        stat,p = fe.statistic, fe.pvalue
        
        dat[ct,name] = [mm10_loss[i][card_endo],card_assn,len(whichc), fe.pvalue,fe.statistic]


# %% [code]
res = pd.DataFrame(dat).T

# %% [code]

from  matplotlib import pyplot as plt

# %% [code]
names_dict = {}
w = res.reset_index()
for clust in w[1].unique():
    names_dict[clust] = '; '.join(sorted(set(w['level_0'][w[1]==clust])))

# %% [code]
res[res[3]<0.05].filter(regex="uter",axis=0)

# %% [code]


# %% [code]

d = res.copy()
d['ld'] = np.log10(d[4])
temp= d[(d[3]<0.05) & (d[4]>1)].reset_index()[[1,'level_1','ld']].drop_duplicates()
temp['name'] = temp[1].map(names_dict)
plot_df = temp.set_index(['name','level_1'])['ld'].unstack().fillna(0)
plot_df = plot_df.loc[plot_df.sum(axis=1)>0]

cg = sns.clustermap(plot_df,cmap = 'Reds',vmax = 4,figsize = (0.1,0.1))
plt.figure(figsize = (1,5))
sns.heatmap(plot_df.iloc[cg.dendrogram_row.reordered_ind,:].iloc[:,cg.dendrogram_col.reordered_ind],cmap = 'Reds',vmax = 4,yticklabels=True,cbar = False)

# %% [code]
cts = tinfo[6].unique()
ct_mc = {ct:np.where(tabsap.index == tinfo[tinfo[6].str.contains(ct)][11].value_counts().index[0])[0][0] for ct in cts}
ct_mc_assn = {k:tinfo[6][tabsap.index[v]==tinfo[11]].value_counts().iloc[:5] for k,v in ct_mc.items()}


# %% [code]
ct_mc_assn['alveolar fibroblast']

# %% [code]
from tqdm.notebook import tqdm

# %% [code]
mm10_kmer = pd.read_pickle('mm10_enh_kmer_counts')

ens['gc'] = (mm10_kmer.loc[['G','C']].sum(axis=0)/mm10_kmer.loc[['G','C','A','T']].sum(axis=0)).values

# %% [code]

lab = 1
labs = mm10_labels[lab]
perm_dat = {}
# np.unique(labels[12])
for clust in tqdm(np.unique(mm10_coup[lab].argmax(axis=1))):
    stats = {}
    whichc = np.where(labs==clust)[0]
    bg_size = len(whichc)

    bins = np.histogram(ens['gc'].loc[whichc],bins=10)
    

    for perm in range(500):
        if perm==0:
            query_set = whichc
        else:
            query_set=  np.concatenate([np.random.choice(np.where((ens['gc']>bin_start)&(ens['gc']<=bin_end))[0],size =int(bg_size*bin_size/bins[0].sum())) for bin_start,bin_end,bin_size in zip(bins[1],bins[1][1:], bins[0])])
        for name,islet_df in beds.items():
            n = np.isin(query_set,islet_df['Name_b']).sum()
            stats.setdefault(name,[]).append(n)
            
    for name in beds:
        perm_dat[lab,clust,name] = [len(whichc), stats[name][0], (stats[name][0]> np.array(stats[name][1:])).mean(), np.mean(stats[name][1:]),np.max(stats[name][1:])]


# %% [code]
w = pd.DataFrame(perm_dat).T.loc[1].reset_index()
temp = w[(w[2]>0.95) & (w[3]<w[1])]

# %% [code]
w['name'] = w['level_0'].map(names_dict)

# %% [code]

w = w.dropna()

# %% [code]
nd = pd.Series({i: mm10_coup[1].argmax(axis=1)[v] for i,v in ct_mc.items()})

# %% [code]
names_dict = {}

for clust in nd.unique():
    names_dict[clust] = '; '.join(sorted(set(nd.index[nd==clust])))

# %% [code]
w.dropna()[w.dropna()['name'].str.contains('cd4')].sort_values(2)

# %% [code]

temp['ld'] = -np.log10(1/501 + 1-w[2])
temp['name'] = temp['level_0'].map(names_dict)
temp = temp[temp['level_1']!='cebpa']
# temp['name2'] = temp['level_0'].apply(lambda w : tinfo[6][tinfo[11].isin(tabsap.index[np.where(mm10_coup[1].argmax(axis=1)==w)[0] ])].value_counts().index[0])
# temp['name'][temp['name'].isna()] =temp['name2']

plot_df = temp.groupby(['level_1','name'])['ld'].max().unstack().fillna(0).T
cg = sns.clustermap(plot_df, cmap = 'Reds',vmax = 2,figsize = (0.1,0.1),metric = 'correlation')
plt.figure(figsize = (3,10))

sns.heatmap(plot_df.iloc[cg.dendrogram_row.reordered_ind,:].iloc[:,cg.dendrogram_col.reordered_ind], cmap = 'Reds',vmax = 3,cbar = False)

# %% [code]
ct_mc_assn['keratinocyte']

# %% [code]
beds = {'gata3':gata3_bed.join(enh_bed).as_df(),
 'cebpa':cebpa_bed.join(enh_bed).as_df(),
'gata2':gata_bed.join(enh_bed).as_df()}

# %% [code]
```
