## Supplementary material for "Deriving genetic codes for molecular phenotypes from first principles": Code: Fig4_eqtl_priors.ipynb

```
# %% [code]
# from scanpy import read_h5ad
import numpy as np
import pandas as pd
import pickle 
from tqdm.notebook import tqdm
import torch
from torch import nn


import os
from matplotlib import pyplot as plt
%pylab inline
import seaborn as sns

from numpy.lib.stride_tricks import sliding_window_view
from tqdm.notebook import tqdm


from scipy.stats import mannwhitneyu,ttest_ind
from scipy.stats import t as tdist
from statsmodels.stats.multitest import fdrcorrection
from sklearn.metrics import roc_curve,auc


from models.PSALearners import LatentPSALearner

torch.cuda.set_device(3)

# %% [code]
renamed = pd.read_pickle('../data/downstream/pbmc_renamed')

# %% [code]
renamed.index.values

# %% [code]
eqtls = pd.read_pickle('../data/downstream/eqtls')
eqtl_cols = eqtls.iloc[:,:20]
#map the cell types from sceqtl study to pbmc_atlas
cell_type_map = {'Classic Monocyte':  'CD14 Mono',
'Non-classic Monocyte': 'CD16 Mono',
'Naïve/Immature B Cell':'B naive',
'Memory B Cell':'B memory',
'Plasma Cell':'Plasmablast',
'CD8 Effector memory':'CD8 TEM',
'CD4 Effector memory/TEMRA':'CD4 TEM',
'Natural Killer Cell':'NK'}
sat_to_eq = {v:k for k,v in cell_type_map.items()}

eqtl_cols['SIG'] = eqtl_cols['FDR']<0.1
eqtl_cols['NOT_SIG'] = eqtl_cols['FDR']>0.5
gt = eqtl_cols[eqtl_cols['GENOTYPED'] == 'Genotyped']


# %% [code]
gct = eqtl_cols[['GENE','CELL_TYPE']].drop_duplicates()

# %% [code]
renamed.index

# %% [code]
tc = gct[gct['CELL_TYPE'].isin(renamed.index)].groupby("GENE").size().sort_values()
[a for a in tc.index[tc==8] if 'CD' in a]

# %% [code]
sig = good_qtl[(good_qtl['SIG']) & good_qtl['CELL_TYPE'].isin(renamed.index)]

# %% [code]
sel_genes = sig[['GENE','CELL_TYPE']].drop_duplicates().groupby('GENE').size().sort_values()
sel_genes = sel_genes.index[sel_genes>=3]

# %% [code]
good_qtl = eqtl_cols[eqtl_cols['GENE'].isin(tc.index[tc==8])]
good_qtl = good_qtl[good_qtl['GENE'].isin(sel_genes) & good_qtl['CELL_TYPE'].isin(renamed.index)]

# %% [code]
from scipy.spatial.distance import pdist,cdist

# %% [code]
gene_sig_counts = good_qtl[good_qtl['SIG']].groupby(['GENE','CELL_TYPE']).size().reset_index().iloc[:,:2].drop_duplicates().groupby('GENE').size()

# %% [code]
from scipy.stats import spearmanr
from tqdm import tqdm

# %% [code]


# %% [code]
res = {}

for gene in sel_genes:
    
    test_inds = good_qtl[good_qtl['GENE']==gene][['CELL_TYPE','RSID','SPEARMANS_RHO']].iloc[:,:2].drop_duplicates().index.values
    print(len(test_inds))
    gene_effects = good_qtl.loc[test_inds][['CELL_TYPE','RSID','SPEARMANS_RHO']].set_index(['CELL_TYPE','RSID'])['SPEARMANS_RHO'].unstack().fillna(0)
    com = sorted(set(gene_effects.index.values).intersection(renamed.index.values))
    
    i,j = np.triu_indices(len(com),1)    
    ed = cdist(gene_effects.loc[com],gene_effects.loc[com])
    gd = cdist(renamed.loc[com],renamed.loc[com])
    e = ed[(i,j)]
    
    
    shuf = np.arange(len(com))
    nperm = 1000
    stats = np.zeros(nperm+1)
    for perm in range(nperm+1):
        if perm>0:
            np.random.shuffle(shuf)
        g = gd[(shuf[i], shuf[j])]
        stats[perm] = spearmanr(e, g)[0]

        # if perm==0:
        #     plt.scatter(g,e)
        #     plt.title('{}, {}'.format(gene,stats[perm]))
        #     plt.show()
    res[gene] = {'r': stats[0],'perm_p':(stats[0]<=stats[1:]).mean(),'n_rs': len(test_inds)//len(com), 'nct':len(com)}


# %% [code]
dat = pd.concat([gene_sig_counts,pd.DataFrame(res).T.sort_values('perm_p')],axis=1).dropna().sort_values('perm_p')

# %% [code]
dat['sig'] = dat['perm_p']<=0.05

# %% [code]


# %% [code]
sns.stripplot(data = dat,
              x = 0, y = 'r',hue = 'sig',legend=False)
plt.hlines(dat.groupby(0)['r'].median(), (np.arange(6)-0.25) ,(np.arange(6)+0.25),colors = 'black')
plt.xlabel('number of cell types with significant eQTL')
plt.ylabel('correlation between dP and dG')
plt.title('Phenotype consistency holds approximately within cis windows for individual genes')

# %% [code]
plt.scatter(gd,ed)

# %% [code]
good_qtl.head()

# %% [code]
cd45 =0

# %% [code]
cd47= eqtl_cols[eqtl_cols['GENE']=='CD47']

# %% [code]
cd47[cd47['FDR']<0.2]

# %% [code]
genes = gt['GENE'].unique()

# %% [code]
cds = sorted([g for g in genes if 'CD' in g])

# %% [code]
sig = gt[gt['FDR']<0.1]

# %% [code]
gene_ct_counts = (sig.groupby(['GENE','CELL_TYPE']).size().unstack().fillna(0)[renamed.index.values]>0).sum(axis = 1)

# %% [code]
from scipy.spatial.distance import cdist
from scipy.stats import spearmanr

# %% [code]
variable_genes = gene_ct_counts[gene_ct_counts >4].index.values
selected_hits = sig[sig['GENE'].isin(variable_genes) & sig['CELL_TYPE'].isin(renamed.index.values)]
gene_groups = selected_hits.groupby('GENE').groups

gene_ct_rs = {}
for gene,inds in gene_groups.items():
    gene_ct_rs[gene] = {}
    for ct in renamed.index.values:
        gene_ct_rs[gene][ct] = set()
    for rs, ct in zip(selected_hits['RSID'].loc[inds], selected_hits['CELL_TYPE'].loc[inds]):
        gene_ct_rs[gene][ct].add(rs)
        
gene_ct_ious = {}
for gene,ct_rs in gene_ct_rs.items():
    gene_ct_ious[gene] = {}
    for ct1 in renamed.index.values:
        for ct2 in renamed.index.values:
            u = ct_rs[ct1].union(ct_rs[ct2])
            i = ct_rs[ct1].intersection(ct_rs[ct2])
            if len(u)>0:
                gene_ct_ious[gene][ct1,ct2] = 1-len(i)/len(u)
            
        

# %% [code]
cors = {}
ct_dist = cdist(renamed,renamed,metric = 'cosine')
triu_inds = np.triu_indices_from(ct_dist,1)
ctd = ct_dist[triu_inds]
for gene in gene_ct_ious.keys():
    
    iou_dist = pd.Series(gene_ct_ious[gene]).unstack().fillna(1).loc[renamed.index.values,:].loc[:,renamed.index.values].values
    np.random.seed(45)
    nperm = 1000
    stats = np.zeros(nperm+1)
    perm = np.arange(ct_dist.shape[0])
    for i in range(nperm+1):
        if i>0:
            np.random.shuffle(perm)
        ioud = iou_dist[perm,:][:,perm][triu_inds]
        stats[i] = spearmanr(ioud,ctd)[0]
    cors[gene] = (stats[0],(stats[0]>stats[1:]).mean())

# %% [code]

df = pd.DataFrame(cors).dropna(axis =1).T
df['lp'] = -np.log10(np.maximum(1-df[1],1/1001))
df['is_sig'] = df[1]>0.95
sns.scatterplot(data =df, x = 0, y = 'lp', hue = 'is_sig')

# %% [markdown]
### Try with effect sizes

# %% [code]
unique_gene_rs = gt[['GENE','RSID']].drop_duplicates()

# %% [code]
len(unique_gene_rs)

# %% [code]
rand_gene_rs = unique_gene_rs.sample(5000,random_state = 5)

# %% [code]
ct_gt = gt[gt['CELL_TYPE'].isin(renamed.index.values)]
good_gene_rs = {}
for gene,rs in tqdm(zip(*rand_gene_rs.values.T),total = len(rand_gene_rs)):
    gene_ctgt = ct_gt[(ct_gt['GENE']==gene)]
    gene_ctgt_rs = gene_ctgt[gene_ctgt['RSID'] == rs]
    if (len(gene_ctgt_rs))==8:
        good_gene_rs[(gene,rs)] = gene_ctgt_rs.copy()


# %% [code]
scores = []
for k, v in good_gene_rs.items():
#     scores.append(-np.log10(v.set_index('CELL_TYPE').loc[renamed.index.values]['P_VALUE']))
    scores.append(v.set_index('CELL_TYPE').loc[renamed.index.values]['SPEARMANS_RHO'])

# %% [code]
from scipy.sparse.csgraph import connected_components, shortest_path

# %% [code]

ct_dist = cdist(renamed,renamed,metric = 'cosine')
# rpcs = np.matmul(renamed, pcs.values)
# ct_dist = cdist(rpcs,rpcs,metric = 'euclidean')
# ct_dist = shortest_path(kneighbors_graph( renamed,n_neighbors=2,metric = 'cosine',mode= 'distance').toarray(),directed = False)

triu_inds = np.triu_indices_from(ct_dist,1)
ctd = ct_dist[triu_inds]
lpv_df = pd.concat(scores,axis = 1)
lpv_dist = cdist(lpv_df,lpv_df, metric = 'cityblock')

np.random.seed(45)
nperm = 10000
stats = np.zeros(nperm+1)
perm = np.arange(ct_dist.shape[0])
for i in range(nperm+1):
    if i>0:
        np.random.shuffle(perm)
    lpvd = lpv_dist[perm,:][:,perm][triu_inds]
    stats[i] = spearmanr(lpvd,ctd)[0]
    
print(stats[0],(stats[0]>stats[1:]).mean())

plt.scatter(ctd, lpv_dist[triu_inds])
plt.xlabel('cosine distance in phenotype space')
plt.ylabel('L1. distance in functionals')
plt.title('each point is a pair of cell types')

# %% [code]
for metric1 in ['euclidean','cosine']:
    for metric2 in ['euclidean','cosine','cityblock']:
        ct_dist = cdist(renamed,renamed,metric = metric1)
        # rpcs = np.matmul(renamed, pcs.values)
        # ct_dist = cdist(rpcs,rpcs,metric = 'euclidean')
        # ct_dist = shortest_path(kneighbors_graph( renamed,n_neighbors=2,metric = 'cosine',mode= 'distance').toarray(),directed = False)

        triu_inds = np.triu_indices_from(ct_dist,1)
        ctd = ct_dist[triu_inds]
        lpv_df = pd.concat(scores,axis = 1)
        lpv_dist = cdist(lpv_df,lpv_df, metric = metric2)

        np.random.seed(45)
        nperm = 10000
        stats = np.zeros(nperm+1)
        perm = np.arange(ct_dist.shape[0])
        for i in range(nperm+1):
            if i>0:
                np.random.shuffle(perm)
            lpvd = lpv_dist[perm,:][:,perm][triu_inds]
            stats[i] = spearmanr(lpvd,ctd)[0]

        print(stats[0],(stats[0]>stats[1:]).mean())

        plt.scatter(ctd, lpv_dist[triu_inds])
        plt.xlabel('{} distance in phenotype space'.format(metric1))
        plt.ylabel('{} distance in functionals'.format(metric2))
        plt.title('corr: {}, pval: {}'.format(stats[0], (stats[0]>stats[1:]).mean()))
        plt.show()

# %% [markdown]
# Syntax

# %% [code]
import seaborn as sns
from scipy.stats import spearmanr

# %% [code]
dbsnp = pd.read_table('chm13v2.0_GWASv1.0rsids_e100_r2022-03-08.vcf', header = None,sep = '\t', comment='#')

# %% [code]
snps = pd.read_pickle('../data/downstream/snp151')
non_patch = snps[~snps[1].str.contains('_')]
lifted_ctgt = ct_gt[ct_gt['RSID'].isin(dbsnp[2])]
t2t = pd.merge(lifted_ctgt, dbsnp[[0,1,2]].rename({2:'RSID'},axis = 1),on = 'RSID').rename({0:'t2t_chr',1:'t2t_pos'},axis = 1)
common = set(t2t['RSID']).intersection(non_patch[4])
np2 = non_patch[non_patch[4].isin(common)]
hg38 = np2[[1,2,4]].rename({1:'hg38_chr',2:'hg38_pos',4:'RSID'},axis = 1)

# %% [code]
dat = pd.merge(t2t, hg38, on = 'RSID')
gene_rsid_groups = dat.groupby(['GENE','RSID']).groups

# %% [code]
sel_gene_rs = []
for gene,rs in gene_rsid_groups:
    if len(gene_rsid_groups[gene,rs])==8:
        sel_gene_rs.append((gene,rs))
        

# %% [code]
gene_rs_features = {}
for k in tqdm(sel_gene_rs):
    cur_df = dat.loc[gene_rsid_groups[k]].set_index('CELL_TYPE').loc[renamed.index.values]
    t2t_chrom = cur_df['t2t_chr'].iloc[0]
    t2t_pos = cur_df['t2t_pos'].iloc[0]//1000
    t2t_feats = np.load('../data/seq/GENA_2k_precomputed/{}_precomputed_features.npy'.format(t2t_chrom), mmap_mode = 'r')[t2t_pos].copy()
    
    hg38_chrom = cur_df['hg38_chr'].iloc[0]
    hg38_pos = cur_df['hg38_pos'].iloc[0]
    chr_dat = np.load('../data/seq/precomputed/{}_precomputed_features_512.npy'.format(hg38_chrom), mmap_mode = 'r')
    chr_pos = np.load('../data/seq/precomputed/{}_CHROM_positions_512.npy'.format(hg38_chrom))
    hg38_feats = chr_dat[np.where(hg38_pos>chr_pos)[0].argmin()].copy()
    

    gene_rs_features[k] = (t2t_feats,hg38_feats,
                           cur_df['P_VALUE'].values.copy(),
                          cur_df['SPEARMANS_RHO'].values.copy())
                           
                           

# %% [code]
rsid_positions = dat[['RSID','t2t_chr','t2t_pos']].drop_duplicates().set_index('RSID')

# %% [code]
from sklearn.neighbors import kneighbors_graph

# %% [code]
gene2rs = {}
for gene,rs in sel_gene_rs:
    if (gene_rs_features[gene,rs][2]<0.01).sum()>2:
        gene2rs.setdefault(gene, set()).add(rs)
        
#filter gene/rs pairs that have multiple 

# %% [code]

# chosen_features = [(k,np.random.choice(list(v))) for k,v in gene2rs.items()]
chosen_features = []
for k,v in gene2rs.items():
    for j in v:
        chosen_features.append((k,j))


N_chosen = len(chosen_features)

cho_np2 = np2[np2[4].isin([a[1] for a in chosen_features])]

dvs = []
for k,v in cho_np2.groupby(1).groups.items():
    if len(v) <2:
        continue
    pos = cho_np2[2].loc[v].values

    dvs.append(np.abs(pos[:,None]-pos[None,:])[np.triu_indices(len(v),1)].min())
dist_val = np.min(dvs)


gena_syn_features = np.stack([gene_rs_features[k][0] for k in chosen_features])
gena_syn_d = cdist(gena_syn_features,gena_syn_features,metric = 'cosine')
gena_syn_g = kneighbors_graph(gena_syn_d,n_neighbors = 5, metric = 'precomputed').toarray()

syn_features = np.stack([gene_rs_features[k][1] for k in chosen_features])
syn_d = cdist(syn_features,syn_features,metric = 'cosine')
syn_g = kneighbors_graph(syn_d,n_neighbors = 5, metric = 'precomputed').toarray()

assoc = ((-np.log10(np.stack([gene_rs_features[k][2] for k in chosen_features])))>1).astype(float)
assoc_d = cdist(assoc,assoc,metric = 'jaccard')


res = {}


inds = np.triu_indices(5,1)
for n_neighbors in range(5,40):
    
    perm_syn_g = kneighbors_graph(gena_syn_d,n_neighbors = n_neighbors, metric = 'precomputed').toarray()
    w = np.zeros(N_chosen)
    for j in range(N_chosen):
        q = np.where(perm_syn_g[j])[0]
        sub = np.zeros(1000)
        for k in range(len(sub)):
            subq = np.random.choice(q, 5, replace = False)
            sub[k] = assoc_d[subq,:][:,subq][inds].mean()
        w[j] = np.median(sub)
    res[n_neighbors]=w.copy()

figsize(5,5)
sns.lineplot(data = pd.concat([pd.DataFrame(res[k],columns = [k]).melt() for k in res], axis =0, ignore_index = True),
            x = 'variable', y = 'value',errorbar = 'se')

plt.show()

res = {}
for n_neighbors in range(5,40):
    inds = np.triu_indices(n_neighbors,1)    
    perm_syn_g = kneighbors_graph(gena_syn_d,n_neighbors = n_neighbors, metric = 'precomputed').toarray()
    w = np.zeros(N_chosen)
    for j in range(N_chosen):
        q = np.where(perm_syn_g[j])[0]
        w[j] = np.median(assoc_d[q,:][:,q][inds].mean())
    res[n_neighbors]=w.copy()
        
figsize(5,5)
sns.lineplot(data = pd.concat([pd.DataFrame(res[k],columns = [k]).melt() for k in res], axis =0, ignore_index = True),
            x = 'variable', y = 'value',errorbar = 'se')

plt.show()

# %% [code]

# chosen_features = [(k,np.random.choice(list(v))) for k,v in gene2rs.items()]
chosen_features = []
for k,v in gene2rs.items():
    for j in v:
        chosen_features.append((k,j))


N_chosen = len(chosen_features)

cho_np2 = np2[np2[4].isin([a[1] for a in chosen_features])]

dvs = []
for k,v in cho_np2.groupby(1).groups.items():
    if len(v) <2:
        continue
    pos = cho_np2[2].loc[v].values

    dvs.append(np.abs(pos[:,None]-pos[None,:])[np.triu_indices(len(v),1)].min())
dist_val = np.min(dvs)


gena_syn_features = np.stack([gene_rs_features[k][0] for k in chosen_features])
gena_syn_d = cdist(gena_syn_features,gena_syn_features,metric = 'cosine')
gena_syn_g = kneighbors_graph(gena_syn_d,n_neighbors = 5, metric = 'precomputed').toarray()

syn_features = np.stack([gene_rs_features[k][1] for k in chosen_features])
syn_d = cdist(syn_features,syn_features,metric = 'cosine')
syn_g = kneighbors_graph(syn_d,n_neighbors = 5, metric = 'precomputed').toarray()

assoc = ((-np.log10(np.stack([gene_rs_features[k][2] for k in chosen_features])))>1).astype(float)
assoc_d = cdist(assoc,assoc,metric = 'jaccard')


orig = gena_syn_features
new = orig.copy()

n_neighbors = 5

perm = np.arange(syn_g.shape[0])
nperm = 10000
stats = np.zeros((nperm+1,len(perm)))
inds = np.triu_indices(n_neighbors,1)
for i in tqdm(range(nperm+1)):
#         if i>0:
#             new = np.random.randn(*orig.shape)

#         perm_syn_d = cdist(new,new,metric = 'cosine')
#         perm_syn_g = kneighbors_graph(perm_syn_d,n_neighbors = n_neighbors, metric = 'precomputed').toarray()

    if i==0:
        perm_syn_g = kneighbors_graph(gena_syn_d,n_neighbors = n_neighbors, metric = 'precomputed').toarray()
    if i>0:
        np.random.shuffle(perm)


    perm_syn_g = perm_syn_g[:,perm]


#     stats[i] = (syn_g*assoc_g).sum(axis =1).mean()
    w = np.zeros(N_chosen)
    for j in range(N_chosen):
 
        q = np.where(perm_syn_g[j])[0]
        
        w[j] = assoc_d[q,:][:,q][inds].mean()

#         w[j] = np.square(assoc[j][None]-assoc[q]).mean(axis = 1).mean()
#         w[j] = np.square(assoc[j]-assoc[q].mean(axis =0)).mean()
    stats[i] = w.copy()
    
figsize(2,5)
sns.stripplot(stats[1:].mean(axis = 1), c = 'grey', s = 1)
plt.hlines(stats[0].mean(),-0.5,0.5 , colors = 'red')
plt.title('gena features, pval = {}'.format((stats[0].mean()<stats[1:].mean(axis = 1)).mean()))

# %% [code]
chosen_features = []
for k,v in gene2rs.items():
    for j in v:
        chosen_features.append((k,j))
sub_p = rsid_positions.loc[[v[1] for v in chosen_features]].sort_values(['t2t_chr','t2t_pos'])
is_close = []
thresh = 5000
for j in range(1,len(sub_p)-1):
     is_close.append(((sub_p['t2t_chr'].iloc[j] == sub_p['t2t_chr'].iloc[j-1]) & (np.abs(sub_p['t2t_pos'].iloc[j] - sub_p['t2t_pos'].iloc[j-1]) < thresh))|((sub_p['t2t_chr'].iloc[j] == sub_p['t2t_chr'].iloc[j+1]) & (np.abs(sub_p['t2t_pos'].iloc[j] - sub_p['t2t_pos'].iloc[j+1]) < thresh)))
        
        
        
sub_p['is_close'] = [is_close[0].copy()]+is_close + [is_close[-1].copy()]


chosen_features = [(k,v) for k,v in chosen_features if sub_p['is_close'].loc[v]==False]

N_chosen = len(chosen_features)

cho_np2 = np2[np2[4].isin([a[1] for a in chosen_features])]

dvs = []
for k,v in cho_np2.groupby(1).groups.items():
    if len(v) <2:
        continue
    pos = cho_np2[2].loc[v].values

    dvs.append(np.abs(pos[:,None]-pos[None,:])[np.triu_indices(len(v),1)].min())
dist_val = np.min(dvs)


gena_syn_features = np.stack([gene_rs_features[k][0] for k in chosen_features])
gena_syn_d = cdist(gena_syn_features,gena_syn_features,metric = 'cosine')
gena_syn_g = kneighbors_graph(gena_syn_d,n_neighbors = 5, metric = 'precomputed').toarray()

syn_features = np.stack([gene_rs_features[k][1] for k in chosen_features])
syn_d = cdist(syn_features,syn_features,metric = 'cosine')
syn_g = kneighbors_graph(syn_d,n_neighbors = 5, metric = 'precomputed').toarray()

assoc = ((-np.log10(np.stack([gene_rs_features[k][2] for k in chosen_features])))>1).astype(float)
assoc_d = cdist(assoc,assoc,metric = 'jaccard')


new = orig.copy()

n_neighbors = 5

perm = np.arange(syn_g.shape[0])
nperm = 10000
stats = np.zeros((nperm+1,len(perm)))
inds = np.triu_indices(n_neighbors,1)
for i in tqdm(range(nperm+1)):
#         if i>0:
#             new = np.random.randn(*orig.shape)

#         perm_syn_d = cdist(new,new,metric = 'cosine')
#         perm_syn_g = kneighbors_graph(perm_syn_d,n_neighbors = n_neighbors, metric = 'precomputed').toarray()

    if i==0:
        perm_syn_g = kneighbors_graph(syn_d,n_neighbors = n_neighbors, metric = 'precomputed').toarray()
    if i>0:
        np.random.shuffle(perm)


    perm_syn_g = perm_syn_g[:,perm]


#     stats[i] = (syn_g*assoc_g).sum(axis =1).mean()
    w = np.zeros(N_chosen)
    for j in range(N_chosen):
 
        q = np.where(perm_syn_g[j])[0]
        
        w[j] = assoc_d[q,:][:,q][inds].mean()

#         w[j] = np.square(assoc[j][None]-assoc[q]).mean(axis = 1).mean()
#         w[j] = np.square(assoc[j]-assoc[q].mean(axis =0)).mean()
    stats[i] = w.copy()
    
figsize(2,5)
sns.stripplot(stats[1:].mean(axis = 1), c = 'grey', s = 1)
plt.hlines(stats[0].mean(),-0.5,0.5 , colors = 'red')
plt.title((stats[0].mean()<stats[1:].mean(axis = 1)).mean())

# %% [code]


# %% [code]


# %% [code]


# %% [code]


# %% [code]


# %% [markdown]
### DNABert features

# %% [code]

# chosen_features = [(k,np.random.choice(list(v))) for k,v in gene2rs.items()]
chosen_features = []
for k,v in gene2rs.items():
    for j in v:
        chosen_features.append((k,j))


N_chosen = len(chosen_features)

cho_np2 = np2[np2[4].isin([a[1] for a in chosen_features])]

dvs = []
for k,v in cho_np2.groupby(1).groups.items():
    if len(v) <2:
        continue
    pos = cho_np2[2].loc[v].values

    dvs.append(np.abs(pos[:,None]-pos[None,:])[np.triu_indices(len(v),1)].min())
dist_val = np.min(dvs)


gena_syn_features = np.stack([gene_rs_features[k][0] for k in chosen_features])
gena_syn_d = cdist(gena_syn_features,gena_syn_features,metric = 'cosine')
gena_syn_g = kneighbors_graph(gena_syn_d,n_neighbors = 5, metric = 'precomputed').toarray()

syn_features = np.stack([gene_rs_features[k][1] for k in chosen_features])
syn_d = cdist(syn_features,syn_features,metric = 'cosine')
syn_g = kneighbors_graph(syn_d,n_neighbors = 5, metric = 'precomputed').toarray()

assoc = ((-np.log10(np.stack([gene_rs_features[k][2] for k in chosen_features])))>1).astype(float)
assoc_d = cdist(assoc,assoc,metric = 'jaccard')


orig = gena_syn_features
new = orig.copy()

n_neighbors = 5

perm = np.arange(syn_g.shape[0])
nperm = 10000
stats = np.zeros((nperm+1,len(perm)))
inds = np.triu_indices(n_neighbors,1)
for i in tqdm(range(nperm+1)):
#         if i>0:
#             new = np.random.randn(*orig.shape)

#         perm_syn_d = cdist(new,new,metric = 'cosine')
#         perm_syn_g = kneighbors_graph(perm_syn_d,n_neighbors = n_neighbors, metric = 'precomputed').toarray()

    if i==0:
        perm_syn_g = kneighbors_graph(syn_d,n_neighbors = n_neighbors, metric = 'precomputed').toarray()
    if i>0:
        np.random.shuffle(perm)


    perm_syn_g = perm_syn_g[:,perm]


#     stats[i] = (syn_g*assoc_g).sum(axis =1).mean()
    w = np.zeros(N_chosen)
    for j in range(N_chosen):
 
        q = np.where(perm_syn_g[j])[0]
        
        w[j] = assoc_d[q,:][:,q][inds].mean()

#         w[j] = np.square(assoc[j][None]-assoc[q]).mean(axis = 1).mean()
#         w[j] = np.square(assoc[j]-assoc[q].mean(axis =0)).mean()
    stats[i] = w.copy()
    
figsize(2,5)
sns.stripplot(stats[1:].mean(axis = 1), c = 'grey', s = 1)
plt.hlines(stats[0].mean(),-0.5,0.5 , colors = 'red')
plt.title('distilbert features, pval = {}'.format((stats[0].mean()<stats[1:].mean(axis = 1)).mean()))
```
