## Supplementary material for "Deriving genetic codes for molecular phenotypes from first principles": Code: Fig4_finemapped_tabsap.ipynb

```
# %% [code]
import numpy as np
import pandas as pd
from scanpy import read_h5ad
import torch
from matplotlib import pyplot as plt
from tqdm.notebook import tqdm
from sklearn.neighbors import NearestNeighbors
from scipy.spatial.distance import cdist
import seaborn as sns
from scipy.cluster.hierarchy import fcluster
from sklearn.neighbors import kneighbors_graph

import torch
from torch import nn
from models.PSALearners import LatentPSALearner
from sklearn.metrics import roc_auc_score
from scipy.stats import spearmanr,ttest_1samp,ttest_ind
DEVICE = 7

# %% [code]
!wget https://storage.googleapis.com/gtex_analysis_v8/single_tissue_qtl_data/GTEx_v8_finemapping_CAVIAR.tar

# %% [code]
!tar -xvf GTEx_v8_finemapping_CAVIAR.tar

# %% [code]
range(40,100,2)

# %% [code]
anns = pd.read_pickle('../data/sc/processed/tabsap_SEACell_assignments')
sc = pd.read_pickle('../data/sc/processed/tabsap_SEACELL_means')/1000
tissue_counts = anns.groupby([11,0]).size().unstack().fillna(0)
tissue_dists = pd.DataFrame(tissue_counts.values/tissue_counts.values.sum(axis = 0,keepdims=True), index = tissue_counts.index, columns = tissue_counts.columns)
tissue_genex_means = pd.DataFrame(np.matmul(tissue_dists.values.T, sc.values), index = tissue_dists.columns, columns = sc.columns)


# %% [code]
from liftover import get_lifter
converter = get_lifter('hg38','GCA_009914755.4')

# %% [code]
eqtls =pd.read_table('GTEx_v8_finemapping_CaVEMaN/GTEx_v8_finemapping_CaVEMaN.txt')

# %% [code]
t2ts = [converter[chrom[3:]][pos] for chrom, pos in zip(eqtls['CHROM'],eqtls['POS'])]

# %% [code]
CP_chroms = ['CP068277.2',
'CP068276.2',
'CP068275.2',
'CP068274.2',
'CP068273.2',
'CP068272.2',
'CP068271.2',
'CP068270.2',
'CP068269.2',
'CP068268.2',
'CP068267.2',
'CP068266.2',
'CP068265.2',
'CP068264.2',
'CP068263.2',
'CP068262.2',
'CP068261.2',
'CP068260.2',
'CP068259.2',
'CP068258.2',
'CP068257.2',
'CP068256.2',
'CP068255.2',
'CP086569.2']
chroms = ['chr{}'.format(j) for j in list(range(1,23))+['X','Y']]
chrom_map = {ch:cp for cp,ch in zip(chroms,CP_chroms)}

# %% [code]
gene_info = pd.read_pickle('../data/sc/processed/tabsap_gene_info').loc[sc.columns]
gene_info['ensembl_split'] = [a.split('.')[0] for a in gene_info['ensemblid']]
good_genes = gene_info['ensembl_split'][~gene_info['ensemblid'].str.contains('PAR')]
gene_info = gene_info[gene_info['ensembl_split'].isin(good_genes)]

# %% [code]
converted_chr = {}
converted_pos = {}

for i,a in enumerate(t2ts):
    if len(a)==0:
        continue
    if a[0][0] in chrom_map:
        converted_chr[i] = chrom_map[a[0][0]]
        converted_pos[i] = a[0][1]
converted_chr = pd.Series(converted_chr)
converted_pos = pd.Series(converted_pos)


t2t_added = pd.concat([eqtls, pd.DataFrame({'t2t_chr':converted_chr}), pd.DataFrame({'t2t_pos':converted_pos})],axis = 1).dropna()
common_genes = list(set(gene_info['ensemblid']).intersection(t2t_added['GENE']))
t2t_added = t2t_added[t2t_added['GENE'].isin(common_genes)]
t2t_added['gene_symbol'] = gene_info.set_index('ensemblid')['gene_symbol'].loc[t2t_added['GENE']].values
t2t_added['t2t_ind'] = t2t_added['t2t_pos'].astype(int)//1000


# %% [code]
len(set(eqtls[eqtls['GENE'].isin(common_genes)]['eQTL']))

# %% [code]
len(set(t2t_added['eQTL']))

# %% [code]
1-144940/145778 #0.5 per cent were lost in liftover

# %% [code]
max_pips = {}
for tissue, gene, t2t_chr,t2t_ind,pip in zip(t2t_added['TISSUE'],t2t_added['gene_symbol'],t2t_added['t2t_chr'],t2t_added['t2t_ind'], t2t_added['Probability']):
    max_pips.setdefault((tissue, gene, t2t_chr,t2t_ind), []).append(pip) 
    
max_pip = pd.Series({k: np.max(v) for k,v in max_pips.items()}).reset_index()
max_pip.columns = ['TISSUE','GENE','t2t_chr', 't2t_ind', 'PIP']
max_pip = max_pip[max_pip['t2t_chr']!='chrY']

common_tissues = set(tissue_genex_means.index).intersection(max_pip['TISSUE'].unique())

renamed = tissue_genex_means.loc[sorted(common_tissues)]

# %% [code]
valid_long = max_pip[max_pip['TISSUE'].isin(common_tissues)]
thresh = 0.9
good_tg_pairs = (valid_long.groupby(['TISSUE','GENE'])['PIP'].min() <thresh)&(valid_long.groupby(['TISSUE','GENE'])['PIP'].max() >thresh)

gtg = (valid_long.groupby(['TISSUE','GENE']).size().sort_values()>3)
good_tissue_gene_pairs = set(good_tg_pairs[good_tg_pairs].index.values).intersection(gtg[gtg].index.values)

valid_long = valid_long.loc[[i for i,a,b in zip(valid_long.index.values, valid_long['TISSUE'], valid_long['GENE']) if (a,b) in good_tissue_gene_pairs]]
valid_long = valid_long[valid_long['GENE'].isin(sc.columns)]


def prepare_sc_batch(ct, gene_to_add, amount = 0.0001):
    ct_mean = renamed.loc[ct]
    edited_mean = ct_mean.copy()
    edited_mean[gene_to_add]+=amount
    return torch.tensor(np.stack([ct_mean.values.copy(), edited_mean.values]),device = 'cuda:{}'.format(DEVICE),dtype = torch.float)

# %% [code]
unique_pos = valid_long[['t2t_chr','t2t_ind']].drop_duplicates()
chrom_inds = unique_pos.groupby('t2t_chr').groups
cis_seq_batches = []
dat_inds =[]
for chrom,inds in chrom_inds.items():
    chm13_pos = unique_pos['t2t_ind'].loc[inds]
    chr_dat = np.load('../data/seq/GENA_2k_precomputed/{}_precomputed_features.npy'.format(chrom), mmap_mode = 'r')
    cis_seq_batches.append(chr_dat[chm13_pos.values]/50)
    dat_inds+= list(inds)

cis_seq_batch = torch.tensor(np.concatenate(cis_seq_batches)).float().to('cuda:{}'.format(DEVICE))

qtl_to_seq_map = {ch_ind:i for i, ch_ind in enumerate (zip(unique_pos['t2t_chr'].loc[dat_inds],unique_pos['t2t_ind'].loc[dat_inds]))}


ct_gene_groups = valid_long.groupby(['TISSUE','GENE']).groups
cis_seq_inds = {}
pips = {}
for key,inds in ct_gene_groups.items():
    CELLTYPE,GENE = key
    cis_seq_inds[CELLTYPE, GENE] = np.array([qtl_to_seq_map[a] for a in zip(valid_long['t2t_chr'].loc[inds],valid_long['t2t_ind'].loc[inds])])
    pips[CELLTYPE,GENE] = valid_long['PIP'].loc[inds]

# %% [code]
def permutation_test(path,perm_gene = False,perm_ct= False,perm_rs = False,group_perm_gene = False,rand_init = False, nperm =1000):
    n_learn = 1
    n_control = 40
    dropout= 0
    softmax = True
    bn = False
    multi_learner = nn.ModuleList([LatentPSALearner(sc = torch.from_numpy(renamed.values),dropout=dropout, softmax=softmax, genome = cis_seq_batch, n_control = n_control, bias = False,batch_norm = bn) for _ in range(n_learn)]).to('cuda:{}'.format(DEVICE))

    checkpoint = torch.load(path)
    sd = checkpoint['learner_state_dict']
    multi_learner.load_state_dict(sd)
    multi_learner.eval();
    learner = multi_learner[0]
    learner.eval();
    

    # initialize maps that will be shuffled during permutations
    orig_genes = valid_long['GENE'].unique()
    shuf_genes = orig_genes.copy()
    orig_cts = renamed.index.values.copy()
    shuf_cts = orig_cts.copy()
    shuf_idx = np.arange(len(cis_seq_batch))

    # want to enable shuffling only genes that are expressed in the same set of cell types, hence this shuffle
    ct_set_to_g = {}
    for g, ct_set in {g: tuple(sorted(renamed[g][renamed[g]>5e-5].index.values)) for g in orig_genes}.items():
        ct_set_to_g.setdefault(ct_set, []).append(g)

    orig_group_genes = [np.array(gs) for _,gs in ct_set_to_g.items()]
    shuf_group_genes = [arr.copy() for arr in orig_group_genes]
    print([len(a) for a in orig_group_genes])


    assert (group_perm_gene&perm_gene)==False

    perm_stats = {}
    PRINT_EVERY = min(nperm,20)
    np.random.seed(45)

    for perm in tqdm(range(1+nperm)):
        # first permutation is always test statistic (i.e., no shuffle)
        if perm>0:
            if group_perm_gene:
                for arr in shuf_group_genes:
                    np.random.shuffle(arr) #only shuffle within groups
            if perm_gene:
                np.random.shuffle(shuf_genes)

            if perm_ct:
                np.random.shuffle(orig_cts)

            if perm_rs:
                np.random.shuffle(shuf_idx)

            if rand_init:
                multi_learner = nn.ModuleList([LatentPSALearner(sc = torch.from_numpy(renamed.values),dropout=dropout, softmax=softmax, genome = cis_seq_batch, n_control = n_control, bias = False,batch_norm = bn) for _ in range(n_learn)]).to('cuda:{}'.format(DEVICE))
                multi_learner.eval();
                learner = multi_learner[0]
                learner.eval();


        # if perm==0, shuf_gene_map, shuf_idx, shuf_ct_map are all identity
        if group_perm_gene:
            shuf_gene_map = {i:j for i,j in zip(np.concatenate(orig_group_genes), np.concatenate(shuf_group_genes))}
        else:
            shuf_gene_map = {i:j for i,j in zip(orig_genes,shuf_genes)}

        shuf_ct_map = {i:j for i,j in zip(orig_cts,shuf_cts)}


        # compute AUCs for the discrepancy
        for CELLTYPE, GENE in ct_gene_groups.keys():
            sub_seq = cis_seq_batch[shuf_idx][cis_seq_inds[CELLTYPE,GENE]]


            shuf_ct = shuf_ct_map[CELLTYPE]
            shuf_gene = shuf_gene_map[GENE]
            sc_batch = prepare_sc_batch(shuf_ct, shuf_gene,amount=1e-4)

            Txs =  learner.T_xs(seq_batch= sub_seq, sc_batch = sc_batch)
            y_pred = torch.abs(Txs[1]-Txs[0]).detach().cpu().numpy()
            y_true = pips[CELLTYPE,GENE]
            
            y_true = (pips[CELLTYPE,GENE]>0.9).astype(int).values
            perm_stats[perm,CELLTYPE,GENE] = roc_auc_score(y_true= y_true, y_score = y_pred)

#             perm_stats[perm,CELLTYPE,GENE] = spearmanr(y_true,y_pred)[0]
#             perm_stats[perm,CELLTYPE,GENE] = (y_pred[y_true>0.9].mean() - y_pred[y_true<0.9]).mean()/(1e-5+y_pred.std())


    res = pd.Series(perm_stats).reset_index()
    res.columns = ['perm', 'ct', 'gene', 'auc']
    res['perm_gene'] = perm_gene
    res['perm_ct'] = perm_ct
    res['group_perm_gene'] = group_perm_gene
    res['rand_init'] = rand_init
    res['perm_rs'] = perm_rs
    return res


# %% [code]
NPERM = 501


results = []


path  = 'results/apr10/tabsap_meansyntax_repeat/repeat_0_epoch_52.pt'
perm_gene = True
perm_ct= False
perm_rs = False
group_perm_gene = False
rand_init = False
results.append(permutation_test(path,perm_gene = perm_gene,perm_ct= perm_ct,perm_rs = perm_rs,group_perm_gene = group_perm_gene,rand_init = rand_init,
                 nperm =NPERM))

perm_gene = False
perm_ct= False
perm_rs = False
group_perm_gene = True
rand_init = False
results.append(permutation_test(path,perm_gene = perm_gene,perm_ct= perm_ct,perm_rs = perm_rs,group_perm_gene = group_perm_gene,rand_init = rand_init,
                 nperm =NPERM))


perm_gene = False
perm_ct= False
perm_rs = True
group_perm_gene = False
rand_init = False
results.append(permutation_test(path,perm_gene = perm_gene,perm_ct= perm_ct,perm_rs = perm_rs,group_perm_gene = group_perm_gene,rand_init = rand_init,
                 nperm =NPERM))


perm_gene = False
perm_ct= True
perm_rs = False
group_perm_gene = False
rand_init = False
results.append(permutation_test(path,perm_gene = perm_gene,perm_ct= perm_ct,perm_rs = perm_rs,group_perm_gene = group_perm_gene,rand_init = rand_init,
                 nperm =NPERM))


perm_gene = True
perm_ct= False
perm_rs = False
group_perm_gene = False
rand_init = False
results.append(permutation_test(path,perm_gene = perm_gene,perm_ct= perm_ct,perm_rs = perm_rs,group_perm_gene = group_perm_gene,rand_init = rand_init,
                 nperm =NPERM))


combined = pd.concat(results, axis = 0,ignore_index = True)

combined.to_pickle('../data/downstream/Nature_tabsap_perm_results_redo')


# %% [code]

combined = pd.read_pickle('../data/downstream/Nature_tabsap_perm_results_redo')
a = combined.groupby(['perm','perm_gene','perm_ct','group_perm_gene','perm_rs','rand_init','gene'])['auc'].mean().reset_index()


keys = ['perm_gene','perm_ct','group_perm_gene','perm_rs','rand_init']
perm_conds = a.groupby(keys).groups
pvalues = {}
avgs = {}
for cond, inds in perm_conds.items():
    cond_perms = a.loc[inds]
    cond_avg = cond_perms.groupby('perm')['auc'].mean()
    key = [a for a,b in zip(keys, cond) if b==True][0]
#     pvalues[key] = 1- (np.abs(cond_avg[0]-.5)>np.abs(.5 - cond_avg[1:])).mean()
    pvalues[key] = 1-(cond_avg[0]> cond_avg.iloc[1:]).mean()
    avgs[key] = a.loc[inds]
    avgs[key]['perm_key'] = key
    

res = pd.concat(avgs).groupby(['perm_key','perm'])['auc'].mean().reset_index()
# one_cond = a.loc[perm_conds[(True, False, False, False, False)]]
# plt.hist(one_cond[one_cond['perm']==0]['auc'], color = 'darkgreen')
# plt.show()

print(pvalues)
plt.figure(figsize = (5,5))
test_stat = res['auc'].iloc[0]
order = ['perm_rs', 'perm_ct', 'perm_gene', 'group_perm_gene']
dat = sns.stripplot(data = res[res['perm']!=0], x = 'perm_key', y = 'auc',
                   s = 1.5, color = 'grey',order = order)
# plt.xticks(rotation = 90);
plt.hlines(test_stat, -0.2, 3.2,colors = 'red')
plt.title(pvalues,fontsize = 5)
plt.hlines(res.groupby('perm_key')['auc'].mean().loc[order], np.arange(4)-0.1, np.arange(4)+0.1,colors = 'black',zorder = 20)


# %% [code]
one_cond = a.loc[perm_conds[(True, False, False, False, False)]]
ttest_1samp(one_cond[one_cond['perm']==0]['auc'],0.5)

# %% [code]
plt.figure(figsize = (1,3))
df = one_cond[one_cond['perm']==0]
sns.stripplot(df['auc'], dodge = True,s = 2.5, c = 'darkgreen')
plt.hlines(df['auc'].mean(), -0.2, 0.2, colors = 'black')
plt.title('finemapping auc')

# %% [code]
```
