## Supplementary material for "Deriving genetic codes for molecular phenotypes from first principles": Code: Fig4_PBMC_allsnps.ipynb

```
# %% [code]
import pandas as pd
import torch
import numpy as np
from tqdm import notebook
from torch import nn
from models.PSALearners import LatentPSALearner
from sklearn.metrics import roc_auc_score
from tqdm.notebook import tqdm

import seaborn as sns
from matplotlib import pyplot as plt

import pickle
DEVICE =6

# %% [code]
eqtls = pd.read_pickle('../data/downstream/eqtls')
eqtl_cols = eqtls.iloc[:,:20]

# %% [code]
eqtls.head()

# %% [code]
len(t2t_eqtls)

# %% [code]
with open('../data/downstream/eqtl_t2tpos','rb') as f:
    t2t_eqtls = pickle.load(f)

# %% [code]
CP_chroms = ['CP068277.2',
'CP068276.2',
'CP068275.2',
'CP068274.2',
'CP068273.2',
'CP068272.2',
'CP068271.2',
'CP068270.2',
'CP068269.2',
'CP068268.2',
'CP068267.2',
'CP068266.2',
'CP068265.2',
'CP068264.2',
'CP068263.2',
'CP068262.2',
'CP068261.2',
'CP068260.2',
'CP068259.2',
'CP068258.2',
'CP068257.2',
'CP068256.2',
'CP068255.2',
'CP086569.2']
chroms = ['chr{}'.format(j) for j in list(range(1,23))+['X','Y']]
chrom_map = {cp:ch for cp,ch in zip(chroms,CP_chroms)}
cpchrom_to_chr = {ch:cp for cp,ch in zip(chroms,CP_chroms)}
t2t_pos = {k:int(v[1]) for k,v in t2t_eqtls.items()if v!='NA' }
t2t_chr = {k:cpchrom_to_chr[v[0]] for k,v in t2t_eqtls.items() if v!='NA'}

# %% [code]


# unique_rsid = eqtl_cols['RSID'].unique()

# with open('../data/downstream/eqtl_rsid', 'wb') as f:
#     pickle.dump(unique_rsid.tolist(), f)

#map the cell types from sceqtl study to pbmc_atlas
cell_type_map = {'Classic Monocyte':  'CD14 Mono',
'Non-classic Monocyte': 'CD16 Mono',
'Naïve/Immature B Cell':'B naive',
'Memory B Cell':'B memory',
'Plasma Cell':'Plasmablast',
'CD8 Effector memory':'CD8 TEM',
'CD4 Effector memory/TEMRA':'CD4 TEM',
'Natural Killer Cell':'NK'}
sat_to_eq = {v:k for k,v in cell_type_map.items()}

eqtl_cols['SIG'] = eqtl_cols['FDR']<0.1
eqtl_cols['NOT_SIG'] = eqtl_cols['FDR']>0.5


# %% [code]
renamed = pd.read_pickle('../data/downstream/pbmc_renamed')
expressed_genes = set(renamed.columns[renamed.max(axis = 0)>0])
print(len(expressed_genes))
ct_gt = eqtl_cols[(eqtl_cols['CELL_TYPE'].isin(renamed.index.values))  & (eqtl_cols['GENE'].isin(expressed_genes))]

# %% [code]
lifted.head()

# %% [code]
lifted['RSID'].drop_duplicates()

# %% [code]
4591503/4603355

# %% [code]
ct_gt['RSID'].drop_duplicates()

# %% [code]
ct_gt['t2t_pos'] = ct_gt['RSID'].map(t2t_pos)
ct_gt['t2t_chr'] = ct_gt['RSID'].map(t2t_chr)
lifted = ct_gt[~ct_gt['t2t_pos'].isna()]

lifted['t2t_ind'] = (lifted['t2t_pos']//1000).astype(int)
lifted['effect_size'] = lifted['SPEARMANS_RHO'].abs()

# %% [code]
chr_names = np.array(list(range(1,23))+['X'])
chrom_lens = np.cumsum([np.load('../data/seq/GENA_2k_precomputed/chr{}_precomputed_features.npy'.format(i),mmap_mode = 'r').shape[0] for i in chr_names])
lens = pd.Series({chrom: v for chrom,v in zip(chroms, [0]+chrom_lens.tolist())})

# %% [code]
lifted['SIG'] = lifted['SIG'].astype(float)

# %% [code]
bin_effect_sizes = lifted.groupby(['t2t_chr','t2t_ind','GENE','CELL_TYPE'])[['effect_size','SIG']].max().reset_index()
bin_effect_sizes['overall_ind'] = bin_effect_sizes['t2t_chr'].map(lens) + bin_effect_sizes['t2t_ind']

# %% [code]
from models.PSALearners import LatentPSALearner
def load_checkpoint(cp):
    checkpoint = torch.load(cp,map_location = 'cpu')
    n_control = checkpoint['learner_state_dict']['0.sc_q.1.weight'].shape[0]
    psa = nn.ModuleList([LatentPSALearner(sc = torch.from_numpy(renamed.values),genome = genome,n_control = n_control,dropout = False,softmax = True)])
    psa.load_state_dict(checkpoint['learner_state_dict'])
    psa.eval()
    return psa[0]

# %% [code]
genome = torch.from_numpy(np.load('../data/seq/GENA_2k_precomputed/concatenated_genome.npy')/10).float() # same as training

# %% [code]
torch.cuda.set_device(6)

# %% [code]

psa = load_checkpoint('results/apr5/metacell_fixed_seq_targ/repeat_0_epoch_76.pt').cuda()


# %% [code]
seq_inds = bin_effect_sizes['overall_ind'].drop_duplicates()

inds_map = {j:i for i,j in enumerate(seq_inds.values)}
seq_batch = genome[seq_inds.values]

# %% [code]
with torch.no_grad():
    ks = psa.seq_k(seq_batch.cuda()).cpu()

# %% [code]
ct_map = {j:i for i,j in enumerate(renamed.index.values)}
with torch.no_grad():
    ct_q = psa.sc_q(torch.from_numpy(renamed.values).float().cuda()).cpu()
    ct = torch.from_numpy(renamed.values).float()

# %% [code]
gene_inds = {k:i for i,k in enumerate(renamed.columns)}

qgenes = bin_effect_sizes['GENE'].map(gene_inds).drop_duplicates()


# %% [code]
qgene_map = {j:i for i,j in enumerate(qgenes.values)}
with torch.no_grad():
    delta = torch.zeros(len(qgenes),renamed.shape[1])
    delta[(np.arange(len(qgenes)), qgenes.values)] += 0.0001

# %% [code]
bin_effect_sizes['gene_index'] = bin_effect_sizes['GENE'].map(gene_inds)
bin_effect_sizes['gene_batch_index'] = bin_effect_sizes['gene_index'].map(qgene_map)
bin_effect_sizes['ct_batch_index'] = bin_effect_sizes['CELL_TYPE'].map(ct_map)
bin_effect_sizes['seq_batch_index'] = bin_effect_sizes['overall_ind'].map(inds_map)


# %% [code]
with torch.no_grad():
    gd = torch.from_numpy(renamed.values[:,bin_effect_sizes['gene_index'].unique()].T).float().cuda()
    gd /= torch.norm(gd, dim = 1, keepdim = True)
    gd = np.maximum(0,(1-torch.abs(gd@gd.T)).cpu().numpy())
    

# %% [code]
from sklearn.neighbors import kneighbors_graph
from scipy.sparse.csgraph import connected_components

# %% [code]
np.random.seed(2322)
knng = kneighbors_graph(gd,n_neighbors = 1, metric = 'precomputed')
concoms = connected_components(knng)[1]
orig_batch = [np.where(concoms==i)[0] for i in np.unique(concoms)]
shuf_batch = [np.where(concoms==i)[0] for i in np.unique(concoms)]
for arr in shuf_batch:
    np.random.shuffle(arr)

shuf3 = {k:v for k,v in zip(np.concatenate(orig_batch), np.concatenate(shuf_batch))}
bin_effect_sizes['shuf3_gb_index'] = bin_effect_sizes['gene_batch_index'].map(shuf3)

# %% [code]
with torch.no_grad():
    inp = ((ct[:,None,:] + delta[None,:,:]).cuda()).reshape(-1,20729)
    
    all_pairs = torch.cat([psa.sc_q(inp[idx]) for idx in np.array_split(np.arange(len(inp)), 10000) ]).view(len(ct),len(delta),-1).cpu()

    
    
    

# %% [code]
@torch.no_grad()
def get_vectorized_discs(gene_batch_inds, seq_batch_inds, ct_batch_inds):
    discs = []
    for inds in np.array_split(np.arange(len(gene_batch_inds)), 100):
        gi = gene_batch_inds[inds]
        si = seq_batch_inds[inds]
        ci = ct_batch_inds[inds]
        new_pos = all_pairs[(ci,gi)]
        discs.append(torch.abs(torch.tanh((new_pos*ks[si]).sum(axis = -1)) - torch.tanh((ct_q[ci]*ks[si]).sum(axis = -1))).numpy())
        
    return np.concatenate(discs)

# %% [code]
all_pairs.shape

# %% [code]
np.random.seed(2322)
knng = kneighbors_graph(gd,n_neighbors = 1, metric = 'precomputed')
concoms = connected_components(knng)[1]

# %% [code]
nperm = 200
out = np.zeros((1+4*nperm,len(bin_effect_sizes)))
orig_batch = [np.where(concoms==i)[0] for i in np.unique(concoms)]
shuf_batch = [np.where(concoms==i)[0] for i in np.unique(concoms)]
nn_cols = ['nn_{}'.format(i) for i in range(nperm)] 
all_cols = ['all_{}'.format(i) for i in range(nperm)]
nn_ct_cols = ['nn_ct_{}'.format(i) for i in range(nperm)]
all_ct_cols = ['all_ct_{}'.format(i) for i in range(nperm)]
perm_cols = ['obs'] + nn_cols + all_cols + nn_ct_cols + all_ct_cols

for N in tqdm(range(1+nperm)):
    if N>0:
        for arr in shuf_batch:
            np.random.shuffle(arr)

    shuf = {k:v for k,v in zip(np.concatenate(orig_batch), np.concatenate(shuf_batch))}
    gene_index = bin_effect_sizes['gene_batch_index'].map(shuf)
    out[N] = get_vectorized_discs(gene_index.values,bin_effect_sizes['seq_batch_index'].values,bin_effect_sizes['ct_batch_index'].values)
    
    
orig_batch = np.arange(len(gd))
shuf_batch = np.arange(len(gd))
for N in tqdm(range(nperm)):
    np.random.shuffle(shuf_batch)
    gene_index = shuf_batch[bin_effect_sizes['gene_batch_index'].values]
    out[nperm+1+N] = get_vectorized_discs(gene_index,bin_effect_sizes['seq_batch_index'].values,bin_effect_sizes['ct_batch_index'].values)

orig_cts = [[0,1],[2,3,4],[5,6,7]]
shuf_cts = [[0,1],[2,3,4],[5,6,7]]
for N in tqdm(range(nperm)):
    for arr in shuf_cts:
        np.random.shuffle(arr)
    shuf = {k:v for k,v in zip(np.concatenate(orig_cts), np.concatenate(shuf_cts))}
    ct_index = bin_effect_sizes['ct_batch_index'].map(shuf).values
    out[2*nperm+1+N] = get_vectorized_discs(bin_effect_sizes['gene_batch_index'].values,bin_effect_sizes['seq_batch_index'].values,ct_index)

    
orig_cts = np.arange(8)
shuf_cts = np.arange(8)
for N in tqdm(range(nperm)):
    np.random.shuffle(shuf_cts)
    ct_index = shuf_cts[bin_effect_sizes['ct_batch_index'].values]
    out[3*nperm+1+N] = get_vectorized_discs(bin_effect_sizes['gene_batch_index'].values,bin_effect_sizes['seq_batch_index'].values,ct_index)


# %% [code]
perms_added = pd.concat([bin_effect_sizes,pd.DataFrame(out[:len(perm_cols)].T)],axis =1 , ignore_index = True)
perms_added.columns = list(bin_effect_sizes.columns) + perm_cols
counts = perms_added.groupby(['GENE','CELL_TYPE']).size()
stds = perms_added.groupby(['GENE','CELL_TYPE'])[perm_cols].std()
means = perms_added.groupby(['GENE','CELL_TYPE'])[perm_cols].mean()

# %% [code]
biggest = perms_added.groupby(['GENE','CELL_TYPE']).apply(lambda w: (w[perm_cols].iloc[np.argsort(w['effect_size'].values)[-3:]].mean(axis = 0)))

# %% [code]
means.shape

# %% [code]
stds.shape

# %% [code]
biggest.shape

# %% [code]
wm = means.copy().reset_index()
wm['dat'] = 'mean'
ws = stds.copy().reset_index()
ws['dat'] = 'std'
wb = biggest.copy().reset_index()
wb['dat'] = 'top3'
pd.concat([wm,ws,wb],axis = 0, ignore_index = True).set_index(['dat','GENE','CELL_TYPE']).to_pickle('../data/downstream/nature_effect_sizes_583_data')


# %% [code]
from scipy.stats import ttest_1samp

# %% [code]
res = {}
ngenes = {}
for ncount in np.arange(5,200,1):
    leads = (biggest[counts>ncount] - means[counts>ncount])/stds[counts>ncount]
    leadz = leads.reset_index().groupby(['GENE'])[perm_cols].mean()


#     res[ncount,'nn'] = (leadz[nn_cols].values >= leadz['obs'].values[:,None]).mean(axis = 1)#list(ttest_1samp(leadz[col],0))
#     res[ncount,'nn_ct'] = (leadz[nn_ct_cols].values >= leadz['obs'].values[:,None]).mean(axis = 1)#list(ttest_1samp(leadz[col],0))
#     res[ncount, 'all'] = (leadz[all_cols].values >= leadz['obs'].values[:,None]).mean(axis = 1)#list(ttest_1samp(leadz[col],0))
#     res[ncount, 'all_ct'] = (leadz[all_ct_cols].values >= leadz['obs'].values[:,None]).mean(axis = 1)#list(ttest_1samp(leadz[col],0))
    ngenes[ncount] = len(leadz)
    res[ncount,'nn'] = (leadz[nn_cols].median(axis =0).values >= leadz['obs'].median()).mean()
    res[ncount,'nn_ct'] = (leadz[nn_ct_cols].median(axis =0).values >= leadz['obs'].median()).mean()
    res[ncount, 'all'] = (leadz[all_cols].median(axis =0).values >= leadz['obs'].median()).mean()
    res[ncount, 'all_ct'] = (leadz[all_ct_cols].median(axis =0).values >= leadz['obs'].median()).mean()

# %% [code]
es_pvals = (-np.log10(np.maximum(1/201, pd.Series(res))).unstack())
es_pvals.plot(legend = False,figsize = (5,5))

plt.hlines(-np.log10(0.05), 0, 200, colors  = 'grey')
idx = np.arange(5,200,10)
plt.title('effect size pvals, y[0,2,5]')
plt.yticks(np.linspace(0,2,5),[]);

plt.show()

plt.figure(figsize = (5,1))
plt.plot(idx, pd.Series(ngenes).loc[idx],c = 'black')
plt.yticks(np.linspace(2500,5000,5),[]);
plt.title('numgenes, y[2.5k,5k,500]')
plt.xticks([])
plt.show()

# %% [code]
res = {}
groups = bin_effect_sizes.reset_index().groupby(['GENE','CELL_TYPE']).groups
y_trues = bin_effect_sizes['SIG'].astype(int).values
for g,v in tqdm(groups.items(),total = len(groups)):
    if len(v)<3:
        continue
    y_true = y_trues[v]
    ytm = y_true.mean()
    if max(ytm,1-ytm)>0.99:
        continue
    res[g] = [roc_auc_score(y_true, y_score = out[j][v]) for j in range(out.shape[0])]
auc_df = pd.DataFrame(res).T
auc_df.columns = perm_cols
ct_avg_aucs = auc_df.reset_index().groupby('level_0')[perm_cols].mean()

# %% [code]
auc_df.to_pickle('../data/downstream/nature_allauc_results')

# %% [code]
pvals = {}
ngene_auc = {}
obss_auc = {}
for num_min in range(5,200,1):
    sub_df = auc_df.loc[[k for k in auc_df.index.values if len(groups[k])>num_min]]
    ngene_auc[num_min] = len(sub_df.index.values)
    flstats = sub_df.reset_index().groupby('level_0').mean(numeric_only=True).median(axis=0)
    obs = flstats.loc['obs']
    obss_auc[num_min] = obs.copy()
    stats =flstats.iloc[1:].reset_index()
    stats['type'] = ['_'.join(a[:-1]) for a in stats['index'].str.split('_')]
    pvals[num_min] = stats.groupby('type').apply(lambda w: (w[0]>=obs).mean()).sort_values()

# %% [code]

(-np.log10(np.maximum(1/201, pd.DataFrame(pvals) )  )).T.plot(legend = None,figsize = (5,5))
plt.hlines(-np.log10(0.05), 0, 200, colors  = 'grey')
idx = np.arange(5,200,10)
plt.title('auc pvals, y[0,2,5]')
plt.yticks(np.linspace(0,2,5),[]);

plt.show()

plt.figure(figsize = (5,1))
plt.plot(idx, pd.Series(ngene_auc).loc[idx],c = 'black')
plt.yticks(np.linspace(500,2000,5),[]);
plt.title('numgenes, y[500,2000,5]')
plt.xticks([])
plt.show()

# %% [markdown]
### Plotting the figures

# %% [code]


# %% [code]
## effect sizes
plt.figure(figsize = (4,4))
ncount = 100
leads = (biggest[counts>ncount] - means[counts>ncount])/stds[counts>ncount]
leadz = leads.reset_index().groupby(['GENE'])[perm_cols].mean()

plot_es = pd.DataFrame(leadz.median(axis = 0))
obs_es = plot_es.loc['obs']
plot_es['perm_type'] = 'obs'
plot_es['perm_type'].loc[nn_cols] = 'grouped gene'
plot_es['perm_type'].loc[all_cols] = 'gene'
plot_es['perm_type'].loc[all_ct_cols] = 'ct'
plot_es['perm_type'].loc[nn_ct_cols] = 'grouped ct'

order = ['ct', 'grouped ct', 'gene', 'grouped gene']
sns.stripplot(data = plot_es.iloc[1:], x = 'perm_type', y = 0, c = 'grey', s = 2.5, order = order)
plt.ylabel('normalized es increase on lead eqtls')
plt.xlabel('permutation type')
plt.hlines(plot_es.groupby('perm_type').mean().loc[order], np.arange(4)-0.2, np.arange(4)+0.2, colors = 'black',zorder = 5)
plt.hlines(obs_es,-0.2, 3.2, colors = 'red')
plt.title((10**(-(es_pvals.loc[ncount]))).to_dict(),fontsize = 5)

# %% [code]
#aucs
plt.figure(figsize = (4,4))
num_min = 10
sub_df = auc_df.loc[[k for k in auc_df.index.values if len(groups[k])>num_min]]
ngene_auc[num_min] = len(sub_df.index.values)


plot_auc = pd.DataFrame(sub_df.reset_index().groupby('level_0').mean(numeric_only=True).median(axis=0))
obs_auc = plot_auc.loc['obs']
plot_auc['perm_type'] = 'obs'
plot_auc['perm_type'].loc[nn_cols] = 'grouped gene'
plot_auc['perm_type'].loc[all_cols] = 'gene'
plot_auc['perm_type'].loc[all_ct_cols] = 'ct'
plot_auc['perm_type'].loc[nn_ct_cols] = 'grouped ct'

order = ['ct', 'grouped ct', 'gene', 'grouped gene']
sns.stripplot(data = plot_auc.iloc[1:], x = 'perm_type', y = 0, c = 'grey', s = 2.5, order = order)
plt.ylabel('normalized auc')
plt.xlabel('permutation type')
plt.hlines(plot_auc.groupby('perm_type').mean().loc[order], np.arange(4)-0.2, np.arange(4)+0.2, colors = 'black')
plt.hlines(obs_auc,-0.2, 3.2, colors = 'red')

plt.title(pvals[num_min].to_dict(),fontsize = 5)

# %% [code]
```
