## Supplementary material for "Deriving genetic codes for molecular phenotypes from first principles": Code: Fig4_PBMC_CRISPR_IFNG.ipynb

```
# %% [code]
import pandas as pd
import torch
import numpy as np
from tqdm import notebook
from torch import nn
from models.PSALearners import LatentPSALearner
from sklearn.metrics import roc_auc_score
from tqdm.notebook import tqdm

import seaborn as sns
from matplotlib import pyplot as plt

import pickle
import itertools


# %% [code]
hits = pd.read_excel('../data/downstream/science.abj4008_table_s2.xlsx')

# %% [code]
crisprcd8 = hits[ (hits['CD4_or_CD8']=='CD8')]

# %% [code]
#download the screen data
# hits = pd.read_excel('../data/downstream/science.abj4008_table_s2.xlsx')
# crisprcd8 = hits[ (hits['CD4_or_CD8']=='CD8')]
# crisprcd8[crisprcd8['CRISPRa_or_i']=='CRISPRa'].loc[:,['Gene','zscore']].groupby('Gene').mean().to_pickle('../data/downstream/cd8_ifng_scores')

scores = pd.read_pickle('../data/downstream/cd8_ifng_scores')

# preparing the gene information
genes = pd.read_table('../data/seq/GENA_2k_precomputed/hs1.110.20220412.ncbiRefSeq.gtf',header = None)

#map CP chroms to chrs
CP_chroms = ['CP068277.2', 'CP068276.2', 'CP068275.2', 'CP068274.2', 'CP068273.2', 'CP068272.2', 'CP068271.2', 'CP068270.2', 'CP068269.2', 'CP068268.2', 'CP068267.2', 'CP068266.2', 'CP068265.2', 'CP068264.2', 'CP068263.2', 'CP068262.2', 'CP068261.2', 'CP068260.2', 'CP068259.2', 'CP068258.2', 'CP068257.2', 'CP068256.2', 'CP068255.2', 'CP086569.2']
chroms = ['chr{}'.format(j) for j in list(range(1,23))+['X','Y']]
chrom_map = {cp:ch for cp,ch in zip(chroms,CP_chroms)}
cp_to_chr = {cp:ch for cp,ch in zip(CP_chroms,chroms)}

#select TSS positions
transcripts = genes[genes[2]=='transcript']
transcripts['id'] =[a.split(';')[0].split(' ')[-1][1:-1] for a in transcripts[8]]
transcripts['chrom'] = transcripts[0].replace(cp_to_chr)
data = transcripts[['chrom',3,4,6,'id']].drop_duplicates()
data = data[data['chrom']!='chrY']
data.columns = ['chrom', 'start', 'stop', 'strand', 'gene_symbol']
data['tss'] = [data['start'].iloc[i] if (s== '+') else data['stop'].iloc[i] for i,s in enumerate(data['strand'])]
data = data.sort_values(['chrom','tss'])
chrs = {chrom:data.loc[val] for chrom,val in data.groupby('chrom').groups.items()}


#build the batch of sequence features
chroms = list(range(1,23)) + ['X']
output_features = []
gene_symbols = []


for chrom_num in tqdm(chroms):
    chrom = 'chr{}'.format(chrom_num)
    chr_dat = np.load('../data/seq/GENA_2k_precomputed/{}_precomputed_features.npy'.format(chrom), mmap_mode = 'r')
    starts = (chrs[chrom]['tss']//1000)
    output_features.append(np.stack([chr_dat[starts+i] for i in range(-2,3)],axis = 1)/50)
#     output_features.append(np.stack([chr_dat[starts]],axis = 1))
    gene_symbols+= (chrs[chrom]['gene_symbol'].tolist())
output_features = np.concatenate(output_features,axis =0)
# seq_batch = torch.from_numpy(output_features).to('cuda:{}'.format(DEVICE)).float().reshape(-1, output_features.shape[2])

# %% [code]
(scores.loc[common_scores].abs()>4).mean()

# %% [code]
common_scores = sorted(set(gene_symbols).intersection(scores.index.values))
q = scores.loc[common_scores]>2
inds= common_scores.copy()
np.random.seed(1234)
np.random.shuffle(inds)
val_genes = inds[:4000]
test_genes = inds[4000:]

np.random.seed(15)
np.random.shuffle(val_genes)    
val1 = val_genes[:1000]
val2 = val_genes[1000:2000]
val3 = val_genes[2000:3000]
val4 = val_genes[3000:]
from sklearn.metrics import roc_curve

def get_aucs(preds,return_curve = False):
    '''
    input: predictions series (indexed by gene symbol)
    output: aucs for the val splits and the overall
    '''
 
    if return_curve:
        return (roc_auc_score(y_true=q, y_score=preds), roc_curve(q, preds))
    return roc_auc_score(y_true=q, y_score=preds)

def znorm(df,axis = 0):
    a = df.values
    return pd.DataFrame((a - a.mean(axis =axis,keepdims = True))/(1e-6+ a.std(axis = axis,keepdims = True)), index = df.index.values, columns = df.columns)

@torch.no_grad()
def eval_learner(learner,sc_batch,return_preds = False):
    learner.eval()
        

    T_xs = learner.T_xs(seq_batch=seq_batch,sc_batch = sc_batch)

    abs_grads = torch.abs(T_xs[1]-T_xs[0])

    tss_mean_abs_grad = abs_grads.reshape(-1,output_features.shape[1]).mean(axis = -1)

    grad_scores = pd.DataFrame({'abs_grad':tss_mean_abs_grad.detach().cpu().numpy(), 
              'gene':gene_symbols}).groupby('gene').mean()
    
    return grad_scores

def get_sc_batch(cd8_mean,shift_size = 1e-4,gene = 'IFNG'):
    orig = torch.from_numpy(cd8_mean.values).float().to('cuda:{}'.format(DEVICE))
    shift = cd8_mean.copy()
    shift[gene] += shift_size
    shifted = torch.from_numpy(shift.values).float().to('cuda:{}'.format(DEVICE))
    return torch.cat([orig[None,:],shifted[None,:]],0)

# %% [code]
from sklearn.decomposition import PCA

# %% [code]
pca = PCA(n_components = 40)

# %% [code]
pc_space = pca.fit_transform(output_features.reshape(-1, output_features.shape[-1])).reshape(*(list(output_features.shape[:2])+[40]))

# %% [code]
pc_space.shape

# %% [code]
import pyranges as pr

# %% [code]
hg38_canon = pd.read_table('nCP6(2).bed',comment = '#',header = None)

# %% [code]
hg38_canon = hg38_canon[hg38_canon[5].isin(common_scores) & hg38_canon[0].isin(pd.read_table('canonical_tx.bed',header = None,comment = '#').values.flatten())]

# %% [code]
hg38_canon['tss'] = [a if strand =='+' else b for a,b,strand in zip(hg38_canon[3],hg38_canon[4], hg38_canon[2])]

# %% [code]
from pyfaidx import Fasta

# %% [code]
hg38 = Fasta('../data/seq/hg38.fa')

# %% [code]
import re

# %% [code]
gcs = {}
for chrom, start,stop,index in tqdm(zip(hg38_canon[1],hg38_canon['tss']-5000, hg38_canon['tss']+5000,hg38_canon.index),total = len(hg38_canon)):
    st = str(hg38[chrom][start:stop]).upper()
    if len(st)>0:
        
        gcs[index] = len(re.findall('G|C',st))/len(st)

# %% [code]
hg38_canon['tss_gc'] = pd.Series(gcs)
gcs = hg38_canon.groupby(5)['tss_gc'].mean()

# %% [code]
seq_batch = torch.from_numpy(output_features).to('cuda:{}'.format(2)).float().reshape(-1, output_features.shape[2])

# %% [code]
n_learn=1
l1_means = pd.read_pickle('../data/downstream/pbmc_l1_means')
DEVICE =2
multi_learner = nn.ModuleList([LatentPSALearner(sc = torch.from_numpy(l1_means.values), softmax=True, dropout = 0, genome = seq_batch, n_control = 40, bias = False,batch_norm = False) for _ in range(n_learn)]).to('cuda:{}'.format(DEVICE))

path  = 'results/apr5/metacell_fixed_seq_targ/repeat_0_epoch_76.pt'
checkpoint = torch.load(path,map_location = 'cpu')
sd = checkpoint['learner_state_dict']
multi_learner.load_state_dict(sd)
learner = multi_learner[0]
ifng_aucs = {}
ifng_preds = {}
sel_genes = sorted(set(library['Regulation Of Type II Interferon-Mediated Signaling Pathway (GO:0060334)']).intersection(l1_means.columns))
for gene in tqdm(l1_means.columns):
    y_pred = eval_learner(learner = learner, sc_batch = get_sc_batch(l1_means.loc['CD8 T'],0.0001,gene=gene)).loc[common_scores].copy()
    ifng_preds[gene] = y_pred.copy()['abs_grad']
    ifng_aucs[gene]=  get_aucs(y_pred,return_curve = False)

# %% [markdown]
## Compute scores at different thresholds

# %% [code]
library_21 = blitz.enrichr.get_library("GO_Biological_Process_2021")

# %% [code]
q = scores.loc[common_scores]>4
ifng_aucs4 = {}
for k,v in tqdm(ifng_preds.items()):
    ifng_aucs4[k] = roc_auc_score(y_score = ifng_preds[k], y_true = q)
    

# %% [code]
q = scores.loc[common_scores]>3
ifng_aucs3 = {}
for k,v in tqdm(ifng_preds.items()):
    ifng_aucs3[k] = roc_auc_score(y_score = ifng_preds[k], y_true = q)
    

# %% [code]
q = scores.loc[common_scores]>2
ifng_aucs2 = {}
for k,v in tqdm(ifng_preds.items()):
    ifng_aucs2[k] = roc_auc_score(y_score = ifng_preds[k], y_true = q)
    

# %% [code]
q = scores.loc[common_scores].abs()>3
ifng_aucs_abs= {}
for k,v in tqdm(ifng_preds.items()):
    ifng_aucs_abs[k] = roc_auc_score(y_score = ifng_preds[k], y_true = q)
    

# %% [code]
from statsmodels.stats.multitest import fdrcorrection

# %% [code]
gs_rocs = {}
q = scores.loc[common_scores]>3
lib2 = {k:sorted(set(v).intersection(ifng_preds.keys())) for k,v in library.items() if (len(v)<100)}
for go,gs in tqdm(lib2.items()):
    if len(gs)<=5:
        continue
        
    w = pd.concat({k:v for k,v in ifng_preds.items() if k in gs}).unstack()
    gs_rocs[go] = roc_auc_score(y_score = pd.DataFrame(rankdata(w,axis=1),index = w.index, columns = w.columns).mean(axis=0),y_true=q.loc[w.columns])

# %% [code]
all_rocs = pd.Series(gs_rocs)

# %% [code]
(all_rocs>all_rocs.filter(regex='Interfer').sort_values().iloc[-1]).mean()

# %% [code]
pd.Series(ifng_aucs).hist()

# %% [code]
datt = pd.DataFrame({'exp':l1_means.loc['CD8 T'], 'auc':pd.Series(ifng_aucs4)})
sns.scatterplot(data = datt, x = 'exp',y = 'auc')
spearmanr(datt['exp'],datt['auc'])

# %% [code]
av_exp = {}
lib2 = {k:sorted(set(v).intersection(ifng_preds.keys())) for k,v in library.items() if (len(v)<100)}
for go,gs in tqdm(lib2.items()):
    if len(gs)<=5:
        continue
    av_exp[go] = l1_means.loc['CD8 T'].reindex(gs).dropna().mean()
    

# %% [code]
sel_genes = library['Regulation Of Type II Interferon-Mediated Signaling Pathway (GO:0060334)']
w = pd.Series(ifng_aucs3)
w = pd.DataFrame({'auc': w, 'ifn':w.index.isin(sel_genes)})
w['ifn'] = w['ifn'].astype('category')
w['size'] = w['ifn'].map({True:200, False:1})
plt.figure(figsize = (5,10))


ax =sns.stripplot(data = w, hue = 'ifn',y = 'auc',x = 'ifn',dodge = True,palette = ['grey','red'],s = 1)
ax =sns.stripplot(data = w[w['ifn']], hue = 'ifn',y = 'auc',x = 'ifn',dodge = True,palette = ['grey','red'],s = 5)
ax.legend_.remove()

# %% [code]


# %% [code]
plt.figure(figsize = (5,5))
datt = pd.DataFrame({'exp':av_exp, 'auc':gs_rocs})
sns.scatterplot(data = datt, x = 'exp',y = 'auc',s = 2.5, c= 'grey')

# %% [code]
pd.Series(gs_rocs).filter(regex = 'Regulation Of Type II Interferon').sort_values()

# %% [code]
# pd.Series(gs_rocs).filter(regex = 'Type II Interferon').sort_values()
top_gs = pd.Series(gs_rocs).sort_values().iloc[-10:].index
    

# %% [code]
set(library['Vacuolar Acidification (GO:0007035)']).intersection(library['Programmed Necrotic Cell Death (GO:0097300)'])

# %% [code]
lm_pvalues = {}
rocs = {}

# sel_genes = sorted(set(library['Positive Regulation Of Type II Interferon Production (GO:0032729)']).intersection(l1_means.columns))
# sel_genes = sorted(set(library['Regulation Of Type II Interferon-Mediated Signaling Pathway (GO:0060334)']).intersection(l1_means.columns))

sel_genes = sorted(set(library['Regulation Of Type II Interferon-Mediated Signaling Pathway (GO:0060334)']).intersection(l1_means.columns))
# sel_genes = sorted(set(library['Programmed Necrotic Cell Death (GO:0097300)']).intersection(l1_means.columns))
# sel_genes = ['IFNG']
for t in np.linspace(2,4,20):
    q = scores.loc[common_scores]>t
    # w = pd.concat(ifng_preds).unstack()
    w = pd.concat({k:v for k,v in ifng_preds.items() if k in sel_genes}).unstack()
    psa_roc = roc_auc_score(y_score = pd.DataFrame(rankdata(w,axis=1),index = w.index, columns = w.columns).mean(axis=0),y_true=q.loc[w.columns])
    gc_roc = roc_auc_score(y_true = q.loc[gcs.index], y_score = gcs)
    
    dd = pd.concat([pd.DataFrame(rankdata(w,axis=1),index = w.index, columns = w.columns).mean(axis=0)/1000,np.log10(gcs)],axis=1).dropna()
    X = sm.add_constant(dd)
    y = q.loc[X.index]
    lm2 = Logit(y,X).fit(maxiter = 10000)
    # lm1 = Logit(y,sm.add_constant(dd[['tss_gc']])).fit(maxiter = 10000)
    lm_pvalues[t] = -np.log10(lm2.pvalues.drop('const').rename({0:'psa'}).copy())
    rocs[t] = pd.Series({'psa':psa_roc, 'gc':gc_roc})

# %% [code]
plt.figure(figsize = (5,5))
plot_df = pd.concat(lm_pvalues).unstack()
plot_df['psa'].plot(label = 'psa', color = 'red')
plot_df['tss_gc'].plot(label = 'gc', color = 'grey')
plt.title('multivariate logistic regression pvalues')
plt.xlabel('z-score threshold for hits')
plt.ylabel('-log10 p')
plt.hlines(-np.log10(0.05), 2,4,colors = 'grey',linewidth = 1,linestyle = '--')
plt.show()
plt.figure(figsize = (5,5))
plot_df = pd.concat(rocs).unstack()
plot_df['psa'].plot(label = 'psa', color = 'red')
plot_df['gc'].plot(label = 'gc', color = 'grey')

plt.ylabel('AUROC')
plt.xlabel('z-score threshold for hits')
plt.legend()


# %% [code]
from scipy.stats import spearmanr
from scipy.stats import rankdata

# %% [code]


# %% [code]


# %% [markdown]
###  Null distributions

# %% [code]
lib2 = {k:sorted(set(v).intersection(ifng_preds.keys())) for k,v in library.items() if (len(v)<100)}

# %% [code]
q = scores.loc[common_scores]>4

orig_genes = np.array(list(ifng_preds.keys()))
shuf_genes = orig_genes.copy()
perm_gs_rocs = {}

for perm in tqdm(range(500)):
    if perm>0:
        np.random.shuffle(shuf_genes)
    shuf_dict = {i:j for i,j in zip(orig_genes,shuf_genes)}
    
    for go in top_gs.tolist() + ['Regulation Of Type II Interferon-Mediated Signaling Pathway (GO:0060334)','Cellular Response To Type II Interferon (GO:0071346)']:
        gs = lib2[go]
        
        w = pd.concat({k:v for k,v in ifng_preds.items() if shuf_dict[k] in gs}).unstack()
        perm_gs_rocs[go,perm] = roc_auc_score(y_score = pd.DataFrame(rankdata(w,axis=1),index = w.index, columns = w.columns).mean(axis=0),y_true=q.loc[w.columns])

# %% [code]
w = pd.Series(perm_gs_rocs).unstack()
w = pd.Series((w.iloc[:,1:].values>= w.values[:,0][:,None]).mean(axis = 1),index = w.index)
new_top = w.index[w==0].tolist()

# %% [code]
q = scores.loc[common_scores]>4

orig_genes = np.array(list(ifng_preds.keys()))
shuf_genes = orig_genes.copy()
perm_gs_rocs2 = {}

for perm in tqdm(range(5000)):
    if perm>0:
        np.random.shuffle(shuf_genes)
    shuf_dict = {i:j for i,j in zip(orig_genes,shuf_genes)}
    
    for go in new_top:
        gs = lib2[go]
        
        w = pd.concat({k:v for k,v in ifng_preds.items() if shuf_dict[k] in gs}).unstack()
        perm_gs_rocs2[go,perm] = roc_auc_score(y_score = pd.DataFrame(rankdata(w,axis=1),index = w.index, columns = w.columns).mean(axis=0),y_true=q.loc[w.columns])

# %% [code]
w = pd.Series(perm_gs_rocs2).unstack()
w = pd.Series((w.iloc[:,1:].values>= w.values[:,0][:,None]).mean(axis = 1),index = w.index)


# %% [markdown]
### Here we do the figures
- threshold three

# %% [code]
orig_genes = np.array(list(ifng_preds.keys()))
shuf_genes = orig_genes.copy()
ifng_roc_perm = {}

for t in [3]:
    sel_genes = lib2['Regulation Of Type II Interferon-Mediated Signaling Pathway (GO:0060334)']
    q = scores.loc[common_scores]>t
    for perm in tqdm(range(200)):
        if perm>0:
            sel_genes = np.random.choice(orig_genes,len(sel_genes),replace = False)
        
            
        w = pd.concat({k:ifng_preds[k] for k in sel_genes}).unstack()
        
        ifng_roc_perm[perm,t] = roc_auc_score(y_score = pd.DataFrame(rankdata(w,axis=1),index = w.index, columns = w.columns).mean(axis=0),y_true=q.loc[w.columns])

# %% [code]
n_learn=1
l1_means = pd.read_pickle('../data/downstream/pbmc_l1_means')
DEVICE =2

sel_genes = sorted(set(library['Regulation Of Type II Interferon-Mediated Signaling Pathway (GO:0060334)']).intersection(l1_means.columns))
path  = 'results/apr5/metacell_fixed_seq_targ/repeat_0_epoch_76.pt'
checkpoint = torch.load(path,map_location = 'cpu')
sd = checkpoint['learner_state_dict']
stats = []
q = scores.loc[common_scores]>3
for perm in tqdm(range(200)):
    multi_learner = nn.ModuleList([LatentPSALearner(sc = torch.from_numpy(l1_means.values), softmax=True, dropout = 0, genome = seq_batch, n_control = 40, bias = False,batch_norm = False) for _ in range(n_learn)]).to('cuda:{}'.format(DEVICE))
    if perm==0:
        multi_learner.load_state_dict(sd)
    learner = multi_learner[0]
    y_preds ={}
    for gene in sel_genes:
        y_preds[gene] =eval_learner(learner = learner, sc_batch = get_sc_batch(l1_means.loc['CD8 T'],0.0001,gene=gene)).loc[common_scores]['abs_grad'].copy()
    w = pd.concat({k:y_preds[k] for k in sel_genes}).unstack()
    stats.append(roc_auc_score(y_score = pd.DataFrame(rankdata(w,axis=1),index = w.index, columns = w.columns).mean(axis=0),y_true=q.loc[w.columns]))
        
null_init_stats = np.array(stats)        

# %% [code]
kk = pd.concat({'random_genes':pd.Series(ifng_roc_perm).reset_index()[0],'null_init':pd.Series(null_init_stats),'other_geneset':pd.Series(other_geneset_aucs)}).unstack().T
plt.figure(figsize = (3,5))

sns.stripplot(data = kk.iloc[1:].melt().dropna(), x ='variable',y = 'value', c= 'grey',s = 2)
plt.hlines(kk.iloc[0,0],-.5,2.5,color = 'red')
plt.xticks(rotation = 90);
plt.ylabel('AUC')
pvs = {j:'%.4f'%(kk[j].dropna().iloc[0]<=kk[j].dropna().iloc[1:]).mean() for j in kk.columns}

pvs


# %% [code]
import blitzgsea as blitz
library = blitz.enrichr.get_library("GO_Biological_Process_2023")
```
