## Supplementary material for "Deriving genetic codes for molecular phenotypes from first principles": Code: Fig4_PBMC_eqtl_evaluation.ipynb

```
# %% [markdown]
## GWAS SNP AUCs
- We will do the permutation tests and baselines for GWAS SNP AUC prediction
- We will do corruption as well

# %% [code]
import pandas as pd
import torch
import numpy as np
from tqdm import notebook
from torch import nn
from models.PSALearners import LatentPSALearner
from sklearn.metrics import roc_auc_score
from tqdm.notebook import tqdm

import seaborn as sns
from matplotlib import pyplot as plt

import pickle
import itertools
DEVICE =3

# %% [code]
renamed = pd.read_pickle('../data/downstream/pbmc_renamed')

# %% [code]
eqtls = pd.read_pickle('../data/downstream/eqtls')
eqtl_cols = eqtls.iloc[:,:20]

#map the cell types from sceqtl study to pbmc_atlas
cell_type_map = {'Classic Monocyte':  'CD14 Mono',
'Non-classic Monocyte': 'CD16 Mono',
'Naïve/Immature B Cell':'B naive',
'Memory B Cell':'B memory',
'Plasma Cell':'Plasmablast',
'CD8 Effector memory':'CD8 TEM',
'CD4 Effector memory/TEMRA':'CD4 TEM',
'Natural Killer Cell':'NK'}
sat_to_eq = {v:k for k,v in cell_type_map.items()}

eqtl_cols['SIG'] = eqtl_cols['FDR']<0.1
eqtl_cols['NOT_SIG'] = eqtl_cols['FDR']>0.5


renamed = pd.read_pickle('../data/downstream/pbmc_renamed')
expressed_genes = set(renamed.columns[renamed.max(axis = 0)>0])
print(len(expressed_genes))
ct_gt = eqtl_cols[(eqtl_cols['CELL_TYPE'].isin(renamed.index.values))  & (eqtl_cols['GENE'].isin(expressed_genes))]


dbsnp = pd.read_table('chm13v2.0_GWASv1.0rsids_e100_r2022-03-08.vcf', header = None,sep = '\t', comment='#')
lifted_ctgt = ct_gt[ct_gt['RSID'].isin(dbsnp[2])]
t2t_trans = pd.merge(lifted_ctgt, dbsnp[[0,1,2]].rename({2:'RSID'},axis = 1),on = 'RSID').rename({0:'t2t_chr',1:'t2t_pos'},axis = 1)
t2t_trans['t2t_ind'] = t2t_trans['t2t_pos']//1000


input_df = t2t_trans
input_df['hg38_ind'] = input_df['POS']//1000
# pool the association over segments
pos_df = input_df.groupby(['t2t_chr','t2t_ind','GENE','CELL_TYPE'])['FDR'].min().reset_index()

# select positions with associations evaluated only against a single gene
rs_gene_counts = pos_df[['t2t_chr','t2t_ind','GENE']].drop_duplicates().groupby(['t2t_chr','t2t_ind']).size()
unique_rs = set(rs_gene_counts[rs_gene_counts<2].index.values)
unique_rs_pos = pos_df.iloc[[i for i,a in enumerate(zip(pos_df['t2t_chr'],pos_df['t2t_ind'])) if a in unique_rs]]
max_fdr = unique_rs_pos.groupby(['GENE','CELL_TYPE']).apply(lambda w: (w['FDR']>0.1).sum())

min_fdr = unique_rs_pos.groupby(['GENE','CELL_TYPE']).apply(lambda w: (w['FDR']<0.1).sum())
counts = unique_rs_pos.groupby(['GENE','CELL_TYPE']).size()
# select gene_ct pairs with at least one significant and one not significant qtl
gene_ct_pairs = set(min_fdr[(min_fdr>0)&(max_fdr>0)].index.values)

valid_long = pos_df.iloc[[i for i,a in enumerate(zip(pos_df['GENE'],pos_df['CELL_TYPE']) )if a in gene_ct_pairs]]
valid_long = valid_long[valid_long['GENE'].isin(expressed_genes)]
valid_long = valid_long[valid_long['t2t_chr']!='chrY']


# %% [code]
# preparing the gene information
genes = pd.read_table('../data/seq/GENA_2k_precomputed/hs1.110.20220412.ncbiRefSeq.gtf',header = None)

#map CP chroms to chrs
CP_chroms = ['CP068277.2', 'CP068276.2', 'CP068275.2', 'CP068274.2', 'CP068273.2', 'CP068272.2', 'CP068271.2', 'CP068270.2', 'CP068269.2', 'CP068268.2', 'CP068267.2', 'CP068266.2', 'CP068265.2', 'CP068264.2', 'CP068263.2', 'CP068262.2', 'CP068261.2', 'CP068260.2', 'CP068259.2', 'CP068258.2', 'CP068257.2', 'CP068256.2', 'CP068255.2', 'CP086569.2']
chroms = ['chr{}'.format(j) for j in list(range(1,23))+['X','Y']]
chrom_map = {cp:ch for cp,ch in zip(chroms,CP_chroms)}
cp_to_chr = {cp:ch for cp,ch in zip(CP_chroms,chroms)}

#select TSS positions
transcripts = genes[genes[2]=='transcript']
transcripts['id'] =[a.split(';')[0].split(' ')[-1][1:-1] for a in transcripts[8]]
transcripts['chrom'] = transcripts[0].replace(cp_to_chr)
data = transcripts[['chrom',3,4,6,'id']].drop_duplicates()
data = data[data['chrom']!='chrY']
data.columns = ['chrom', 'start', 'stop', 'strand', 'gene_symbol']
data['tss'] = [data['start'].iloc[i] if (s== '+') else data['stop'].iloc[i] for i,s in enumerate(data['strand'])]
data = data.sort_values(['chrom','tss'])


chrs = {chrom:data.loc[val] for chrom,val in data.groupby('chrom').groups.items()}

# %% [markdown]
## Computing baselines
- Distance to TSS
- GC content
- ENCODE marks
- Supervised classifier (per gene)
- Supervised classifier (overall)

# %% [code]
baseline_aucs = {}

# %% [markdown]
### Distance to TSS

# %% [code]
# need to get tss data from below
df = data.groupby(['gene_symbol','chrom'])['tss'].mean().reset_index().set_index('gene_symbol')
common = set(valid_long['GENE'].values).intersection(df.index.values)
cvl =  valid_long[valid_long['GENE'].isin(common)]

CP_chroms = ['CP068277.2',
'CP068276.2',
'CP068275.2',
'CP068274.2',
'CP068273.2',
'CP068272.2',
'CP068271.2',
'CP068270.2',
'CP068269.2',
'CP068268.2',
'CP068267.2',
'CP068266.2',
'CP068265.2',
'CP068264.2',
'CP068263.2',
'CP068262.2',
'CP068261.2',
'CP068260.2',
'CP068259.2',
'CP068258.2',
'CP068257.2',
'CP068256.2',
'CP068255.2',
'CP086569.2']
chroms = ['chr{}'.format(j) for j in list(range(1,23))+['X','Y']]
chrom_map = {cp:ch for cp,ch in zip(chroms,CP_chroms)}
cpchrom_to_chr = {ch:cp for cp,ch in zip(chroms,CP_chroms)}

# %% [code]
cvl['tss'] = 0
cvl['chr2'] = 'a'
cvl['tss'] = df['tss'].loc[cvl['GENE']].values
cvl['chr2'] = df['chrom'].loc[cvl['GENE']].values


# %% [code]
cvl['diff'] = np.abs(cvl['t2t_ind']-cvl['tss']//1000)
groups = cvl.groupby(['GENE','CELL_TYPE']).groups
res = {}
for k,v in groups.items():
    y_true =cvl['FDR'].loc[v]< 0.1
    y_score = cvl['diff'].loc[v]
    res[k] = roc_auc_score(y_true = y_true, y_score = -y_score)
    
tss_aucs = pd.Series(res).reset_index().groupby('level_0')[0].mean()

# %% [code]
baseline_aucs['distance to tss'] = tss_aucs.mean()

# %% [markdown]
### GC content

# %% [code]
from pyfaidx import Fasta
GENOME = '../GENA_LM/data/genomes/ncbi_dataset/data/GCA_009914755.4/GCA_009914755.4_T2T-CHM13v2.0_genomic.fna'
data = Fasta(GENOME, sequence_always_upper=True)


# %% [code]
gps = valid_long.groupby(['t2t_chr','t2t_ind']).groups
gcs = {}
for chrom, pos in tqdm(gps.keys()):
    seq = data[chrom_map[chrom]][pos:pos+2000].seq
    gc = (seq.count('G') + seq.count('C'))/len(seq)
    gcs[chrom,pos] = gc
gcs = pd.Series(gcs)

mvl = pd.merge(valid_long, gcs.reset_index(),left_on = ['t2t_chr','t2t_ind'], right_on = ['level_0','level_1'])


groups = mvl.groupby(['GENE','CELL_TYPE']).groups
mres = {}
for k,v in groups.items():
    y_true =mvl['FDR'].loc[v]< 0.1
    y_score = mvl[0].loc[v]
    mres[k] = roc_auc_score(y_true = y_true, y_score = -y_score)
    
gc_aucs = pd.Series(mres).reset_index().groupby('level_0')[0].mean()
baseline_aucs['gc content'] = gc_aucs.mean()

# %% [markdown]
## ENCODE marks

# %% [code]
#get the combined encode data
# !wget http://hgdownload.soe.ucsc.edu/gbdb/hg38/encode3/ccre/encodeCcreCombined.bb
# !wget http://hgdownload.soe.ucsc.edu/admin/exe/linux.x86_64/bigBedToBed 
# !chmod a+x bigBedToBed 

# !./bigBedToBed encodeCcreCombined.bb encode.bed

# %% [code]
encode = pd.read_csv('encode.bed', sep='\t', header=None)
dummies = pd.get_dummies(encode[10])
feat_names =list(dummies.columns)

# %% [code]
from numpy.lib.stride_tricks import sliding_window_view

# %% [code]


# %% [code]
cre_aucs = {}
for radius in 1000*np.arange(1,11):
    d = t2t_trans[['CHR','POS']].drop_duplicates() #hg38
    for feat in feat_names:
        d[feat] = False

    for chrom in tqdm(list(range(1,23)) + ['X']):
        dq = d['CHR']==chrom
        if dq.sum()==0:
            continue

        a = d['POS'].loc[dq].values.astype(int)
        enc_q = (encode[0] == 'chr{}'.format(chrom))
        chrom_cre = encode.loc[enc_q]

        for feat in feat_names:
            allarr = np.zeros(a.max()+2*radius, dtype = bool)
            feat_pos = chrom_cre[(chrom_cre[10]==feat)]
            for start,end in zip(feat_pos[1], feat_pos[2]):
                allarr[start:end ]= 1

            sliding = sliding_window_view(allarr,2*radius,axis = 0)
            #1 if the windows of length 2*r starting at pos - r intersect the features
            d[feat].loc[dq] = sliding[a-radius].max(axis = -1).copy() 


    cre_added = t2t_trans.merge(d, on = ['CHR','POS'])
    for feat in feat_names:
        cre_added['not'+feat] = ~cre_added[feat].copy()

    input_df = cre_added
    # pool the association over segments
    pos_df = input_df.groupby(['t2t_chr','t2t_ind','GENE','CELL_TYPE'])[['FDR'] +['not'+feat for feat in feat_names]] .min().reset_index()

    # select positions with associations evaluated only against a single gene
    rs_gene_counts = pos_df[['t2t_chr','t2t_ind','GENE']].drop_duplicates().groupby(['t2t_chr','t2t_ind']).size()
    unique_rs = set(rs_gene_counts[rs_gene_counts<2].index.values)
    unique_rs_pos = pos_df.iloc[[i for i,a in enumerate(zip(pos_df['t2t_chr'],pos_df['t2t_ind'])) if a in unique_rs]]
    max_fdr = unique_rs_pos.groupby(['GENE','CELL_TYPE']).apply(lambda w: (w['FDR']>0.1).sum())

    min_fdr = unique_rs_pos.groupby(['GENE','CELL_TYPE']).apply(lambda w: (w['FDR']<0.1).sum())
    counts = unique_rs_pos.groupby(['GENE','CELL_TYPE']).size()
    # select gene_ct pairs with at least one significant and one not significant qtl
    gene_ct_pairs = set(min_fdr[(min_fdr>0)&(max_fdr>0)].index.values)

    cre_vl = pos_df.iloc[[i for i,a in enumerate(zip(pos_df['GENE'],pos_df['CELL_TYPE']) )if a in gene_ct_pairs]]
    cre_vl = cre_vl[cre_vl['GENE'].isin(expressed_genes)]
    cre_vl = cre_vl[cre_vl['t2t_chr']!='chrY']

    for feat in feat_names:
        cre_vl[feat] = -cre_vl['not'+feat].copy()


    groups = cre_vl.groupby(['GENE','CELL_TYPE']).groups

    assert len(cre_vl)==len(valid_long)

    for feat in feat_names:
        feat_res = {}
        for k,v in groups.items():
            y_true =cre_vl['FDR'].loc[v]< 0.1
            y_score = cre_vl[feat].loc[v]
            feat_res[k] = roc_auc_score(y_true = y_true, y_score = y_score)

        cre_aucs[feat+'_{}'.format(radius)] = pd.Series(feat_res).reset_index().groupby('level_0')[0].mean()

# %% [code]
cre_b = pd.DataFrame(pd.Series({k:v.mean() for k,v in cre_aucs.items()}).sort_values())


# %% [code]
cre_b['cre_type'] = [(a[0]) for a in cre_b.index.str.split('_')]
cre_b['distance'] = [(a[-1]) for a in cre_b.index.str.split('_')]
top_cre_results = pd.DataFrame(cre_b.reset_index().groupby('cre_type').apply(lambda w: w.iloc[w[0].argmax()]))[['index',0]].set_index('index')[0].to_dict()

# %% [code]
baseline_aucs.update(top_cre_results)

# %% [code]
baseline_aucs

# %% [markdown]
## Supervised classifier

# %% [code]
train_data = renamed.loc[:,~renamed.columns.isin(valid_long['GENE'].unique())]

# preparing the gene information
genes = pd.read_table('../data/seq/GENA_2k_precomputed/hs1.110.20220412.ncbiRefSeq.gtf',header = None)

#map CP chroms to chrs
CP_chroms = ['CP068277.2', 'CP068276.2', 'CP068275.2', 'CP068274.2', 'CP068273.2', 'CP068272.2', 'CP068271.2', 'CP068270.2', 'CP068269.2', 'CP068268.2', 'CP068267.2', 'CP068266.2', 'CP068265.2', 'CP068264.2', 'CP068263.2', 'CP068262.2', 'CP068261.2', 'CP068260.2', 'CP068259.2', 'CP068258.2', 'CP068257.2', 'CP068256.2', 'CP068255.2', 'CP086569.2']
chroms = ['chr{}'.format(j) for j in list(range(1,23))+['X','Y']]
chrom_map = {cp:ch for cp,ch in zip(chroms,CP_chroms)}
cp_to_chr = {cp:ch for cp,ch in zip(CP_chroms,chroms)}

#select TSS positions
transcripts = genes[genes[2]=='transcript']
transcripts['id'] =[a.split(';')[0].split(' ')[-1][1:-1] for a in transcripts[8]]
transcripts['chrom'] = transcripts[0].replace(cp_to_chr)
data = transcripts[['chrom',3,4,6,'id']].drop_duplicates()
data = data[data['chrom']!='chrY']
data.columns = ['chrom', 'start', 'stop', 'strand', 'gene_symbol']
data['tss'] = [data['start'].iloc[i] if (s== '+') else data['stop'].iloc[i] for i,s in enumerate(data['strand'])]
data = data.sort_values(['chrom','tss'])


chrs = {chrom:data.loc[val] for chrom,val in data.groupby('chrom').groups.items()}


#build the batch of sequence features
chroms = list(range(1,23)) + ['X']
output_features = []
gene_symbols = []


for chrom_num in tqdm(chroms):
    chrom = 'chr{}'.format(chrom_num)
    chr_dat = np.load('../data/seq/GENA_2k_precomputed/{}_precomputed_features.npy'.format(chrom), mmap_mode = 'r')
    starts = (chrs[chrom]['tss']//1000)
    
    sub = ((starts + 50 )< len(chr_dat) ) & ((starts - 50 )>= 0)
    
    
    
    output_features.append(np.stack([chr_dat[starts.loc[sub]+i] for i in range(-50,50)],axis = 1))
#     output_features.append(np.stack([chr_dat[starts]],axis = 1))
    gene_symbols+= (chrs[chrom]['gene_symbol'].loc[sub].tolist())
output_features = np.concatenate(output_features,axis =0)

gs = pd.Series(gene_symbols)
common = gs[gs.isin(train_data.columns)].reset_index().groupby(0).min()

def znorm(df,axis = 0):
    a = df.values
    return pd.DataFrame((a - a.mean(axis =axis,keepdims = True))/(1e-6+ a.std(axis = axis,keepdims = True)), index = df.index.values, columns = df.columns)


expr =train_data.loc[:,common.index.values].T

Y = znorm(expr[expr.std(axis = 1)>0],axis = 1)


# %% [code]


# %% [code]
size = 10
X = output_features[common['index'].loc[Y.index.values].values].transpose(0,2,1)[:,:,(50-size):(50+size)]

idx = np.arange(X.shape[0])
np.random.seed(123)
np.random.shuffle(idx)

tr = idx[:6000]
val = idx[6000:]
X_tr = torch.from_numpy(X[tr]).float().cuda()
X_val = torch.from_numpy(X[val]).float().cuda()
Y_tr = torch.from_numpy(Y.values[tr]).float().cuda()
Y_val = torch.from_numpy(Y.values[val]).float().cuda()

# model = nn.Sequential(nn.Conv1d(768,24,5), nn.ReLU(), nn.Conv1d(24,8,1)).cuda()
model = nn.Sequential(nn.Conv1d(768,8,1)).cuda()
bias = torch.zeros(1,8, requires_grad = True,device = 'cuda')

bce = nn.BCEWithLogitsLoss()
opt = torch.optim.Adam(list(model.parameters()) + [bias], lr = 0.001)

tr_inds = np.arange(X_tr.shape[0])
lh = []
vlh = []
rh = []


avg = torch.from_numpy(Y.values).mean(axis =0,keepdim = True).cuda()
def get_logits(inp):
    return bias + avg+0.001*model(inp).mean(axis = -1)

def get_loss(x,y):
    return torch.square(x-y).sum(axis = 1).mean()

for epoch in tqdm(range(5000)):
    np.random.shuffle(tr_inds)
    for batch in np.array_split(tr_inds, 10):
        
        x = X_tr[batch]
        y = Y_tr[batch]
        
        reg = torch.abs(model[0].weight).mean() + torch.abs(model[-1].weight).mean()
        logits = get_logits(x)
        loss = get_loss(logits,y)
        opt.zero_grad()
        (loss+1e-4*reg).backward()
        opt.step()
        lh.append(loss.detach().cpu().numpy())
        rh.append(reg.detach().cpu().numpy())
        
    vlh.append(get_loss(get_logits(X_val),Y_val).detach().cpu().numpy())

        
    if (epoch+1)%50==0:
        plt.figure(figsize=(9,3))
        plt.subplot(1,3,1)
        plt.plot(lh)
        plt.ylim(5,6)
        plt.subplot(1,3,2)
        plt.plot(vlh)
        plt.ylim(5.2,5.6)
        plt.subplot(1,3,3)
        plt.plot(rh)
        plt.ylim(0,1e-5)
        plt.show()

    if epoch>20:
        if (np.mean(vlh[-20:-10])-np.mean(vlh[-10:]) <0):
            break


# %% [code]
std = get_loss(Y_val.mean(axis =0,keepdim=True),Y_val).item()

# %% [code]
Y_val

# %% [code]
std

# %% [code]
1- np.mean(vlh[-10:])/std

# %% [code]
1-np.mean(vlh[-10:])/std

# %% [code]
# extract features for all the unique positions in the filtered associations
unique_pos = valid_long[['t2t_chr','t2t_ind']].drop_duplicates()
chrom_inds = unique_pos.groupby('t2t_chr').groups
cis_seq_batches = []
dat_inds =[]
for chrom,inds in chrom_inds.items():
    chm13_pos = unique_pos['t2t_ind'].loc[inds]
    chr_dat = np.load('../data/seq/GENA_2k_precomputed/{}_precomputed_features.npy'.format(chrom), mmap_mode = 'r')
    cis_seq_batches.append(chr_dat[chm13_pos.values]/10)
    dat_inds+= list(inds)
cis_seq_batch = torch.tensor(np.concatenate(cis_seq_batches)).float().to('cuda:{}'.format(DEVICE))

# construct a map assigning chromosome inds to the features
qtl_to_seq_map = {ch_ind:i for i, ch_ind in enumerate (zip(unique_pos['t2t_chr'].loc[dat_inds],unique_pos['t2t_ind'].loc[dat_inds]))}


#construct maps assigning each cell type/gene to its indices in the seq batch and preds
ct_gene_groups = valid_long.groupby(['CELL_TYPE','GENE']).groups
seq_inds = {}
fdrs = {}
for key,inds in ct_gene_groups.items():
    CELLTYPE,GENE = key
    seq_inds[CELLTYPE, GENE] = np.array([qtl_to_seq_map[a] for a in zip(valid_long['t2t_chr'].loc[inds],valid_long['t2t_ind'].loc[inds])])
    fdrs[CELLTYPE,GENE] = valid_long['FDR'].loc[inds]
    
classifier_preds = pd.DataFrame((cis_seq_batch.cuda()@model[0].weight[:,:,0].T).detach().cpu().numpy(), columns = Y.columns)
vl = valid_long.copy()
groups = vl.groupby(['GENE','CELL_TYPE']).groups
cpres = {}
for k,v in groups.items():
    y_true =(vl['FDR'].loc[v]< 0.1).values
    y_score = classifier_preds[k[1]].loc[[qtl_to_seq_map[a,b] for a,b in zip(vl['t2t_chr'].loc[v], vl['t2t_ind'].loc[v])]].values

    cpres[k] = roc_auc_score(y_true = y_true, y_score = -y_score)
    
cpaucs = pd.Series(cpres).reset_index().groupby('level_0')[0].mean()


# %% [code]
baseline_aucs['expression_classifier'] = cpaucs.mean()

# %% [code]
baseline_aucs

# %% [markdown]
### eQTL classifier

# %% [code]
from sklearn.linear_model import LogisticRegressionCV,LogisticRegression

# %% [code]
genes = valid_long['GENE'].unique().copy()
np.random.seed(1234)
out = {}
for gene in tqdm(genes):
    test_genes = [gene]    


    tvl = valid_long.loc[~valid_long['GENE'].isin(test_genes)].copy()
    test = valid_long.loc[valid_long['GENE'].isin(test_genes)].copy()


    lr = LogisticRegression(max_iter=10000)
    train = tvl.groupby(['t2t_chr','t2t_ind'])['FDR'].min().reset_index()
    xtr = cis_seq_batch[[qtl_to_seq_map[a,b] for a,b in zip(train['t2t_chr'], train['t2t_ind'])]].cpu().numpy()
    ytr = torch.from_numpy((train['FDR']<0.1).astype(float).values)
    lr.fit(xtr,ytr)

    vl = test.copy()
    groups = vl.groupby(['GENE','CELL_TYPE']).groups
    cres = {}
    preds = lr.predict_proba(cis_seq_batch.detach().cpu().numpy())[:,1]
    for k,v in groups.items():
        y_true =(vl['FDR'].loc[v]< 0.1).values
        y_score = preds[[qtl_to_seq_map[a,b] for a,b in zip(vl['t2t_chr'].loc[v], vl['t2t_ind'].loc[v])]]

        cres[k] = roc_auc_score(y_true = y_true, y_score = -y_score)

    caucs = pd.Series(cres).reset_index().groupby('level_0')[0].mean()

    out[gene]= np.mean(caucs)

# %% [code]
pd.Series(out).mean()

# %% [code]
baseline_dat

# %% [markdown]
# Permutation tests

# %% [code]
unique_pos = valid_long[['t2t_chr','t2t_ind']].drop_duplicates()
chrom_inds = unique_pos.groupby('t2t_chr').groups
cis_seq_batches = []
dat_inds =[]
for chrom,inds in chrom_inds.items():
    chm13_pos = unique_pos['t2t_ind'].loc[inds]
    chr_dat = np.load('../data/seq/GENA_2k_precomputed/{}_precomputed_features.npy'.format(chrom), mmap_mode = 'r')
    cis_seq_batches.append(chr_dat[chm13_pos.values]/50)
    dat_inds+= list(inds)
cis_seq_batch = torch.tensor(np.concatenate(cis_seq_batches)).float().to('cuda:{}'.format(DEVICE))

# construct a map assigning chromosome inds to the features
qtl_to_seq_map = {ch_ind:i for i, ch_ind in enumerate (zip(unique_pos['t2t_chr'].loc[dat_inds],unique_pos['t2t_ind'].loc[dat_inds]))}


#construct maps assigning each cell type/gene to its indices in the seq batch and preds
ct_gene_groups = valid_long.groupby(['CELL_TYPE','GENE']).groups
seq_inds = {}
y_trues = {}
for key,inds in ct_gene_groups.items():
    CELLTYPE,GENE = key
    seq_inds[CELLTYPE, GENE] = np.array([qtl_to_seq_map[a] for a in zip(valid_long['t2t_chr'].loc[inds],valid_long['t2t_ind'].loc[inds])])
    y_trues[CELLTYPE,GENE] = (valid_long['FDR'].loc[inds]<0.1)
    
    


# %% [code]
ct_set_to_g = {}
for g, ct_set in {g: tuple(sorted(renamed[g][renamed[g]>2e-5].index.values)) for g in orig_genes}.items():
    ct_set_to_g.setdefault(ct_set, []).append(g)

orig_group_genes = [np.array(gs) for _,gs in ct_set_to_g.items()]

# %% [code]
plt.figure(figsize = (3,2))
plt.bar(np.arange(len(orig_group_genes)), pd.Series([len(v) for v in orig_group_genes]).sort_values())
plt.xticks(np.arange(len(orig_group_genes)),rotation =90);
plt.title('size of gene groups')

# %% [code]


# %% [code]


# %% [code]
# auxiliary function to prepare the original and edited batch
def prepare_sc_batch(ct, gene_to_add, amount = 0.0001):
    ct_mean = renamed.loc[ct]
    edited_mean = ct_mean.copy()
    edited_mean[gene_to_add]+=amount
    return torch.tensor(np.stack([ct_mean.values.copy(), edited_mean.values]),device = 'cuda:{}'.format(DEVICE),dtype = torch.float)
    
    
def permutation_test(path,perm_gene = False,perm_ct= False,perm_rs = False,group_perm_gene = False,group_perm_rs = False,rand_init = False,group_perm_ct = False,
                     nperm =1000,thresh = -1,n_control = 40,init_one_half = False):
    softmax = True
    bn = False
    n_learn = 1
    n_control = n_control
    dropout= 0
    softmax = True
    bn = False
    multi_learner = nn.ModuleList([LatentPSALearner(sc = torch.from_numpy(renamed.values),dropout=dropout, softmax=softmax, genome = cis_seq_batch, n_control = n_control, bias = False,batch_norm = bn) for _ in range(n_learn)]).to('cuda:{}'.format(DEVICE))

    checkpoint = torch.load(path)
    sd = checkpoint['learner_state_dict']
    multi_learner.load_state_dict(sd)
    multi_learner.eval();
    learner = multi_learner[0]
    learner.eval();
    
    assert (group_perm_gene&perm_gene)==False
    assert (group_perm_rs&perm_rs)==False
    assert (group_perm_ct&perm_ct)==False
    assert (rand_init&init_one_half)==False
    
    
    ct_lists = [['Classic Monocyte','Non-classic Monocyte' ], ['Naïve/Immature B Cell' ,'Memory B Cell', 'Plasma Cell'] ,['CD4 Effector memory/TEMRA','Natural Killer Cell','CD8 Effector memory']]
    group_ct_shufs = []
    for l in list(itertools.product(*[list(itertools.permutations(c)) for c in ct_lists])):
        group_ct_shufs.append({i:j for i,j in zip(np.concatenate(ct_lists), np.concatenate([[k for k in ll] for ll in l]))})

    
    # initialize maps that will be shuffled during permutations
    orig_genes = valid_long['GENE'].unique()
    shuf_genes = orig_genes.copy()
    orig_cts = renamed.index.values.copy()
    shuf_cts = orig_cts.copy()
    shuf_idx = np.arange(len(cis_seq_batch))

    # want to enable shuffling only genes that are expressed in the same set of cell types, hence this shuffle
    ct_set_to_g = {}
    for g, ct_set in {g: tuple(sorted(renamed[g][renamed[g]>2e-5].index.values)) for g in orig_genes}.items():
        ct_set_to_g.setdefault(ct_set, []).append(g)

    orig_group_genes = [np.array(gs) for _,gs in ct_set_to_g.items()]
    shuf_group_genes = [arr.copy() for arr in orig_group_genes]

    #want to enable shuffling to get sense for resolution
    if group_perm_rs:
        concom = ccs[thresh]
        orig_group_rs = [np.where(concom[1]==k)[0] for k in range(concom[0])]
        assert (np.concatenate(orig_group_rs)-np.arange(len(concom[1]))).max()==0    
        shuf_group_rs = [arr.copy() for arr in orig_group_rs]


    assert (group_perm_gene&perm_gene)==False
    assert (group_perm_rs&perm_rs)==False
    assert (group_perm_ct&perm_ct)==False
    
    if group_perm_ct:
        nperm = len(group_ct_shufs)-1
    

    perm_stats = {}
    PRINT_EVERY = min(nperm,10)
    np.random.seed(45)

    for perm in tqdm(range(1+nperm)):
        # first permutation is always test statistic (i.e., no shuffle)
        if perm>0:
            if group_perm_gene:
                for arr in shuf_group_genes:
                    np.random.shuffle(arr) #only shuffle within groups
            if group_perm_rs:
                for arr in shuf_group_rs:
                    np.random.shuffle(arr) #only shuffle within groups
            if perm_gene:
                np.random.shuffle(shuf_genes)

            if perm_ct:
                np.random.shuffle(shuf_cts)
                


            if perm_rs:
                np.random.shuffle(shuf_idx)

            if rand_init:
                multi_learner = nn.ModuleList([LatentPSALearner(sc = torch.from_numpy(renamed.values),dropout=dropout, softmax=softmax, genome = cis_seq_batch, n_control = n_control, bias = False,batch_norm = bn) for _ in range(n_learn)]).to('cuda:{}'.format(DEVICE))
                multi_learner.eval();
                learner = multi_learner[0]
                learner.eval();
                
            if init_one_half:
                multi_learner = nn.ModuleList([LatentPSALearner(sc = torch.from_numpy(renamed.values),dropout=dropout, softmax=softmax, genome = cis_seq_batch, n_control = n_control, bias = False,batch_norm = bn) for _ in range(n_learn)]).to('cuda:{}'.format(DEVICE))
                multi_learner.eval();
                ll = multi_learner[0]
                ll.eval();
                learner.seq_k = ll.seq_k
                


        # if perm==0, shuf_gene_map, shuf_idx, shuf_ct_map are all identity
        if group_perm_gene:
            shuf_gene_map = {i:j for i,j in zip(np.concatenate(orig_group_genes), np.concatenate(shuf_group_genes))}
        else:
            shuf_gene_map = {i:j for i,j in zip(orig_genes,shuf_genes)}


        if group_perm_rs:
            shuf_idx = np.concatenate(shuf_group_rs)
            
        shuf_ct_map = {i:j for i,j in zip(orig_cts,shuf_cts)}
        
        if group_perm_ct:
            shuf_ct_map = group_ct_shufs[perm]


        # compute AUCs for the discrepancy
        for CELLTYPE, GENE in ct_gene_groups.keys():
            sub_seq = cis_seq_batch[shuf_idx][seq_inds[CELLTYPE,GENE]]
            

            shuf_ct = shuf_ct_map[CELLTYPE]
            shuf_gene = shuf_gene_map[GENE]
            sc_batch = prepare_sc_batch(shuf_ct, shuf_gene)

            Txs =  learner.T_xs(seq_batch= sub_seq, sc_batch = sc_batch)
            y_true = (y_trues[CELLTYPE,GENE]).astype(int).values
            y_pred = torch.abs(Txs[1]-Txs[0]).detach().cpu().numpy()
            perm_stats[perm,CELLTYPE,GENE] = roc_auc_score(y_true= y_true, y_score = y_pred)

    res = pd.Series(perm_stats).reset_index()
    res.columns = ['perm', 'ct', 'gene', 'auc']
    res['perm_gene'] = perm_gene
    res['group_perm_gene'] = group_perm_gene
    res['perm_rs'] = perm_rs
    res['perm_ct'] = perm_ct
    res['group_perm_rs'] = group_perm_rs
    res['rand_init'] = rand_init
    res['thresh'] = thresh
    res['group_perm_ct'] = group_perm_ct
    return res


# %% [code]


# %% [code]


# %% [code]


# %% [code]
from scipy.sparse.csgraph import connected_components

# %% [code]
# SLOW way to compute grouped pairwise distances...
pw_distances = {}
ccs = {}
for chrom1,pos1,ind1 in tqdm(zip(unique_pos['t2t_chr'].loc[dat_inds],unique_pos['t2t_ind'].loc[dat_inds],np.arange(len(dat_inds))), total = len(dat_inds)):
    for chrom2,pos2,ind2  in zip(unique_pos['t2t_chr'].loc[dat_inds],unique_pos['t2t_ind'].loc[dat_inds],np.arange(len(dat_inds))):
        if chrom1!=chrom2:
            continue
        pw_distances[ind1,ind2] = np.abs(pos1-pos2)
distances = pd.Series(pw_distances).unstack().fillna(1e6)

# connected components to define which RS can be permuted
from scipy.sparse.csgraph import connected_components
for threshold in [5,6,7,8,9,10,50,100,1000]:
    if threshold in ccs:
        continue
    ccs[threshold] = connected_components(1*(distances.values<threshold))


# %% [code]
path  = 'results/apr5/metacell_fixed_seq_targ/repeat_0_epoch_76.pt'
NPERM = 2000
results = []
    
perm_gene = False
perm_ct= False
perm_rs = False
group_perm_gene = False
rand_init = True ###
group_perm_rs = False
group_perm_ct = False
thresh = -1
results.append(permutation_test(path,perm_gene = perm_gene,perm_ct= perm_ct,perm_rs = perm_rs,group_perm_gene = group_perm_gene,rand_init = rand_init,
                 group_perm_rs = group_perm_rs,thresh = thresh,
                 nperm =NPERM))

perm_gene = False
perm_ct= False
perm_rs = False
group_perm_gene = True ###
rand_init = False
group_perm_rs = False
group_perm_ct = False
thresh = -1
results.append(permutation_test(path,perm_gene = perm_gene,perm_ct= perm_ct,perm_rs = perm_rs,group_perm_gene = group_perm_gene,rand_init = rand_init,
                 group_perm_rs = group_perm_rs,thresh = thresh,
                 nperm =NPERM))


perm_gene = False
perm_ct= False
perm_rs = True ###
group_perm_gene = False
rand_init = False
group_perm_ct = False
group_perm_rs = False
thresh = -1
results.append(permutation_test(path,perm_gene = perm_gene,perm_ct= perm_ct,perm_rs = perm_rs,group_perm_gene = group_perm_gene,rand_init = rand_init,
                 group_perm_rs = group_perm_rs,thresh = thresh,
                 nperm =NPERM))


perm_gene = False
perm_ct= True
perm_rs = False
group_perm_gene = False
rand_init = False
group_perm_rs = False
group_perm_ct = False
thresh = -1
results.append(permutation_test(path,perm_gene = perm_gene,perm_ct= perm_ct,perm_rs = perm_rs,group_perm_gene = group_perm_gene,rand_init = rand_init,
                 group_perm_rs = group_perm_rs,thresh = thresh,
                 nperm =NPERM))


perm_gene = True
group_perm_gene = False
perm_rs = False
group_perm_rs = False
rand_init = False
perm_ct= False
group_perm_ct = False
thresh = -1
results.append(permutation_test(path,perm_gene = perm_gene,perm_ct= perm_ct,perm_rs = perm_rs,group_perm_gene = group_perm_gene,rand_init = rand_init,
                 group_perm_rs = group_perm_rs,thresh = thresh,
                 nperm =NPERM))


perm_gene = False
perm_ct= False
perm_rs = False
group_perm_gene = False
rand_init = False
group_perm_rs = True ####
group_perm_ct = False
for thresh in list(ccs.keys()):
    results.append(permutation_test(path,perm_gene = perm_gene,perm_ct= perm_ct,perm_rs = perm_rs,group_perm_gene = group_perm_gene,rand_init = rand_init,
                     group_perm_rs = group_perm_rs,thresh = thresh,
                     nperm =NPERM))
    
    
    
perm_gene = False
perm_ct= False
perm_rs = False
group_perm_gene = False
rand_init = False###
group_perm_rs = False
group_perm_ct = True
thresh = -1
results.append(permutation_test(path,perm_gene = perm_gene,perm_ct= perm_ct,perm_rs = perm_rs,group_perm_gene = group_perm_gene,rand_init = rand_init,
                 group_perm_rs = group_perm_rs,thresh = thresh,group_perm_ct = group_perm_ct,
                 nperm =NPERM))


combined = pd.concat(results, axis = 0,ignore_index = True)

combined.to_pickle('../data/downstream/Nature_cis_perm_results_583model')


# %% [markdown]
### checking one init

# %% [code]
check_72[check_72['perm']==0].groupby('gene')['auc'].mean().mean()

# %% [code]
path  = 'results/apr5/metacell_fixed_seq_targ/repeat_0_epoch_76.pt'
NPERM = 200

    
perm_gene = False
perm_ct= False
perm_rs = False
group_perm_gene = False
rand_init = False ###
group_perm_rs = False
group_perm_ct = False
thresh = -1
one_res = permutation_test(path,perm_gene = perm_gene,perm_ct= perm_ct,perm_rs = perm_rs,group_perm_gene = group_perm_gene,rand_init = rand_init,
                 group_perm_rs = group_perm_rs,thresh = thresh,
                 nperm =NPERM,init_one_half = True)


# %% [code]
aa = one_res.groupby(['perm','gene'])['auc'].mean().reset_index().groupby('perm').mean()
(aa.iloc[1:]>aa.iloc[0]).mean()

# %% [code]
combined.groupby(['perm','gene'])

# %% [code]
combined = pd.read_pickle('../data/downstream/Nature_cis_perm_results_583model')
a = combined.groupby(['perm','perm_gene','perm_ct','group_perm_ct','group_perm_gene','group_perm_rs', 'perm_rs','rand_init','thresh','gene'])['auc'].mean().reset_index()


perm_conds = a.groupby(['perm_gene','perm_ct','group_perm_ct','group_perm_gene','perm_rs','group_perm_rs', 'rand_init','thresh']).groups
pvalues = {}
avgs = {}
for cond, inds in perm_conds.items():
    cond_perms = a.loc[inds]
    cond_avg = cond_perms.groupby('perm')['auc'].mean()
    keys = ['perm_gene','perm_ct','group_perm_ct','group_perm_gene','perm_rs','group_perm_rs', 'rand_init','thresh']
    if cond[-1]==-1:
        key = [a for a,b in zip(keys, cond) if b==True][0]
    else:
        key = 'group_perm_rs_{}'.format(cond[-1])    
    pvalues[key] = 1- (cond_avg[0]>cond_avg[1:]).mean()
    avgs[key] = a.loc[inds]
    avgs[key]['perm_key'] = key
    
res = pd.concat(avgs,ignore_index = True).groupby(['perm_key','perm'])['auc'].mean().reset_index()
test_stat = res['auc'].iloc[0]


# %% [code]
test_stat

# %% [code]


display(pd.Series(pvalues))


order = ['perm_rs','rand_init', 'perm_gene','group_perm_gene','perm_ct','group_perm_ct']

plt.figure(figsize = (6,5))
dat = sns.stripplot(data = res[(res['perm_key'].isin(order) )& (res['perm']!=0)], x = 'perm_key', y = 'auc',
                   s = 1, color = 'grey',order = order)
plt.xticks(rotation = 90)
plt.hlines(test_stat, -0.2, len(order)-1 + 0.2,colors = 'red',zorder  = 5)

mean_res = res[res['perm']!=0].groupby('perm_key').mean()['auc'].loc[order]

plt.hlines(mean_res, np.arange(len(order))-0.2, np.arange(len(order))+0.2, colors = 'black',zorder = 5)

plt.title([v for o,v in pd.Series(pvalues).loc[order].to_dict().items()], fontsize = 6)

# %% [code]


# %% [code]


# %% [markdown]
### Evaluating resolution

# %% [code]
resolution = res[(res['perm']!=0) & (res['perm_key'].str.contains('group_perm_rs'))].copy()
resolution['res'] = [int(pk.split('_')[-1]) for pk in resolution['perm_key']]
resolution = resolution[~resolution['res'].isin([5,6])]
display(1-resolution.groupby('res')['auc'].apply(lambda w: (w< test_stat).mean()))
plt.figure(figsize = (5,5))
pcs = resolution.groupby('res').apply(lambda w: np.percentile(w['auc'],95))
medians = resolution.groupby('res').apply(lambda w: np.percentile(w['auc'],50))
sns.stripplot(data = resolution, y = 'auc', x = 'res', c = 'grey', s = 1)
plt.xticks(rotation = 90)
plt.hlines(test_stat, -0.5, len(pcs)-0.5,colors = 'red',zorder = 5)


plt.hlines(pcs, np.arange(len(pcs))-0.2,np.arange(len(pcs))+0.2, colors = 'black',linestyle = 'dotted',zorder = 25)
plt.hlines(medians, np.arange(len(pcs))-0.2,np.arange(len(pcs))+0.2, colors = 'black',zorder = 25)

# %% [code]


# %% [code]


# %% [markdown]
### Plotting baselines

# %% [code]
baseline_dat = {'tss':tss_aucs,'gc':gc_aucs, 'expr_class':cpaucs,'eqtl_class': out}
baseline_dat.update(cre_aucs)#{k:cre_aucs[k] for k in top_cre_results})
baseline_dat['psa'] = avgs['perm_rs'][avgs['perm_rs']['perm']==0].set_index('gene')['auc']

plt.figure(figsize = (20,5))
baseline_df = pd.DataFrame(baseline_dat)
baseline_means = baseline_df.mean(axis =0)
order = baseline_means.sort_values().index.values
sns.stripplot(data = baseline_df.melt(), x = 'variable',y  = 'value',order = order, c = 'navy')
plt.xticks(rotation = 90);
psa_pos = np.sum(baseline_means<baseline_means['psa'])
plt.hlines(baseline_dat['psa'].mean(), psa_pos-0.3, psa_pos+0.3, colors = 'red',zorder = 25)
plt.hlines(baseline_df.mean(axis =0).loc[order], np.arange(len(baseline_means))-0.3, np.arange(len(baseline_means))+0.3 ,colors = 'black',zorder = 2)

# %% [code]
np.where(order=='psa')

# %% [code]
baseline_dat = {'tss':tss_aucs,'gc':gc_aucs, 'expr_class':cpaucs,'eqtl_class': out}
baseline_dat.update({k:cre_aucs[k] for k in top_cre_results})
baseline_dat['psa'] = avgs['perm_rs'][avgs['perm_rs']['perm']==0].set_index('gene')['auc']

plt.figure(figsize = (5,5))
baseline_df = pd.DataFrame(baseline_dat)
baseline_means = np.abs(0.5-baseline_df.mean(axis =0))
order = np.array(['tss','gc','CTCF-only_1000', 'PLS_1000', 'pELS_7000',
       'DNase-H3K4me3_6000','dELS_3000',  'eqtl_class','expr_class','psa'])
sns.stripplot(data = baseline_df.melt(), x = 'variable',y  = 'value',order = order, s = 2.5, c = 'navy')
plt.xticks(rotation = 90);
plt.hlines(baseline_df.mean(axis =0).loc[order], np.arange(len(baseline_means))-0.3, np.arange(len(baseline_means))+0.3 ,colors = 'black',zorder = 20)
plt.hlines(baseline_df.mean(axis =0).loc['psa'],  np.where(order=='psa')[0]-0.3, np.where(order=='psa')[0]+0.3 ,colors = 'red',zorder = 20)

# %% [code]
from scipy.stats import ttest_ind,ttest_1samp

# %% [code]
1/2000

# %% [code]
pd.Series({col: ttest_1samp(baseline_df[col].dropna(), 0.5)[1] for col in baseline_df.columns}).sort_values()


# %% [markdown]
### Random seed


# %% [code]
NPERM = 200
path  = 'results/apr10/pbmc_583_redo_110_detach/repeat_0_epoch_76.pt'
results = []


perm_gene = False
perm_ct= False
perm_rs = False
group_perm_gene = True ###
rand_init = False
group_perm_rs = False
thresh = -1
results.append(permutation_test(path,perm_gene = perm_gene,perm_ct= perm_ct,perm_rs = perm_rs,group_perm_gene = group_perm_gene,rand_init = rand_init,
                 group_perm_rs = group_perm_rs,thresh = thresh,
                 nperm =NPERM))

perm_gene = False
perm_ct= True
perm_rs = False
group_perm_gene = False
rand_init = False
group_perm_rs = False
thresh = -1
results.append(permutation_test(path,perm_gene = perm_gene,perm_ct= perm_ct,perm_rs = perm_rs,group_perm_gene = group_perm_gene,rand_init = rand_init,
                 group_perm_rs = group_perm_rs,thresh = thresh,
                 nperm =NPERM))


perm_gene = True
group_perm_gene = False
perm_rs = False
group_perm_rs = False
rand_init = False
perm_ct= False
thresh = -1
results.append(permutation_test(path,perm_gene = perm_gene,perm_ct= perm_ct,perm_rs = perm_rs,group_perm_gene = group_perm_gene,rand_init = rand_init,
                 group_perm_rs = group_perm_rs,thresh = thresh,
                 nperm =NPERM))
perm_gene = False
group_perm_gene = False
perm_rs = True
group_perm_rs = False
rand_init = False
perm_ct= False
thresh = -1
results.append(permutation_test(path,perm_gene = perm_gene,perm_ct= perm_ct,perm_rs = perm_rs,group_perm_gene = group_perm_gene,rand_init = rand_init,
                 group_perm_rs = group_perm_rs,thresh = thresh,
                 nperm =NPERM))


perm_gene = False
group_perm_gene = False
perm_rs = True
group_perm_rs = False
rand_init = False
perm_ct= False
group_perm_ct= True
thresh = -1
results.append(permutation_test(path,perm_gene = perm_gene,perm_ct= perm_ct,perm_rs = perm_rs,group_perm_gene = group_perm_gene,rand_init = rand_init,
                 group_perm_rs = group_perm_rs,thresh = thresh,
                 nperm =NPERM))


seed_combined = pd.concat(results, axis = 0,ignore_index = True)

seed_combined.to_pickle('../data/downstream/Nature_cis_perm_results_random_seed')


# %% [code]
combined = pd.read_pickle('../data/downstream/Nature_cis_perm_results_random_seed')
a = combined.groupby(['perm','perm_gene','perm_ct','group_perm_gene','group_perm_rs', 'perm_rs','rand_init','thresh','gene'])['auc'].mean().reset_index()


perm_conds = a.groupby(['perm_gene','perm_ct','group_perm_gene','perm_rs','group_perm_rs', 'rand_init','thresh']).groups
pvalues = {}
avgs = {}
for cond, inds in perm_conds.items():
    cond_perms = a.loc[inds]
    cond_avg = cond_perms.groupby('perm')['auc'].mean()
    keys = ['perm_gene','perm_ct','group_perm_gene','perm_rs','group_perm_rs', 'rand_init','thresh']
    if cond[-1]==-1:
        key = [a for a,b in zip(keys, cond) if b==True][0]
    else:
        key = 'group_perm_rs_{}'.format(cond[-1])    
    pvalues[key] = 1- (cond_avg[0]>cond_avg[1:]).mean()
    avgs[key] = a.loc[inds]
    avgs[key]['perm_key'] = key
    
res = pd.concat(avgs,ignore_index = True).groupby(['perm_key','perm'])['auc'].mean().reset_index()
test_stat = res['auc'].iloc[0]


display(pd.Series(pvalues))


order = ['perm_rs','perm_ct','perm_gene','group_perm_gene']

plt.figure(figsize = (6,5))
dat = sns.stripplot(data = res[(res['perm_key'].isin(order) )& (res['perm']!=0)], x = 'perm_key', y = 'auc',
                   s = 1.2, color = 'grey',order = order)
plt.xticks(rotation = 90)
plt.hlines(test_stat, -0.5, 4.5,colors = 'red')

mean_res = res[res['perm']!=0].groupby('perm_key').mean()['auc'].loc[order]

plt.hlines(mean_res, np.arange(len(order))-0.2, np.arange(len(order))+0.2, colors = 'black',zorder = 5)

# %% [markdown]
### Biological replicates
- Located in other notebook

# %% [markdown]
## Ablations

# %% [code]
np.array([r.groupby('gene')['auc'].mean().mean() for r in results]).max()

# %% [code]


NPERM = 0
results = []
for i in range(4, 104, 4):
    path  = 'results/apr20/metacell_no_sc/repeat_0_epoch_{}.pt'.format(i)
    n_control = 40


    perm_gene = False
    perm_ct= False
    perm_rs = False
    group_perm_gene = True ###
    rand_init = False
    group_perm_rs = False
    thresh = -1
    results.append(permutation_test(path,perm_gene = perm_gene,perm_ct= perm_ct,perm_rs = perm_rs,group_perm_gene = group_perm_gene,rand_init = rand_init,
                     group_perm_rs = group_perm_rs,thresh = thresh,
                     nperm =NPERM))

# %% [code]


NPERM = 0
path  = 'results/apr5/metacell_fixed_seq_targ/repeat_0_epoch_76.pt'
n_control = 40


perm_gene = False
perm_ct= False
perm_rs = False
group_perm_gene = True ###
rand_init = False
group_perm_rs = False
thresh = -1
r = permutation_test(path,perm_gene = perm_gene,perm_ct= perm_ct,perm_rs = perm_rs,group_perm_gene = group_perm_gene,rand_init = rand_init,
                     group_perm_rs = group_perm_rs,thresh = thresh,
                     nperm =NPERM)

# %% [code]
r.groupby('gene')['auc'].mean().mean()

# %% [code]


NPERM = 0
results = []
for i in range(4, 104, 4):
    path  = 'results/apr5/metacell_nosyn_sparsity/repeat_0_epoch_{}.pt'.format(i)
    n_control = 40


    perm_gene = False
    perm_ct= False
    perm_rs = False
    group_perm_gene = True ###
    rand_init = False
    group_perm_rs = False
    thresh = -1
    results.append(permutation_test(path,perm_gene = perm_gene,perm_ct= perm_ct,perm_rs = perm_rs,group_perm_gene = group_perm_gene,rand_init = rand_init,
                     group_perm_rs = group_perm_rs,thresh = thresh,
                     nperm =NPERM))

# %% [code]
4*120*(np.argmax([r.groupby('gene')['auc'].mean().mean() for r in results])+1)

# %% [code]
r.groupby('gene')['auc'].mean().mean()

# %% [markdown]
## Corruption results

# %% [markdown]
### One-by-one corrupted

# %% [code]
import os

# %% [code]
GENES = [a.split('_')[-1] for a in os.listdir('results/sep29/corruption/')]

# %% [code]
# load the learner
single_stats = {}
nperm = 0
for gene in GENES:
    path = 'results/sep29/corruption/single_{}/repeat_0_epoch_76.pt'.format(gene)
    
    softmax = True
    bn = False
    n_learn = 1
    n_control = 40
    dropout= 0
    softmax = True
    bn = False
    multi_learner = nn.ModuleList([LatentPSALearner(sc = torch.from_numpy(renamed.values),dropout=dropout, softmax=softmax, genome = cis_seq_batch, n_control = n_control, bias = False,batch_norm = bn) for _ in range(n_learn)]).to('cuda:{}'.format(DEVICE))

    checkpoint = torch.load(path)
    sd = checkpoint['learner_state_dict']
    multi_learner.load_state_dict(sd)
    multi_learner.eval();
    learner = multi_learner[0]
    learner.eval();


    # initialize maps that will be shuffled during permutations
    orig_genes = valid_long['GENE'].unique()
    shuf_genes = orig_genes.copy()
    orig_cts = renamed.index.values.copy()
    shuf_cts = orig_cts.copy()
    shuf_idx = np.arange(len(cis_seq_batch))

    # want to enable shuffling only genes that are expressed in the same set of cell types, hence this shuffle
    ct_set_to_g = {}
    for g, ct_set in {g: tuple(sorted(renamed[g][renamed[g]>2e-5].index.values)) for g in orig_genes}.items():
        ct_set_to_g.setdefault(ct_set, []).append(g)

    orig_group_genes = [np.array(gs) for _,gs in ct_set_to_g.items()]
    shuf_group_genes = [arr.copy() for arr in orig_group_genes]

    # select which kind of permutation
    perm_gene = False
    perm_ct= False
    perm_rs = False
    group_perm_gene = False
    group_perm_rs = False

    assert (group_perm_gene&perm_gene)==False
    assert (group_perm_rs&perm_rs)==False


    nperm = 0
    np.random.seed(45)

    for perm in tqdm(range(1+nperm)):
        # first permutation is always test statistic (i.e., no shuffle)
        if perm>0:
            if group_perm_gene:
                for arr in shuf_group_genes:
                    np.random.shuffle(arr) #only shuffle within groups
            if group_perm_rs:
                for arr in shuf_group_rs:
                    np.random.shuffle(arr) #only shuffle within groups
            if perm_gene:
                np.random.shuffle(shuf_genes)

            if perm_ct:
                np.random.shuffle(orig_cts)

            if perm_rs:
                np.random.shuffle(shuf_idx)

        # if perm==0, shuf_gene_map, shuf_idx, shuf_ct_map are all identity
        if group_perm_gene:
            shuf_gene_map = {i:j for i,j in zip(np.concatenate(orig_group_genes), np.concatenate(shuf_group_genes))}
        else:
            shuf_gene_map = {i:j for i,j in zip(orig_genes,shuf_genes)}


        if group_perm_rs:
            shuf_idx = np.concatenate(shuf_group_rs)
        shuf_ct_map = {i:j for i,j in zip(orig_cts,shuf_cts)}


        # compute AUCs for the discrepancy
        for CELLTYPE, GENE in ct_gene_groups.keys():
            sub_seq = cis_seq_batch[shuf_idx][seq_inds[CELLTYPE,GENE]]
            y_true = (fdrs[CELLTYPE,GENE]<0.1).astype(int).values

            shuf_ct = shuf_ct_map[CELLTYPE]
            shuf_gene = shuf_gene_map[GENE]
            sc_batch = prepare_sc_batch(shuf_ct, shuf_gene,amount = 1e-4)

            Txs =  learner.T_xs(seq_batch= sub_seq, sc_batch = sc_batch)
            y_pred = torch.abs(Txs[1]-Txs[0]).detach().cpu().numpy()
            single_stats[gene,CELLTYPE,GENE] = roc_auc_score(y_true= y_true, y_score = y_pred)


# %% [code]
psa_aucs = baseline_dat['psa'].copy()

# %% [code]
single_res = pd.Series(single_stats).reset_index()
single_res.columns = ['corrupted_gene', 'ct', 'eval_gene', 'auc']

# %% [code]
orig_aucs = psa_aucs.sort_values()
order = orig_aucs.index.values

# %% [code]
df = single_res.groupby(['corrupted_gene','eval_gene'])['auc'].mean().reset_index()

df['orig_auc'] = psa_aucs.loc[df['eval_gene'].values].values
df['diff'] = df['auc'] - df['orig_auc']

order = orig_aucs[orig_aucs>0.50].index.values
diff_df = df.set_index(['corrupted_gene', 'eval_gene'])['auc'].unstack().loc[order,:].loc[:,order]

sns.heatmap(znorm(diff_df,axis = 0).T,cmap = 'bwr',vmin = -5, vmax = 5)
plt.xticks([])
plt.yticks([])

# %% [code]
mat = diff_df.values.copy()

orig = np.arange(mat.shape[0])
perm = np.arange(mat.shape[0])
nperm = 100000
stats = np.zeros(1+nperm)
for i in tqdm(range(1+nperm)):
    stats[i] = np.median(mat[(orig,perm)])
    np.random.shuffle(perm)
    
(stats[0]<stats[1:]).mean()

# %% [code]
plt.figure(figsize = (5,5))
sns.scatterplot(df[(df['orig_auc']>0.5)&(df['eval_gene']==df['corrupted_gene'])],
                y = 'orig_auc', x = 'auc')
plt.ylim(0,1)

plt.xlim(0,1)
plt.plot([0,1], [0,1], c = 'black',linewidth = 0.4)
plt.xlabel('auc with corrupted input')
plt.ylabel('auc with original input')

plt.hlines(0.5,0,1, colors = 'black', linewidth = 0.4,linestyle = 'dashed')
plt.vlines(0.5,0,1, colors = 'black', linewidth = 0.4,linestyle = 'dashed')

# %% [code]
from scipy.stats import spearmanr

# %% [code]
I = df[df['corrupted_gene']==df['eval_gene']].set_index('corrupted_gene').loc[GENES]['diff'].values
J = 1-geneps.loc[GENES].values
spearmanr(I,J)

# %% [code]
plt.figure(figsize = (5,5))
plt.scatter(J,I,c = 'navy')
plt.xlabel('gene permutation p value')
plt.ylabel('z-score on diagonal')
```
