## Supplementary material for "Deriving genetic codes for molecular phenotypes from first principles": Code: Fig4_PBMC_replicates.ipynb

```
# %% [code]
import pandas as pd
import torch
import numpy as np
from tqdm import notebook
from torch import nn
from models.PSALearners import LatentPSALearner
from sklearn.metrics import roc_auc_score
from tqdm.notebook import tqdm
import seaborn as sns
from  matplotlib import pyplot as plt
import pickle
from scipy.stats import ttest_1samp
DEVICE =3

# %% [code]
valid_long = pd.read_pickle('../data/downstream/pbmc_valid_long')

# %% [code]
info = pd.read_pickle('../data/sc/processed/pbmc2_SEACELL_means')

# %% [code]
assn = pd.read_pickle('../data/sc/processed/pbmc2_SEACell_assignments')

# %% [code]
cell_type_map = {'Classic Monocyte':  'CD14 Mono',
'Non-classic Monocyte': 'CD16 Mono',
'Naïve/Immature B Cell':'B naive',
'Memory B Cell':'B memory',
'Plasma Cell':'Plasmablast',
'CD8 Effector memory':'CD8 TEM',
'CD4 Effector memory/TEMRA':'CD4 TEM',
'Natural Killer Cell':'NK'}
sat_to_eq = {v:k for k,v in cell_type_map.items()}


# %% [code]
allmeans = info.loc[assn[14]].groupby(assn[9].values).mean()

# %% [code]
renamed= pd.DataFrame({k: allmeans.loc[v] for k,v in cell_type_map.items()}).T

# %% [code]
# extract features for all the unique positions in the filtered associations
unique_pos = valid_long[['t2t_chr','t2t_ind']].drop_duplicates()
chrom_inds = unique_pos.groupby('t2t_chr').groups
cis_seq_batches = []
dat_inds =[]
for chrom,inds in chrom_inds.items():
    chm13_pos = unique_pos['t2t_ind'].loc[inds]
    chr_dat = np.load('../data/seq/GENA_2k_precomputed/{}_precomputed_features.npy'.format(chrom), mmap_mode = 'r')
    cis_seq_batches.append(chr_dat[chm13_pos.values])
    dat_inds+= list(inds)
cis_seq_batch = torch.tensor(np.concatenate(cis_seq_batches)).float().to('cuda:{}'.format(DEVICE))

# construct a map assigning chromosome inds to the features
qtl_to_seq_map = {ch_ind:i for i, ch_ind in enumerate (zip(unique_pos['t2t_chr'].loc[dat_inds],unique_pos['t2t_ind'].loc[dat_inds]))}


#construct maps assigning each cell type/gene to its indices in the seq batch and preds
ct_gene_groups = valid_long.groupby(['CELL_TYPE','GENE']).groups
seq_inds = {}
fdrs = {}
for key,inds in ct_gene_groups.items():
    CELLTYPE,GENE = key
    seq_inds[CELLTYPE, GENE] = np.array([qtl_to_seq_map[a] for a in zip(valid_long['t2t_chr'].loc[inds],valid_long['t2t_ind'].loc[inds])])
    fdrs[CELLTYPE,GENE] = valid_long['FDR'].loc[inds]
    
    


# %% [code]
# auxiliary function to prepare the original and edited batch
def prepare_sc_batch(ct, gene_to_add, amount = 0.0001):
    ct_mean = renamed.loc[ct]
    edited_mean = ct_mean.copy()
    edited_mean[gene_to_add]+=amount
    return torch.tensor(np.stack([ct_mean.values.copy(), edited_mean.values]),device = 'cuda:{}'.format(DEVICE),dtype = torch.float)
    

# %% [code]
@torch.no_grad()
def permutation_test(path,perm_gene = False,perm_ct= False,perm_rs = False,group_perm_gene = False,group_perm_rs = False,rand_init = False, nperm =1000,thresh = -1,n_control = 40,
                    collect_preds = False):
    softmax = True
    bn = False
    n_learn = 1
    n_control = n_control
    dropout= 0
    softmax = True
    bn = False
    multi_learner = nn.ModuleList([LatentPSALearner(sc = torch.from_numpy(renamed.values),dropout=dropout, softmax=softmax, genome = cis_seq_batch, n_control = n_control, bias = False,batch_norm = bn) for _ in range(n_learn)]).to('cuda:{}'.format(DEVICE))

    checkpoint = torch.load(path)
    sd = checkpoint['learner_state_dict']
    multi_learner.load_state_dict(sd)
    multi_learner.eval();
    learner = multi_learner[0]
    learner.eval();
    
    assert (group_perm_gene&perm_gene)==False
    assert (group_perm_rs&perm_rs)==False
    
    
    
    # initialize maps that will be shuffled during permutations
    orig_genes = valid_long['GENE'].unique()
    shuf_genes = orig_genes.copy()
    orig_cts = renamed.index.values.copy()
    shuf_cts = orig_cts.copy()
    shuf_idx = np.arange(len(cis_seq_batch))

    # want to enable shuffling only genes that are expressed in the same set of cell types, hence this shuffle
    ct_set_to_g = {}
    for g, ct_set in {g: tuple(sorted(renamed[g][renamed[g]>2e-5].index.values)) for g in orig_genes}.items():
        ct_set_to_g.setdefault(ct_set, []).append(g)

    orig_group_genes = [np.array(gs) for _,gs in ct_set_to_g.items()]
    shuf_group_genes = [arr.copy() for arr in orig_group_genes]

    #want to enable shuffling to get sense for resolution
    if group_perm_rs:
        concom = ccs[thresh]
        orig_group_rs = [np.where(concom[1]==k)[0] for k in range(concom[0])]
        assert (np.concatenate(orig_group_rs)-np.arange(len(concom[1]))).max()==0    
        shuf_group_rs = [arr.copy() for arr in orig_group_rs]


    assert (group_perm_gene&perm_gene)==False
    assert (group_perm_rs&perm_rs)==False


    perm_stats = {}
    PRINT_EVERY = min(nperm,10)
    np.random.seed(45)
    if collect_preds:
        y_preds = []
    for perm in tqdm(range(1+nperm)):
        # first permutation is always test statistic (i.e., no shuffle)
        if perm>0:
            if group_perm_gene:
                for arr in shuf_group_genes:
                    np.random.shuffle(arr) #only shuffle within groups
            if group_perm_rs:
                for arr in shuf_group_rs:
                    np.random.shuffle(arr) #only shuffle within groups
            if perm_gene:
                np.random.shuffle(shuf_genes)

            if perm_ct:
                np.random.shuffle(orig_cts)

            if perm_rs:
                np.random.shuffle(shuf_idx)

            if rand_init:
                multi_learner = nn.ModuleList([LatentPSALearner(sc = torch.from_numpy(renamed.values),dropout=dropout, softmax=softmax, genome = cis_seq_batch, n_control = n_control, bias = False,batch_norm = bn) for _ in range(n_learn)]).to('cuda:{}'.format(DEVICE))
                multi_learner.eval();
                learner = multi_learner[0]
                learner.eval();


        # if perm==0, shuf_gene_map, shuf_idx, shuf_ct_map are all identity
        if group_perm_gene:
            shuf_gene_map = {i:j for i,j in zip(np.concatenate(orig_group_genes), np.concatenate(shuf_group_genes))}
        else:
            shuf_gene_map = {i:j for i,j in zip(orig_genes,shuf_genes)}


        if group_perm_rs:
            shuf_idx = np.concatenate(shuf_group_rs)
        shuf_ct_map = {i:j for i,j in zip(orig_cts,shuf_cts)}


        # compute AUCs for the discrepancy
        for CELLTYPE, GENE in ct_gene_groups.keys():
            sub_seq = cis_seq_batch[shuf_idx][seq_inds[CELLTYPE,GENE]]/50
            y_true = (fdrs[CELLTYPE,GENE]<0.1).astype(int).values

            shuf_ct = shuf_ct_map[CELLTYPE]
            shuf_gene = shuf_gene_map[GENE]
            sc_batch = prepare_sc_batch(shuf_ct, shuf_gene,amount = 1e-4)

            Txs =  learner.T_xs(seq_batch= sub_seq, sc_batch = sc_batch)
#             Txs =  learner.sc_q(sc_batch) @ learner.seq_k(sub_seq).T
            y_pred = torch.abs(Txs[1]-Txs[0]).detach().cpu().numpy()
            if (collect_preds) & (perm==0):
                y_preds.append(y_pred)
            perm_stats[perm,CELLTYPE,GENE] = roc_auc_score(y_true= y_true, y_score = y_pred)

    res = pd.Series(perm_stats).reset_index()
    res.columns = ['perm', 'ct', 'gene', 'auc']
    res['perm_gene'] = perm_gene
    res['group_perm_gene'] = group_perm_gene
    res['perm_rs'] = perm_rs
    res['perm_ct'] = perm_ct
    res['group_perm_rs'] = group_perm_rs
    res['rand_init'] = rand_init
    res['thresh'] = thresh
    if collect_preds:
        return res,y_preds
    else:
        return res


# %% [markdown]
### This is with the model with 40

# %% [code]
NPERM = 200
path = 'results/apr20/metacell_pbmc2_initchange/repeat_0_epoch_124.pt'
n_control = 40


results = []
    
perm_gene = False
perm_ct= False
perm_rs = False
group_perm_gene = False
rand_init = True ###
group_perm_rs = False
thresh = -1
results.append(permutation_test(path,perm_gene = perm_gene,perm_ct= perm_ct,perm_rs = perm_rs,group_perm_gene = group_perm_gene,rand_init = rand_init,
                 group_perm_rs = group_perm_rs,thresh = thresh,
                 nperm =NPERM, n_control = n_control))

perm_gene = False
perm_ct= False
perm_rs = False
group_perm_gene = True ###
rand_init = False
group_perm_rs = False
thresh = -1
results.append(permutation_test(path,perm_gene = perm_gene,perm_ct= perm_ct,perm_rs = perm_rs,group_perm_gene = group_perm_gene,rand_init = rand_init,
                 group_perm_rs = group_perm_rs,thresh = thresh,
                 nperm =NPERM, n_control = n_control))


perm_gene = False
perm_ct= False
perm_rs = True ###
group_perm_gene = False
rand_init = False
group_perm_rs = False
thresh = -1
results.append(permutation_test(path,perm_gene = perm_gene,perm_ct= perm_ct,perm_rs = perm_rs,group_perm_gene = group_perm_gene,rand_init = rand_init,
                 group_perm_rs = group_perm_rs,thresh = thresh,
                 nperm =NPERM, n_control = n_control))


perm_gene = False
perm_ct= True
perm_rs = False
group_perm_gene = False
rand_init = False
group_perm_rs = False
thresh = -1
results.append(permutation_test(path,perm_gene = perm_gene,perm_ct= perm_ct,perm_rs = perm_rs,group_perm_gene = group_perm_gene,rand_init = rand_init,
                 group_perm_rs = group_perm_rs,thresh = thresh,
                 nperm =NPERM, n_control = n_control))


perm_gene = True
group_perm_gene = False
perm_rs = False
group_perm_rs = False
rand_init = False
perm_ct= False
thresh = -1
results.append(permutation_test(path,perm_gene = perm_gene,perm_ct= perm_ct,perm_rs = perm_rs,group_perm_gene = group_perm_gene,rand_init = rand_init,
                 group_perm_rs = group_perm_rs,thresh = thresh,
                 nperm =NPERM, n_control = n_control))

combined = pd.concat(results, axis = 0,ignore_index = True)
combined.to_pickle('../data/downstream/Nature_pbmc2_cis_perm_results')


a = combined.groupby(['perm','perm_gene','perm_ct','group_perm_gene','group_perm_rs', 'perm_rs','rand_init','thresh','gene'])['auc'].mean().reset_index()


perm_conds = a.groupby(['perm_gene','perm_ct','group_perm_gene','perm_rs','group_perm_rs', 'rand_init','thresh']).groups
pvalues = {}
avgs = {}
for cond, inds in perm_conds.items():
    cond_perms = a.loc[inds]
    cond_avg = cond_perms.groupby('perm')['auc'].mean()
    keys = ['perm_gene','perm_ct','group_perm_gene','perm_rs','group_perm_rs', 'rand_init','thresh']
    if cond[-1]==-1:
        key = [a for a,b in zip(keys, cond) if b==True][0]
    else:
        key = 'group_perm_rs_{}'.format(cond[-1])    
    pvalues[key] = 1- (cond_avg[0]>cond_avg[1:]).mean()
    avgs[key] = a.loc[inds]
    avgs[key]['perm_key'] = key

    
res = pd.concat(avgs,ignore_index = True).groupby(['perm_key','perm'])['auc'].mean().reset_index()
test_stat = res['auc'].iloc[0]
test_stat

# %% [code]
print(ttest_1samp(avgs['perm_gene'].set_index(['perm','gene']).loc[0]['auc'],0.5))
print(pvalues)
print(test_stat)
dat = sns.stripplot(data = res[(~(res['perm_key'].str.contains('group_perm_rs'))) & (res['perm']!=0)], x = 'perm_key', y = 'auc',
                   s = 2.5, color = 'grey')
plt.xticks(rotation = 90)
plt.hlines(test_stat, -0.5, 4.5,colors = 'red',zorder = -1)

# %% [code]
plt.figure()
plt.figure(figsize = (.5,2))
dat = avgs['perm_gene'].set_index(['perm','gene']).loc[0]['auc']
sns.stripplot(dat,s = 2.5, c= 'darkgreen')
plt.hlines(dat.mean(), -0.2,0.2, colors = 'black')
plt.title('z40 model')

# %% [markdown]
### This is model with 80

# %% [code]
NPERM = 200

path  = 'results/apr20/metacell_pbmc2_80/repeat_0_epoch_83.pt' 
n_control = 80


results = []
    
perm_gene = False
perm_ct= False
perm_rs = False
group_perm_gene = False
rand_init = True ###
group_perm_rs = False
thresh = -1
results.append(permutation_test(path,perm_gene = perm_gene,perm_ct= perm_ct,perm_rs = perm_rs,group_perm_gene = group_perm_gene,rand_init = rand_init,
                 group_perm_rs = group_perm_rs,thresh = thresh,
                 nperm =NPERM, n_control = n_control))

perm_gene = False
perm_ct= False
perm_rs = False
group_perm_gene = True ###
rand_init = False
group_perm_rs = False
thresh = -1
results.append(permutation_test(path,perm_gene = perm_gene,perm_ct= perm_ct,perm_rs = perm_rs,group_perm_gene = group_perm_gene,rand_init = rand_init,
                 group_perm_rs = group_perm_rs,thresh = thresh,
                 nperm =NPERM, n_control = n_control))


perm_gene = False
perm_ct= False
perm_rs = True ###
group_perm_gene = False
rand_init = False
group_perm_rs = False
thresh = -1
results.append(permutation_test(path,perm_gene = perm_gene,perm_ct= perm_ct,perm_rs = perm_rs,group_perm_gene = group_perm_gene,rand_init = rand_init,
                 group_perm_rs = group_perm_rs,thresh = thresh,
                 nperm =NPERM, n_control = n_control))


perm_gene = False
perm_ct= True
perm_rs = False
group_perm_gene = False
rand_init = False
group_perm_rs = False
thresh = -1
results.append(permutation_test(path,perm_gene = perm_gene,perm_ct= perm_ct,perm_rs = perm_rs,group_perm_gene = group_perm_gene,rand_init = rand_init,
                 group_perm_rs = group_perm_rs,thresh = thresh,
                 nperm =NPERM, n_control = n_control))


perm_gene = True
group_perm_gene = False
perm_rs = False
group_perm_rs = False
rand_init = False
perm_ct= False
thresh = -1
results.append(permutation_test(path,perm_gene = perm_gene,perm_ct= perm_ct,perm_rs = perm_rs,group_perm_gene = group_perm_gene,rand_init = rand_init,
                 group_perm_rs = group_perm_rs,thresh = thresh,
                 nperm =NPERM, n_control = n_control))

combined = pd.concat(results, axis = 0,ignore_index = True)
combined.to_pickle('../data/downstream/Nature_pbmc2_cis_perm_results')


a = combined.groupby(['perm','perm_gene','perm_ct','group_perm_gene','group_perm_rs', 'perm_rs','rand_init','thresh','gene'])['auc'].mean().reset_index()


perm_conds = a.groupby(['perm_gene','perm_ct','group_perm_gene','perm_rs','group_perm_rs', 'rand_init','thresh']).groups
pvalues = {}
avgs = {}
for cond, inds in perm_conds.items():
    cond_perms = a.loc[inds]
    cond_avg = cond_perms.groupby('perm')['auc'].mean()
    keys = ['perm_gene','perm_ct','group_perm_gene','perm_rs','group_perm_rs', 'rand_init','thresh']
    if cond[-1]==-1:
        key = [a for a,b in zip(keys, cond) if b==True][0]
    else:
        key = 'group_perm_rs_{}'.format(cond[-1])    
    pvalues[key] = 1- (cond_avg[0]>cond_avg[1:]).mean()
    avgs[key] = a.loc[inds]
    avgs[key]['perm_key'] = key

    
res = pd.concat(avgs,ignore_index = True).groupby(['perm_key','perm'])['auc'].mean().reset_index()
test_stat = res['auc'].iloc[0]
test_stat

# %% [code]
print(ttest_1samp(avgs['perm_gene'].set_index(['perm','gene']).loc[0]['auc'],0.5))
print(pvalues)
print(test_stat)
dat = sns.stripplot(data = res[(~(res['perm_key'].str.contains('group_perm_rs'))) & (res['perm']!=0)], x = 'perm_key', y = 'auc',
                   s = 2.5, color = 'grey')
plt.xticks(rotation = 90)
plt.hlines(test_stat, -0.5, 4.5,colors = 'red',zorder = -1)

# %% [code]
plt.figure()
plt.figure(figsize = (.5,2))
dat = avgs['perm_gene'].set_index(['perm','gene']).loc[0]['auc']
sns.stripplot(dat,s = 2.5, c= 'darkgreen')
plt.hlines(dat.mean(), -0.2,0.2, colors = 'black')
plt.title('z80 model')

# %% [code]
print(pvalues)

# %% [code]
```
