## Supplementary material for "Deriving genetic codes for molecular phenotypes from first principles": Code: Fig4_PSA_attribution.ipynb

```
# %% [code]
import pandas as pd
import torch
import numpy as np
from tqdm import notebook
from torch import nn
from models.PSALearners import LatentPSALearner
from sklearn.metrics import roc_auc_score
from tqdm.notebook import tqdm

import seaborn as sns
from matplotlib import pyplot as plt

import pickle
import itertools


# %% [code]
#read the wherry atac profiles + lift to T2T
dat = pd.read_excel('wherry_atac.xlsx',sheet_name=None)
from pyliftover import LiftOver
CP_chroms = ['CP068277.2', 'CP068276.2', 'CP068275.2', 'CP068274.2', 'CP068273.2', 'CP068272.2', 'CP068271.2', 'CP068270.2', 'CP068269.2', 'CP068268.2', 'CP068267.2', 'CP068266.2', 'CP068265.2', 'CP068264.2', 'CP068263.2', 'CP068262.2', 'CP068261.2', 'CP068260.2', 'CP068259.2', 'CP068258.2', 'CP068257.2', 'CP068256.2', 'CP068255.2', 'CP086569.2']
chroms = ['chr{}'.format(j) for j in list(range(1,23))+['X','Y']]
chrom_map = {cp:ch for cp,ch in zip(chroms,CP_chroms)}
cp_to_chr = {cp:ch for cp,ch in zip(CP_chroms,chroms)}

chrom_starts = {'chr1': 0,
 'chr2': 248388,
 'chr3': 491085,
 'chr4': 692191,
 'chr5': 885766,
 'chr6': 1067812,
 'chr7': 1239939,
 'chr8': 1400507,
 'chr9': 1546767,
 'chr10': 1697385,
 'chr11': 1832144,
 'chr12': 1967272,
 'chr13': 2100597,
 'chr14': 2214164,
 'chr15': 2315326,
 'chr16': 2415080,
 'chr17': 2511411,
 'chr18': 2595688,
 'chr19': 2676231,
 'chr20': 2737939,
 'chr21': 2804150,
 'chr22': 2849241,
 'chrX': 2900566}

lo = LiftOver('hg19-chm13v2.chain')

def liftover_region(chrom, pos):
    result = lo.convert_coordinate(chrom, pos)
    return result[0][1] if result else None

for key, df in dat.items():
    # Apply liftover to start and end
    df['t2t_start'] = df.apply(lambda row: liftover_region(row['chr'], row['start']), axis=1)
    df['t2t_end'] = df.apply(lambda row: liftover_region(row['chr'], row['end']), axis=1)
    
    # Drop rows that failed liftover
    df = df.dropna(subset=['t2t_start', 't2t_end'])
    df['ind'] = (df['chr'].map(chrom_starts)+df['t2t_start']//1000)
    df = df.dropna()
    df['ind'] = df['ind'].astype(int)
    dat[key] = df
    

# %% [code]
genome = (torch.from_numpy(np.load('../data/seq/GENA_2k_precomputed/concatenated_genome.npy'))/10).float()

# %% [code]
from sklearn.decomposition import PCA
    

# %% [code]
means = pd.read_pickle('../data/sc/processed/pbmc_SEACELL_means')
assns = pd.read_pickle('../data/sc/processed/pbmc_SEACell_assignments')
freqs = assns.groupby(14).apply(lambda w : w[8].value_counts(normalize=True)).unstack().fillna(0)

n_learn=1
l1_means = pd.read_pickle('../data/downstream/pbmc_l1_means')
# compute first pc for each cell type
pca = PCA(n_components = 1)
pc0 = {}
for ct in freqs.columns:
    pca.fit (means.loc[freqs[ct][freqs[ct]>0.5].index])
    pc0[ct] = pd.Series(pca.components_[0],index = means.columns)

pc_df = pd.concat(pc0).unstack().loc[l1_means.index]


DEVICE =1
multi_learner = nn.ModuleList([LatentPSALearner(sc = torch.from_numpy(l1_means.values), softmax=True, dropout = 0, genome = genome, n_control = 40, bias = False,batch_norm = False) for _ in range(n_learn)]).to('cuda:{}'.format(DEVICE))

path  = 'results/apr5/metacell_fixed_seq_targ/repeat_0_epoch_76.pt'
checkpoint = torch.load(path,map_location = 'cpu')
sd = checkpoint['learner_state_dict']
multi_learner.load_state_dict(sd)
learner = multi_learner[0].to('cuda:{}'.format(DEVICE))

# define edits as increasing/decreasing PCs
sc_batch = torch.from_numpy(l1_means.values).to('cuda:{}'.format(DEVICE))
cop = l1_means.copy().values
cop+= 0.0001*pc_df.values
sc_batch_pos = torch.from_numpy(cop).to('cuda:{}'.format(DEVICE))
cop = l1_means.copy().values
cop-= 0.0001*pc_df.values
sc_batch_neg = torch.from_numpy(cop).to('cuda:{}'.format(DEVICE))

#get the discrepancies increasing/decreasing each PC
with torch.no_grad():
    T_xs = learner.T_xs(seq_batch=genome.to('cuda:{}'.format(DEVICE)),sc_batch = (sc_batch).float()).cpu().numpy()
    T_xs_pos = learner.T_xs(seq_batch=genome.to('cuda:{}'.format(DEVICE)),sc_batch = (sc_batch_pos).float()).cpu().numpy()
    T_xs_neg = learner.T_xs(seq_batch=genome.to('cuda:{}'.format(DEVICE)),sc_batch = (sc_batch_neg).float()).cpu().numpy()

pos_diffs = pd.DataFrame(T_xs_pos-T_xs, index = l1_means.index).abs()
neg_diffs = pd.DataFrame(T_xs_neg-T_xs, index = l1_means.index).abs()

# %% [code]
plt.figure(figsize = (3,1))
genes = l1_means.columns[~l1_means.columns.str.contains('RPS|RPL')]
(pc0['CD8 T'].loc[genes].sort_values().iloc[np.arange(5).tolist()+ np.arange(-5,0).tolist()]).plot.bar(color = 'grey')

# %% [code]
from scipy.stats import ttest_ind,mannwhitneyu

# %% [code]
# compute ttest to evaluate pos vs neg diffs
df = dat['SCMR3+']
q = df[df['subset_comparison'] =='scmR3+_vs_temra']

scm_inds = q['ind'][(q['log2FoldChange']>0) & (q['padj']<0.00001)]
eff_inds = q['ind'][(q['log2FoldChange']<0)& (q['padj']<0.00001)]
q_chroms_scm= q['chr'][(q['log2FoldChange']>0)& (q['padj']<0.00001)]
q_chroms_eff= q['chr'][(q['log2FoldChange']<0)& (q['padj']<0.00001)]


diff = {}
ranges = np.arange(-25,25).tolist() 
for ct in l1_means.index:
    for shift in ranges :
        shift1 = shift
        shift2 = shift
        shift_scm_inds = np.maximum(np.minimum(scm_inds + shift1, len(genome)-1),0)
        shift_eff_inds = np.maximum(np.minimum(eff_inds + shift2, len(genome)-1),0)

        a1 = (pos_diffs-neg_diffs).loc[ct].loc[shift_scm_inds]
        a2 = (pos_diffs-neg_diffs).loc[ct].loc[shift_eff_inds]

        a,b = ttest_ind(a1,a2)
        c,d = mannwhitneyu(a1,a2)
        diff[ct,shift] = {'t':a,'lp_t':-np.log10(b), 'mw_p':-np.log10(d)}


for shift in ranges:
    shift1 = shift
    shift2 = shift
    shift_scm_inds = np.maximum(np.minimum(scm_inds + shift1, len(genome)-1),0)
    shift_eff_inds = np.maximum(np.minimum(eff_inds + shift2, len(genome)-1),0)
    a1 = []
    for chrom,ind in zip(q_chroms_scm,shift_scm_inds):
        start = 1000*(ind - chrom_starts[chrom])
        seq = str(records[chrom][start:(start+2000)]).upper()
        a1.append(len(re.findall('G|C',seq)))
    a2 = []
    for chrom,ind in zip(q_chroms_eff,shift_eff_inds):
        start = 1000*(ind - chrom_starts[chrom])
        seq = str(records[chrom][start:(start+2000)]).upper()
        a2.append(len(re.findall('G|C',seq)))
        
    a,b = ttest_ind(a1,a2)
    c,d = mannwhitneyu(a1,a2)
    diff['gc',shift] = {'t':a,'lp_t':-np.log10(b), 'mw_p':-np.log10(d)}

# %% [code]
plt.figure(figsize = (5,5))
diff_df = pd.DataFrame(diff).T
for ct,marker,color in zip(['CD8 T', 'CD4 T','B','NK'],['-','--','-.',':'],['red'] + 3*['grey']):
    diff_df.loc[ct]['lp_t'].sort_index().plot(label= ct,color = color,linestyle = marker)
plt.legend()
plt.ylabel('-log10p')
plt.xlabel('kb from peak')
    

# %% [code]
plt.figure(figsize = (5,5))
diff_df = pd.DataFrame(diff).T
for ct,marker  in zip(['Mono', 'DC'] + ['gc'], ['--','-.',':']):
    diff_df.loc[ct]['lp_t'].sort_index().plot(label= ct,linestyle = marker,color = 'grey')
plt.legend()
    

# %% [code]


# %% [code]
from pyfaidx import Fasta
t2t_seq = Fasta('../GENA_LM/data/genomes/ncbi_dataset/data/GCA_009914755.4/GCA_009914755.4_T2T-CHM13v2.0_genomic.fna')

# %% [code]
records = {cp_to_chr[cp]:v for cp,v in t2t_seq.items() if cp in cp_to_chr} 

# %% [code]
#get seqs
seqs = {}
for idx,ind,chrom in zip(q.index,q['ind'], q['chr']):
    start = 1000*(ind - chrom_starts[chrom])
    seq = str(records[chrom][start:(start+2000)]).upper()
    seqs[idx] = seq

q['seq'] = pd.Series(seqs)    

    

# %% [code]
import re
q['gc'] = [len(re.findall('G|C',a)) for a in  q['seq']]

# %% [code]
#shuffle bins
np.random.seed(123)
all_seqs = []
selq = q[(q['log2FoldChange']>0) & (q['padj']<0.00001)]
for seq in tqdm(selq['seq']):
    all_seqs.append(seq)
    for i in np.arange(0,1975,25):
        ss = np.array(list(seq))
        np.random.shuffle(ss[i:i+50])
        all_seqs.append(''.join(ss))
        


# %% [code]
from transformers import AutoTokenizer, AutoModel

tokenizer = AutoTokenizer.from_pretrained('AIRI-Institute/gena-lm-bert-base')
model = AutoModel.from_pretrained('AIRI-Institute/gena-lm-bert-base', trust_remote_code=True)

model = model.to('cuda:{}'.format(DEVICE))

# %% [code]
corr_feats = []
model.eval()
with torch.no_grad():
    for seq in tqdm(all_seqs):
        corr_feats.append ((model(torch.LongTensor([[0] + tokenizer(seq)['input_ids']]).to('cuda:{}'.format(DEVICE)), output_hidden_states  = True)['hidden_states'][-1][0][1:].mean(axis = 0).to(torch.float16).detach().cpu().numpy())/50)        

        
pd.DataFrame(np.stack(corr_feats)).to_pickle('sliding_corrupt_jun9')
    
    


# %% [code]
corr_feats = pd.read_pickle('sliding_corrupt_jun9')

# %% [code]
corr_feats.shape

# %% [code]
motif.weblogo('hi.png')

# %% [code]
from Bio import motifs
from Bio.Seq import Seq


with open("JASPAR2020_CORE_vertebrates_non-redundant_pfms_jaspar.txt") as handle:
    motif_list = list(motifs.parse(handle, "jaspar"))

results = []

all_pwm_scores = {}
for motif in motif_list:
    if motif.name.upper() not in ['TCF7']:
        continue
    
    pwm = motif.counts.normalize(pseudocounts=0.1).log_odds()
    motif_len = len(pwm['A'])
    pwm_scores = []
    for i, seq_str in tqdm(enumerate(selq['seq'])):
        seq = Seq(seq_str)

        a_f = np.pad(pwm.calculate(seq), (0,motif_len-1))
        a_r = np.pad(pwm.calculate(seq.reverse_complement()),(0,motif_len-1))[::-1]
        pwm_scores.append(np.stack([a_f,a_r]).max(axis = 0))
        
        

    all_pwm_scores[motif.name] = np.stack(pwm_scores)

# %% [code]
sc_batch = torch.from_numpy(l1_means.values).to('cuda:{}'.format(DEVICE))
cop = l1_means.copy().values
cop+= 0.0001*pc_df.values
sc_batch_pos = torch.from_numpy(cop).to('cuda:{}'.format(DEVICE))
cop = l1_means.copy().values
cop-= 0.0001*pc_df.values
sc_batch_neg = torch.from_numpy(cop).to('cuda:{}'.format(DEVICE))


def get_disc(feats):
    seq_batch=torch.from_numpy(feats).float().to('cuda:{}'.format(DEVICE))
    with torch.no_grad():
        T_xs = learner.T_xs(seq_batch=seq_batch,sc_batch = (sc_batch).float()).cpu().numpy()
        T_xs_pos = learner.T_xs(seq_batch=seq_batch,sc_batch = (sc_batch_pos).float()).cpu().numpy()
        T_xs_neg = learner.T_xs(seq_batch=seq_batch,sc_batch = (sc_batch_neg).float()).cpu().numpy()
    
    
    pos_diffs_sub = pd.DataFrame(T_xs_pos-T_xs, index = l1_means.index).abs()
    neg_diffs_sub = pd.DataFrame(T_xs_neg-T_xs, index = l1_means.index).abs()
    return  (pos_diffs_sub-neg_diffs_sub)


# disc_orig = get_disc(feats = np.stack([feats_orig[a] for a in q2.index]))
# disc_shuf = get_disc(feats = np.stack([feats_rem[a] for a in q2.index]))


# %% [code]
disc_corr = get_disc(corr_feats.values)
R = disc_corr.loc['CD8 T'].values.reshape(-1, 80)
scores = R[:,1:] - (R[:,0][:,None])
scores = (scores-scores.mean(axis=1,keepdims=True))/scores.std(axis = 1,keepdims=True)

# %% [code]
res_unique = {}

for num_seq in np.arange(100,700,100):
    for thr in [-2,-2.5,-3]:
        for score_thr in [5]:
            for name in ['TCF7']:
                sel_inds = np.argsort(R[:,0])[-num_seq:]
                pwm_scores=  all_pwm_scores[name][sel_inds]
                perm_scores = scores.copy()[sel_inds]
                stat = []
                for perm in range(2000):
                    if perm>0:
                        for i in range(len(perm_scores)):
                            np.random.shuffle(perm_scores[i])
                    motif_scores = []
                    ii,jj = np.where(perm_scores<thr)
                    a,b = np.unique(ii,return_counts = True)
                    uniques = a[b<3]
                    is_unique_i = np.isin(ii, uniques)
                    for i,j in zip(ii[is_unique_i],jj[is_unique_i]):
                        motif_scores.append(pwm_scores[i,(j*25):(j*25+50)].max())
                    
                        
                    stat.append((np.array(motif_scores)>score_thr).sum())
                res_unique[name,num_seq,thr,score_thr]= np.array(stat)

# %% [code]
res = {}

for num_seq in np.arange(100,700,100):
    for thr in [-2,-2.5,-3]:
        for score_thr in [5]:
            for name in ['TCF7']:
                sel_inds = np.argsort(R[:,0])[-num_seq:]
                pwm_scores=  all_pwm_scores[name][sel_inds]
                perm_scores = scores.copy()[sel_inds]
                stat = []
                for perm in range(2000):
                    if perm>0:
                        for i in range(len(perm_scores)):
                            np.random.shuffle(perm_scores[i])
                    motif_scores = []
                    ii,jj = np.where(perm_scores<thr)
                    for i,j in zip(ii,jj):
                        motif_scores.append(pwm_scores[i,(j*25):(j*25+50)].max())
                    
                        
                    stat.append((np.array(motif_scores)>score_thr).sum())
                res[name,num_seq,thr,score_thr]= np.array(stat)

# %% [code]
plt.figure(figsize = (5,5))
tcf7_res = pd.Series({k: (v[0]>v[1:]).mean() for k,v in res_unique.items()})
r = tcf7_res.reorder_levels([0,3,2,1]).loc['TCF7',5].loc[[-2.5,-3]]
(-np.log10(1-r.loc[-2.5])).plot(label = '2.5sd',color = 'red',linestyle = '--')
(-np.log10(1-r.loc[-3])).plot(label = '3sd',color = 'red', linestyle = '-')
plt.legend()
plt.ylabel('-log10p TCF7 motif enrichment')
plt.xlabel('top n peaks')
plt.hlines(-np.log10(0.05),100,600, color = 'grey', linestyle = ':', linewidth = 2)
plt.title('peaks w unique high-importance region')

# %% [code]
plt.figure(figsize = (5,5))
tcf7_res = pd.Series({k: (v[0]>v[1:]).mean() for k,v in res.items()})
r = tcf7_res.reorder_levels([0,3,2,1]).loc['TCF7',5].loc[[-2.5,-3]]
(-np.log10(1-r.loc[-2.5])).plot(label = '2.5sd',color = 'red',linestyle = '--')
(-np.log10(1-r.loc[-3])).plot(label = '3sd',color = 'red', linestyle = '-')
plt.legend()
plt.ylabel('-log10p TCF7 motif enrichment')
plt.xlabel('top n peaks')
plt.hlines(-np.log10(0.05),100,600, color = 'grey', linestyle = ':', linewidth = 2)
plt.title('all peaks')

# %% [code]


# %% [code]


# %% [code]


# %% [code]
```
