## Supplementary material for "Deriving genetic codes for molecular phenotypes from first principles": Code: Fig5_crc_subset_codes.ipynb

```
# %% [code]
#from scanpy import read_h5ad
import numpy as np
import pandas as pd
import pickle 
from tqdm.notebook import tqdm
import torch
from torch import nn


import os
from matplotlib import pyplot as plt
%pylab inline
import seaborn as sns

from numpy.lib.stride_tricks import sliding_window_view
from tqdm.notebook import tqdm


from scipy.stats import mannwhitneyu,ttest_ind,spearmanr
from statsmodels.stats.multitest import fdrcorrection
from sklearn.metrics import roc_curve,auc
from sklearn.neighbors import NearestNeighbors
from scipy.spatial.distance import cdist
import seaborn as sns
from scipy.cluster.hierarchy import fcluster


from models.PSALearners import LatentPSALearner

from sklearn.metrics import roc_auc_score
torch.cuda.set_device(3)
DEVICE = 3

# %% [code]
from sklearn.neighbors import NearestNeighbors
from tqdm.notebook import tqdm
x = pd.read_csv('../data/sc/raw/crc.csv')
x['Cluster name2'] = x['ClusterName'].astype('category')
freq_col = 'Cluster name2'
num_neighbors = 20
window_dataset = []
knn = NearestNeighbors(n_neighbors=num_neighbors)
xinds = []
for exp in tqdm(x['spots'].unique()):
    exp_cells = x[x['spots']==exp]
    dummies = pd.get_dummies(exp_cells[freq_col])
    knn.fit(exp_cells[['X:X','Y:Y']])
    knng = knn.kneighbors_graph()
    window_dataset.append(dummies.values[knng.nonzero()[1].reshape(-1,num_neighbors).flatten()].reshape(-1,num_neighbors,dummies.shape[1]).sum(axis = 1))
    xinds.append(exp_cells.index.values.copy())

all_windows = np.concatenate(window_dataset, axis = 0)
sc = pd.DataFrame(np.log(1+all_windows), index = np.concatenate(xinds),columns = dummies.columns)

# %% [code]
cd8mbk = MiniBatchKMeans(n_clusters = 100)
cd8mbk.fit(sc.loc[x.index[(x['Cluster name2']=='CD8+ T cells')]])
cd8_neighs = pd.DataFrame(cd8mbk.cluster_centers_,columns = dummies.columns)
# cd8_neighs.to_pickle('crc_ot_intermediate_files/cd8_neighs')

# %% [code]
cd8_neighs = pd.read_pickle('crc_ot_intermediate_files/cd8_neighs')

# %% [code]
mbk_labels = np.load('crc_ot_intermediate_files/mbk_labels.npy')
mbk_centers = np.load('crc_ot_intermediate_files/mbk_centers.npy')

# %% [code]
genome = np.load('../data/seq/GENA_2k_precomputed/concatenated_genome.npy')

# %% [code]
from sklearn.cluster import MiniBatchKMeans
from scipy.sparse.csgraph import shortest_path
from sklearn.neighbors import kneighbors_graph

# %% [code]
from ot.gromov import semirelaxed_gromov_wasserstein as gw

# %% [code]
mbk = MiniBatchKMeans(n_clusters = 20)
mbk.fit(genome)
np.save(os.path.join('crc_ot_intermediate_files','mbk_labels'), mbk.labels_)
np.save(os.path.join('crc_ot_intermediate_files','mbk_centers'), mbk.cluster_centers_)

# %% [code]


# %% [markdown]
### Plot the schematic

# %% [code]
from scipy.cluster import hierarchy

# %% [code]
link = sns.clustermap(mbk_centers,figsize = (0.01,0.01)).dendrogram_row.linkage;

# %% [code]
plt.figure(figsize = (5,1))
plt.axis('off')
plt.hist(np.random.randn(4500),color = 'grey',bins = 50)

# %% [code]


# %% [code]
link2 = sns.clustermap(shortest_path(kneighbors_graph(sub_mbk_centers[8,9],metric = 'cosine',n_neighbors = 10),directed = False),figsize = (.1,.1)).dendrogram_row.linkage


# %% [code]
hierarchy.dendrogram(link2,truncate_mode = 'lastp')
plt.show()


# %% [code]
plt.figure(figsize = (10,1))
with plt.rc_context({'lines.linewidth': 2.5, 'figure.facecolor':(0,0,0,0)}):
    ll = hierarchy.dendrogram(link2,link_color_func= lambda w: 'black',truncate_mode = 'lastp')
    

    w = np.concatenate([np.array(ic)[np.where(np.array(dc)==0)] for ic,dc in zip(ll['icoord'], ll['dcoord']) if (np.array(dc)==0).sum()>0])
    plt.scatter(w, [0]*len(w),color = 'black',zorder = 2, s= 100)
    plt.ylim(-5)
    plt.axis('off')
    plt.show()

# %% [code]


# %% [code]
plt.figure(figsize = (10,1))
with plt.rc_context({'lines.linewidth': 2.5}):
    hierarchy.dendrogram(link,link_color_func= lambda w: 'black')
    plt.scatter(range(20),[0]*20)
    plt.show()

# %% [markdown]
### Run OT multiple times per superclustering with reproducible seeds

# %% [code]
sd = shortest_path(kneighbors_graph(cd8_neighs,n_neighbors=10,metric = 'cosine'),directed=False)
sub_mbk_labels = {}
sub_mbk_centers = {}
sub_mbk_assns = {}
base = {}

np.random.seed(1851)
seeds =  np.unique(np.random.randint(0,100000, size = 20000))[:5000]
seed_index = 0
for up_clust in tqdm(range(20)):
    if up_clust ==2:
        continue
    for cl in range(100):
        seed_index +=1
        sub_mbk = MiniBatchKMeans(n_clusters = 500,random_state = seeds[seed_index])
        sub_mbk.fit(genome[mbk_labels==up_clust])
        sub_mbk_labels[up_clust,cl] = sub_mbk.labels_.copy()
        sub_mbk_centers[up_clust, cl]= sub_mbk.cluster_centers_.copy()
        r = 20
        maxr = 10000
        while maxr>1000:
            sub_cd = shortest_path(kneighbors_graph(sub_mbk.cluster_centers_,n_neighbors = r),directed=False)
            maxr = sub_cd.max()
            r+=1
        
        a = torch.from_numpy(sd/sd.max()).float().cuda()
        
        b = torch.from_numpy(sub_cd/sub_cd.max()).float().cuda()
        res = gw(a,b,log = True)
        base[up_clust,cl] = res[1]['srgw_dist'].item()
        assns = {}
        for i in range(len(sd)):
            inds = [a for a in range(len(sd)) if a!=i]
            a = torch.from_numpy(sd[inds,:][:,inds]/sd.max()).float().cuda()
            
            res = gw(a,b,log = True)
            assns[i] = res[0].argmax(axis=1).cpu().numpy()
        sub_mbk_assns[up_clust,cl] = assns.copy()
        counts = {}
        for i,v in assns.items():
            for j in v:
                counts.setdefault(j,set()).add(i)
    

# %% [code]
print('hi')

# %% [code]
for k,v in sub_mbk_assns.items():
    np.save(os.path.join('crc_ot_intermediate_files', 'sub_mbk_assns{}'.format(k)), v)
for k,v in sub_mbk_centers.items():
    np.save(os.path.join('crc_ot_intermediate_files', 'sub_mbk_centers{}'.format(k)), v)
for k,v in sub_mbk_labels.items():
    np.save(os.path.join('crc_ot_intermediate_files', 'sub_mbk_labels{}'.format(k)), v)
pd.Series(base).to_pickle(os.path.join('crc_ot_intermediate_files','srgw_loss'))

# %% [code]
base = pd.read_pickle(os.path.join('crc_ot_intermediate_files/','srgw_loss'))

# %% [code]
pd.Series(base).loc[8].sort_values()

# %% [code]
pd.Series(base).loc[8].sort_values()

# %% [code]
print('hi')

# %% [code]
b = pd.Series(base).sort_values().dropna().reset_index()

# %% [code]
b['clust'] = b['level_0'].astype(str)

# %% [code]

orders = b.groupby('clust')[0].median().sort_values().index.values

# %% [code]
orders

# %% [code]
from scipy.stats import ttest_ind,mannwhitneyu

# %% [code]
diffs = {}
for i in range(20):
    for j in range(20):
        s, p = ttest_ind(base.loc[i],base.loc[j])
        if p<0.05:
            diffs[i,j] = s

# %% [code]
plt.figure(figsize = (5,2))
sns.stripplot(data = b,x = 'clust',y = 0, order = orders,s= 2, c = 'black')
plt.xlabel('')
plt.ylabel('alignment loss')

# %% [markdown]
### Lift to hg38 for ENCODE analysis

# %% [code]
from liftover import get_lifter

converter = get_lifter('hs1', 'hg38', one_based=True)
converter_hg19 = get_lifter('hs1', 'hg19', one_based=True)
chrom = '1'
pos = 103786442
converter[chrom][pos]

chr_names = np.array(list(range(1,23))+['X'])
chrom_ends = np.cumsum([np.load('../data/seq/GENA_2k_precomputed/chr{}_precomputed_features.npy'.format(i),mmap_mode = 'r').shape[0] for i in chr_names])

chrom_starts = np.concatenate([[0], chrom_ends])

names = []
for chrom, start,stop in tqdm(zip(chr_names,chrom_starts,chrom_ends)):
    print(chrom,start,stop)
    for pos in np.arange(stop-start):
        names.append('chr{}:{}-{}'.format(chrom, 1000*pos, 1000*pos+2000))


to_lift = [(a.split(':')[0], int(a.split(':')[1].split('-')[0])) for a in names]
lifted_hg38 = [[name,a[0],a[1]-1000, a[1]+1000] for name,a in [(name,z[0]) for (name,z) in [(name,converter[a][b]) for name,(a,b) in zip(names,to_lift)] if len(z)>0]]


# %% [code]
import pyranges as pr

# %% [code]

hg38_encode = pd.read_table('encode.bed',header = None)[[0,1,2,10]].rename({0:'Chromosome',1:'Start',2:'End',10:'type'},axis=1)
encode_beds = {}
for k,v in hg38_encode.groupby('type').groups.items():
    encode_beds[k] = pr.PyRanges(hg38_encode.loc[v])
    
    

# %% [code]
# hg38 = pd.concat([pd.DataFrame(lifted_hg38).set_index(0),pd.DataFrame({'label':mbk_labels},index =names)],axis=1).dropna()
joined_all = pr.PyRanges(hg38.rename({1:'Chromosome',2:'Start',3:'End'},axis=1).reset_index()).join(pr.PyRanges(hg38_encode)).as_df()


# %% [code]
from scipy.stats import fisher_exact

# %% [code]
mbk_ccre = {}
for typ in joined_all['type'].unique():
    N = len(set(joined_all['index']))
    n = len(set(joined_all['index'][joined_all['type']==typ]))
    for i in range(20):
        m = len(set(joined_all['index'][(joined_all['type']==typ) & (joined_all['label']==i)]))
        M = len(set(joined_all['index'][(joined_all['label']==i)]))
        fe = fisher_exact([[m,M-m],[n,N-n]])
        if fe.pvalue<0.000001:
            mbk_ccre[typ,i] = fe.statistic


# %% [code]
orders

# %% [code]
plot_dat = np.log2(1e-5+pd.Series( mbk_ccre)).unstack().fillna(0)
plot_dat = plot_dat.reindex(np.array(orders).astype(int),axis=1).dropna(axis=1)
cg = sns.clustermap(plot_dat,cmap = 'bwr', vmin = -2, vmax = 2,col_cluster = False,figsize = (0.1,0.1))
plt.figure(figsize = (5,1))
sns.heatmap(plot_dat.loc[['dELS','pELS','PLS','DNase-H3K4me3', 'CTCF-only']],yticklabels = True,xticklabels = False, cbar = False, cmap = 'bwr',vmin = -2, vmax = 2)

# %% [code]


# %% [code]


# %% [code]


# %% [markdown]
### Empirical null distribution for phenotype corruption

# %% [code]

assn = {}
pv = {}
for k, centers in tqdm(sub_mbk_centers.items()):
    if k[0] not in [8,9,1]:
        continue
    if k[1] not in pd.Series(base).loc[k[0]].sort_values().index[:5]:
        continue
    r = 20
    maxr = 10000
    while maxr>1000:
        sub_cd = shortest_path(kneighbors_graph(centers,n_neighbors = r),directed=False)
        maxr = sub_cd.max()
        r+=1
    
    
    b = torch.from_numpy(sub_cd/sub_cd.max()).float().cuda()
    
    
    
    
    
    losses = []
    sds = []
    for perm in range(2000):
        mat = cd8_neighs.values.copy()
        if perm>0:
            i,j = np.random.choice(np.arange(len(mat)),size = 2, replace=False)
            t = 0.25 *np.random.rand()
            mat[i] =  t*(mat[j] - mat[i]) + (1-t)*mat[i]
        sd = shortest_path(kneighbors_graph(mat,n_neighbors=10,metric='cosine'),directed=False)
        sds.append(sd)
        a = torch.from_numpy(sd/sd.max()).float().cuda()
        res = gw(a,1*b,log = True)
        if perm==0:
            assn[k] = res[0].argmax(axis = 1).cpu().numpy()
        losses.append(res[1]['srgw_dist'].item())
        # aa = pd.Series(res[0].argmax(axis=1).cpu().numpy())
        # sel_inds = np.unique(aa)
    losses=np.array(losses)
    diff = np.where(np.abs(np.stack(sds[1:])-sds[0][None]).max(axis = (1,2))>0)
    pv[k] =(losses[1:][diff]>losses[0]).mean()
    

# %% [code]
pd.Series(base).loc[8].sort_values()

# %% [code]
dat[0]

# %% [code]
dat = pd.concat([-np.log10(1-pd.Series(pv)),pd.Series(base)],axis=1).dropna().sort_values(0).reset_index()

dat['cluster'] = dat['level_0'].astype(str)
plt.figure(figsize = (5,5))
sns.scatterplot(data = dat, x = 1 ,y =0, hue = 'cluster',hue_order = ['8','9','1'])
plt.xlabel('alignment loss')
plt.xticks(np.linspace(0.0102,0.0106,5),np.linspace(0.0102,0.0106,5).round(5));
plt.ylabel('-log10p over corrupt input')


# %% [code]
dat.sort_values(0)

# %% [markdown]
### Analyse the enrichment of TF motifs for top code

# %% [code]
import numpy as np

# %% [code]
chr_names = np.array(list(range(1,23))+['X'])
chrom_lens = np.cumsum([np.load('../data/seq/GENA_2k_precomputed/chr{}_precomputed_features.npy'.format(i),mmap_mode = 'r').shape[0] for i in chr_names])
chrom_starts = {'chr{}'.format(chrom): v for chrom,v in zip(chr_names, [0]+chrom_lens.tolist())}

# %% [code]
'../data/seq/GENA_2k_precomputed/concatenated_genome.npy'

# %% [code]
chrom_starts

# %% [code]
#select TSS positions
transcripts = genes[genes[2]=='transcript']
transcripts['id'] =[a.split(';')[0].split(' ')[-1][1:-1] for a in transcripts[8]]
transcripts['chrom'] = transcripts[0].replace(cp_to_chr)
data = transcripts[['chrom',3,4,6,'id']].drop_duplicates()
data = data[data['chrom']!='chrY']
data.columns = ['chrom', 'start', 'stop', 'strand', 'gene_symbol']
data['tss'] = [data['start'].iloc[i] if (s== '+') else data['stop'].iloc[i] for i,s in enumerate(data['strand'])]
data = data.sort_values(['chrom','tss'])
chrs = {chrom:data.loc[val] for chrom,val in data.groupby('chrom').groups.items()}
chr_names = np.array(list(range(1,23))+['X'])
chrom_lens = np.cumsum([np.load('../data/seq/GENA_2k_precomputed/chr{}_precomputed_features.npy'.format(i),mmap_mode = 'r').shape[0] for i in chr_names])

lens = pd.Series({chrom: v for chrom,v in zip(chroms, [0]+chrom_lens.tolist())})
chrom_starts = np.array([0] + chrom_lens.tolist())[:-1]
data['idx'] = lens.loc[data['chrom']].values + data['tss']//1000
data = data[~((data['gene_symbol'].str.contains('LOC')) |(data['gene_symbol'].str.contains('MIR')) |(data['gene_symbol'].str.contains('-'))|(data['gene_symbol'].str.contains('LINC')) |(data['gene_symbol'].str.contains('LINC|RPL|RPS'))|(data['gene_symbol'].str.contains('-')))]


# %% [code]


# %% [code]
sub_mbk_assns[8,9]= np.load('crc_ot_intermediate_files/sub_mbk_assns(8, 9).npy',allow_pickle = True).astype(dict).flatten()[0]
sub_mbk_centers[8,9]= np.load('crc_ot_intermediate_files/sub_mbk_centers(8, 9).npy')
sub_mbk_labels[8,9] = np.load('crc_ot_intermediate_files/sub_mbk_labels(8, 9).npy')

# %% [code]
assn = {}
key= (8,9)
for i,v in sub_mbk_assns[key].items():
    for ind,j in enumerate([k for k in range(len(cd8_neighs)) if k!=i]):
        assn.setdefault(j, []).append(v[ind])
count = {}
for k,v in assn.items():
    for j in v:
        count.setdefault(j,[]).append(k)
        alls = []
for i,v in count.items():
    for j in v:cd
        alls.append((i,j))


count = pd.Series(alls).value_counts()
good_clusts = np.unique([a[0] for a in count[count>20].index])

# %% [code]
sd = shortest_path(kneighbors_graph(cd8_neighs,n_neighbors=10,metric = 'cosine'),directed=False)

r = 20
maxr = 10000
while maxr>1000:
    sub_cd = shortest_path(kneighbors_graph(sub_mbk_centers[key],n_neighbors = r),directed=False)
    maxr = sub_cd.max()
    r+=1

a = torch.from_numpy(sd/sd.max()).float().cuda()

b = torch.from_numpy(sub_cd/sub_cd.max()).float().cuda()
overall_assn = gw(a,b,log = True)[0].argmax(axis=1).cpu().numpy()

# %% [code]
top_pairs = pd.Series({(overall_assn[i],i): np.sum(v ==overall_assn[i]) for i,v in assn.items()}).sort_values()

# %% [code]
#sanity check
key = (8,9)
w = []
k = 55
for j in range(100):
    if j<k:
        w.append(sub_mbk_assns[key][j][k-1])
    if j>k:
        w.append(sub_mbk_assns[key][j][k])

# %% [code]
(np.array(w)==253).sum()

# %% [code]
key

# %% [code]
hg38_clust8 = pd.concat([pd.DataFrame(lifted_hg38).set_index(0),pd.DataFrame({'label':sub_mbk_labels[key]},index = np.array(names)[np.where(mbk_labels==key[0])[0]])],axis=1).dropna()


# %% [code]
t2t_clust8 = pd.DataFrame({'label':sub_mbk_labels[key]},index = np.where(mbk_labels==key[0])[0])
t2t_clust8['chrom'] = [a.split(':')[0] for a in np.array(names)[t2t_clust8.index]]

# %% [code]
label_groups_idx = t2t_clust8.reset_index().groupby('label').groups

label_genes = {}
for label,inds in tqdm(t2t_clust8.groupby(['label']).groups.items()):
    closest_ind = np.abs(data['idx'].values[:,None] - inds.values[None,:])
    neq_chrom = data['chrom'].values[:,None] != t2t_clust8['chrom'].loc[inds].values[None,:]
    closest_ind[neq_chrom] = 1e10
    label_genes.setdefault(label, set()).update(data['gene_symbol'].iloc[np.where((closest_ind<20))[0]])

# %% [markdown]
#### Generate HOMER input commands

# %% [code]
for clust in good_clusts:
    dat = hg38_clust8[hg38_clust8['label']==clust].reset_index()[[1,2,3]]
    # dat = hg38_clust8.reset_index()[[1,2,3]]
    dat[2] = dat[2].astype(int)
    dat[3] = dat[3].astype(int)
    if len(dat)>100:
        print ("findMotifsGenome.pl bed_topclust_{} hg38 test_output{} -size 1000 -mask -p 8 ".format(clust,clust))
        # print ("findMotifsGenome.pl temp_bed_{} hg38 background_output{} -size 1000 -mask -p 8 -bg clust8_background".format(clust,clust))
    # print(clust,len(dat))
        dat.to_csv('bed_topclust_{}'.format(clust),sep = '\t',header=None,index=None)

# %% [code]
len(np.unique(overall_assn))

# %% [code]
for clust in overall_assn:
    if clust in good_clusts:
        continue
    
    dat = hg38_clust8[hg38_clust8['label']==clust].reset_index()[[1,2,3]]
    # dat = hg38_clust8.reset_index()[[1,2,3]]
    dat[2] = dat[2].astype(int)
    dat[3] = dat[3].astype(int)
    if len(dat)>100:
        print ("findMotifsGenome.pl bed_topclust_{} hg38 test_output{} -size 1000 -mask -p 8 ".format(clust,clust))
        # print ("findMotifsGenome.pl temp_bed_{} hg38 background_output{} -size 1000 -mask -p 8 -bg clust8_background".format(clust,clust))
    # print(clust,len(dat))
        dat.to_csv('bed_topclust_{}'.format(clust),sep = '\t',header=None,index=None)

# %% [code]
from models.utils import znorm

# %% [code]
sns.clustermap(znorm(cd8_neighs.T,axis=1),xticklabels=True,figsize = (20,5),cmap = 'bwr',vmin = -2, vmax = 2)

# %% [code]
x = pd.read_csv('../data/sc/raw/crc.csv')
x['Cluster name2'] = x['ClusterName'].astype('category')


# %% [code]
cluster_counts = x['Cluster name2'].value_counts(normalize = True)

# %% [code]
cluster_counts

# %% [code]
top_cts = x['Cluster name2'].value_counts(normalize = True).index

# %% [code]
from scipy.stats import rankdata
from scipy.spatial.distance import cdist


# %% [code]
len(np.unique(overall_assn))

# %% [markdown]
### Attribution via motif enrichment consistency specificity

# %% [code]
#build up homer log fold change data fram
good_seq_clust = top_pairs[top_pairs>-1].reset_index()['level_0'].unique()
# this is not filtering clusters at all

enr_clust = {}

for clust in overall_assn:
    if clust not in good_seq_clust:
        continue
    try:
        enr_clust[clust] = pd.read_table('test_output{}/knownResults.txt'.format(clust))
    except FileNotFoundError:
        continue
print(len(enr_clust))
clust_neigh = {}
for clust in enr_clust:
    clust_neigh[clust] = cd8_neighs.iloc[np.where(overall_assn==clust)[0]].mean(axis = 0)
a1 = pd.concat(clust_neigh).unstack().T #this is the dataframe which averages a genome cluster by the CNs that are assigned to it

homer = pd.concat(enr_clust)
homer['fdr'] = fdrcorrection(np.exp(homer['Log P-value']))[1]
homer['pc'] = [float(a.split('%')[0]) for a in homer['% of Target Sequences with Motif']]
homer['bgpc'] = [float(a.split('%')[0]) for a in homer['% of Background Sequences with Motif']]
homer['lfc'] = np.log2((1e-4+homer['pc'])/(1e-4+homer['bgpc']))


# %% [code]
len(good_seq_clust)

# %% [code]
from scipy.sparse.csgraph import laplacian

# %% [code]
graph = kneighbors_graph(cd8_neighs,n_neighbors=10,metric = 'cosine').toarray()

# %% [code]
num_sig_mots = enr.reset_index().groupby(['Motif Name']).size().sort_values()

# %% [code]
#permutation tests for overall smoothness
enr = homer.reset_index().groupby(['Motif Name','level_0'])['lfc'].min().unstack().fillna(0)
assn_map = {i:j for i,j in enumerate(overall_assn) if j in enr.columns}

mapped_inds = cd8_neighs.index.map(assn_map)
good_inds = ~np.isnan(mapped_inds)
V = enr[mapped_inds[good_inds]].values


laps ={}
normV = (V-V.mean(axis = 1,keepdims=True))/np.linalg.norm(V,axis=1,keepdims=True)
for i in range(normV.shape[1]):
    sub = np.array([a for a in range(normV.shape[1]) if a!=i])
    
    
    L = laplacian(graph[good_inds,:][:,good_inds],symmetrized = True)
    
    vL = (normV[:,sub]@L[sub,:][:,sub])
    laps[i] = pd.Series((vL*vL).sum(axis=1),index = enr.index)


perm_laps ={}
normV = (V-V.mean(axis = 1,keepdims=True))/np.linalg.norm(V,axis=1,keepdims=True)

shuf = np.arange(normV.shape[1])
L = laplacian(graph[good_inds,:][:,good_inds],symmetrized = True)
for i in range(50000):
    if i>0:
        np.random.shuffle(shuf)
    vL = (normV[:,shuf])@L
    perm_laps[i] = pd.Series((vL*vL).sum(axis=1),index = enr.index)

# %% [code]
#permutation test for smoothness, but that preserves repetition when assigned clusters are mapped to same points in the coupling (sanity check)
enr = homer.reset_index().groupby(['Motif Name','level_0'])['lfc'].min().unstack().fillna(0)
assn_map = {i:j for i,j in enumerate(overall_assn) if j in enr.columns}

mapped_inds = cd8_neighs.index.map(assn_map)
good_inds = ~np.isnan(mapped_inds)


equiv_perm_laps ={}

enr_take_cols = mapped_inds[good_inds].map({j:i for i,j in enumerate(enr.columns)})
shuf = np.arange(enr.shape[1])
L = laplacian(graph[good_inds,:][:,good_inds],symmetrized = True)
for i in tqdm(range(50000)):
    
    if i>0:
        np.random.shuffle(shuf)

    V = enr.values[:,shuf][:,enr_take_cols]
    normV = (V-V.mean(axis = 1,keepdims=True))/np.linalg.norm(V,axis=1,keepdims=True)
    vL = normV@L
    equiv_perm_laps[i] = pd.Series((vL*vL).sum(axis=1),index = enr.index)

# %% [code]
equiv_pvs = pd.concat(equiv_perm_laps).reset_index().groupby('Motif Name').apply(lambda w : (w[0].iloc[0]<w[0].iloc[1:].values).mean()).sort_values()

# %% [code]
#this is clustering within groups - not shown in paper
clusters = fcluster(cg.dendrogram_row.linkage,criterion = 'maxclust',t =30)[good_inds]
clust_arrs = [np.where(clusters==clust)[0] for clust in np.unique(clusters)]
clust_perm_laps ={}
normV = (V-V.mean(axis = 1,keepdims=True))/np.linalg.norm(V,axis=1,keepdims=True)

shuf = np.arange(normV.shape[1])

L = laplacian(graph[good_inds,:][:,good_inds],symmetrized = True)
for i in range(2000):
    if i>0:
        for arr in clust_arrs:
            np.random.shuffle(arr)
        shuf = np.concatenate(clust_arrs)
    vL = (normV[:,shuf])@L
    clust_perm_laps[i] = pd.Series((vL*vL).sum(axis=1),index = enr.index)

clust_pvs = pd.concat(clust_perm_laps).reset_index().groupby('Motif Name').apply(lambda w : (w[0].iloc[0]<w[0].iloc[1:].values).mean()).sort_values()

# %% [code]
w = pd.concat(laps).reset_index()
order = w.groupby('Motif Name')[0].max().sort_values().index

# %% [code]
pvs = pd.concat(perm_laps).reset_index().groupby('Motif Name').apply(lambda w : (w[0].iloc[0]<w[0].iloc[1:].values).mean()).sort_values()


# %% [code]
# this is corrupting the graph, so testing specificity for graph
enr = homer.reset_index().groupby(['Motif Name','level_0'])['lfc'].min().unstack().fillna(0)
assn_map = {i:j for i,j in enumerate(overall_assn) if j in enr.columns}

mapped_inds = cd8_neighs.index.map(assn_map)
good_inds = ~np.isnan(mapped_inds)
V = enr[mapped_inds[good_inds]].values # make sure we recompute this since it's getting reassigned somewhere


normV = (V-V.mean(axis = 1,keepdims=True))/np.linalg.norm(V,axis=1,keepdims=True)
cor_laps = {}
co = 0
for perm in tqdm(range(50000)):
    mat = cd8_neighs.values.copy()
    if perm>0:
        for _ in range(5):
            i,j = np.random.choice(np.arange(len(mat)),size = 2, replace=False)
            t = 0.25*np.random.rand()
            mat[i] =  t*(mat[j] - mat[i]) + (1-t)*mat[i]

    gr = kneighbors_graph(mat,n_neighbors=10,metric = 'cosine')[good_inds,:][:,good_inds]
    if ((gr.toarray()==graph[good_inds,:][:,good_inds]).min() & (perm>0)):
        continue
    
    L = laplacian(gr,symmetrized = True)
    
    vL = (normV@L)
    cor_laps[co] = pd.Series((vL*vL).sum(axis=1),index = enr.index)
    co+=1
  
cor_pvs = pd.concat(cor_laps).reset_index().groupby('Motif Name').apply(lambda w : (w[0].iloc[0]<w[0].iloc[1:].values).mean()).sort_values()  

# %% [code]
import networkx as nx

# %% [code]
w =cor_pvs[(equiv_pvs>0.99)& (pvs>0.99)].sort_values().iloc[-10:]
plt.figure(figsize = (5,2))
(-np.log10(1-equiv_pvs + 1e-6)).hist( color = 'black')
plt.xlabel('-log10 smoothness p-value')
plt.ylabel('number of motifs')

# %% [code]
w =cor_pvs[(equiv_pvs>0.99)& (pvs>0.99)].sort_values().iloc[-10:]
plt.figure(figsize = (2,5))
(-np.log10(1-equiv_pvs + 1e-6)).hist( color = 'black',orientation = 'horizontal')
plt.ylabel('-log10 smoothness p-value')
plt.xlabel('number of motifs')

# %% [code]
plt.figure(figsize = (5,2))
w.rename({i:i.split('(')[0] for i in w.index}).sort_values().iloc[-5:].plot.bar(color = 'black')
plt.ylim(0.6,0.85)
plt.ylabel('smoothness score')

# %% [code]
plt.figure(figsize = (2,5))
w.rename({i:i.split('(')[0] for i in w.index}).sort_values().iloc[-5:].plot.barh(color = 'black')
plt.xlim(0.6,0.85)
plt.xlabel('smoothness score')

# %% [code]
enr = homer[homer['fdr']<1e-5].reset_index().groupby(['Motif Name','level_0'])['lfc'].mean()
k = enr.loc[w.index[-5:]].unstack().fillna(0)

# %% [code]
k

# %% [code]
com = list(set(a1.columns).intersection(k.columns)) #a1 is the assigned cluster cn means, k is the enrichment for the most graph-specific smooth

# %% [code]
plot_cts = top_cts[:10].tolist() 

# %% [code]
from models.utils import znorm

# %% [markdown]
### Annotate the neighborhoods grouped by assignment

# %% [code]
# fg = sns.clustermap(znorm(a1[com].loc[plot_cts],axis=1),metric = 'correlation',figsize = (1,1))

fg = sns.clustermap(znorm(a1[com],axis=1).loc[plot_cts+['CD11c+ DCs']],metric = 'correlation',figsize = (1,1))
assn_clusts = fcluster(fg.dendrogram_col.linkage, t = 4,criterion = 'maxclust')
colors = np.array(sns.color_palette('tab20',20))[assn_clusts]
# sns.clustermap(znorm(a1[com].loc[plot_cts],axis=1),metric = 'correlation', col_colors = colors,figsize = (12,5),cmap = 'bwr',vmin = -2.5, vmax = 2.5,yticklabels=True)
pp = znorm(a1[com],axis = 1).loc[top_cts]
sns.clustermap(pp.rename(colnames,axis=1),metric = 'correlation',col_colors = pp.columns.map(colnames_colors),figsize = (12,10),cmap = 'Reds',vmin = 0, vmax = 2.5,yticklabels=True,
              )


# %% [code]
# colnames = {215:'plasma cell rich', 309:'tls', 485: 'vasculature/stroma', 218: 'macrophage/stroma', 449: 'tls border',  301: 'vasculature', 443:'tumor/T cell', 343:'tumor/stroma',223: 'tumor boundary', 350: 'CD8/macrophage', 404:'macrophage', 218:'macrophage/stroma/granulocyte', 133:'granulocyte/dc'}


colnames = {215:'macrophage: plasma cell', 309:'tls', 485: 'perivascular, stroma', 218: 'macrophage/stroma', 449: 'perivascular, B cell',  301: 'perivascular, macrophage', 443:'stroma: tumor/T cell', 343:'tumor',223: 'tumor: boundary', 350: 'macrophage, CD8', 404:'macrophage', 218:'stroma: macrophage/granulocyte/CD8', 133:'perivascular, granulocyte/dc'}

colnames_colors = {i:j for i,j in zip(colnames,sns.color_palette('tab20',20))}

# %% [code]
plot_hm = k[com].rename({i: i.split('(')[0] for i in k.index},axis=0).rename({i: i.split('(')[0] for i in k.index},axis=0)


# %% [code]
motif_heatmap = sns.clustermap(plot_hm.rename(colnames,axis=1).T,metric = 'cosine', row_colors = plot_hm.columns.map(colnames_colors),figsize = (6,10),cmap = 'Reds',vmin = 0, vmax = 0.5,xticklabels=True,
              row_linkage = fg.dendrogram_col.linkage)


# %% [markdown]
### Compute proximal TSS

# %% [code]
label_groups_idx = t2t_clust8.reset_index().groupby('label').groups

dist_labs = {}
for dist in [5,10,15,20,25,30,35,40,45,50]:
    if dist in dist_labs:
        continue
    dist_labs[dist] = {}
    for label,inds in tqdm(t2t_clust8.groupby(['label']).groups.items()):
        if label not in colnames:
            continue
        closest_ind = np.abs(data['idx'].values[:,None] - inds.values[None,:])
        neq_chrom = data['chrom'].values[:,None] != t2t_clust8['chrom'].loc[inds].values[None,:]
        closest_ind[neq_chrom] = 1e10
        dist_labs[dist].setdefault(label, set()).update(data['gene_symbol'].iloc[np.where((closest_ind<dist))[0]])

# %% [code]
#!wget https://storage.googleapis.com/public-download-files/hgnc/tsv/tsv/locus_types/gene_with_protein_product.txt
protein_coding_genes = pd.read_table('gene_with_protein_product.txt')['symbol']
gs = pd.Series(sorted(set(dist_labs[20][343] - set(dist_labs[5][343])).intersection(protein_coding_genes)))
print('\n'.join(gs[~gs.str.contains('^OR')]))

# %% [code]


for dist in sorted(dist_labs.keys()):
    gene_clust_counts = {}
    for clust, geneset in dist_labs[dist].items():
        for gene in geneset:
            gene_clust_counts.setdefault(gene, set()).add(clust)

    is4 = {k:v for k,v in gene_clust_counts.items() if len(v)>2}
    is5 = {k:v for k,v in gene_clust_counts.items() if len(v)>5}
    
    if len(is5)>0:
        print(dist)
        break

# %% [code]
is5

# %% [code]
tss_lens[(tss_lens-tss_lens['ZBTB20']).abs()<50000]

# %% [code]
tss_lens[is4.keys()].sort_values().iloc[-10:]

# %% [code]
clust8_t2t

# %% [code]


# %% [code]
clust8_t2t = pd.Series(sub_mbk_labels[key], index=np.where(mbk_labels==key[0])[0]).reset_index()
clust8_t2t['label'] = clust8_t2t[0].astype(str)
clust8_t2t['name'] = np.array(names)[np.where(mbk_labels==key[0])[0]]

# %% [code]
clust8_t2t['chrom'] = [a[0] for a in clust8_t2t['name'].str.split(':')]

# %% [code]
sub = clust8_t2t[clust8_t2t[0].isin(colnames)]

# %% [code]


# %% [code]
# preparing the gene information
genes = pd.read_table('../data/seq/GENA_2k_precomputed/hs1.110.20220412.ncbiRefSeq.gtf',header = None)

#map CP chroms to chrs
CP_chroms = ['CP068277.2', 'CP068276.2', 'CP068275.2', 'CP068274.2', 'CP068273.2', 'CP068272.2', 'CP068271.2', 'CP068270.2', 'CP068269.2', 'CP068268.2', 'CP068267.2', 'CP068266.2', 'CP068265.2', 'CP068264.2', 'CP068263.2', 'CP068262.2', 'CP068261.2', 'CP068260.2', 'CP068259.2', 'CP068258.2', 'CP068257.2', 'CP068256.2', 'CP068255.2', 'CP086569.2']
chroms = ['chr{}'.format(j) for j in list(range(1,23))+['X','Y']]
chrom_map = {cp:ch for cp,ch in zip(chroms,CP_chroms)}
cp_to_chr = {cp:ch for cp,ch in zip(CP_chroms,chroms)}

#select TSS positions
transcripts = genes[genes[2]=='transcript']
transcripts['id'] =[a.split(';')[0].split(' ')[-1][1:-1] for a in transcripts[8]]
transcripts['chrom'] = transcripts[0].replace(cp_to_chr)
data = transcripts[['chrom',3,4,6,'id']].drop_duplicates()
data = data[data['chrom']!='chrY']
data.columns = ['chrom', 'start', 'stop', 'strand', 'gene_symbol']
data['tss'] = [data['start'].iloc[i] if (s== '+') else data['stop'].iloc[i] for i,s in enumerate(data['strand'])]
data = data.sort_values(['chrom','tss'])
chrs = {chrom:data.loc[val] for chrom,val in data.groupby('chrom').groups.items()}

tss_lens = data.groupby('gene_symbol')['tss'].apply(lambda w: w.max() - w.min())

# %% [code]


# %% [markdown]
### Finding genes close to multiple cluster assignments

# %% [code]
from torch.nn import AvgPool1d,MaxPool1d
biglabs = pd.DataFrame(mbk_labels)
biglabs['sublab'] = -1
biglabs['sublab'][biglabs[0]==key[0]] = sub_mbk_labels[key]
rolling_counts = pd.get_dummies(biglabs['sublab'])[colnames.keys()].astype(float)
halfN = 25
mp = MaxPool1d(kernel_size=2*halfN+1, padding=halfN,stride = 1)
rc = np.concatenate([mp(torch.from_numpy(rolling_counts.iloc[start:end].values.T)).numpy().T for start,end in zip(chrom_starts,chrom_ends)],axis=0) #max pools with kernel size 2*halfN to get proximity in each chrom, have to split to avoid reading thru
window_add = pd.DataFrame(rc, columns = rolling_counts.columns)
data['close'] = window_add.sum(axis = 1).loc[data['idx'].values].values
biglabs['count'] = window_add.sum(axis = 1)
gene_symbol_prox = data.groupby('gene_symbol')['idx'].apply(lambda w: window_add.loc[w].max(axis = 0).sum())

# %% [code]
close_genes

# %% [code]
plt.figure(figsize = (30,2))
sns.scatterplot(data = biglabs[biglabs['sublab']>=0].reset_index(),x = 'index', y = 'count')
close_genes = data[data['gene_symbol'].isin(gene_symbol_prox.index[gene_symbol_prox>3])]
close_genes = close_genes[close_genes['gene_symbol'].isin(protein_coding_genes)]
gene_means = close_genes.groupby('gene_symbol')['idx'].mean()

plt.vlines(close_genes['idx'].values, 0, 6,colors='red')
plt.ylim(2.5,6.1)
print(gene_means.sort_values())

# %% [code]
from matplotlib.patches import Rectangle

# %% [markdown]
#### Plotting the tracks along genome

# %% [code]
plt.figure(figsize = (40,10))
sub = window_add[(biglabs['sublab']>=0) & (biglabs['count']>2)][plot_hm.columns[motif_heatmap.dendrogram_row.reordered_ind]]
for i in range(12):
    print(i,sub.columns[i])
    plt.scatter(sub.index, (i+1)*sub.iloc[:,i], color = colnames_colors[sub.columns[i]],s=200,alpha = 1)
    plt.text(0, i+1, colnames[sub.columns[i]],ha = 'right', va = 'center',color = colnames_colors[sub.columns[i]])
plt.vlines(gene_means.values, 0, 12.5,colors='black',zorder = -1)
plt.ylim(0.5,12.5)

chrom_ind = 0
for x0, width in [(i, j-i) for i,j in zip(chrom_starts,chrom_ends)]:
    
    plt.gca().add_patch(Rectangle([x0,0], height=12.5,width = width, color = ['black' if chrom_ind%2==0 else 'grey'][0],zorder = -1,alpha = 0.2,linewidth = 0))
    chrom_ind+=1
plt.xlim(-1e6, 3.5e6)

# %% [code]
from scipy.stats import mannwhitneyu
enr = homer[homer['fdr']<0.0001].reset_index().groupby(['Motif Name','level_0'])['lfc'].mean()
not_enr = homer[homer['P-value']>0.05].reset_index().groupby(['Motif Name','level_0'])['lfc'].mean()
cors = {}
tt = {}
uniques = {}
for mot in enr.reset_index()['Motif Name'].unique():
    
    com = sorted(enr.loc[mot].index)
    not_com =sorted(not_enr.loc[mot].index)
    if len(com)==1:
        uniques.setdefault(com[0],set()).add(mot)

    if (len(com)<5):
        continue

    a = enr.loc[mot].loc[com]
    b = a1[a.index].T
    
    rs,ps = ttest_ind(cd8_neighs.iloc[np.isin(overall_assn,com)],cd8_neighs.iloc[np.isin(overall_assn,not_com)])
    _,mwps = mannwhitneyu(cd8_neighs.iloc[np.isin(overall_assn,com)],cd8_neighs.iloc[np.isin(overall_assn,not_com)])
    
    for ct,(r,p,mwp) in zip(a1.index,zip(rs,ps,mwps)):
        tt[ct,mot] = [r,p,mwp]
        
    
    for ct in top_cts:
        r,p = spearmanr(b[ct].values,a.values)
        cors[ct,mot] = [r,p,len(a)]
    
    
cors = pd.DataFrame(cors).T
tt = pd.DataFrame(tt).T
tt['fdr'] = fdrcorrection(tt[2])[1]
cors['fdr'] = fdrcorrection(cors[1])[1]

cors['p'] = cors[1]
cors['p'][(cors[2]==5) & (cors[0].abs()>0.99)] = 2/120
cors['p'][(cors[2]==6) & (cors[0].abs()==1)] = 2/720
cors['fdr2'] = fdrcorrection(cors['p'])[1]

# %% [code]
gene_means.sort_values()

# %% [code]
sub['posbin'] = sub['index']//200
posbin_counts = sub.groupby('posbin')['label'].apply(lambda w: len(np.unique(w))).sort_values()

# %% [code]
data['dist2peak'] = np.abs(data['idx'].values[:,None] - 200*posbin_counts[posbin_counts>2].index.values[None,:]).min(axis = 1)

# %% [code]
data.groupby('gene_symbol')['dist2peak'].min().sort_values().loc['ZBTB20']

# %% [code]
2062*200

# %% [code]
data[(data['idx']-(2062*200)).abs()<200]

# %% [code]
new_cts = ['tumor cells','smooth muscle','stroma','granulocytes','vasculature','CD8+ T cells', 'CD4+ T cells CD45RO+','CD11c+ DCs','Tregs','B cells','plasma cells']

# %% [markdown]
### Investigating some of the enrichment

# %% [code]
from scipy.stats import pearsonr

# %% [code]
mot = 'Zac1(Zf)/Neuro2A-Plagl1-ChIP-Seq(GSE75942)/Homer'
com = sorted(enr.loc[mot].index)
not_com =sorted(not_enr.loc[mot].index)
if len(com)==1:
    uniques.setdefault(com[0],set()).add(mot)

a = enr.loc[mot].loc[com]
b = a1[a.index].T
plt.figure(figsize = (5,5))
u = a
v = b['B cells']
plt.scatter(v,u,c = 'black')
plt.xlabel('log B cell count')
plt.ylabel('Zac1 motif enrichment')
print(spearmanr(u,v), pearsonr(u,v))

# %% [code]
mot = 'Tbx21(T-box)/GM12878-TBX21-ChIP-Seq(Encode)/Homer'
com = sorted(enr.loc[mot].index)
not_com =sorted(not_enr.loc[mot].index)
if len(com)==1:
    uniques.setdefault(com[0],set()).add(mot)

a = enr.loc[mot].loc[com]
b = a1[a.index].T
plt.figure(figsize = (5,5))
u = a
v1 = b['lymphatics']
v2 = b['smooth muscle']
plt.scatter(v2,u,c = b['vasculature'])
# plt.ylabel('Tbx21 motif enrichment')
print(spearmanr(v1,v2), pearsonr(v1,v2))

# %% [code]
from scipy.cluster.hierarchy import fcluster

# %% [code]


# %% [code]
w = pd.concat([laps,num_sig_mots],axis=1).dropna()

# %% [code]
w['rat'] = w[1]/w[0]
```
