## Supplementary material for "Deriving genetic codes for molecular phenotypes from first principles": Code: GraphPSA.py

```
import torch
from torch import nn
import numpy as np


class GraphPSABase():
    def __init__(self, syntax_graph, genome_tensor, phenotype_graph,phenotype_tensor, sc_target,val_sc, val_target, learner, device = 'cpu:0',**learner_kwargs):

        '''
        provides base methods: extract batches, eval neighbors, loss 
        _sc_consistency_loss, _syntax_consistency_loss and _prox_consistency_loss have to be overriden 
        syntax graph and phenotype graph have to be num_neighbors x num nodes
        '''
        
        self.genome = genome_tensor
        self.syntax = syntax_graph
        self.sc = phenotype_tensor
        self.sc_knn = phenotype_graph
        self.sc_dim = phenotype_tensor.shape[1]
        self.seq_dim = genome_tensor.shape[1]
        self.sc_target = sc_target
        self.num_sc_neighbors = self.sc_knn.shape[0]
        self.num_seq_neighbors = self.syntax.shape[0]
        self.val_sc = val_sc
        self.val_target = val_target
        self.learner = learner(**learner_kwargs)
        self.device = device

    def to(self, device = 'cpu:0'):
        self.learner.to(device=device)
        self.device = device

    def extract_batches(self, sc_inds, seq_inds):
        sc_knn_inds = self.sc_knn[:,sc_inds]
        sc_batch = self.sc[sc_knn_inds]
        seq_knn_inds = self.syntax[:,seq_inds]
        seq_proximity_inds = torch.stack([seq_inds-1, seq_inds+1],dim = 0)
        seq_batch = self.genome[torch.cat([seq_knn_inds,seq_proximity_inds], dim = 0)]
        return sc_batch.to(self.device),seq_batch.to(self.device).float()

    def eval_neighbors(self,sc_batch,seq_batch,sc_inds,seq_inds):
        '''
        outputs a tensor T dim (num_sc_neighbors, num_sc_inds, num_seq_neighbors, num_seq_inds) 
        containing values of T(x,s) for the sc at sc_graph[(i,j)] and seq at seq_graph[(k,l)] 
        and tensor L of dim (num_sc_neighbors, num_sc_inds, 2,num_seq_inds)
        containing values of T(x,s) for sc at (i,j) and either side in genome of sc
        '''
        Txs = self.learner.T_xs(sc_batch=sc_batch.view(-1, self.sc_dim), seq_batch=seq_batch.view(-1, self.seq_dim))
        Txs_view = Txs.view(self.num_sc_neighbors,len(sc_inds), self.num_seq_neighbors +2, len(seq_inds))
        
        T = Txs_view[:,:,:-2,:] # split because last two are adjacent sequences
        L = Txs_view[:,:,-2:,:]

        return T,L


    def _sc_target_loss(self,sc_pred, sc_target):
        return torch.square(sc_pred - sc_target).mean()
    
    def _seq_targets(self):
        return 0

    def _seq_entropy(self, sc_syntax_vals):
        #sc_syntax_vals has ksc x nsc x kseq x nseq
        seq0 = sc_syntax_vals[:,:,0,:]
        v = torch.abs(seq0)
        probs = v/v.sum(axis = -1,keepdim = True)
        return (-probs*torch.log(1e-10 + probs)).mean()

    def loss(self, sc_inds, seq_inds, return_vals = False):
        '''
        graph consistency losses (sc, syntax, proximity)
        sc_targets losses (for integral over seq)
        seq_targets losses (for integral over sc)
        marking this for removal since we switched to multicomponent
        '''
        pass
        # sc_batch, seq_batch = self.extract_batches(sc_inds = sc_inds,seq_inds=seq_inds)
        # sc_syntax_vals,sc_L_vals = self.eval_neighbors(seq_batch = seq_batch, sc_batch = sc_batch, sc_inds = sc_inds, seq_inds=seq_inds)
        # sc_targets = self.sc_target[sc_inds].to(self.device)
        # seq_targets = self._seq_targets()

        # sc_consistency_loss = self._sc_consistency_loss(sc_syntax_vals,sc_batch = sc_batch)
        # syntax_consistency_loss = self._syntax_consistency_loss(sc_syntax_vals,seq_batch = seq_batch)
        # prox_consistency_loss = self._prox_consistency_loss(sc_syntax_vals, sc_L_vals)

        # sc_preds = self.learner.Txs_to_sc_pred(sc_syntax_vals[0,:,0,:],sc_batch) # zeroth neighbor for sc and syntax
        # sc_target_loss = self._sc_target_loss(sc_pred=sc_preds,sc_target=sc_targets)

        # seq_mean_est = sc_syntax_vals[0,:,0,:].mean(axis = 0)
        # seq_target_loss = torch.square(seq_mean_est - seq_targets).mean()

        # entropy = self.ent_reg(sc_syntax_vals=sc_syntax_vals)
        
        # loss_dict = {'sc_con':sc_consistency_loss, 'syn_con':syntax_consistency_loss, 'prox_con': prox_consistency_loss,'sc_targ': sc_target_loss, 'seq_targ':seq_target_loss, 'seq_entropy':entropy}
        
        # scxseq = sc_syntax_vals[0,:,0,:1000].detach()
        # if return_vals == True:
        #     return loss_dict, scxseq.std(axis = 1).mean(), scxseq.std(axis = 0).mean(), sc_syntax_vals[0,:,0,:]

        # return loss_dict, scxseq.std(axis = 1).mean(), scxseq.std(axis = 0).mean()

    def validate(self, seq_inds):
        self.learner.eval()
        with torch.no_grad():
            seq_batch = self.genome[seq_inds].to(self.device).float()
            sc_batch = self.val_sc.to(self.device)
            preds = self.learner.T_xs(seq_batch=seq_batch,sc_batch=sc_batch).mean(axis = -1)
            out = torch.square(preds - self.val_target.to(self.device)).mean()

        self.learner.train()
        return out
    


class GraphLaplacianPSA(GraphPSABase):
    '''
    minimizes intra-local neighborhood variance on sc/syntax/proximity
    '''
    def __init__(self, syntax_graph, genome_tensor, phenotype_graph, phenotype_tensor, sc_target, val_sc, val_target, learner,syntax_type, device='cpu:0', **learner_kwargs):
        super().__init__(syntax_graph, genome_tensor, phenotype_graph, phenotype_tensor, sc_target, val_sc, val_target, learner, device, **learner_kwargs)
        self.syntax_type = syntax_type

    def _sc_consistency_loss(self,sc_syntax_vals, sc_batch = None):
        sc_center= sc_syntax_vals[0,:,:,:]
        square_norm = torch.square(sc_center).sum(axis = 0)
        return (torch.square(sc_center[None,:,:,:] - sc_syntax_vals[1:,:,:,:])/(1e-5 + square_norm[None,None,:,:])).sum(axis = (0,1)).mean()
        ''' old
        sc_center = sc_syntax_vals[0:,:,:] #len(sc_inds) x  k_syn x len(seq_inds)
        sc_neighborhood_means = sc_syntax_vals[1:].mean(axis = 0) #len(sc_inds)x  k_syn x len(seq_inds)
        square_norm = torch.square(sc_center).mean(axis = (1,2),keepdim = True)
        return (torch.square((sc_neighborhood_means.detach() - sc_center))/(1e-3+square_norm)).mean(axis = (1,2)).mean()
        '''
        

    def _syntax_consistency_loss(self, sc_syntax_vals, detach = True,seq_batch = None):

        # '''
        # Strongest: that for any given cell, its associations are consistent with the graph - i.e. if it's associated with a locus,
        # then given the syntactic neighbors of that locus, the cell should also be associated with that locus
        # '''
        # seq_center = sc_syntax_vals[:,:,0,:] #T((sc), source)
        # square_norm = torch.square(seq_center).sum(axis = -1) #T(sc)
        # return (torch.square(seq_center[:,:,None,:] - sc_syntax_vals[:,:,1:])/(1e-5 + square_norm[:,:,None,None])).sum(axis = (2,3)).mean() # for each (sc), have laplacian discr. for it

        '''
        Intermediate
        this says that for any given sequence, its neighboring sequences are associated with similar cells
        intermediate one (e.g. run 596)
        '''
        
        if detach:
            seq_center = sc_syntax_vals[:,:,0,:].detach()
        else:
            seq_center = sc_syntax_vals[:,:,0,:]

        if self.syntax_type =='distance':
             #T(x,i, source)
            square_norm = torch.square(seq_center).mean(axis = (0,1), keepdim=True)
            sq_diff = torch.square(seq_center[:,:,None,:] - sc_syntax_vals[:,:,1:,:])/(1e-5 + square_norm[:,:,None,:])
            return sq_diff.sum(axis = (0,1)).mean(axis = 0).mean() #first sum across the scs, then average across neighbors, then average across all seq
        if self.syntax_type =='mean':
            square_norm = torch.square(seq_center).mean(axis = (0,1), keepdim=True)
            syntax_neighborhood_means = sc_syntax_vals[:,:,1:,:].mean(axis =2) # k_sc x sc_inds x num_seq
            return (torch.square((seq_center - syntax_neighborhood_means))/(1e-3+ square_norm)).mean(axis = (0,1)).mean()
        # if self.syntax_type =='pw_distance':
          # TODO: implement so that for each   
            # square_norm = torch.square(seq_center[0]).mean(axis = 1, keepdim=True)
            # txs_nb = sc_syntax_vals[0,:,:,:]/(1e-3+square_norm) #sc_b x 20 x seq_b
            # num_genome_neighbors = sc_syntax_vals.shape[2]
            # triu_inds = torch.triu_indices(num_genome_neighbors, num_genome_neighbors ,offset=1,device = self.device)
            # return torch.square(txs_nb[:,None,:,:]) -torch.square(txs_nb[:,:,None,:]).sum(axis = 0)[triu_inds].mean()

            #torch.square(txs_nb[None,:,:,:] - txs_nb[:,None,:,:])[triu_inds].sum(axis = 0)
        #     torch.triu_indices(batch_size,batch_size,offset=1,device = self.device)


        #     square_norm = torch.square(seq_center).mean(axis = (0,1), keepdim=True)
        #     syntax_neighborhood_means = sc_syntax_vals[:,:,1:,:].mean(axis =2) # k_sc x sc_inds x num_seq
        #     return (torch.square((seq_center - syntax_neighborhood_means))/(1e-3+ square_norm)).mean(axis = (0,1)).mean()  
        
        # '''
        # Weakest
        # this was the old one , which says that for any given sequence, its activations are close to the average across its neighbors
        # '''
        # # old: following lines takes the squared difference from the neighborhood average
        
        
        
        
        
        
        
    def _prox_consistency_loss(self, sc_syntax_vals, sc_L_vals):
        seq_center = sc_syntax_vals[:,:,0,:].detach()
        # abs_norm = torch.abs(seq_center).mean(axis = (0,1), keepdim=True) #average activations across all of the sc (shame to compute these multiple times)
        square_norm = torch.square(seq_center).mean(axis = (0,1), keepdim=True)
        prox_neighborhood_means = sc_L_vals.mean(axis = 2)
        return (torch.square((seq_center - prox_neighborhood_means))/(1e-3 + square_norm)).mean(axis = (0,1)).mean()


class GraphTemplatePSA(GraphLaplacianPSA):
    '''
    like GraphLaplacianPSA but matches cosine similarities between pairs of scs and their corresponding functionals
    '''
    def __init__(self, syntax_graph, genome_tensor, phenotype_graph, phenotype_tensor, sc_target, val_sc, val_target, learner, syntax_type = 'mean',pool_fun ='mean', device='cpu:0', **learner_kwargs):
        super().__init__(syntax_graph, genome_tensor, phenotype_graph, phenotype_tensor, sc_target, val_sc, val_target, learner, device = device,syntax_type = syntax_type, **learner_kwargs)

        if pool_fun =='mean':
            self.pool_fun = torch.mean
        if pool_fun =='max':
            self.pool_fun = torch.amax


    def _discrepancy(self, flat_txs, flat_batch,eps = 1e-10):
        orig_norm = flat_batch/(eps+torch.norm(flat_batch, p = 2, dim =1, keepdim=True))
        orig_dist = 1-torch.abs(torch.matmul(orig_norm,orig_norm.T)) #cosine distance
        
        batch_size =flat_txs.shape[0]
        row,col = torch.triu_indices(batch_size,batch_size,offset=1,device = self.device)
        txs_dist = torch.abs(flat_txs[row]-flat_txs[col]).mean(axis = -1) #L1 distance


        discrepancy = self.pool_fun(torch.abs(orig_dist[(row,col)] - txs_dist))
        return discrepancy

    def _sc_consistency_loss(self, sc_syntax_vals, sc_batch,eps = 1e-10):
        
        flat_batch = sc_batch[0]
        flat_txs = sc_syntax_vals[0,:,0,:]

        return self._discrepancy(flat_txs=flat_txs, flat_batch=flat_batch)
    

  

class MultiComponentGraphPSA():
    def __init__(self, n_components, psa_type, syntax_graph, genome_tensor, phenotype_graph, phenotype_tensor, sc_target, val_sc, val_target, learner, detach  = True, syntax_type = 'mean',pool_fun = 'mean', device='cpu:0', **learner_kwargs):
        self.detach = detach
        self.psa_list = []
        self.n_components = n_components
        for i in range(n_components):
            self.psa_list.append(psa_type(syntax_graph, genome_tensor, phenotype_graph, phenotype_tensor, sc_target, val_sc, val_target, detach = detach, learner=learner, pool_fun=pool_fun,syntax_type = syntax_type,device='cpu:0', **learner_kwargs))
        self.device = device
        self.learner = nn.ModuleList(psa.learner for psa in self.psa_list)

    def to(self, device='cpu:0'):
        for psa  in self.psa_list:
            psa.to(device) 
        self.device = device

    
    def loss(self, sc_inds, seq_inds, return_vals=False):
        psa0 = self.psa_list[0]
        sc_batch, seq_batch = psa0.extract_batches(sc_inds = sc_inds,seq_inds=seq_inds)
        sc_targets = psa0.sc_target[sc_inds].to(self.device)
        seq_targets = 0


        loss_dict = {'sc_con':[], 'syn_con':[], 'prox_con': [],'sc_targ': [], 'seq_targ':[], 'seq_entropy': []}
        Txs_list = []

        for psa in self.psa_list:
            sc_syntax_vals, sc_L_vals = psa.eval_neighbors(seq_batch = seq_batch, sc_batch = sc_batch, sc_inds = sc_inds, seq_inds=seq_inds)
            Txs_list.append(sc_syntax_vals[0,:,0,:])
            sc_con =psa._sc_consistency_loss( sc_syntax_vals = sc_syntax_vals,sc_batch = sc_batch)
            

            loss_dict['sc_con'].append(sc_con)
            loss_dict['syn_con'].append(psa._syntax_consistency_loss(sc_syntax_vals,seq_batch = seq_batch,detach = self.detach))
            loss_dict['prox_con'].append(psa._prox_consistency_loss(sc_syntax_vals, sc_L_vals))

            sc_preds = psa.learner.Txs_to_sc_pred(sc_syntax_vals[0,:,0,:],sc_batch) # zeroth neighbor for sc and syntax
            loss_dict['sc_targ'].append(psa._sc_target_loss(sc_pred=sc_preds,sc_target=sc_targets))

            # seq_mean_est = sc_syntax_vals[0,:,0,:].mean(axis = 0)
            # loss_dict['seq_targ'].append(torch.abs(seq_mean_est - seq_targets).mean())

            # change seq targ so that tis the mean absolute value, not the absolute mean value
            loss_dict['seq_targ'].append(torch.abs(sc_syntax_vals[0,:,0,:]).mean(axis = 0).mean())

            loss_dict['seq_entropy'].append( psa._seq_entropy(sc_syntax_vals))

            
        Txsl = torch.stack(Txs_list,dim = -1)
        nseq = Txsl.shape[1]
        nx = Txsl.shape[0]


        eye = torch.eye(1+self.n_components,device = self.device)
        
        Tsl = Txsl.sum(axis = 0)
        Tsl_norm = torch.cat([torch.ones(nseq,1,dtype = torch.float, device = self.device)/np.sqrt(nseq),
                              Tsl/(1e-6+torch.norm(Tsl, p = 2,dim =0,keepdim=True))], dim = 1) # normalize and concatenate unit constant vectors
        s_off_diag = torch.abs((1-eye)* torch.matmul(Tsl_norm.T, Tsl_norm))

        Txl = Txsl.sum(axis = 1)
        Txl_norm = torch.cat([torch.ones(nx,1,dtype = torch.float, device = self.device)/np.sqrt(nx),
                              Txl/(1e-6+torch.norm(Txl, p = 2,dim =0,keepdim=True))], dim = 1)
        x_off_diag = torch.abs((1-eye)* torch.matmul(Txl_norm.T, Txl_norm))
        
        #orthogonality loss
        
       
        loss_dict['ortho_loss'] = ( s_off_diag.mean() + x_off_diag.mean())

        return loss_dict,Txs_list

    def validate(self, seq_inds):
        with torch.no_grad():
            val_losses = []
            seq_batch = self.psa_list[0].genome[seq_inds].to(self.device).float()
            sc_batch = self.psa_list[0].val_sc.to(self.device)
            for psa in self.psa_list:
                psa.learner.eval()
                losses = []
                bs = min(512, len(sc_batch))
                num_its = max(1, bs/len(sc_batch))
                noised_losses = []
                for _ in range(num_its):
                    sc_inds = np.random.choice(np.arange(len(sc_batch)), size = bs, replace = False)
                    flat_sc_batch = sc_batch[sc_inds]
                    # flat_txs = psa.learner.T_xs(seq_batch=seq_batch,sc_batch=sc_batch[sc_inds]).mean(axis = -1) #why was this taking the  mean?
                    flat_txs = psa.learner.T_xs(seq_batch=seq_batch,sc_batch=sc_batch[sc_inds]) 
                    losses.append(psa._discrepancy(flat_txs=flat_txs, flat_batch=flat_sc_batch)) #this will break with laplcian since it requires discrepancy to be defined
                out = torch.mean(torch.stack(losses))
                val_losses.append(out)
                psa.learner.train()
        return val_losses
    
```
