## Supplementary material for "Deriving genetic codes for molecular phenotypes from first principles": Code: PSALearners.py

```
import torch
from torch import nn
import numpy as np

class PSALearnerBase(nn.Module):
    def __init__(self) -> None:
        super(PSALearnerBase, self).__init__()

    def T_xs(self, sc_batch, seq_batch):
        '''
        override to output a matrix with value of function T(x,s)
        '''
        pass

    
    def Txs_to_sc_pred(self, txs,sc_batch):
        ''' returns the predicted regression value from functionals by averaging over s
        '''
        return txs.mean(axis = -1)


class LatentPSALearner(PSALearnerBase):
    '''
    uses distances in sc space as weights
    '''
    def __init__(self, **kwargs):
        sc = kwargs['sc']
        genome = kwargs['genome']
        n_control = kwargs['n_control']
        if 'n_control' not in kwargs:
            n_control = 5
        
        bias = False
        bn_layer = nn.Identity()
        sc_norm_layer = nn.Identity()
        self.shift = 0

        if 'bias' in kwargs:
            if kwargs['bias'] ==True:
                bias = True
        if 'shift' in kwargs:
            self.shift = kwargs['shift']

        if 'batch_norm' in kwargs:
            if kwargs['batch_norm'] == True:
                bn_layer = nn.BatchNorm1d(n_control)

        if 'layer_norm' in kwargs:
            if kwargs['layer_norm'] == True:
                bn_layer = nn.LayerNorm(n_control)

        if 'softmax' in kwargs:
            if kwargs['softmax']==True:
                sc_norm_layer =nn.Softmax(dim = 1)
        


        
        super(LatentPSALearner, self).__init__()
        self.sc_q = nn.Sequential(nn.Dropout(p = kwargs['dropout']), nn.Linear(sc.shape[1], n_control, bias = False),sc_norm_layer) #removed dropout,add batch norm as specified by kwargs
        self.seq_k= nn.Sequential(nn.Linear(genome.shape[1], n_control,bias = bias),bn_layer) 
        self._nonlinearity = torch.tanh

    
    def T_xs(self, sc_batch,seq_batch):
        # sc_probs = torch.softmax(self.sc_q.forward(sc_batch), dim = 1) #sc_batch x n_control
        sc_probs = self.sc_q.forward(sc_batch) #SOFTMAX specified by kwarg 
        seq_k = self.seq_k(seq_batch) #seq_batch x n_control
        txs_before_nonlinearity  = torch.matmul(sc_probs,seq_k.T) #sc x seq
        # txs = (1+torch.abs(self.scale))*(1+torch.tanh(txs_before_nonlinearity))
        # txs = (1+torch.tanh(txs_before_nonlinearity))
        # txs = (1+torch.tanh(txs_before_nonlinearity))
        txs = self.shift + self._nonlinearity(txs_before_nonlinearity)
        return txs
    
```
