## Supplementary material for "Deriving genetic codes for molecular phenotypes from first principles": Code: corrupt_pbmc_data_prep.ipynb

```
# %% [code]
from scanpy import read_h5ad
import numpy as np
import pandas as pd
import pickle 
from tqdm.notebook import tqdm
import torch
from torch import nn


import os
from matplotlib import pyplot as plt
%pylab inline
import seaborn as sns

from numpy.lib.stride_tricks import sliding_window_view
from tqdm.notebook import tqdm


from scipy.stats import mannwhitneyu,ttest_ind
from scipy.stats import t as tdist
from statsmodels.stats.multitest import fdrcorrection
from sklearn.metrics import roc_curve,auc


from models.PSALearners import LatentPSALearner


# %% [markdown]
# Generating the corrupt datasets

# %% [code]
graph = np.load('../data/sc/graphs/metacell_pbmc_train_graph.npy')
train_inds= np.load('../data/sc/graphs/metacell_pbmc_train_inds.npy')
train_features = np.load('../data/sc/graphs/metacell_pbmc_train_features.npy')
val_features = np.load('../data/sc/graphs/metacell_pbmc_val_features.npy')


# %% [code]
import os

# %% [code]
path = '../data/sc/graphs/nature_corrupt_metacell_pbmc'

os.mkdir(path)

# %% [code]
vl = pd.read_pickle('../data/downstream/pbmc_valid_long')
GENES = sorted(vl['GENE'].unique())
sc = pd.read_pickle('../data/sc/processed/pbmc_SEACELL_means')
gene_inds_map = {j:i for i,j in enumerate(sc.columns)}
permutation_inds = [gene_inds_map[j] for j in GENES]
for i,gene in zip(permutation_inds, GENES):
    assert sc.columns[i]==gene 

# %% [code]
np.random.seed(24)
for shuf, i in tqdm(enumerate(permutation_inds)):
    perm_train_feats = train_features.copy()
    perm_val_feats = val_features.copy()
    np.random.shuffle(perm_train_feats[:,i])
    np.random.shuffle(perm_val_feats[:,i])
    np.save(os.path.join(path, 'single_{}_train_features'.format(sc.columns[i])), perm_train_feats)
    np.save(os.path.join(path, 'single_{}_val_features'.format(sc.columns[i])), perm_val_feats)
    np.save(os.path.join(path, 'single_{}_train_graph'.format(sc.columns[i])), graph)
    np.save(os.path.join(path, 'single_{}_train_inds'.format(sc.columns[i])), train_inds)


# %% [code]
files = [path+'/'+'_'.join(a.split('_')[:-2]) for a in os.listdir(path) if 'train_features' in a]

# %% [code]
os.mkdir('results/sep29')

# %% [code]
COMMAND = "python pbmc_corruption.py --device {} --out_dir results/sep29/corruption/{} --num_repeat 1 --save_every 4 --num_epochs 100 --seq_batch_size 100000 --psa_type template --learner_type latent --num_component 1 --ortho_weight 0 --n_control 40 --precomputed_genome_prefix ../data/seq/GENA_2k_precomputed --sc_batch_size 512 --ignore_density --softmax --overwrite True --seq_target_weight 0.01 --manual_seed 125 --sc_W_init ../data/sc/inits/averaged_PBMC_pca_init --seq_W_init ../data/sc/inits/averaged_GENA_pca_init --gradient_accumulation_steps 1 --dropout 0 --sc_prefix {}"

file_lists = {}
for i,f in enumerate(files):
    file_lists.setdefault(i%8,[]).append(COMMAND.format(i%8,f.split('/')[-1],f))

# %% [code]
for i in range(8):
    print ('if [ "$1" = {} ]\nthen'.format(i))
    print('\t'+'\n\n\t'.join(sorted(file_lists[i])))
    print('fi\n\n')

# %% [code]
print('\n\n'.join(np.concatenate([sorted(file_lists[i]) for i in range(8)])))

# %% [code]
print('(' + ' ' .join(().astype(str))  + ')')

# %% [code]
```
