## Supplementary material for "Deriving genetic codes for molecular phenotypes from first principles": Code: metacells_pbmc.ipynb

```
# %% [code]
!pip install SEACells

# %% [code]
from scanpy import read_h5ad
import numpy as np
import pandas as pd
import pickle 
from tqdm.notebook import tqdm
import torch
from torch import nn


import os
from matplotlib import pyplot as plt
%pylab inline
import seaborn as sns

from numpy.lib.stride_tricks import sliding_window_view
from tqdm.notebook import tqdm


from scipy.stats import mannwhitneyu,ttest_ind
from scipy.stats import t as tdist
from statsmodels.stats.multitest import fdrcorrection
from sklearn.metrics import roc_curve,auc


from models.PSALearners import LatentPSALearner

import SEACells

# %% [code]
x =read_h5ad('../data/sc/raw/pbmc_multimodal.h5ad')
sc_df = x.to_df()
ann = x.obs
t0 =sc_df.loc[x.obs['time']=='0']
cells = t0.loc[x.obs['donor'].loc[t0.index.values].isin(['P1','P3','P5'])]/1000

# %% [code]
pcs.shape

# %% [code]
sc.shape

# %% [code]
torch.cuda.set_device(0)
with torch.no_grad():
    sc = torch.from_numpy(cells.values).cuda()
    pcs = torch.pca_lowrank(sc, q = 200)[2]
    pc_df = pd.DataFrame(pcs.detach().cpu().numpy(), index = cells.columns)
    pc_cells = pd.DataFrame(torch.matmul(sc - sc.mean(axis = 0,keepdim = True), pcs).detach().cpu().numpy(), index = cells.index.values)

# %% [code]


# %% [code]
from scanpy import AnnData

# %% [code]
data = AnnData(X= cells.values,obs=ann.loc[cells.index.values].values,var= x.var,varm={'PCs':pc_df.values}, obsm = {'PC':pc_cells.values})

# %% [code]
model = SEACells.core.SEACells(data, 
                  build_kernel_on='PC',
                      n_SEACells=200, 
                  n_waypoint_eigs=20,
                  convergence_epsilon = 1e-5)


# %% [code]


model.construct_kernel_matrix()

# %% [code]
model.n_waypoint_eigs = 15

# %% [code]


# Initialize archetypes
model.initialize_archetypes()


# %% [code]
model.fit(min_iter=10, max_iter=50)

# %% [code]
data.obs[['SEACell']]

# %% [code]
pd.DataFrame(data.obs.values,index = cells.index.values).to_pickle('../data/sc/processed/pbmc_SEACell_assignments')

# %% [code]
data

# %% [code]
seacell_means = cells.groupby(data.obs['SEACell'].values).mean()

# %% [code]
seacell_means.to_pickle('../data/sc/processed/pbmc_SEACELL_means')
```
