## Supplementary material for "Deriving genetic codes for molecular phenotypes from first principles": Code: metacells_tabulasap.ipynb

```
# %% [code]
from scanpy import read_h5ad
import numpy as np
import pandas as pd
import pickle 
from tqdm.notebook import tqdm
import torch
from torch import nn


import os
from matplotlib import pyplot as plt
%pylab inline
import seaborn as sns

from numpy.lib.stride_tricks import sliding_window_view
from tqdm.notebook import tqdm


from scipy.stats import mannwhitneyu,ttest_ind
from scipy.stats import t as tdist
from statsmodels.stats.multitest import fdrcorrection
from sklearn.metrics import roc_curve,auc


from models.PSALearners import LatentPSALearner

import SEACells

# %% [code]
x =read_h5ad('../data/sc/raw/TabulaSapiens.h5ad')

# %% [code]
inds_10x = np.where(x.obs['method']=='10X')[0]
np.random.seed(23)
np.random.shuffle(inds_10x)

# %% [code]
samp = x[inds_10x[:100000]]

# %% [code]
sc_df = samp.to_df()

# %% [code]
sc_df.shape

# %% [code]
cells = sc_df

# %% [code]
torch.cuda.set_device(7)
with torch.no_grad():
    sc = torch.log(1+torch.from_numpy(cells.values).cuda())
    pcs = torch.pca_lowrank(sc, q = 200)[2]
    pc_df = pd.DataFrame(pcs.detach().cpu().numpy(), index = cells.columns)
    pc_cells = pd.DataFrame(torch.matmul(sc - sc.mean(axis = 0,keepdim = True), pcs).detach().cpu().numpy(), index = cells.index.values)

# %% [code]


# %% [code]
from scanpy import AnnData

# %% [code]
data = AnnData(X= cells.values,obs=x.obs.loc[cells.index.values].values,var= x.var,varm={'PCs':pc_df.values}, obsm = {'PC':pc_cells.values})

# %% [code]
model = SEACells.core.SEACells(data, 
                  build_kernel_on='PC',
                      n_SEACells=400, 
                  n_waypoint_eigs=15,
                  convergence_epsilon = 1e-5)


# %% [code]
model.construct_kernel_matrix()

# %% [code]
model.n_waypoint_eigs = 15

# %% [code]


# Initialize archetypes
model.initialize_archetypes()


# %% [code]
model.fit(min_iter=10, max_iter=50)

# %% [code]
seacell_means = np.log(1+cells).groupby(data.obs['SEACell'].values).mean()

# %% [code]
pd.DataFrame(data.obs.values,index = cells.index.values).to_pickle('../data/sc/processed/tabsap_SEACell_assignments')

# %% [code]


# %% [code]
print('hi')

# %% [code]
seacell_means

# %% [code]


# %% [code]
means = seacell_means.mean(axis =0)
good_genes = set(seacell_means.columns) - set([a for a in seacell_means.columns if ((('MT-' in a) or ('MIR' in a) or ('SNO' in a) or ('LINC' in a))or (means[a] <1e-5 ) or ('.' in a))])


# %% [code]
good_gene_inds  = [i for i,j in enumerate(seacell_means.columns) if j in good_genes]


# %% [code]
seacell_means.iloc[:,good_gene_inds]

# %% [code]
pcs.shape

# %% [code]
torch.save({'vt':pcs[good_gene_inds]}, '../data/sc/inits/tabsap_pca_init')

# %% [code]
seacell_means.iloc[:,good_gene_inds].to_pickle('../data/sc/processed/tabsap_SEACELL_means')

# %% [code]
x.var.to_pickle('../data/sc/processed/tabsap_gene_info')

# %% [code]
```
