## Supplementary material for "Deriving genetic codes for molecular phenotypes from first principles": Code: sc_data_prep_metacells_pbmc.ipynb

```
# %% [code]
from scanpy import read_h5ad
import numpy as np
import pandas as pd
import pickle 
from tqdm.notebook import tqdm
import torch
from torch import nn


import os
from matplotlib import pyplot as plt
%pylab inline
import seaborn as sns

from numpy.lib.stride_tricks import sliding_window_view
from tqdm.notebook import tqdm


from scipy.stats import mannwhitneyu,ttest_ind
from scipy.stats import t as tdist
from statsmodels.stats.multitest import fdrcorrection
from sklearn.metrics import roc_curve,auc


from models.PSALearners import LatentPSALearner


# %% [code]
asn = pd.read_pickle('../data/sc/processed/pbmc_SEACell_assignments')

counts = asn.groupby(9).apply(lambda w: w[14].value_counts(normalize = True)).unstack().fillna(0).sort_index().T

# %% [code]


# %% [code]
sc = pd.read_pickle('../data/sc/processed/pbmc_SEACELL_means').sort_index()

# %% [code]
from scipy.spatial.distance import cdist
from sklearn.neighbors import kneighbors_graph
from scipy.sparse.csgraph import shortest_path

# %% [code]
sns.clustermap(shortest_path(kneighbors_graph(sc,metric = 'cosine', n_neighbors= 5,mode = 'connectivity'),directed= False), cmap = 'Reds')

# %% [code]
f = sns.clustermap(cdist(sc,sc,metric = 'cosine'),cmap = 'Reds')

# %% [code]
from sklearn.neighbors import kneighbors_graph

# %% [code]
idx = np.arange(200)
np.random.seed(23)
np.random.shuffle(idx)
train_inds = idx[:-20]
val_inds = idx[-20:]

# %% [code]
features = sc.iloc[idx].values

# %% [code]

graph = kneighbors_graph(features,n_neighbors = 3,metric = 'cosine',include_self=True).nonzero()[1].reshape(-1,3).T

# %% [code]
train_inds = np.arange(200)[graph.max(axis = 0)<180]

# %% [code]
val_inds = np.sort(list(set(list(range(200))) - set(train_inds.tolist())))

# %% [code]

train_features = features[:-20]
val_features = features[-20:]


# %% [code]
np.save('../data/sc/graphs/metacell_pbmc_train_graph',graph.astype(np.int64))
np.save('../data/sc/graphs/metacell_pbmc_train_inds',train_inds)
np.save('../data/sc/graphs/metacell_pbmc_train_features', train_features)
np.save('../data/sc/graphs/metacell_pbmc_val_features', val_features)


# %% [code]
sns.clustermap(counts, row_linkage=f.dendrogram_row.linkage)

# %% [markdown]
# Generating the corrupt datasets

# %% [code]
np.save('../data/sc/graphs/metacell_pbmc_train_graph',graph.astype(np.int64))
np.save('../data/sc/graphs/metacell_pbmc_train_inds',train_inds)
np.save('../data/sc/graphs/metacell_pbmc_train_features', train_features)
np.save('../data/sc/graphs/metacell_pbmc_val_features', val_features)


# %% [code]
import os

# %% [code]
path = '../data/sc/graphs/corrupt_metacell_pbmc'

os.mkdir(path)

# %% [code]
GENES = ['USP36', 'SH3GLB1', 'PDS5B', 'USP25', 'ZNF800', 'UBA3', 'TOMM7','PREP', 'SLC4A7', 'SLMAP', 'QDPR', 'MYLIP', 'SYNE1', 'RRAS2','CHD6', 'YEATS4', 'PTPRE', 'YWHAE', 'UTRN', 'ZC3H7A']
permutation_inds = [13628,715,10623,16028,6275,3145,5838,5467,2842,3110,3735,4958,5659,8335,14196,10196,9625,12728,5623,12119]
for i,gene in zip(permutation_inds, GENES):
    assert sc.columns[i]==gene 

# %% [code]
gr = graph.astype(np.int64)

# %% [code]
perm_train_feats = train_features.copy()
perm_val_feats = val_features.copy()
np.random.seed(24)
for shuf in tqdm(range(5)):
    for i in permutation_inds:
        np.random.shuffle(perm_train_feats[:,i])
        np.random.shuffle(perm_val_feats[:,i])
    
    np.save(os.path.join(path, 'all_shuf_{}_train_features'.format(shuf)), perm_train_feats)
    np.save(os.path.join(path, 'all_shuf_{}_val_features'.format(shuf)), perm_val_feats)
    np.save(os.path.join(path, 'all_shuf_{}_train_graph'.format(shuf)), gr)
    np.save(os.path.join(path, 'all_shuf_{}_train_inds'.format(shuf)), train_inds)


# %% [code]
np.random.seed(24)
for shuf, i in tqdm(enumerate(permutation_inds)):
    perm_train_feats = train_features.copy()
    perm_val_feats = val_features.copy()
    np.random.shuffle(perm_train_feats[:,i])
    np.random.shuffle(perm_val_feats[:,i])
    np.save(os.path.join(path, 'single_{}_train_features'.format(shuf)), perm_train_feats)
    np.save(os.path.join(path, 'single_{}_val_features'.format(shuf)), perm_val_feats)
    np.save(os.path.join(path, 'single_{}_train_graph'.format(shuf)), gr)
    np.save(os.path.join(path, 'single_{}_train_inds'.format(shuf)), train_inds)


# %% [code]
files = [path+'/'+'_'.join(a.split('_')[:-2]) for a in os.listdir(path) if 'train_features' in a]

# %% [code]
COMMAND = "python pbmc_corruption.py --device {} --out_dir results/mar31/corruption/{} --num_repeat 1 --save_every 2 --num_epochs 200 --seq_batch_size 100000 --psa_type template --learner_type latent --learner_bias False --num_component 1 --ortho_weight 0 --n_control 40 --precomputed_genome_prefix ../data/seq/GENA_2k_precomputed --sc_batch_size 512 --ignore_density --softmax --overwrite True --seq_target_weight 0.01 --manual_seed 125 --sc_W_init ../data/sc/inits/averaged_PBMC_pca_init --seq_W_init ../data/sc/inits/averaged_GENA_pca_init --gradient_accumulation_steps 1 --dropout 0 --sc_prefix {}"

file_lists = {}
for i,f in enumerate(files):
    
    file_lists.setdefault(i%8,[]).append(COMMAND.format(i%8,f.split('/')[-1],f))

# %% [code]
for i in range(8):
    print ('if [ "$1" = {} ]\nthen'.format(i))
    print('\t'+'\n\n\t'.join(sorted(file_lists[i])))
    print('fi\n\n')

# %% [code]
print('\n\n'.join(np.concatenate([sorted(file_lists[i]) for i in range(8)])))

# %% [code]
print('(' + ' ' .join(().astype(str))  + ')')

# %% [code]
```
