## Supplementary material for "Deriving genetic codes for molecular phenotypes from first principles": Code: sc_data_prep_metacells_pbmc_replicates.ipynb

```
# %% [code]
from scanpy import read_h5ad
import numpy as np
import pandas as pd
import pickle 
from tqdm.notebook import tqdm
import torch
from torch import nn


import os
from matplotlib import pyplot as plt
%pylab inline
import seaborn as sns

from numpy.lib.stride_tricks import sliding_window_view
from tqdm.notebook import tqdm


from scipy.stats import mannwhitneyu,ttest_ind
from scipy.stats import t as tdist
from statsmodels.stats.multitest import fdrcorrection
from sklearn.metrics import roc_curve,auc


from models.PSALearners import LatentPSALearner


# %% [code]
asn = pd.read_pickle('../data/sc/processed/pbmc2_SEACell_assignments')

# %% [code]
counts = asn.groupby(9).apply(lambda w: w[14].value_counts(normalize = True)).unstack().fillna(0).sort_index().T

# %% [code]
sc = pd.read_pickle('../data/sc/processed/pbmc2_SEACELL_means').sort_index()

# %% [code]
from scipy.spatial.distance import cdist

# %% [code]
f = sns.clustermap(cdist(sc,sc,metric = 'cosine'),cmap = 'Reds',vmax = 0.2)

# %% [code]
from sklearn.neighbors import kneighbors_graph

# %% [code]
idx = np.arange(200)
np.random.seed(23)
np.random.shuffle(idx)
train_inds = idx[:-20]
val_inds = idx[-20:]
features = sc.iloc[idx].values

graph = kneighbors_graph(features,n_neighbors = 3,metric = 'cosine',include_self=True).nonzero()[1].reshape(-1,3).T
train_inds = np.arange(200)[graph.max(axis = 0)<180]
val_inds = np.sort(list(set(list(range(200))) - set(train_inds.tolist())))

train_features = features[:-20]
val_features = features[-20:]

np.save('../data/sc/graphs/metacell_pbmc2_train_graph',graph.astype(np.int64))
np.save('../data/sc/graphs/metacell_pbmc2_train_inds',train_inds)
np.save('../data/sc/graphs/metacell_pbmc2_train_features', train_features)
np.save('../data/sc/graphs/metacell_pbmc2_val_features', val_features)


# %% [code]


# %% [code]


# %% [code]


# %% [code]


# %% [code]


# %% [code]
sns.clustermap(counts, row_linkage=f.dendrogram_row.linkage)

# %% [code]
```
