## Supplementary material for "Deriving genetic codes for molecular phenotypes from first principles": Code: sc_data_prep_tabsap.ipynb

```
# %% [code]
import numpy as np
import pandas as pd
from scanpy import read_h5ad
import torch
from matplotlib import pyplot as plt
from tqdm.notebook import tqdm
from sklearn.neighbors import NearestNeighbors
from scipy.spatial.distance import cdist
import seaborn as sns
from scipy.cluster.hierarchy import fcluster
from sklearn.neighbors import kneighbors_graph

# %% [code]
anns = pd.read_pickle('../data/sc/processed/tabsap_SEACell_assignments')
sc = pd.read_pickle('../data/sc/processed/tabsap_SEACELL_means')


# %% [code]
sc.max(axis = 0).sort_values()/1000

# %% [code]


# %% [code]
idx = np.arange(400)
np.random.seed(2382)
np.random.shuffle(idx)
num_val = 50
train_inds = idx[:-num_val]
val_inds = idx[-num_val:]
features = sc.iloc[idx].values/1000

graph = kneighbors_graph(features,n_neighbors = 3,metric = 'cosine',include_self=True).nonzero()[1].reshape(-1,3).T
train_inds = np.arange(len(idx))[graph.max(axis = 0)<(len(idx)-num_val)]
val_inds = np.sort(list(set(list(range(400))) - set(train_inds.tolist())))

train_features = features[:-num_val]
val_features = features[-num_val:]

np.save('../data/sc/graphs/metacell_tabsap_train_graph',graph.astype(np.int64))
np.save('../data/sc/graphs/metacell_tabsap_train_inds',train_inds)
np.save('../data/sc/graphs/metacell_tabsap_train_features', train_features)
np.save('../data/sc/graphs/metacell_tabsap_val_features', val_features)


# %% [code]
tissue_counts = anns.groupby([11,0]).size().unstack().fillna(0)
tissue_dists = pd.DataFrame(tissue_counts.values/tissue_counts.values.sum(axis = 0,keepdims=True), index = tissue_counts.index, columns = tissue_counts.columns)


# %% [code]
tissue_genex_means = pd.DataFrame(np.matmul(tissue_dists.values.T, sc.values), index = tissue_dists.columns, columns = sc.columns)

# %% [code]
tissue_genex_means.max(axis = 0).sort_values()

# %% [code]
dists = cdist(sc,sc, metric = 'cosine')

# %% [code]
sns.clustermap(dists,cmap = 'Reds')

# %% [code]
!wget https://raw.githubusercontent.com/eQTL-Catalogue/eQTL-Catalogue-resources/master/tabix/tabix_ftp_paths.tsv

# %% [code]
catalog = pd.read_table('tabix_ftp_paths.tsv.1',sep = '\t')
query = catalog[(catalog['study_label']=='GTEx') & catalog['tissue_label'].str.upper().isin(tissues) & (catalog['quant_method'] == 'ge')]
commands = []
for i in range(len(query)):
    tissue = query['tissue_label'].iloc[i].upper()
    dataset_id = query['dataset_id'].iloc[i]
    commands.append('wget ftp://ftp.ebi.ac.uk/pub/databases/spot/eQTL/susie/QTS000015/{}/{}.credible_sets.tsv.gz -O {}.tsv.gz'.format(dataset_id, dataset_id,tissue))
print('\n'.join(commands)) 

    

# %% [code]
tissues =['LIVER',
 'LUNG',
 'MUSCLE',
 'PANCREAS',
 'PROSTATE',
 'SKIN',
 'SPLEEN',
 'UTERUS']

# %% [code]
dfs =[]
for tis in tissues:
    df = pd.read_table('../data/downstream/eqtl_database/{}.tsv'.format(tis))
    df['TISSUE'] = tis
    dfs.append(df)
df = pd.concat(dfs,axis = 0,ignore_index = True)

# %% [code]
dbsnp = pd.read_table('chm13v2.0_GWASv1.0rsids_e100_r2022-03-08.vcf', header = None,sep = '\t', comment='#')

# %% [code]
lifted_ctgt = df[df['rsid'].isin(dbsnp[2])]
t2t = pd.merge(df, dbsnp[[0,1,2]].rename({2:'rsid'},axis = 1),on = 'rsid').rename({0:'t2t_chr',1:'t2t_pos'},axis = 1)
t2t['t2t_ind'] = t2t['t2t_pos']//1000


# %% [code]
gene_info = pd.read_pickle('../data/sc/processed/tabsap_gene_info').loc[sc.columns]
gene_info['ensembl_split'] = [a.split('.')[0] for a in gene_info['ensemblid']]
good_genes = gene_info['ensembl_split'][~gene_info['ensemblid'].str.contains('PAR')]
gene_info = gene_info[gene_info['ensembl_split'].isin(good_genes)].set_index('ensembl_split')

# %% [code]
t2t = t2t[t2t['gene_id'].isin(good_genes)]

# %% [code]
t2t['GENE'] = gene_info['gene_symbol'].loc[t2t['gene_id'].values].values.astype(str)

# %% [code]


# %% [code]
genome = np.load('../data/seq/GENA_2k_precomputed/concatenated_genome.npy')
genome_tensor = (torch.from_numpy(genome)/10).float()
genome_mean = genome_tensor.mean(axis = 0).to('cuda:{}'.format(DEVICE))

# %% [code]
t2t.sort_values('pvalue')

# %% [code]
pos_df = t2t.groupby(['t2t_chr','t2t_ind','GENE','TISSUE'])['cs_size'].min().reset_index()

# %% [code]
pos_df

# %% [code]

valid_long = pos_df[pos_df['t2t_chr']!='chrY']
good_tg_pairs = (valid_long.groupby(['TISSUE','GENE'])['cs_size'].max()>20) & (valid_long.groupby(['TISSUE','GENE'])['cs_size'].min()<20)
gtg = valid_long.groupby(['TISSUE','GENE']).size()>3
good_tissue_gene_pairs = set(good_tg_pairs[good_tg_pairs].index.values).intersection(gtg[gtg].index.values)

valid_long = valid_long.loc[[i for i,a,b in zip(valid_long.index.values, valid_long['TISSUE'], valid_long['GENE']) if (a,b) in good_tissue_gene_pairs]]
valid_long = valid_long[valid_long['GENE'].isin(sc.columns)]
print(valid_long['GENE'].unique())

# %% [code]
unique_pos = valid_long[['t2t_chr','t2t_ind']].drop_duplicates()
chrom_inds = unique_pos.groupby('t2t_chr').groups
cis_seq_batches = []
dat_inds =[]
for chrom,inds in chrom_inds.items():
    chm13_pos = unique_pos['t2t_ind'].loc[inds]
    chr_dat = np.load('../data/seq/GENA_2k_precomputed/{}_precomputed_features.npy'.format(chrom), mmap_mode = 'r')
    cis_seq_batches.append(chr_dat[chm13_pos.values]/10)
    dat_inds+= list(inds)

cis_seq_batch = torch.tensor(np.concatenate(cis_seq_batches)).float().to('cuda:{}'.format(DEVICE))

qtl_to_seq_map = {ch_ind:i for i, ch_ind in enumerate (zip(unique_pos['t2t_chr'].loc[dat_inds],unique_pos['t2t_ind'].loc[dat_inds]))}


ct_gene_groups = valid_long.groupby(['TISSUE','GENE']).groups
cis_seq_inds = {}
pips = {}
for key,inds in ct_gene_groups.items():
    CELLTYPE,GENE = key
    cis_seq_inds[CELLTYPE, GENE] = np.array([qtl_to_seq_map[a] for a in zip(valid_long['t2t_chr'].loc[inds],valid_long['t2t_ind'].loc[inds])])
    pips[CELLTYPE,GENE] = valid_long['cs_size'].loc[inds]

# %% [code]
def prepare_sc_batch(ct, gene_to_add, amount = 0.0001):
    ct_mean = renamed_susy.loc[ct]
    edited_mean = ct_mean.copy()
    edited_mean[gene_to_add]+=amount
    return torch.tensor(np.stack([ct_mean.values.copy(), edited_mean.values]),device = 'cuda:{}'.format(DEVICE),dtype = torch.float)

# %% [code]
renamed_susy = tissue_genex_means.set_index(tissue_genex_means.index.str.upper()).loc[tissues]/1000

# %% [code]
import torch
from torch import nn
from models.PSALearners import LatentPSALearner
from sklearn.metrics import roc_auc_score
from scipy.stats import spearmanr
DEVICE = 0

# %% [code]

path  = 'results/apr7/tabsap_meansyntax_hilr/repeat_0_epoch_20.pt'.format(pt)
softmax = True
bn = False
n_learn = 1
n_control = 40
dropout= 0
softmax = True
bn = False
multi_learner = nn.ModuleList([LatentPSALearner(sc = torch.from_numpy(renamed_susy.values),dropout=dropout, softmax=softmax, genome = cis_seq_batch, n_control = n_control, bias = False,batch_norm = bn) for _ in range(n_learn)]).to('cuda:{}'.format(DEVICE))

checkpoint = torch.load(path)
sd = checkpoint['learner_state_dict']
multi_learner.load_state_dict(sd)
multi_learner.eval();
learner = multi_learner[0]
learner.eval();


# initialize maps that will be shuffled during permutations
orig_genes = valid_long['GENE'].unique()
shuf_genes = orig_genes.copy()
orig_cts = renamed_susy.index.values
shuf_cts = orig_cts.copy()
shuf_idx = np.arange(len(cis_seq_batch))

# want to enable shuffling only genes that are expressed in the same set of cell types, hence this shuffle
ct_set_to_g = {}
for g, ct_set in {g: tuple(sorted(renamed_susy[g][renamed_susy[g]>1e-3].index.values)) for g in orig_genes}.items():
    ct_set_to_g.setdefault(ct_set, []).append(g)

orig_group_genes = [np.array(gs) for _,gs in ct_set_to_g.items()]
shuf_group_genes = [arr.copy() for arr in orig_group_genes]
print([len(a) for a in orig_group_genes])


# select which kind of permutation
perm_gene = False
perm_ct= False
perm_rs = True
group_perm_gene = False
group_perm_rs = False
rand_init = False

assert (group_perm_gene&perm_gene)==False
assert (group_perm_rs&perm_rs)==False


nperm = 1000
perm_stats = {}
PRINT_EVERY = min(nperm,20)
np.random.seed(45)

for perm in tqdm(range(1+nperm)):
    # first permutation is always test statistic (i.e., no shuffle)
    if perm>0:
        if group_perm_gene:
            for arr in shuf_group_genes:
                np.random.shuffle(arr) #only shuffle within groups
        if group_perm_rs:
            for arr in shuf_group_rs:
                np.random.shuffle(arr) #only shuffle within groups
        if perm_gene:
            np.random.shuffle(shuf_genes)

        if perm_ct:
            np.random.shuffle(orig_cts)

        if perm_rs:
            np.random.shuffle(shuf_idx)

        if rand_init:
            multi_learner = nn.ModuleList([LatentPSALearner(sc = torch.from_numpy(renamed_susy.values),dropout=dropout, softmax=softmax, genome = cis_seq_batch, n_control = n_control, bias = False,batch_norm = bn) for _ in range(n_learn)]).to('cuda:{}'.format(DEVICE))
            multi_learner.eval();
            learner = multi_learner[0]
            learner.eval();


    # if perm==0, shuf_gene_map, shuf_idx, shuf_ct_map are all identity
    if group_perm_gene:
        shuf_gene_map = {i:j for i,j in zip(np.concatenate(orig_group_genes), np.concatenate(shuf_group_genes))}
    else:
        shuf_gene_map = {i:j for i,j in zip(orig_genes,shuf_genes)}


    if group_perm_rs:
        shuf_idx = np.concatenate(shuf_group_rs)
    shuf_ct_map = {i:j for i,j in zip(orig_cts,shuf_cts)}


    # compute AUCs for the discrepancy
    for CELLTYPE, GENE in ct_gene_groups.keys():
        sub_seq = cis_seq_batch[shuf_idx][cis_seq_inds[CELLTYPE,GENE]]


        shuf_ct = shuf_ct_map[CELLTYPE]
        shuf_gene = shuf_gene_map[GENE]
        sc_batch = prepare_sc_batch(shuf_ct, shuf_gene,amount=1e-4)

        Txs =  learner.T_xs(seq_batch= sub_seq, sc_batch = sc_batch)
        y_pred = torch.abs(Txs[1]-Txs[0]).detach().cpu().numpy()

        y_true = (pips[CELLTYPE,GENE]<20).astype(int).values
        perm_stats[perm,CELLTYPE,GENE] = roc_auc_score(y_true= y_true, y_score = y_pred)

#         y_true = -pips[CELLTYPE,GENE]
#         perm_stats[perm,CELLTYPE,GENE] = spearmanr(y_pred,y_true)[0]


    # occasionally print
    if (1+perm)%PRINT_EVERY==0:
        res = pd.Series(perm_stats).reset_index()
        res.columns = ['perm', 'ct', 'gene', 'auc']
        avgs = res.groupby('perm').apply(lambda w: w.groupby('gene')['auc'].mean()).mean(axis = 1)
        print('pval:', perm, (avgs[0]>avgs.iloc[1:]).mean(), 'test_stat:', avgs[0], 'perm mean:',avgs[1:].mean())


# %% [code]
pd.Series(ptres).sort_values()

# %% [code]
pd.Series(ptres).plot()

# %% [code]


# %% [code]


# %% [code]


# %% [code]


# %% [code]


# %% [code]


# %% [code]


# %% [code]


# %% [code]


# %% [code]
(renamed/1000).to_pickle('../data/downstream/tabsap_renamed')
valid_long.to_pickle('../data/downstream/tabsap_valid_long')

# %% [code]


# %% [code]


# %% [code]


# %% [code]


# %% [code]
from liftover import get_lifter
converter = get_lifter('hg38','GCA_009914755.4')

# %% [code]
eqtls =pd.read_table('GTEx_v8_finemapping_CaVEMaN/GTEx_v8_finemapping_CaVEMaN.txt')

# %% [code]
t2ts = [converter[chrom[3:]][pos] for chrom, pos in zip(eqtls['CHROM'],eqtls['POS'])]

# %% [code]
CP_chroms = ['CP068277.2',
'CP068276.2',
'CP068275.2',
'CP068274.2',
'CP068273.2',
'CP068272.2',
'CP068271.2',
'CP068270.2',
'CP068269.2',
'CP068268.2',
'CP068267.2',
'CP068266.2',
'CP068265.2',
'CP068264.2',
'CP068263.2',
'CP068262.2',
'CP068261.2',
'CP068260.2',
'CP068259.2',
'CP068258.2',
'CP068257.2',
'CP068256.2',
'CP068255.2',
'CP086569.2']
chroms = ['chr{}'.format(j) for j in list(range(1,23))+['X','Y']]
chrom_map = {ch:cp for cp,ch in zip(chroms,CP_chroms)}

# %% [code]
converted_chr = {}
converted_pos = {}

for i,a in enumerate(t2ts):
    if len(a)==0:
        continue
    if a[0][0] in chrom_map:
        converted_chr[i] = chrom_map[a[0][0]]
        converted_pos[i] = a[0][1]
converted_chr = pd.Series(converted_chr)
converted_pos = pd.Series(converted_pos)

# %% [code]
t2t_added = pd.concat([eqtls, pd.DataFrame({'t2t_chr':converted_chr}), pd.DataFrame({'t2t_pos':converted_pos})],axis = 1).dropna()

# %% [code]
common_genes = list(set(gene_info['ensemblid']).intersection(t2t_added['GENE'].values))

# %% [code]
t2t_added = t2t_added[t2t_added['GENE'].isin(common_genes)]

# %% [code]
t2t_added['gene_symbol'] = gene_info.set_index('ensemblid')['gene_symbol'].loc[t2t_added['GENE']].values

# %% [code]
t2t_added['t2t_ind'] = t2t_added['t2t_pos'].astype(int)//1000

# %% [code]
max_pips = {}
for tissue, gene, t2t_chr,t2t_ind,pip in zip(t2t_added['TISSUE'],t2t_added['gene_symbol'],t2t_added['t2t_chr'],t2t_added['t2t_ind'], t2t_added['Probability']):
    max_pips.setdefault((tissue, gene, t2t_chr,t2t_ind), []).append(pip) 
    
max_pip = pd.Series({k: np.max(v) for k,v in max_pips.items()}).reset_index()
max_pip.columns = ['TISSUE','GENE','t2t_chr', 't2t_ind', 'PIP']
max_pip = max_pip[max_pip['t2t_chr']!='chrY']

valid_long = max_pip[max_pip['TISSUE'].isin(common_tissues)]
good_tg_pairs = (valid_long.groupby(['TISSUE','GENE'])['PIP'].min() <0.9)&(valid_long.groupby(['TISSUE','GENE'])['PIP'].max() >0.9)


gtg = valid_long.groupby(['TISSUE','GENE']).size().sort_values()>3
good_tissue_gene_pairs = set(good_tg_pairs[good_tg_pairs].index.values).intersection(gtg[gtg].index.values)

valid_long = valid_long.loc[[i for i,a,b in zip(valid_long.index.values, valid_long['TISSUE'], valid_long['GENE']) if (a,b) in good_tissue_gene_pairs]]
valid_long = valid_long[valid_long['GENE'].isin(sc.columns)]

# %% [code]


# %% [code]
common_

# %% [code]
print(valid_long['GENE'].unique())

# %% [code]
DEVICE = 0

# %% [code]
unique_pos = valid_long[['t2t_chr','t2t_ind']].drop_duplicates()
chrom_inds = unique_pos.groupby('t2t_chr').groups
cis_seq_batches = []
dat_inds =[]
for chrom,inds in chrom_inds.items():
    chm13_pos = unique_pos['t2t_ind'].loc[inds]
    chr_dat = np.load('../data/seq/GENA_2k_precomputed/{}_precomputed_features.npy'.format(chrom), mmap_mode = 'r')
    cis_seq_batches.append(chr_dat[chm13_pos.values]/10)
    dat_inds+= list(inds)

cis_seq_batch = torch.tensor(np.concatenate(cis_seq_batches)).float().to('cuda:{}'.format(DEVICE))

qtl_to_seq_map = {ch_ind:i for i, ch_ind in enumerate (zip(unique_pos['t2t_chr'].loc[dat_inds],unique_pos['t2t_ind'].loc[dat_inds]))}


ct_gene_groups = valid_long.groupby(['TISSUE','GENE']).groups
cis_seq_inds = {}
pips = {}
for key,inds in ct_gene_groups.items():
    CELLTYPE,GENE = key
    cis_seq_inds[CELLTYPE, GENE] = np.array([qtl_to_seq_map[a] for a in zip(valid_long['t2t_chr'].loc[inds],valid_long['t2t_ind'].loc[inds])])
    pips[CELLTYPE,GENE] = valid_long['PIP'].loc[inds]

# %% [code]
renamed = tissue_genex_means.loc[sorted(common_tissues)]/1000
renamed.to_pickle('../data/downstream/tabsap_renamed')
valid_long.to_pickle('../data/downstream/tabsap_valid_long')

# %% [code]
def prepare_sc_batch(ct, gene_to_add, amount = 0.0001):
    ct_mean = renamed.loc[ct]
    edited_mean = ct_mean.copy()
    edited_mean[gene_to_add]+=amount
    return torch.tensor(np.stack([ct_mean.values.copy(), edited_mean.values]),device = 'cuda:{}'.format(DEVICE),dtype = torch.float)

# %% [code]
path  = 'results/apr7/tabsap_meansyntax_hilr/repeat_0_epoch_24.pt'.format(pt)
softmax = True
bn = False
n_learn = 1
n_control = 40
dropout= 0

bias = False
multi_learner = nn.ModuleList([LatentPSALearner(sc = torch.from_numpy(renamed.values),dropout=0, softmax=True, genome = cis_seq_batch, n_control = n_control, bias = bias,batch_norm = False) for _ in range(n_learn)]).to('cuda:{}'.format(DEVICE))

checkpoint = torch.load(path)
sd = checkpoint['learner_state_dict']
multi_learner.load_state_dict(sd)
multi_learner.eval();
learner = multi_learner[0]
learner.eval();


# initialize maps that will be shuffled during permutations
orig_genes = valid_long['GENE'].unique()
shuf_genes = orig_genes.copy()
orig_cts = renamed.index.values
shuf_cts = orig_cts.copy()
shuf_idx = np.arange(len(cis_seq_batch))

# want to enable shuffling only genes that are expressed in the same set of cell types, hence this shuffle
ct_set_to_g = {}
for g, ct_set in {g: tuple(sorted(renamed[g][renamed[g]>1e-3].index.values)) for g in orig_genes}.items():
    ct_set_to_g.setdefault(ct_set, []).append(g)

orig_group_genes = [np.array(gs) for _,gs in ct_set_to_g.items()]
shuf_group_genes = [arr.copy() for arr in orig_group_genes]
print([len(a) for a in orig_group_genes])


# select which kind of permutation
perm_gene = True
perm_ct= False
perm_rs = False
group_perm_gene = False
group_perm_rs = False
rand_init = False

assert (group_perm_gene&perm_gene)==False
assert (group_perm_rs&perm_rs)==False


nperm = 1000
perm_stats = {}
PRINT_EVERY = min(nperm,20)
np.random.seed(45)

for perm in tqdm(range(1+nperm)):
    # first permutation is always test statistic (i.e., no shuffle)
    if perm>0:
        if group_perm_gene:
            for arr in shuf_group_genes:
                np.random.shuffle(arr) #only shuffle within groups
        if group_perm_rs:
            for arr in shuf_group_rs:
                np.random.shuffle(arr) #only shuffle within groups
        if perm_gene:
            np.random.shuffle(shuf_genes)

        if perm_ct:
            np.random.shuffle(orig_cts)

        if perm_rs:
            np.random.shuffle(shuf_idx)

        if rand_init:
            multi_learner = nn.ModuleList([LatentPSALearner(sc = torch.from_numpy(renamed.values),dropout=dropout, softmax=softmax, genome = cis_seq_batch, n_control = n_control, bias = False,batch_norm = bn) for _ in range(n_learn)]).to('cuda:{}'.format(DEVICE))
            multi_learner.eval();
            learner = multi_learner[0]
            learner.eval();


    # if perm==0, shuf_gene_map, shuf_idx, shuf_ct_map are all identity
    if group_perm_gene:
        shuf_gene_map = {i:j for i,j in zip(np.concatenate(orig_group_genes), np.concatenate(shuf_group_genes))}
    else:
        shuf_gene_map = {i:j for i,j in zip(orig_genes,shuf_genes)}


    if group_perm_rs:
        shuf_idx = np.concatenate(shuf_group_rs)
    shuf_ct_map = {i:j for i,j in zip(orig_cts,shuf_cts)}


    # compute AUCs for the discrepancy
    for CELLTYPE, GENE in ct_gene_groups.keys():
        sub_seq = cis_seq_batch[shuf_idx][cis_seq_inds[CELLTYPE,GENE]]


        shuf_ct = shuf_ct_map[CELLTYPE]
        shuf_gene = shuf_gene_map[GENE]
        sc_batch = prepare_sc_batch(shuf_ct, shuf_gene,amount=1e-4)

        Txs =  -learner.T_xs(seq_batch= sub_seq, sc_batch = sc_batch)
        y_pred = (Txs[1]-Txs[0]).detach().cpu().numpy()

        y_true = (pips[CELLTYPE,GENE]>0.9).astype(int).values
        perm_stats[perm,CELLTYPE,GENE] = roc_auc_score(y_true= y_true, y_score = y_pred)

#         y_true = -pips[CELLTYPE,GENE]
#         perm_stats[perm,CELLTYPE,GENE] = spearmanr(y_pred,y_true)[0]


    # occasionally print
    if (1+perm)%PRINT_EVERY==0:
        res = pd.Series(perm_stats).reset_index()
        res.columns = ['perm', 'ct', 'gene', 'auc']
        avgs = res.groupby('perm').apply(lambda w: w.groupby('gene')['auc'].mean()).mean(axis = 1)
        print('pval:', perm, (np.abs(0.5-avgs[0])>np.abs(0.5-avgs.iloc[1:])).mean(), 'test_stat:', avgs[0], 'perm mean:',avgs[1:].mean())


# %% [code]
pd.Series(ptres).sort_values()

# %% [code]
pd.Series(ptres).plot()

# %% [code]


# %% [code]


# %% [code]


# %% [code]


# %% [code]
```
