## Supplementary material for "Deriving genetic codes for molecular phenotypes from first principles": Code: README.md

# PSA_sub
The genome features were extracted from the GENA language model using seq_preprocessing/long_tokens.py.

The genome kNN was constructed using seq_preprocessing/build_genome_knn_gena.ipynb.

The phenotype data was preprocessed using the notebooks in phenotype_preprocessing/, which contains notebooks for metacell construction and constructing the datasets.

PSA was performed using train_psa.py, with the arguments in 'training_arguments.txt'.

Other analysis (including OT etc)  were performed using the notebooks Fig*.ipnyb.
