## Supplementary material for "Deriving genetic codes for molecular phenotypes from first principles": Code: build_genome_knn_GENA.ipynb

```
# %% [code]
import pandas as pd
import numpy as np

# %% [code]
CP_chroms = ['CP068277.2',
'CP068276.2',
'CP068275.2',
'CP068274.2',
'CP068273.2',
'CP068272.2',
'CP068271.2',
'CP068270.2',
'CP068269.2',
'CP068268.2',
'CP068267.2',
'CP068266.2',
'CP068265.2',
'CP068264.2',
'CP068263.2',
'CP068262.2',
'CP068261.2',
'CP068260.2',
'CP068259.2',
'CP068258.2',
'CP068257.2',
'CP068256.2',
'CP068255.2',
'CP086569.2']
chroms = ['chr{}'.format(j) for j in list(range(1,23))+['X','Y']]
chrom_map = {cp:ch for cp,ch in zip(chroms,CP_chroms)}

# %% [code]


# %% [code]
selected_chroms = ['chr{}'.format(j) for j in list(range(1,23))+['X']]
chrom_features = {}
for chrom in selected_chroms:
    chrom_features[chrom] = np.load('../data/../GENA_LM/processed_chrom_{}.npy'.format(chrom_map[chrom]))

# %% [code]
import osos.mkdir('../data/seq/GENA_2k_precomputed/')

# %% [code]
for k,v in chrom_features.items():
    np.save('../data/seq/GENA_2k_precomputed/{}_precomputed_features.npy'.format(k),v)

# %% [code]
chrom_features.keys()

# %% [code]
import numpy as np
import pandas as pd
import faiss
import torch

from tqdm.notebook import tqdm

# %% [code]
chrom_features['chr1'].shape

# %% [code]
chroms = list(range(1,23)) + ['X']
chrom_files =[ '../data/seq/GENA_2k_precomputed/chr{}_precomputed_features.npy'.format(chrom) for chrom in chroms]
genome = np.concatenate([np.load(chrom, mmap_mode='r') for chrom in chrom_files])
genome_tensor = torch.from_numpy(genome)

cpu_index = faiss.IndexFlatIP(768)
co = faiss.GpuMultipleClonerOptions()
co.shard = True


i = 0
N = len(genome_tensor)
bs = 10000
for i in tqdm(range(1+ N//bs), total = N//bs):
    seq_cuda = genome_tensor[bs*i: min(bs*(i+1), N) ].cuda()
    normalized_vectors = seq_cuda/torch.norm(seq_cuda, p =2, dim=1,keepdim=True)
    cpu_index.add(normalized_vectors.detach().cpu().numpy())
gpu_index = faiss.index_cpu_to_all_gpus(cpu_index, co)

knng = np.zeros((N, 50), dtype = np.int64)
for i in tqdm(range(1+ N//bs), total = N//bs):
    seq_cuda = genome_tensor[bs*i: min(bs*(i+1), N) ].cuda()
    normalized_vectors = seq_cuda/torch.norm(seq_cuda, p =2, dim=1,keepdim=True)
    gpu_index.search(normalized_vectors.detach().cpu().numpy(), 50, I = knng[bs*i: min(bs*(i+1), N)])
np.save('../data/seq/GENA_2k_precomputed/full_syntax_knn_graph', knng.T)

# %% [code]
np.save('../data/seq/GENA_2k_precomputed/concatenated_genome.npy', genome/10 )
genome_tensor

# %% [code]
genome = np.load('../data/seq/GENA_2k_precomputed/concatenated_genome.npy')

# %% [code]
np.save('../data/seq/GENA_2k_precomputed/concatenated_genome.npy', genome/5)
```
