## Supplementary material for "Deriving genetic codes for molecular phenotypes from first principles": Code: hg38_ccre.ipynb

```
# %% [code]
import pandas as pd
import torch
import numpy as np
from tqdm import tqdm


import numpy as np
from scipy import sparse


import seaborn as sns
from matplotlib import pyplot as plt

import pickle

from scipy.sparse import csr_matrix as s_csr_matrix
from scipy.sparse.csgraph import laplacian
from scipy.spatial.distance import cdist

import seaborn as sns

# %% [code]
# !wget https://hgdownload.soe.ucsc.edu/gbdb/hg38/encode3/ccre/encodeCcreCombined.bb
# !wget http://hgdownload.soe.ucsc.edu/admin/exe/linux.x86_64/bigBedToBed
# !chmod a+x bigBedToBed
# !./bigBedToBed encodeCcreCombined.bb encode.bed

# %% [code]
!head -n 10 encodeCcreCombined.bb

# %% [code]
ccre = pd.read_table('encode.bed',header = None)

# %% [code]
ccre['start_idx'] = np.floor(ccre[1]/2000).astype(int)
ccre['end_idx'] = np.ceil(ccre[2]/2000).astype(int)
ccre['idx_length'] = ccre['end_idx'] - ccre['start_idx']

# %% [code]
enh = ccre[ccre[10].isin(['pELS','dELS'])].sort_values([0,1])

# %% [code]


# %% [code]
import networkx as nx
g = nx.Graph()
for i,(chrom,start,stop,next_chrom,next_start,next_stop) in enumerate(zip(enh[0],enh[1],enh[2],enh[0].iloc[1:],enh[1].iloc[1:],enh[2].iloc[1:])):
    if chrom!=next_chrom:
        continue
    g.add_node(i)
    if (next_stop-start)<2000:
        g.add_edge(i,i+1)

# %% [code]
concoms = list(nx.connected_components(g))

# %% [code]
start_stops = []
for c in concoms:
    start_stops.append((enh[0].iloc[min(c)], enh[1].iloc[min(c)], enh[2].iloc[max(c)]))

# %% [code]
bin_widths = np.array([b-a for _,a,b in start_stops])

# %% [code]
enh_pos = pd.DataFrame(start_stops)
enh_pos['mean'] = (0.5*(enh_pos[1]+enh_pos[2])).astype(int)

# %% [code]
np.random.seed(123)
from pyfaidx import Fasta
seq = Fasta('../data/seq/hg38.fa')


# %% [code]
kmer_init = {}
for i in range(1,7):
    kmer_init[i] = {}

res = {}
for chrom,inds in enh_pos.groupby(0).groups.items():
    chrom_seq =  str(seq[chrom]).upper()

    for ind,pos in tqdm(zip(inds,enh_pos['mean'].loc[inds]),total= len(inds)):
        kmers = {}        
        s = chrom_seq[pos-500:pos+500]
        for j in range(len(s)):
            for k in range(1,7):
                if (j<1000-k):
                    w = kmers.setdefault(s[j: j+k],0) 
                    kmers[s[j: j+k]] = w+1
             
        res[chrom,ind] = pd.Series(kmers)

# %% [code]
df = pd.DataFrame(res).fillna(0)

# %% [code]
idx_to_concom = {}
for i,c in enumerate(concoms):
    for j in c:
        idx_to_concom[j] = i

# %% [code]
#missing one per chrom

# %% [code]
sub_enh = enh.iloc[[j for j in idx_to_concom]]
sub_enh['concom'] = [v for _,v in idx_to_concom.items()]

# %% [code]
df

# %% [code]
sub_enh.to_pickle('hg38_enhancer')
df.to_pickle('hg38_enh_kmer_counts')

# %% [code]


# %% [code]
df.shape

# %% [code]
(4*300000*5000)/1e9

# %% [code]
torch.manual_seed(123)
torch.cuda.set_device(3)
with torch.no_grad():
    dat = torch.from_numpy(df.values.T).float().cuda()
    dat -= dat.mean(axis = 0,keepdims = True)
    dat /= (1e-6 + torch.std(dat, axis = 0, keepdims = True))
    pcs = torch.pca_lowrank(dat, q = 4000)
    proj = dat @ pcs[2]
    

# %% [code]
!pwd

# %% [code]
enh['concom'].isna().sum()
```
