## Supplementary material for "Deriving genetic codes for molecular phenotypes from first principles": Code: hg38_cre_evs.ipynb

```
# %% [code]
import numpy as np
import torch
import pandas as pd

# %% [code]
df = pd.read_pickle('hg38_enh_kmer_counts')

# %% [code]
torch.manual_seed(123)
torch.cuda.set_device(0)
with torch.no_grad():
    dat = torch.from_numpy(df.values.T).float().cuda()
    dat -= dat.mean(axis = 0,keepdims = True)
    dat /= (1e-6 + torch.std(dat, axis = 0, keepdims = True))
    pcs = torch.pca_lowrank(dat, q = 4000)
    proj = dat @ pcs[2]
    

# %% [code]
genome_pcs = pd.DataFrame(proj[:,:3050].cpu().numpy(), index = df.columns)

# %% [code]
from matplotlib import pyplot as plt
plt.plot(torch.cumsum(pcs[1]/pcs[1].sum(),0).cpu().numpy())
plt.ylim(0.9,1)
plt.vlines(3100, 0,1, colors = 'black')

# %% [code]
genome_pcs.to_pickle('hg38_cre_pcs')

# %% [code]
import seaborn as sns
from scipy.spatial.distance import cdist
samp = genome_pcs.sample(200)
sns.clustermap(cdist(samp, samp,metric = 'cosine'))

# %% [code]
from sklearn.neighbors import kneighbors_graph
knng = kneighbors_graph(genome_pcs,n_neighbors= 10, include_self = False,mode = 'distance',metric = 'cosine')

# %% [code]
from scipy import sparse
sparse.save_npz('hg38_cre_knng', knng)

# %% [code]
from scipy.sparse import csr_matrix as s_csr_matrix
from scipy.sparse.csgraph import laplacian


from cupyx.scipy.sparse import csr_matrix as cp_csr_matrix
from cupy.cuda import Device
from cupyx.scipy.sparse.linalg import eigsh as gpu_eigsh
from scipy.sparse.linalg import eigsh as cpu_eigsh
from scipy import sparse

import numpy as np
import torch
import pandas as pd

from matplotlib import pyplot as plt

from scipy import sparse

# %% [code]
genome_graph = sparse.load_npz('hg38_cre_knng.npz')


# %% [code]
lap = laplacian(genome_graph,normed = False,symmetrized = True)
with Device(1):
    genome_lap = cp_csr_matrix(lap)
    genome_res_gpu = gpu_eigsh(genome_lap,k = 2000,return_eigenvectors=True,which = 'SA') #takes ~ 20 minutes


# %% [code]
hg38_cre_spec = genome_res_gpu[0].get()
hg38_cre_evs = genome_res_gpu[1].get()
np.save('hg38_cre_spec', hg38_cre_spec)
np.save('hg38_cre_evs', hg38_cre_evs)

# %% [code]
```
