## Supplementary material for "Deriving genetic codes for molecular phenotypes from first principles": Code: long_tokens.py

```
from pyfaidx import Fasta
from tqdm.auto import tqdm
from src.gena_lm.modeling_bert import BertForSequenceClassification
from transformers import AutoTokenizer
import numpy as np
import os
import torch

import argparse
parser = argparse.ArgumentParser()
parser.add_argument('--genome', type = str, required = True)
parser.add_argument('--out_path', type = str,required = True)

args = parser.parse_args()

tokenizer = AutoTokenizer.from_pretrained('AIRI-Institute/gena-lm-bert-base')
model = BertForSequenceClassification.from_pretrained('AIRI-Institute/gena-lm-bert-base')
model.cuda()
model.eval()


# GENOME = 'data/genomes/ncbi_dataset/data/GCA_009914755.4/GCA_009914755.4_T2T-CHM13v2.0_genomic.fna'
# OUT_PATH = ''
# GENOME = 'ncbi_dataset/data/GCF_000001635.27/GCF_000001635.27_GRCm39_genomic.fna'
# OUT_PATH = 'data/precomputed/MOUSE'

GENOME = args.genome
OUT_PATH = args.out_path


data = Fasta(GENOME, sequence_always_upper=True)
print(len(data))


with torch.no_grad():
    for chrom in tqdm(data.keys()):
        chrom_len = len(data[chrom])
        splits = range(0, chrom_len,1000)
        chrom_outputs = np.zeros((len(splits),768), dtype = np.float16)
        for j,idx in tqdm(enumerate(splits), total = len(splits)):
            seq = str(data[chrom][idx: min(chrom_len, idx+2000)])
            if seq.count('N')>10:
                continue
            chrom_outputs[j] = (model(torch.LongTensor([[0] + tokenizer(seq)['input_ids']]).cuda(), output_hidden_states  = True)['hidden_states'][-1][0][1:].mean(axis = 0).to(torch.float16).detach().cpu().numpy())
        np.save(os.path.join(OUT_PATH,'processed_chrom_{}.npy'.format(chrom)),chrom_outputs)
        
```
