## Supplementary material for "Deriving genetic codes for molecular phenotypes from first principles": Code: SupFig_comparing_evo.ipynb

```
# %% [code]
!wget http://hgdownload.soe.ucsc.edu/goldenPath/hg38/multiz470way/multiz470waySummary.bb

# %% [code]
!./bigBedToBed multiz470waySummary.bb multiz470.bed

# %% [code]
import pandas as pd

# %% [code]
enh_pos = pd.read_pickle('../enh_pos')

# %% [code]
enh_pos.columns = ['Chromosome','Start','End','mean_pos']

# %% [code]
import pyranges as pr

# %% [code]
multiz = pd.read_table('../multiz470.bed',header=None)
multiz.columns = ['Chromosome','Start','End','Species','Score',5,6]

maf = pr.PyRanges(multiz.iloc[:,:5])

# %% [code]
from scipy.spatial.distance import cdist

# %% [code]
import numpy as np
import seaborn as sns
from scipy.sparse import load_npz

# %% [code]
kmer_counts = pd.read_pickle('../hg38_enh_kmer_counts').T
evs =np.load('../hg38_cre_evs.npy')

# %% [code]
kmer_counts.shape

# %% [code]
enh_pos['Start'] = enh_pos['mean_pos']-1000
enh_pos['End'] = enh_pos['mean_pos']+1000

# %% [code]
enh_sample = enh_pos.sample(10000, random_state = 4).sort_index()
ev_feat = evs[enh_sample.index.values]
kmer_feat = kmer_counts.values[enh_sample.index.values]
species_int = pr.PyRanges(enh_sample.reset_index()).join(maf).as_df()

# %% [code]
info = pd.read_table('../multiz_info.txt',header=None)
ev = species_int.groupby(['index','Species'])['Score'].max().unstack().fillna(0)
cols = []
names = set()
for a in ev.columns:
    
    ids = info[5][info[4].str.contains(a)]
    if len(ids)>0:
        zz = ids.iloc[0]
        
        if zz not in names:
            cols.append(a)
            names.add(zz)
        
# ev = ev[names]
ev = ev[cols]

# %% [code]
min(ev.index ==enh_sample.index)

# %% [code]
shuf = np.arange(ev.shape[0])
np.random.shuffle(shuf)
splits = np.array_split(shuf, 50)

# %% [code]
len(split)

# %% [code]
from scipy.spatial.distance import pdist
from scipy.stats import spearmanr,pearsonr,mannwhitneyu,ttest_rel

# %% [code]
res = {}
for i,split in enumerate(splits):
    ed = pdist((ev.values[split]))
    r1 = pearsonr(pdist(ev_feat[split]),ed)[0]
    r2 = pearsonr(pdist(kmer_feat[split]),ed)[0]
    res[i] = {'spectral':r1,'kmer':r2}

# %% [code]
dat = pd.DataFrame(res).T

# %% [code]
from matplotlib import pyplot as plt

# %% [code]
plt.figure(figsize = (5,5))
sns.stripplot(data = dat.melt(ignore_index = False),x = 'variable',y = 'value', c = 'black')
plt.hlines([dat['spectral'].median(), dat['kmer'].median()], [-0.25,0.75],[0.25,1.25], colors = 'black')
plt.xlabel('metric')
plt.ylabel('correlation with evolutionary distance')

# %% [code]
from scipy.stats import ttest_ind

# %% [code]
mannwhitneyu(dat['spectral'],dat['kmer'])

# %% [code]


# %% [code]


# %% [code]


# %% [code]


# %% [code]
knn = load_npz('hg38_cre_knng.npz')

# %% [code]
neighbors = knn.nonzero()[1].reshape(-1,10)

# %% [code]
neighbors.shape

# %% [code]


# %% [code]
info[4].str.contains('HLailF').sum()

# %% [code]
ev.columns

# %% [code]
best_match

# %% [code]
np.random.seed(1245)
choice = np.random.choice(np.arange(neighbors.shape[0]),50, replace = False)
samp_df = enh_pos.iloc[neighbors[choice].flatten()].reset_index()
p1 = pr.PyRanges(samp_df)
ints = p1.join(maf)
best_match = ints.df.groupby(['index','Species'])['Score'].mean()
ev= best_match.unstack().fillna(0)


cols = []
names = set()
for a in ev.columns:
    
    ids = info[5][info[4].str.contains(a)]
    if len(ids)>0:
        zz = ids.iloc[0]
        
        if zz not in names:
            cols.append(a)
            names.add(zz)
        
# ev = ev[names]
ev = ev[cols]

# %% [code]
ev = ev[cols]

# %% [code]
for a insamp_df[~samp_df['index'].isin(ev.index)]

ev.loc[196764] = 0

# %% [code]
dists = pd.DataFrame(cdist(ev,ev, metric = 'cityblock'),index=ev.index, columns = ev.index)

# %% [code]
for i in range(50):
    dat= samp_df.iloc[(10*i): (10*(i+1))]
    dd = (dat['mean_pos'].values[:,None] - dat['mean_pos'].values[None,:])
    print(np.abs(dd[np.triu_indices_from(dd,1)].min()))

# %% [code]
from tqdm.notebook import tqdm

# %% [code]
sub_neigh = neighbors[choice]

i = 0
rr = []
sn = sub_neigh.copy()
for perm in tqdm(range(200)):
    if perm>0:
        s = sub_neigh.copy().flatten()
        np.random.shuffle(s)
        sn = s.reshape(sub_neigh.shape)
    
    out = {}
    for k in range(3,10):
        for i in range(sub_neigh.shape[0]):
            out[i,k] = dists.loc[(sn[i][:k], sn[i][:k])].values.max()
    rr.append(pd.Series(out).unstack().mean())

# %% [code]
w = pd.concat(rr,axis=1)

# %% [code]


# %% [code]
sns.stripplot(data = w.iloc[:,1:].melt(ignore_index=False).reset_index(),
              x ='index', y = 'value', c= 'grey', s = 2)

plt.hlines(w.iloc[:,0], np.arange(7)-0.25, np.arange(7)+0.25,colors = 'red')

# %% [code]
w.iloc[:,1:].min(axis=1)

# %% [code]

(w.values[:,[0]] >= w.values[:,[a for a in range(700) if a not in [0,200]]]).mean(axis = 1)

# %% [code]
ev.iloc[]

# %% [code]
feats = np.load('hg38_cre_evs.npy')`

# %% [code]
feats.shape

# %% [code]


# %% [code]
d_ev = cdist(ev>0,ev>0,metric = 'cosine')
rr ={}
for n in np.linspace(100,500,20,dtype = int):
    d_f = cdist(feats[ev.index.values,:n],feats[ev.index.values,:n])
    
    qf = d_f[np.triu_indices_from(d_f,1)]
    qev  = d_ev[np.triu_indices_from(d_ev,1)]
    
    rr[n] = spearmanr(qf,qev)[0]

pd.Series(rr).plot()

# %% [code]


# %% [code]
n = 2000
shuf = ev.index.values.copy()
nperm = 1000
perm_res = np.zeros(nperm+1)
for perm in tqdm(range(nperm+1)):
    if perm>0:
        np.random.shuffle(shuf)
    d_f = cdist(feats[shuf,:n],feats[shuf,:n])
    qf = d_f[np.triu_indices_from(d_f,1)]
    qev  = d_ev[np.triu_indices_from(d_ev,1)]
    perm_res[perm] = spearmanr(qf,qev)[0]

plt.hist(perm_res[1:])
plt.vlines(perm_res[0], 0, 200, colors = 'black')
plt.title('p = %.3f'%(1-(perm_res[0]>np.abs(perm_res[1:])).mean()))

# %% [code]
plt.hist(perm_res[1:])
plt.vlines(perm_res[0], 0, 200, colors = 'black')

# %% [code]
perm_res

# %% [code]
rk = {}
for t in np.linspace(0.0001,0.5,50):
    rk[t] = ((qev[(qf>t) &  (qf<(t+1e-2))]<0.02).mean())

rk = pd.Series(rk).iloc[1:]

# %% [code]
plt.scatter(qf, qev,s = 0.1)

# %% [code]
from scipy.stats import spearmanr

# %% [code]
sns.clustermap(df)

# %% [code]

plt.scatter(qf[sel],qev[sel], s = 0.01)

# %% [code]


# %% [code]
from matplotlib import pyplot as plt

# %% [code]
plt.scatter(d_f[:100][:,:100].flatten(),d_ev[:100][:,:100])

# %% [code]
cor_res_250 = pd.DataFrame(outs)

# %% [code]
cor_res_50 = pd.DataFrame(outs)

# %% [code]
cor_res_150 =  pd.DataFrame(outs)

# %% [code]
outs = []
for _ in tqdm(range(10)):
    print(outs)
    choice = np.random.choice(np.arange(enh_pos.shape[0]),150, replace = False)
    samp_df = enh_pos.iloc[choice.flatten()].reset_index()
    p1 = pr.PyRanges(samp_df)
    ints = p1.join(maf)
    best_match = ints.df.groupby(['index','Species'])['Score'].mean()
    ev= best_match.unstack().fillna(0)
    
    
    cols = []
    names = set()
    for a in ev.columns:
        
        ids = info[5][info[4].str.contains(a)]
        if len(ids)>0:
            zz = ids.iloc[0]
            
            if zz not in names:
                cols.append(a)
                names.add(zz)
            
    # ev = ev[names]
    ev = ev[cols]
    
    n = 300
    shuf = ev.index.values.copy()
    d_ev = cdist(ev>0,ev>0,metric = 'cosine')
    nperm = 500
    perm_res = np.zeros(nperm+1)
    for perm in range(nperm+1):
        if perm>0:
            np.random.shuffle(shuf)
        d_f = cdist(feats[shuf,:n],feats[shuf,:n])
        qf = d_f[np.triu_indices_from(d_f,1)]
        qev  = d_ev[np.triu_indices_from(d_ev,1)]
        perm_res[perm] = spearmanr(qf,qev)[0]
    
    plt.hist(perm_res[1:])
    plt.vlines(perm_res[0], 0, 200, colors = 'black')
    pv = (1-(perm_res[0]>np.abs(perm_res[1:])).mean())
    plt.title('p = %.3f'%pv)
    outs.append([perm_res[0],pv])

# %% [code]


# %% [code]


# %% [code]


# %% [code]


# %% [code]


# %% [code]
import itertools

# %% [code]
ev.index.max()

# %% [code]
ntf = np.load('nt_evo_50m_cre_feats.npy')

# %% [code]
ntf.shape

# %% [code]
np.random.seed(1245)
choice = np.random.choice(np.arange(ntf.shape[0]),100, replace = False)
samp_df = enh_pos.iloc[choice.flatten()].reset_index()
p1 = pr.PyRanges(samp_df)
ints = p1.join(maf)
best_match = ints.df.groupby(['index','Species'])['Score'].mean()
ev= best_match.unstack().fillna(0)


cols = []
names = set()
for a in ev.columns:
    
    ids = info[5][info[4].str.contains(a)]
    if len(ids)>0:
        zz = ids.iloc[0]
        
        if zz not in names:
            cols.append(a)
            names.add(zz)
        
# ev = ev[names]
ev = ev[cols]
d_ev = cdist(ev>0,ev>0,metric = 'cosine')

# %% [code]
prop = {}
d_ev = cdist(ev>0,ev>0,metric = 'hamming')
for n in range(50,2000,50):
    d_f = cdist(feats[ev.index.values,:n],feats[ev.index.values,:n])
    outs = np.zeros(5000)
    np.random.seed(123)
    s = 0

    while s < len(outs):
        i = np.random.randint(len(d_f))
        j =  np.random.randint(len(d_f))
        k = np.random.randint(len(d_f))
        if len(set([i,j,k]))<3:
            continue
    
        outs[s] = np.argmax([ d_ev[i,j], d_ev[i,k]])== np.argmax([d_f[i,j], d_f[i,k]])
        s+=1
    
    prop[n] = outs.mean()
pd.Series(prop).plot()
plt.title('laplacian features')

# %% [code]


d_f = cdist(ntf[ev.index.values],ntf[ev.index.values],metric = 'cityblock')
outs = np.zeros(20000)
np.random.seed(123)
s = 0

while s < len(outs):
    i = np.random.randint(500)
    j =  np.random.randint(500)
    k = np.random.randint(500)
    if len(set([i,j,k]))<3:
        continue

    outs[s] = np.argmax([ d_ev[i,j], d_ev[i,k]])== np.argmax([d_f[i,j], d_f[i,k]])
    s+=1
outs.mean()


# %% [code]


# %% [code]
list(itertools.combinations(range(500), 3))[:10]

# %% [code]
d_f = cdist(feats[ev.index.values,:n],feats[ev.index.values,:n])
qf = d_f[np.triu_indices_from(d_f,1)]
qev  = d_ev[np.triu_indices_from(d_ev,1)]
plt.scatter(qf,qev, s = 0.01)

# %% [code]
rk = {}
for t in np.linspace(0.0001,0.5,50):
    rk[t] = ((qev[(qf>t) &  (qf<(t+1e-2))]<0.02).mean())

rk = pd.Series(rk).iloc[1:]

# %% [code]
rk

# %% [code]


# %% [code]
res = best_match.unstack().fillna(0)

# %% [code]
res[('chr1',    102402)]

# %% [code]
res
```
