## Supplementary material for "Deriving genetic codes for molecular phenotypes from first principles": Code: train_psa.py

```
import os
import argparse
parser = argparse.ArgumentParser()
parser.add_argument('--device', type = int, required = True)
parser.add_argument('--out_dir', type = str,required = True)

parser.add_argument('--overwrite', type = bool, default = False)
parser.add_argument('--save_every', type = int, default = 20)
parser.add_argument('--num_repeat', type = int, default = 2)


parser.add_argument('--precomputed_genome_prefix', type = str, default = '../data/seq/GENA_2k_precomputed')
parser.add_argument('--sc_prefix',type = str, default = '../data/sc/graphs/pbmc')
parser.add_argument('--num_chrom', type = int, default = 23)
parser.add_argument('--manual_seed', type = int,default = 11230)

parser.add_argument('--num_component', type = int, required = True)
parser.add_argument('--seq_batch_size',type = int, default=20000)
parser.add_argument('--batch_norm', action = 'store_true')
parser.add_argument('--layer_norm', action = 'store_true')
parser.add_argument('--softmax', action = 'store_true')
parser.add_argument('--num_genome_neighbors',type = int, default = 20)
parser.add_argument('--sc_batch_size',type = int, default=100)
parser.add_argument('--gradient_accumulation_steps',type = int, default=1)
parser.add_argument('--dropout',type = float, default=0)


parser.add_argument('--sc_consistency_weight',type = float, default = 0.1)
parser.add_argument('--seq_consistency_weight',type = float, default = 0.1)
parser.add_argument('--seq_entropy_weight',type = float, default = 0)
parser.add_argument('--ortho_weight',type = float, default = 0)
parser.add_argument('--sc_target_weight',type = float, default = 0)
parser.add_argument('--prox_consistency_weight',type = float, default = 0)
parser.add_argument('--seq_target_weight', type = float, default = 0.0)

parser.add_argument('--weight_decay', type = float, default = 0)
parser.add_argument('--ignore_density', action = 'store_true')
parser.add_argument('--sc_W_init', type = str)
parser.add_argument('--seq_W_init', type = str)
parser.add_argument('--init_noise',type = float, default=0)


parser.add_argument('--learning_rate', type = float, default = 1e-3)


parser.add_argument('--num_epochs',type = int, default=50)
parser.add_argument('--num_cpu', type = int, default = 8)
parser.add_argument('--val_every', type = int, default=10)
parser.add_argument('--psa_type', type = str,default = 'template')
parser.add_argument('--pool_fun', type = str,default = 'mean')
parser.add_argument('--learner_type', type = str, default = 'latent')
parser.add_argument('--learner_bias', type = bool, default = False)
parser.add_argument('--n_control', type = int, default =   5)
parser.add_argument('--condition', type = str,default = 'NA')


args = parser.parse_args()

NUM_COMPONENT = args.num_component
POOL_FUN = args.pool_fun
EPOCHS = args.num_epochs
PATH = args.out_dir
GENOME_PREFIX = args.precomputed_genome_prefix
NUM_CHROM = args.num_chrom


SC_CONSISTENCY_WEIGHT = args.sc_consistency_weight
SC_TARGET_WEIGHT = args.sc_target_weight
SEQ_CONSISTENCY_WEIGHT = args.seq_consistency_weight
PROX_CONSISTENCY_WEIGHT = args.prox_consistency_weight
SEQ_TARGET_WEIGHT = args.seq_target_weight
SEQ_ENTROPY_WEIGHT = args.seq_entropy_weight
ORTHO_WEIGHT = args.ortho_weight
IGNORE_DENSITY = args.ignore_density

SAVE_EVERY = args.save_every
VAL_EVERY = args.val_every
LEARNING_RATE = args.learning_rate
WEIGHT_DECAY = args.weight_decay

bias = args.learner_bias

if args.manual_seed:
    SEED = args.manual_seed


try:
    os.mkdir(PATH)
    print('made dir {}'.format(PATH))
except:
    print("path '{}' already exists".format(PATH))
    if args.overwrite:
        print('overwriting...')
    else:
        print('overwrite is set to false, terminating')
        quit()

import json
from tqdm.auto import tqdm
import torch
import numpy as np
from models.utils import moving_average
from sklearn.metrics import roc_auc_score
import pandas as pd
import wandb


if args.psa_type == "laplacian":
    print("PSA type:", args.psa_type)
    from models.GraphPSA import GraphLaplacianPSA as PSA
elif args.psa_type == "template":
    print("PSA type:", args.psa_type)
    from models.GraphPSA import GraphTemplatePSA as PSA
else:
    print("invalid PSA type, quitting")
    quit()

if args.learner_type == "latent":
    print("Learner type: latent, num control: {}".format(args.n_control))
    from models.PSALearners import LatentPSALearner as Learner
    learner_kwargs = {'n_control':args.n_control}
else:
    print("invalid PSA type, quitting")
    quit()


#save config
with open(os.path.join(PATH, 'config.json'), 'w') as f:
    f.write(json.dumps(vars(args)))


learner_kwargs['bias'] = False
learner_kwargs['dropout'] = args.dropout
if args.batch_norm:
    learner_kwargs['batch_norm'] = True
if args.layer_norm:
    learner_kwargs['layer_norm'] = True
    print('layer norm active, disabling batch norm')
if args.softmax:
    learner_kwargs['softmax'] = True

print(learner_kwargs)
from models.GraphPSA import MultiComponentGraphPSA

print("using genome prefix: ", GENOME_PREFIX)
#load syntax graph
syntax_graph_numpy = np.array(np.load(os.path.join(GENOME_PREFIX,'full_syntax_knn_graph.npy'),mmap_mode = 'r')[:(1+args.num_genome_neighbors)])
syntax_graph = torch.from_numpy(syntax_graph_numpy)

print('loaded syntax graph...')
#load genome
genome = np.load(os.path.join(GENOME_PREFIX,'concatenated_genome.npy'))
genome_tensor = (torch.from_numpy(genome)/10)# remove this - might be weird.to('cuda:{}'.format(args.device))
print('loaded genome features')

# we have to initialize the seq loader to avoid sampling kmers at the edges of each chrom
# this is only relevant for the proximity loss :)!
chroms = list(range(1,NUM_CHROM)) + ['X']
chrom_files =[ os.path.join(GENOME_PREFIX,'chr{}_precomputed_features.npy'.format(chrom)) for chrom in chroms]
chr_lens = [np.load(chrom, mmap_mode='r').shape[0] for chrom in chrom_files]
sums = np.cumsum(chr_lens)
seq_loader = []
for start,end in zip(np.pad(sums+1,1)[:-1],-1+ np.pad(sums,1)[1:-1]):
    seq_loader.append(np.arange(start,end))
seq_loader = np.concatenate(seq_loader)


# load phenotype + knn graph
SC_PREFIX = args.sc_prefix

phenotype_graph = torch.from_numpy(np.load(SC_PREFIX+'_train_graph.npy'))
sc_loader = np.load(SC_PREFIX+'_train_inds.npy')

sc = torch.from_numpy(np.load(SC_PREFIX + '_train_features.npy')).float()
val_sc = torch.from_numpy(np.load(SC_PREFIX+'_val_features.npy')).float()

print('NOT fitting density!')
sc_target = torch.ones(sc.shape[0], dtype = torch.float)
val_target = torch.ones(val_sc.shape[0], dtype = torch.float)


print ('loaded phenotype and knn graph')

### insert evals here


learner_kwargs['sc'] = sc
learner_kwargs['genome'] = genome_tensor
torch.set_num_threads(args.num_cpu)

seq_batch_size = args.seq_batch_size
sc_batch_size = args.sc_batch_size


N_seqbatch = len(seq_loader)//seq_batch_size
N_scbatch = max(1,len(sc_loader)//sc_batch_size)
seq_splits = np.array_split(seq_loader,N_seqbatch)
sc_splits = np.array_split(sc_loader,N_scbatch)


for repeat in range(args.num_repeat):
    total_iter = 0
    j = 0
    run = wandb.init(PATH,reinit=True)
    wandb.config.update(args)
    wandb.config.update({'repeat':repeat})
    torch.manual_seed(SEED+repeat)


    
    psa = MultiComponentGraphPSA(n_components = NUM_COMPONENT, psa_type = PSA, syntax_graph = syntax_graph[:20],
                       genome_tensor = genome_tensor,
                        phenotype_graph = phenotype_graph,
                        phenotype_tensor = sc,
                       sc_target=sc_target,
                        val_sc = val_sc,
                        val_target = val_target,
                        learner = Learner,
                        pool_fun=POOL_FUN,**learner_kwargs)


    psa.to('cuda:{}'.format(args.device))


        
    if args.seq_W_init:
        vt = torch.load(args.seq_W_init,map_location = torch.device('cpu'))['vt'][:,:args.n_control].float().to(args.device)
        seq_init = True
    else:
        print('no seq pca initialization provided, quitting')
        quit()
    if seq_init: 
        for learner in psa.learner:
            W = list(learner.seq_k.parameters())[0] #requires this to exist!
            W.data = vt.T
            print("successfully initialized the seq weights with PCA!, saved in {}".format(args.seq_W_init))


    if args.sc_W_init:
        vt = torch.load(args.sc_W_init,map_location = torch.device('cpu'))['vt'][:,:args.n_control].float().to(args.device)
        sc_init = True
        print('warning, disabling sc pca init since init specified')
    else:
        print('no sc pca initialization provided, quitting')
        sc_init = False


    if sc_init: 
        for learner in psa.learner:
            W = list(learner.sc_q.parameters())[0] #requires this to exist!
            W.data = vt.T
            print("successfully initialized the sc weights with PCA!, saved in {}".format(args.sc_W_init))
            

        

    optimizer = torch.optim.AdamW(list(psa.learner.parameters()), lr = LEARNING_RATE,weight_decay = WEIGHT_DECAY)
    loss_weights = {'sc_con':SC_CONSISTENCY_WEIGHT, 'syn_con':SEQ_CONSISTENCY_WEIGHT, 'prox_con':PROX_CONSISTENCY_WEIGHT,
                     'sc_targ': SC_TARGET_WEIGHT, 'seq_targ': SEQ_TARGET_WEIGHT, 'seq_entropy':SEQ_ENTROPY_WEIGHT}
    wandb.watch(psa.learner, log ='all')


    np.random.seed(SEED+repeat)
    for epoch in tqdm(range(EPOCHS)):
        np.random.shuffle(seq_loader)
        
        for i in tqdm(range(N_seqbatch)):
            
            sc_inds = torch.from_numpy(sc_splits[j])
            seq_inds = torch.from_numpy(seq_splits[i])
            
            
            losses,_= psa.loss(sc_inds=sc_inds, seq_inds = seq_inds)

            loss =0
            for key,weight in loss_weights.items():
                loss+=weight*torch.stack(losses[key]).mean()

            loss += ORTHO_WEIGHT*losses['ortho_loss']
            
        
            
            loss.backward()
            total_iter += 1

            if (total_iter%args.gradient_accumulation_steps)==0:
                log_dict = {}
                for indiv_loss in loss_weights:
                    for l_idx in range(NUM_COMPONENT):
                        log_dict['{}/{}'.format(indiv_loss, l_idx)] = losses[indiv_loss][l_idx]

                log_dict['ortho_loss'] = losses['ortho_loss']
                wandb.log(log_dict, step = total_iter) 
                optimizer.step()
                optimizer.zero_grad()
            if N_scbatch>1:
                j+=1
            if j==N_scbatch:
                j=0
                np.random.shuffle(sc_loader)
                sc_splits = np.array_split(sc_loader,len(sc_loader)/sc_batch_size)

            if (1+total_iter)%VAL_EVERY==0:
                wandb.log({'val_targ_loss/{}'.format(l_idx): val for l_idx,val in enumerate(psa.validate(seq_inds))}, step = total_iter)

            


        if ((1+epoch)%SAVE_EVERY == 0):
            torch.save({'learner_state_dict': psa.learner.state_dict(),
            'optim_state_dict':optimizer.state_dict()}, os.path.join(PATH, 'repeat_{}_epoch_{}.pt'.format(repeat,(1+epoch))))
    run.finish()       
            
```
